# Complex scaffold remodeling in plant triterpene biosynthesis

**DOI:** 10.1101/2022.09.26.509531

**Authors:** Ricardo De La Peña, Hannah Hodgson, Jack C.T. Liu, Michael J. Stephenson, Azahara C. Martin, Charlotte Owen, Alex Harkess, Jim Leebens-Mack, Luis E. Jimenez, Anne Osbourn, Elizabeth S. Sattely

## Abstract

Triterpenes with complex scaffold modifications are widespread in plants yet little is known regarding biosynthesis. Limonoids are an exemplary family that includes the bitter taste in citrus (e.g., limonin) and the active constituents in neem oil, a widely used bioinsecticide (e.g., azadirachtin). Despite limonoid commercial value, a complete biosynthetic route has not been described. Here, we report the discovery of 22 enzymes that catalyze 12 unique reactions including the 4-carbon scission and furan installation that are a signature of the limonoid family and a pair of sterol isomerases previously unknown in plants. This gene set is then used for the total biosynthesis of kihadalactone A and azadirone in a heterologous plant. These results enable access to valuable limonoids and provide a template for discovery and reconstitution of triterpene biosynthetic pathways in plants that require multiple skeletal rearrangements and oxidations.

## Introduction

Among numerous complex triterpenes that are found in the plant kingdom, limonoids are particularly notable given their wide range of biological activities and structural diversity that stems from extensive scaffold modifications (*1, 2*). Produced by mainly two families in the Sapindales, Rutaceae (citrus) and Meliaceae (mahogany) (*3*), these molecules bear a signature furan and include over 2,800 unique structures (*4, 5*). Azadirachtin, a well-studied limonoid, exemplifies the substantial synthetic challenge for this group of molecules, with 16 stereocenters and 7 quaternary carbons. Notably, few synthetic routes to limonoids have been reported (*6*), (*7*), (*8*). More generally, complete biosynthetic pathways to triterpenes with extensive scaffold modifications have remained elusive. This lack of production routes limits utility and biological investigation of clinical candidates from this diverse compound class (*9*).

Around 90 limonoids have also been reported to have anti-insect activity (*2*), and several have also been found to target mammalian receptors and pathways (*4*). For example, azadirachtin (**Figure 1**), the main component of biopesticides derived from the neem tree (*Azadirachta indica*), is a potent antifeedant against >600 insect species (*9*). Perhaps related to antifeedant activity, Rutaceae limonoids such as nomilin, obacunone and limonin (**Figure 1**) that accumulate in *Citrus* species at high levels (*3*) are partially responsible for the “delayed bitterness” of citrus fruit juice, which causes serious economic losses for the citrus juice industry worldwide (*10*). In mammalian systems, several limonoids have shown inhibition of HIV-1 replication (*11*) and anti-inflammatory activity (*12*). Some limonoids of pharmaceutical interest have also been associated with specific mechanisms of action: gedunin (**Figure 1**) and nimbolide (Fig. S1) exert potent anti-cancer activity through Hsp90 inhibition (*13*) and RNF114 blockade (*14, 15*), respectively. Despite extensive interest in the biology and chemistry of complex plant triterpenes over the last half century, few complete biosynthetic pathways have been described. A notable exception is the disease resistance saponin from oat, avenacin A-1, whose pathway consists of 4 cytochrome P450 (CYP)-mediated scaffold modifications and 6 side-chain tailoring steps (*16*). Significant barriers to pathway reconstitution of complex triterpenes include a lack of knowledge of the structures of key intermediates, order of scaffold modification steps, instability of pathway precursors, and the challenge of identifying candidate genes for the anticipated >10 enzymatic transformations required to generate advanced intermediates. Limonoids are no exception; to date, only the first three enzymatic steps to the protolimonoid melianol (**1**) from the primary metabolite 2,3-oxidosqualene have been elucidated (**Figure 1**) (*17*).

**Figure 1.**
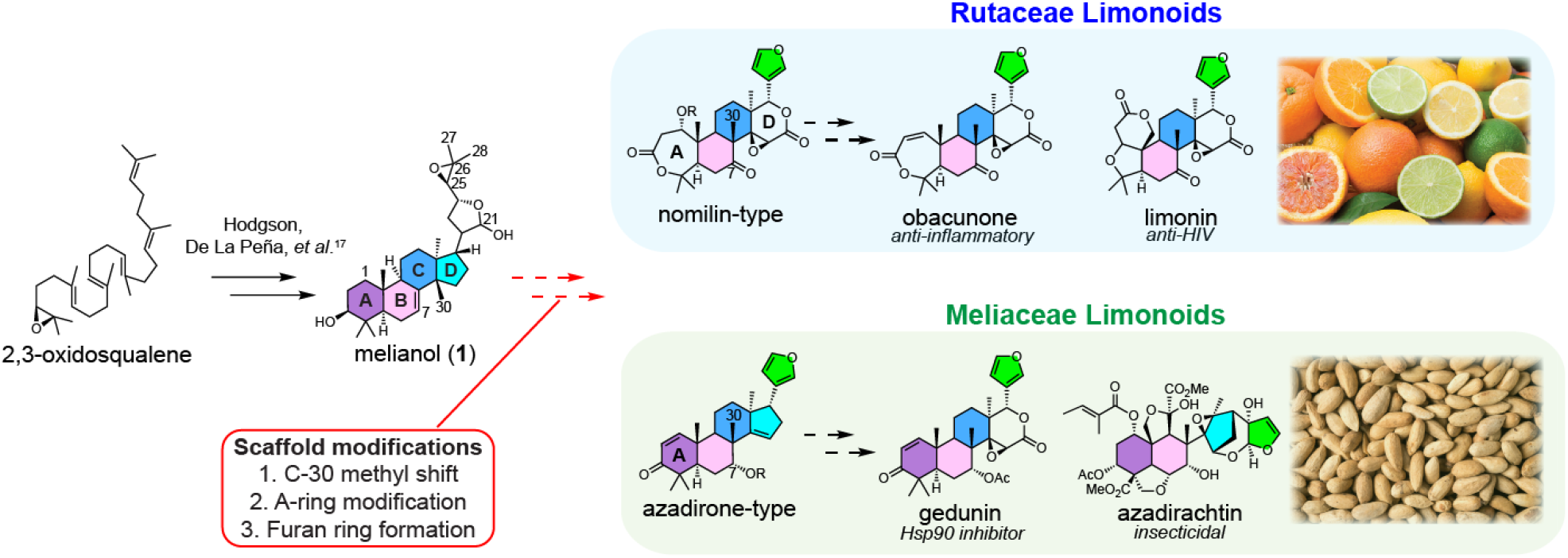
Structures of Rutaceae and Meliaceae limonoids and proposed biosynthetic pathway. We previously characterized three conserved enzymes from both *Citrus* and *Melia* species that catalyze the formation of the protolimonoid melianol (**1**) from 2,3-oxidosqualene (*17*). Additionally conserved scaffold modifications (outlined in the red box) are proposed to convert protolimonoids to true limonoids. Beyond this, Rutaceae limonoids differ from Meliaceae limonoids in two key structural features: *seco*-A,D ring and C-7 modification, which are proposed to be the result of Rutaceae and Meliaceae specific modifications. Exceptions to this rule could potentially arise from late-stage species-specific tailoring (Fig. S40). Rutaceae limonoids are derived from nomilin-type intermediates while Meliaceae limonoids are proposed to originate from azadirone-type intermediates. While the exact point of pathway divergence is unknown, comparative analysis of the various protolimonoid structures suggested that C-1, C-7, C-21 hydroxylation and/or acetoxylation are part of the conserved tailoring process. Obacunone and limonin are commonly found in various *Citrus* species (adapted photo by IgorDutina on iStock with standard license) and are responsible for the bitterness of their seeds. Azadirachtin (the most renowned Meliaceae limonoid) accumulates at high levels in the seeds of neem tree (photo by JIC photography), which are the source of commercial neem biopesticides.

In this work, we used systematic transcriptome and genome mining, coupled with co-expression, phylogenetic and comparative analysis, to identify suites of candidate genes from *Citrus sinensis* and *Melia azedarach* implicated in limonoid biosynthesis. We also generated a *de novo* annotated pseudochromosome-level genome assembly, with accompanying RNA-seq datasets for *M. azedarach* (a close relative of neem, *A. indica*) to facilitate our investigations. Functional enzyme characterization by transient expression and combinatorial biosynthesis in *Nicotiana benthamiana* – which we engineered to produce high levels of the precursor melianol (**1**) – resulted in the identification of a total of 22 biosynthetic genes involved in the biosynthesis of furan-containing limonoids, enabling – for the first time – the heterologous production of two furan-containing limonoids, azadirone (**18**) (Meliaceae) and kihadalactone A (**19**) (Rutaceae). The enzymes encoded by this gene set convert melianol to a limonoid with a core C26 scaffold (**Figure 1**) through a series of steps that include including ring rearrangement, C4 scission, furan installation, and key oxidative modifications common to both Citrus and Melia limonoids.

## Results

### Biosynthetic logic and comparative analysis of *Rutaceae* and *Meliaceae* metabolites

Limonoids are unusual within the triterpene class due to their extensive biosynthetic scaffold rearrangements. They are referred to as tetranortriterpenoids because their signature tetracyclic, triterpene scaffold (protolimonoid) loses four carbons during the formation of a signature furan ring to give rise to the basic C26 limonoid structure (**Figure 1**). A diversity of modifications can then occur to the basic limonoid scaffold through the cleavage of one or more of the four main rings post-cyclization (*18, 19*) (Fig. S1).

Radioactive isotope labeling studies suggest that most Rutaceae limonoids are derived from a nomilin-type intermediate (*seco*-A,D ring scaffolds) whereas Meliaceae limonoids are derived from an azadirone-type intermediate (intact ring scaffold) (**Figure 1**) (*4, 5*), (*20, 21*). It is proposed that at least two main scaffold modifications are conserved in both plant families: a C-30 methyl shift of the protolimonoid scaffold (*apo*-rearrangement) and the conversion of the hemiacetal ring of melianol (**1**) to a mature furan ring with a concomitant loss of the C-25∼C-28 carbon side chain (**Figure 1**) (*17*). Proposed Rutaceae- and Meliaceae-specific modifications would then yield the nomilin- and azadirone-type intermediates. While the exact point of divergence on the biosynthetic pathway between the two families is unknown, the diversity and array of protolimonoid structures isolated beyond melianol (**1**) hint at a series of possible conserved biosynthetic transformations. These include hydroxylation and/or acetoxylation among protolimonoids in C-1,C-7 and C-21 which suggests involvement of CYPs, 2-oxoglutarate-dependent dioxygenases (2-ODDs) and acetyltransferases.

### Identification of candidate limonoid biosynthetic genes

One genome of Rutaceae plants (*C. sinensis* var. Valencia) and several transcriptome resources, including from Citrus and Meliaceae plants (two from *A. indica* and one from *M. azedarach*) were previously used to identify the first three enzymes in the limonoid pathway (*17*). These included an oxidosqualene cyclase (*Cs*OSC1 from *C. sinensis, Ai*OSC1 from *A. indica*, and *Ma*OSC1 from *M. azedarach*), and two CYPs (*Cs*CYP71CD1/*Ma*CYP71CD2 and *Cs*CYP71BQ4/*Ma*CYP71BQ5) that complete the pathway to melianol (*17*). To identify enzymes that further tailor melianol (**1**), we expanded our search to include additional sources. For Rutaceae enzyme identification, we included publicly available microarray data compiled by the Network inference for Citrus Co-Expression (<u>NICCE</u>) (*22*). For Meliaceae enzyme identification, we generated additional RNA-seq data and a reference-quality genome assembly and annotation.

Of publicly available microarray data for Citrus, fruit datasets were selected for in depth analysis as *CsOSC1* expression levels were highest in the fruit and it has been implicated as the site of limonin biosynthesis and accumulation (*21*). Gene co-expression analysis was first performed on the Citrus fruit dataset using only *CsOSC1* as the bait gene. This revealed promising candidate genes exhibiting highly correlated expression with *CsOSC1* (Fig. S2). As we characterized more limonoid biosynthetic genes (as described below) we also included these as bait genes to enhance the stringency of co-expression analysis and further refine the candidate list. The top-ranking candidate list is rich in genes typically associated with secondary metabolism (**Figure 2A**). The list specifically included multiple predicted CYPs, 2-ODDs and acetyltransferases, consistent with the proposed biosynthetic transformations.

**Figure 2.**
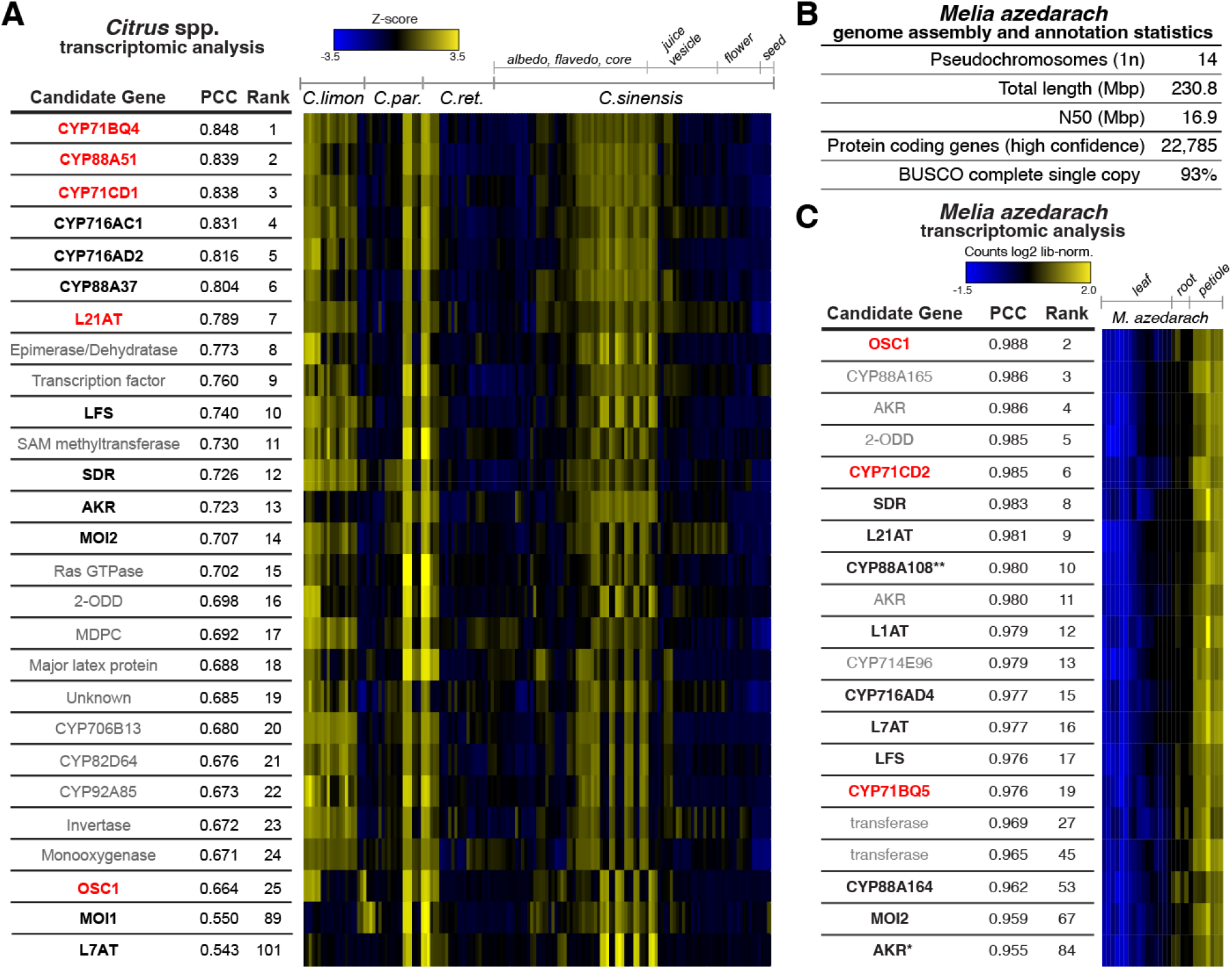
Genomic and transcriptomic analysis of *Citrus* and *Melia* resources. (A) Co-expression analysis of *C. sinensis* publicly available microarray expression data from <u>NICCE</u> (*22*) using *CsOSC1, CsCYP71CD1, CsCYP71BQ4, CsCYP88A51* and *CsL21AT* as bait genes. Linear regression analysis was used to rank the top 25 genes based on Pearson’s correlation coefficient (PCC) to the bait genes of interest. Heat map displays Z-score calculated from log2 normalized expression across the fruit dataset. The reported PCC value corresponds to the average value calculated using each bait gene. Genes in red indicate bait genes used in analysis and genes in black are functional limonoid biosynthetic genes (Table S17). Functional candidates outside of the top 25 genes are also included. For identification of individual bait genes used in this analysis see Fig. S2. (B) Summary of *Melia azedarach* pseudo-chromosome genome assembly and annotation statistics (Fig. S3 to S4, Table S1 to S2). (C) Expression pattern of *M. azedarach* limonoid candidate genes selected based on PCC to melianol biosynthetic genes (*MaOSC1, MaCYP71CD2* and *MaCYP71BQ5* (*17*), shown in red) and biosynthetic annotation. Heatmap (constructed using Heatmap3 V1.1.1 (*46*), with scaling by row (gene)) includes genes that are ranked within the top 87 for co-expression and are annotated with one of six interpro domains of biosynthetic interest (IPR005123 (Oxoglutarate/iron-dependent dioxygenase), IPR020471 (Aldo/keto reductase), IPR002347 (Short-chain dehydrogenase/reductase SDR), IPR001128 (Cytochrome P450), IPR003480 (Transferase) and IPR007905 (Emopamil-binding protein)). Asterisks indicate the following: (*) full-length gene identified in transcriptomic rather than genomic data via homology to *Cs*AKR ((Table S11, Table S18), (**) gene previously identified as homolog of limonoid co-expressed gene from *A. indica (17)*). Genes shown in black are newly identified functional limonoid biosynthetic genes (this study) (Table S10).

Efforts to identify and clone candidate genes from *M. azedarach* have previously been limited by the lack of a reference genome with high-quality gene annotations and by the lack of suitable transcriptomic data for co-expression analysis (i.e. multiple tissues, with replicates). Therefore, in parallel to our search in *Citrus*, we generated genomic and transcriptomic resources for *M. azedarach* to look for orthologous genes. A pseudochromosome level reference-quality *M. azedarach* genome assembly was generated using PacBio long-read and Hi-C sequencing technologies (Table S1, Fig. S3). Although the assembled genome size (230 Mbp) is smaller than available literature predictions for this species of 421 Mbp (*23*), the chromosome number (1n=14) matches literature reports (*23*) and was confirmed by karyotyping (Fig. S4). The genome assembly annotation predicted 22,785 high-confidence protein coding genes (**Figure 2B**, Table S1). BUSCO assessment (*24*) of this annotation confirmed the completeness of the genome, as 93% of expected orthologs are present as complete single copy genes (comparable to 98% in the gold standard *Arabidopsis thaliana*) (**Figure 2B**, Table S1).

Illumina paired-end RNA-seq reads were generated for three different *M. azedarach* tissues (7 different tissues in total, with four replicates of each tissue, Table S2), previously shown to differentially accumulate and express limonoids and their biosynthetic genes (*17*). Read-counts were generated by aligning RNA-Seq reads to the genome annotation, and EdgeR (*25*) was used to identify a subset of 18,151 differentially expressed genes (P-value < 0.05). The known melianol biosynthetic genes *MaOSC1, MaCYP71CD2* and *MaCYP71BQ5* (*17*) were used as bait genes for co-expression analysis across the sequenced tissues and the resulting ranked list was filtered by their Interpro domain annotations to enrich for relevant biosynthetic enzyme-coding genes. This informed the selection of 17 candidate genes for further investigation for functional analysis along with Citrus candidates (**Figure 2C**).

### Citrus CYP88A51 and Melia CYP88A108 act with different melianol oxide isomerases (MOIs) to form distinct proto-limonoid scaffolds

Top-ranking genes from both the *Citrus* and *Melia* candidate lists (**Figure 2A, 2C**) were tested for function by *Agrobacterium*-mediated transient expression in *N. benthamiana* with the previously reported melianol (**1**) biosynthetic enzymes *Cs*OSC1, *Cs*CYP71CD1, and *Cs*CYP71BQ4 or *Ai*OSC1, *Ma*CYP71CD2, and *Ma*CYP71BQ4. LC/MS analysis of crude methanolic extracts from *N. benthamiana* leaves revealed that the expression of either *Cs*CYP88A51 or *Ma*CYP88A108, in combination with their respective melianol biosynthesis genes, lead to the disappearance of melianol (**1**) and the accumulation of multiple mono-oxidized products (**Figure 3D**, Fig. S5 to S6). This result suggested that, while these CYP88A enzymes accept melianol as a substrate, the resulting products could be unstable or undergo further modification by endogenous *N. benthamiana* enzymes.

**Figure 3.**
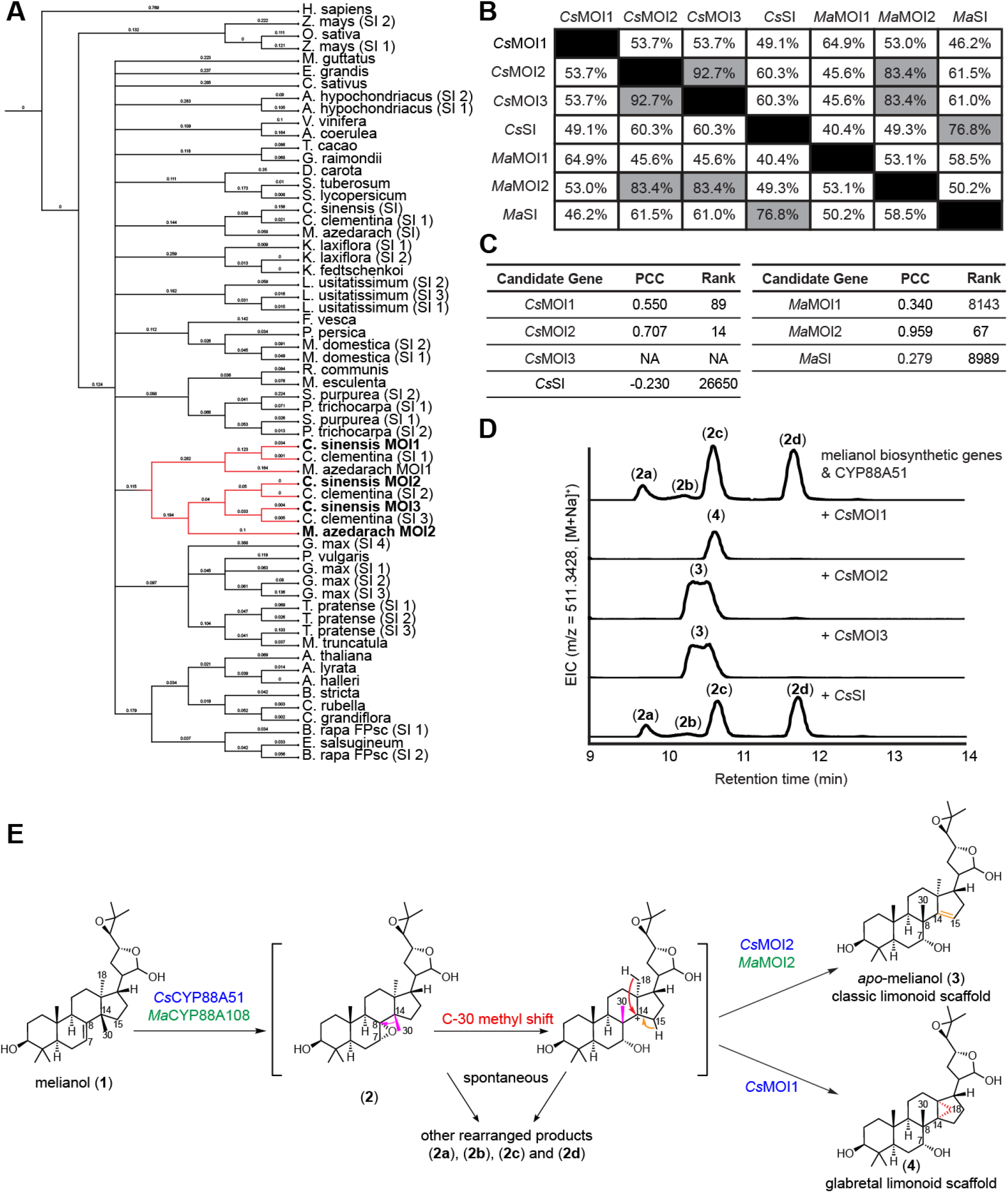
Characterization of MOIs. (A) Phylogenetic tree of SI genes from high-quality plant dicot genomes. SI sequences from ∼35 plant species were identified and downloaded from Phytozome via pFAM assignments. The human SI sequence was used as an outgroup. SI and MOIs from *C. sinensis* and *M. azedarach* selected for further analysis are bolded and their respective tree branches are indicated in red. The phylogenetic tree was built in Geneious with a neighbor-joining method and 100 bootstrap resamplings. Formatting of the tree was done using iTOL. (B) Percentage protein identity of MOIs and SIs from *C. sinensis* and *M. azedarach*, those with percent homology greater than 75% are highlighted in gray. (C) Co-expression of MOIs and SIs from *C. sinensis* and *M. azedarach* displaying rank and PCC as outlined in **Figure 2A, 2C**. (D) Characterization of products generated via overexpression of MOIs and SI using transient gene expression in *N. benthamiana*. Liquid chromatography–mass spectrometry (LC-MS) extracted ion chromatograms (EICs) resulting from overexpression of *At*HMGR, *Cs*OSC1, *Cs*CYP71CD1, *Cs*CYP71BQ4, *Cs*CYP88A51, and *Cs*MOIs and *Cs*SI in *N. benthamiana*. Representative EICs are shown (n=3). (E) Proposed mechanism of *Cs*CYP88A51/*Ma*CYP88A108,*Cs*MOI2/*Ma*MOI2 and *Cs*MOI1. *Cs*CYP88A51/*Ma*CYP88A108 first oxidizes the C7,C8 position of melianol (**1**) to yield an unstable epoxide intermediate (**2**), which undergoes spontaneous C-30 methyl shift from C-14 to C-8 (highlighted in red). Either (**2**) or the methyl shifted product spontaneously form a series of oxidized products (**2a** - **2d**). In the presence of MOIs, the rearrangement of (**2**) is guided to form either (**3**) or (**4**) and no (**2a**), (**2b**), (**2c**), and (**2d**) are observed. Note that structures of (**2**), (**2a**), (**2b**), (**2c**) and (**2d**) are not determined but their MS fragmentation patterns suggest they are isomeric molecules resulting from a single oxidation of melianol (**1**). Genes from *Citrus* are shown in blue and those from *Melia* are shown in green.

Despite the accumulation of multiple related metabolites, we continued to screen additional co-expressed candidate genes for further activity. This screen included homologs of *A. thaliana HYDRA1*, an ER membrane protein known as a sterol isomerase (SI) (two from the *Citrus* candidate list, and one from the *Melia* list). SIs are exclusively associated with phytosterol and cholesterol biosynthesis, where they catalyze double bond isomerization from the C-8 to the C-7 position. They are present in all domains of life and are required for normal development of mammals (*26*), plants (*27*) and yeast (*28*). Testing of these putative *SI*s through transient *Agrobacterium*-mediated gene expression in *N. benthamiana* resulted in a marked change of the metabolite profile with the accumulation of a single mono-oxidized product with no mass change (**Figure 3D**, Fig. S7). We suspected that these enzymes were able to capture unstable intermediates and promote isomerization of the C30 methyl group required to generate mature limonoids. These sterol isomerases are therefore re-named melianol oxide isomerases, *CsMOI1-3* and *MaMOI2*, because of their ability to generate isomers of mono-oxidized melianol products.

*SI*s are typically found as single copy genes in given plant species. Surprisingly, we found additional putative *SI* genes in the *C. sinensis* and *M. azedarach* genomes, four and three, respectively (Fig. S8). Phylogenetic analysis of SIs across a set of diverse plant species revealed that SIs are highly conserved and grouped similarly to plant phylogeny (**Figure 3A**). However, only one of the *C. sinensis* and one of the *M. azedarach* sequences (*CsSI* and *MaSI*) grouped with others of closely related plant species while the remaining *SI*s (*CsMOI1∼3* and *MaMOI1,2*) fell into a separate clade. This suggested that *CsSI* and *MaSI* are the conserved genes involved in phytosterol biosynthesis. Comparison of all *C. sinensis* and *M. azedarach* SI/MOI protein sequences showed that *Cs*MOI2 is ∼93% identical at the protein level to *Cs*MOI3 and ∼83% to *Ma*MOI2, but only ∼54% and ∼60% similar to *Cs*MOI1 and *Cs*SI, respectively (**Figure 3B**). While *CsMOI1, CsMOI2*, and *MaMOI2* ranked among the top 100 genes in our co-expression analysis lists (**Figure 3C**), *CsSI, MaMOI1* and *MaSI* showed no co-expression with limonoid biosynthetic genes. The absence of *CsMOI3* from this list is attributed to the lack of specific microarray probes required for expression monitoring. Notably, screening of *Cs*SI in the *N. benthamiana* expression system did not change the product profile of *Cs*CYP88A51, consistent with its predicted involvement in primary metabolism based on the phylogenetic analysis (**Figure 3D**).

To determine the chemical structures of the isomeric products formed through the action of these MOIs, we carried out large-scale expression experiments in *N. benthamiana* and isolated 13.1 mg of pure product. NMR analysis revealed the product of *Ma*MOI2 to be the epimeric mixture *apo-*melianol (**3**) bearing the characteristic limonoid scaffold with a migrated C-30 methyl group on C-8, a C-14/15 double bond, and C-7 hydroxylation (**Figure 3E**, Table S3) ((*29*). While the structure of the direct product of *Cs*MOI2 was not determined until after the discovery of two additional downstream tailoring enzymes, NMR analysis also confirmed C-8 methyl migration (Table S4). These data indicate that, as predicted by sequence analysis, *Cs*MOI2 and *Ma*MOI2 indeed are functional homologs and catalyze a key step in limonoid biosynthesis by promoting an unprecedented methyl shift. Analysis of the product formed with expression of *Cs*MOI1, indicated the presence of a metabolite with a different retention time relative to *apo-*melianol (**3**) (**Figure 3D**). Isolation and NMR analysis of (**4’**), a metabolite derived from (**4**) after inclusion of two additional tailoring enzymes (Table S5), indicated C-30 methyl group migration to C-8 and cyclopropane ring formation via bridging of the C18 methyl group to C-14.

Based on the characterized structures, we proposed that in the absence of MOIs, the CYP88A homologs form the unstable C-7/8 epoxide (**2**), which may either spontaneously undergo a Wagner-Meerwein rearrangement via C-30 methyl group migration and subsequent epoxide-ring-opening or degrade through other routes to yield multiple rearranged products (**2a**), (**2b**), (**2c**) and (**3**) (**Figure 3E**). MOIs stabilize the unstable carbocation intermediate and isomerize it to two types of limonoids: *Cs*MOI2, *Cs*MOI3 and *Ma*MOI2 form the C-14/15 double bond scaffold (classic limonoids) while *Cs*MOI1 forms the cyclopropane ring scaffold (glabretal limonoids). Glabretal limonoids have been isolated from certain Meliaceae and Rutaceae species before but are less common (*30, 31*). Together, our result suggest that *Cs*CYP88A51, *Ma*CYP88A108 and two different types of MOIs are responsible for rearrangement from melianol (**1**) to either (**3**) or (**4**) through an epoxide intermediate (**2**). These MOIs represent the first known cases of sterol isomerase neofunctionalization from primary metabolism in plants.

### Characterization of conserved tailoring enzymes L21AT and SDR

Having enzymes identified for the prerequisite methyl shift limonoid rearrangement, we continued screening other candidate genes (**Figure 2A, 2C**) for activity on (**3**) towards downstream products. BAHD-type acetyltransferases (named *Cs*L21AT or *Ma*L21AT, limonoid 21-*O*-acetyltransferse) and short-chain dehydrogenase reductases (*Cs*SDR and its homolog *Ma*SDR) were found to be active on (**3**), yielding acetylated and a dehydrogenated products, respectively (Fig. S9 to S12). While the sequence of events can be important for some enzymatic transformations in plant biosynthesis, L21AT and SDR homologs appear to have broad substrate specificity. L21AT can act on (**1**) or (**3**), and SDR is active on all intermediates after the OSC1 product (Fig. S13 to S14), suggesting a flexible reaction order in the early biosynthetic pathway.

Furthermore, the products formed from the modification of (**3**) by both Citrus and Melia L21AT and SDR *homologs* were purified by large-scale *N. benthamiana* expression and structurally determined by NMR to be 21(*S*)-acetoxyl-apo-melianone (**6**) (**Figure 4A**, Table S4, Table S6 to S7, Fig. S15). (**6**) is a protolimonoid previously purified from the Meliaceae species *Chisocheton paniculatus (32*) and is also detectable in *M. azedarach* tissues (Fig. S16). L21AT likely stereoselectively acetylates the 21-(*S*) isomer; a possible role for this transformation is stabilization of the hemiacetal ring observed as an epimeric mixture in melianol (**1**) (*17*) and *apo*-melianol (**3**) (Table S3). Overall, our results indicated that L21AT acetylates the C21 hydroxyl and SDR oxidizes the C3 hydroxyl to the ketone on early protolimonoid scaffolds.

**Figure 4.**
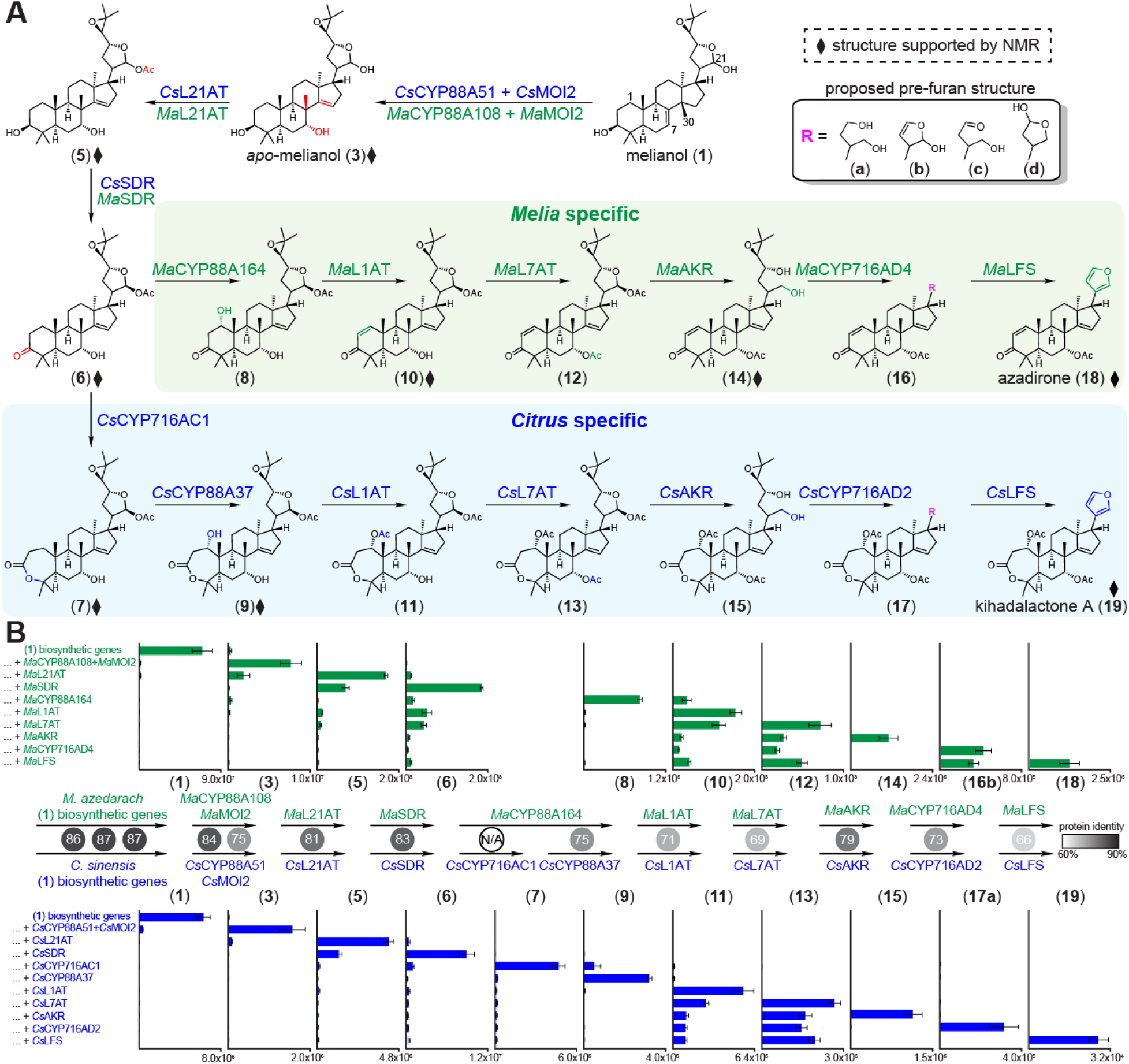
Complete biosynthetic pathway to azadirone (18) and kihadalactone A (19). (A) Gene sets that lead to the production of azadirone (18) and kihadalactone A (19) in *N. benthamiana* leaves. Arrows indicate the proposed sequence of biosynthetic steps based on metabolite data in Panel B. Diamonds represent intermediates whose structures were supported either by NMR analysis of the purified product or comparison with an authentic standard (**18**). Compounds **3, 5, 6, 9, 10**, and **14** were purified from *N. benthamiana* leaf extracts and analyzed by NMR; the structures of **7** and **19** are supported by partial NMR. (B) Integrated peak area of extracted ion chromatogram (EIC) for each pathway intermediates produced in *N. benthamiana* after sequential co-expression of individual enzymes. Values and error bars represent the mean and the standard error of the mean; n=6 biological replicates. Percentage identity between homologous proteins are shown in numbers in the circles and colored in gray scale. (**1**) biosynthetic genes comprise *Ma*OSC1/*Cs*OSC1, *Ma*CYP71CD2/*Cs*CYP71CD1, and *Ma*CYP71BQ5/*Cs*CYP71BQ4. *Cs*CYP88A37 is a homolog to *Ma*CYP88A164 while *Cs*CYP716AC1 has no *Melia* homolog.

### *Citrus* and *Melia* cytochrome P450s catalyze distinct limonoid A-ring modifications

Further *Citrus* and *Melia* candidate screens (**Figure 2A, 2C**) revealed activity of two *Citrus* CYPs, *Cs*CYP716AC1 and *Cs*CYP88A37, each capable of oxidizing (**6**) directly to (**7**) and (**8**) or consecutively to (**9**), and one CYP from *Melia* (*Ma*CYP88A164, a homolog of *Cs*CYP88A37) *a*lso capable of oxidizing (**6**) to (**8**) (**Figure 4A**, Fig. S17 to S20). Purification and NMR analysis of the downstream product (**9**) revealed it to be 1-hydroxy-luvungin A, which bears an A-ring lactone (Table S8). Additional NMR product characterization suggested that *Cs*CYP716AC1 is responsible for A-ring lactone formation and *Cs*CYP88A37 is responsible for C1 hydroxylation (Table S9). Although the exact order of oxidation steps to (**9**) appeared to be interchangeable for *Cs*CYP716AC1 and *Cs*CYP88A37, incomplete disappearance of (**6**) by *Cs*CYP88A37 suggested that oxidation by *Cs*CYP716AC1 takes precedence (Fig. S19).

Interestingly, in the absence of *Cs*SDR, neither *Cs*CYP716AC1 nor *Cs*CYP88A37 can oxidize the protolimonoid scaffold, suggesting the necessary involvement of the C-3 ketone for further processing (Fig. S21). These results, in combination with NMR characterization, indicated that *Cs*CYP716AC1 is likely responsible for Baeyer-Villiger oxidation to the A-ring lactone structure signature of Rutaceae limonoids. Comparative transcriptomics in *M. azedarach* revealed the lack of a *Cs*CYP716AC1 close homolog. The closest Melia homolog to *Cs*CYP716AC1 is truncated, not co-expressed with melianol biosynthetic genes, and only shares 63% protein identity (Table S10). These results highlight a branch point between biosynthetic routes in the Rutaceae and Meliaceae families.

### Acetylations complete tailoring in both *Citrus* and *Melia* protolimonoid scaffolds and set the stage for furan ring biosynthesis

Subsequent *Citrus* and *Melia* gene candidate screens (**Figure 2A, Figure 2C**) revealed further activity of BAHD acetyltransferases. *Cs*L1AT and its homolog *Ma*L1AT (named limonoid 1-*O*-acetyltransferase) are active on (**9**) (Fig. S22 to S23). When *Cs*L1AT was co-expressed with the biosynthetic genes for (**9**), a new molecule (**11**) with mass corresponding to acetylation of (**9**) was observed. When *Cs*CYP88A37 was omitted, acetylation of (**7**) was not observed (Fig. S24), suggesting that *Cs*L1AT acetylates the C-1 hydroxyl of (**9**) to yield (**11**). Surprisingly, when *Cs*CYP716AC1 was omitted from the Citrus candidates or when MaLAT1 was tested, the dehydration scaffold (**10**) accumulated (Fig. S23 to S24). Large-scale transient plant expression, purification, and NMR analysis of the dehydration product showed that the structure (**10**) (Table S11 to S12) contains a C-1/2 double bond and is an epimer to a previously reported molecule from *A. indica* (*33*). (**10**) also accumulated in *M. azedarach* extracts (Fig. S16). Two more co-expressed *Citrus* and *Melia* acetyltransferase homologs, *Cs*L7AT and *Ma*L7AT, (named limonoid 7-*O*-acetyltransferase) were found to result in acetylated scaffolds (**12**) and (**13**) and modification at the C-7 hydroxyl was later confirmed by the purification and NMR analysis of a downstream product (**14**) (**Figure 2A, Figure 2C**, Fig. S25 to S26, Table S13).

Three acetyltransferases (L1AT, L7AT, and L21AT) have been characterized in the biosynthesis for the 1,7,21-*O*-acetyl protolimonid (**13**) (**Figure 4A**). However, even in the presence of the biosynthetic genes to (**13**), two di-acetyl intermediates, (**11**) (1,21-*O*-acetyl) and possibly, the 1,7-*O*-acetyl protolimonoid scaffold, also accumulate (Fig. S27). This observation suggests promiscuity of L7AT and L21AT in terms of substrate specificity, and hints at the possibility that these transformations could represent signatures of a metabolic network.

### Downstream enzymes complete the biosynthesis to the furan-containing products azadirone (18) and kihadalactone A (19)

With acetylation established, the key enzymes involved in the C4 scission implicated in furan ring formation still remained elusive. It was unclear which enzyme classes could catalyze these modifications. We screened gene candidates via combinatorial transient expression in *N. benthamiana* as previously described and ultimately identified three active candidate pairs (one from each species): the aldo-keto reductases (*Cs*AKR/*Ma*AKR), the CYP716ADs (*Cs*CYP716AD2/*Ma*CYP716AD4), and the 2-ODDs (named limonoid furan synthase, *Cs*LFS/*Ma*LFS) (**Figure 2A,2C**). Systematic testing of these gene sets resulted in the accumulation of the furan-containing molecules azadirone (**18**) and khlidalactone A (**19**), two limonoids present in the respective native species. When *Cs*AKR/*Ma*AKR was included in our screens, we first identified the appearance of a new peak with mass corresponding to reductive deacetylation (Fig. S28 to S29). The product generated by expression of the *Melia* gene set in *N. benthamiana* was purified and characterized via NMR analysis to be the 21,23-diol (**14**) (**Figure 4A**, Table S13). Thus, the corresponding *Cs*AKR product (**15**) was proposed to share the same diol motif. It remains to be confirmed whether these AKRs perform C-21 deacetylation and reduction or whether these enzymes intercept a C-21,23 hemiacetal intermediate (that lacks the C-21 -OAc group) to form (**14**) or (**15**) diol product. While the latter is arguably the more likely scenario given the canonical activity of AKRs, it casts doubt on the role of L21AT in limonoid biosynthesis.

Unexpectedly, transient expression of *Ma*CYP716AD4 or *Cs*CYP716AD2 with the biosynthetic genes for (**14**) or (**15**) resulted in two new pairs of peaks, each with C4 loss. Proposed structures indicate a C_4_H_6_O fragment loss (**16a and 17a**) and a C_4_H_10_O fragment loss (**16b and 17b**) from their respective precursors (**Figure 4A**, Fig. S30 to S31). It is unclear whether these observed masses correspond to the true products of CYP716ADs or whether these are further modified by endogenous *N. benthamiana* enzymes. CYP716AD products may contain C-21 hydroxyl and C-23 aldehyde functionalities (**16c and 17c**) or spontaneously form the five-membered hemiacetal ring (**16d and 17d**) (**Figure 4A**, Fig. S32). We found that additional co-expression of LFS with the characterized genes to (**16 and 17**) resulted in the accumulation of products (**18**) and (**19**) (Fig. S33 to S34). Based on the predicted chemical formula, MS fragmentation pattern, and NMR analysis (Fig. S33, Table S14), we proposed the product of *Cs*LFS to be kihadalactone A (**19**), a known furan-containing limonoid (*34*) previously identified in extracts from the Rutaceae plant *Phellodendron amurense*. We detected the presence of (**19**) in *P. amurense* seed samples (Fig. S35), confirming prior reports of accumulation. Similarly, when *Ma*LFS was included in the co-expression, a new product with a mass equivalent to the furan-containing limonoid azadirone (**18**) was observed (Fig. S34). The production of azadirone (**18**) in *N. benthamiana* was confirmed by comparison to an analytical standard (Fig. S36) (isolated from *A. indica* leaf powder, with NMR confirmation (Table S15). In addition, we detected azadirone in extracts from three Meliaceae species (Fig. S36).

Taken together, we have discovered the 10- and 11-step biosynthetic transformations that enable the biosynthesis of two known limonoids, azadirone (**18**) and kihadalactone A (**19**), as well as an enzyme catalyzing the formation of the alternative glabretal scaffold (*Cs*MOI1). Sequential introduction of these enzymes into *N. benthamiana* transient co-expression experiments demonstrate step-wise transformations leading to (**18**) and (**19**) (**Figure 4B**). All of the enzymes involved in the biosynthesis of (**18**) and (**19**), except *Cs*CYP716AC1, are homologous pairs, and show a gradual decreasing trend in protein identity from 86% for the first enzyme pair *Cs*OSC1/*Ma*OSC1 to 66% for *Cs*LFS/*Ma*LFS. Intriguingly, despite the varied protein identities (**Figure 4B**), these homologous enzymes from Melia or Citrus can be used to created functional hybrid pathways comprising a mix of species genes, supporting a promiscuous evolutionary ancestor for the limonoid biosynthetic enzymes (Fig. S37).

## Discussion

In this work, we leverage parallel pathway discovery in two related producers of limonoids – Melia and Citrus – to help overcome many of the bottlenecks that have limited complex triterpene biosynthesis discovery to date. Comparative transcriptomics between these species combined with combinatorial testing of candidate gene sets in a heterologous host and extensive mass spectrometry and chemical structure analysis of intermediates were critical for the identification of a complete set of genes that can be used to produce two core limonoids: azadirone and kihadalactone A.

A major challenge in elucidating pathways that require many (e.g. >10) different metabolic steps is to determine whether observed enzymatic transformations in a heterologous host are “on-pathway” and, if so, in what order they occur. Combinatorial gene expression was particularly helpful in addressing this challenge. For example, we had not anticipated that C21 O-acylation of early intermediate (**3**) might enable the production of the limonoid furan. However, downstream analysis and enzyme drop-out experiments indicated that this transformation is required for production of kihadalactone A. In another example, early C7 *O*-acetylation seems unnecessary for the biosynthesis of elaborated limonoids, many of which have C7 ketone or OH functionality. However, when we omitted *Cs*L7AT in the full kihadalactone A (**19**) pathway, the expected (**19**) deacetylated product was not observed (Fig. S38). Instead, an oxidized intermediate accumulates that still contains the full triterpene scaffold, indicating that C-7 *O*-acetylation is important for C-4 scission. C-7 *O*-acetylation in the Meliaceae pathway is also key for furan biosynthesis, as in its absence an analogous side-product (**20**) is made (Fig. S31 to S32, Fig. S39, Table S16). These data suggest that downstream C-7 O-deacylation is required to reach further elaborated limonoids (e.g. limonin and azadirachtin; Fig. S40).

Another challenge is the potential that a pathway is not truly linear in the producing plant, but instead functions as a metabolic network. Although we track most closely major pathway intermediates that accumulate as we build a heterologous pathway, it is likely that limonoid biosynthesis in the native plants are highly branched and can result in the production of multiple variants. Some of this branching could also occur during pathway reconstitution. For example, MOI1 from Citrus catalyzes the formation of a cyclopropane product rather than a scaffold methyl shift; it is likely that this cyclopropane protolimonoid can also be processed in parallel by downstream enzymes to generate azadirone-like product variants. It is important to note that while all the pathway enzymes described in **Figure 4** are required for production of the final limonoid products, the sequence of enzymatic steps shown by the arrows is inferred from the accumulation of presumed pathway intermediates; a metabolic network that involves a different sequences of steps is also possible. Further study of each individual enzyme *in vitro* will be required to quantify substrate preference.

The 22 enzymes we describe here comprise core pathways to *Citrus* and *Melia* limonoids (**Figure 4B**) that can serve as a foundation for the discovery of additional enzymes for natural and unnatural limonoids. Our core pathway includes chemical transformations that have not previously been reported in plant specialized metabolism. For example, MOI1 and MOI2, which appear to have evolved from sterol isomerases, are capable of catalyzing two different rearrangements despite their conserved active site residues (Fig. S41). These data suggest that plants with more than one copy of sterol isomerase-like gene and homology differences would be interesting subjects for future investigations, since this may suggest a possible role in complex triterpene biosynthesis (**Figure 3A**). The co-location of the limonoid biosynthetic gene *MaMOI2* with two other non-limonoid SI genes in the *M. azedarach* genome is consistent with the origin of *MaMOI2* by tandem duplication and neofunctionalization (Fig. S42); this genomic arrangement is also conserved in *Citrus* on chromosome 5. Recent data also supports the unique role of these enzymes in quassinoid biosynthesis (*35*).

The C-4 scission and furan ring installation are novel transformations that generate an important pharmacophore of the limonoids. Although likely the result of convergent evolution, furan rings are found in a variety of natural products (*36, 37*). Although furan-forming enzymes have been reported from other plants (*38, 39*), (*40*), the AKR, CYP716AD and 2-ODD module described here represents a new mechanism of furan formation via the oxidative cleavage of a C-4 moiety. Along with the sterol isomerases (MOIs), the AKR and 2-ODDs add to the growing pool of enzyme families (*41, 42*) associated with primary sterol metabolism that appear to have been recruited to plant secondary triterpene metabolism, possibly due to the structural similarities between sterols and tetracyclic triterpenes.

Given their structural differences, the Rutaceae and Meliaceae limonoids are likely formed from different biosynthetic intermediates: nomilin-type (7-membered A ring) and azadirone-type (6-membered A ring), respectively (**Figure 1**). We show that their biosynthetic pathways are largely conserved until the production of (**6**), at which point *Cs*CYP716AC1 yields a divergent pathway in citrus. Interestingly, the percentage identities of the homologous genes are higher in the upstream than downstream pathway genes (**Figure 4B**). The lack of a clear *Cs*CYP716AC1 homolog in the *Melia* and *Azadirachta* genome sequences prompts us to speculate that a functional copy of a CYP716AC may have been lost in a Meliaceae ancestor. However, the possibility of neofunctionalization of a *Cs*CYP716AC1 after the evolution of a core limonoid pathway in a Rutaceae ancestor cannot be ruled out.

Limonoids are only one of many families of triterpenes from plants with complex scaffold modifications. Other examples include the *Schisandra* nortriterpenes (*43*), quinonoids (*44*), quassinoids (*45*), and dichapetalins (*44*); each represent a large collection of structurally diverse terpenes that contain several members with potent demonstrated biological activity but no biosynthetic route. Despite the value of these complex plant triterpenes, individual molecular species are typically only available through multi-step chemical synthesis routes or isolation from producing plants, limiting drug development (*15*) and agricultural utility (*9*). Many are only easily accessible in unpurified extract form that contains multiple chemical constituents; for example, azadirachtin, one of the most potent limonoids, can only be obtained commercially as a component of neem oil. Our results demonstrate that pathways to triterpenes with complex scaffold modifications can be reconstituted in a plant host, and the gene sets we describe enable rapid production and isolation of naturally-occurring limonoids. We anticipate that bioproduction of limonoids will serve as an attractive method to generate clinical candidates for evaluation, and that stable engineering of the limonoid pathway could be a viable strategy for sustainable crop protection.

## Supporting information

Supplemental NMR Data

## Acknowledgements

We would like to acknowledge Jasmine Staples for performing the extractions of Meliaceae material sourced from Kew Gardens. Work undertaken in A.O’s laboratory was funded through a Biotechnology and Biological Sciences Research Council (BBSRC) Industrial Partnership Award (IPA) with Syngenta (BB/T015063/1). A.O’s lab is also supported by the John Innes Foundation and the BBSRC Institute Strategic Programme Grant ‘Molecules from Nature - Products and Pathways’ (BBS/E/J/00PR9790). Work undertaken in E.S.S’s laboratory was supported by the National Institutes of Health (NIH) U01 grant GM110699 and NIH R01 grant GM121527.

## Author contributions

R.D.L.P., H.H., A.O. and E.S.S. conceived of the project and, with assistance from J.C.T.L., designed the research. H.H.generated and analyzed *M. azedarach* genome and transcriptome data. R.D.L.P. analyzed *Citrus* gene expression data and selected candidate genes from Citrus and, with J.C.T.L., expressed and characterized biosynthetic genes and metabolic products. L.E.J. assisted with isolation of Citrus intermediates. H.H. analyzed the *Melia* sequence resources and selected, expressed and characterized *Melia* biosynthetic genes and metabolic products. J.C.T.L and M.S. performed NMR analysis on the Citrus and Melia products, respectively. J.L.M advised on *M. azedarach* genomics. A.H. performed chromatin cross-linking and DNA extraction on *M. azedarach* tissues for Hi-C analysis by Phase Genomics. A.C.M. performed karyotyping on *M. azedarach* roots. C.O. combined the pseudo-chromosome level genome assembly with the *M. azedarach* annotation. R.D.L.P, J.C.T.L., H.H., A.O. and E.S.S. analyzed the data and wrote the manuscript.

