## Supplemental Material for "Complex scaffold remodeling in plant triterpene biosynthesis"

**This PDF file includes:**

Materials and Method

Figs. S1 to S42

Tables S1 to S23

Captions for Data S1

**Other Supplementary Materials for this manuscript include the following:**

Data S1 - Full NMR spectral data for isolated compounds

**Materials and Methods**	**6**

Generation of *Melia azedarach* genome assembly, annotation and RNA-seq dataset	6

Transcriptome data mining and analysis of Citrus dataset	7

Mining of resources for gene expression analysis in *M. azedarach*	7

Cloning of candidate genes from *C. sinensis* and *M. azedarach*	8

Characterization of *C. sinensis* and *M. azedarach* candidate genes through transient co-expression in *Nicotiana benthamiana*	8

Extraction and analysis of limonoids and protolimonoids from Rutaceae species and *N. benthamiana* expressing candidate *C. sinensis* biosynthetic genes	9

Extraction and analysis of limonoids and protolimonoids from Meliaceae species and *N. benthamiana* expressing candidate Meliaceae biosynthetic genes	10

General considerations for the purification and characterisation of limonoid intermediates from *N. benthamiana* expressing Citrus biosynthetic genes	10

General considerations for the purification and characterization of limonoid intermediates from *N. benthamiana* expressing *M. azedarach* biosynthetic genes and *A. indica*	11

General considerations for NMR characterizations	11

Purification of apo-melianol (**3**) (via expression of *M. azedarach* genes)	12

Purification of (**6**) (via expression of *C. sinensis* genes)	12

Purification of (**4’**) (via expression of *C. sinensis* genes)	13

Purification of 21(*S*)-acetoxyl-apo-melianone (**6**) (via expression of *M. azedarach* genes)	13

Purification of (**9**) (via expression of *C. sinensis* genes)	13

Purification of epi-neemfruitin B (**10**) (via expression of *M. azedarach* genes)	13

Purification of the *Ma*L7AT and *Ma*AKR product (**14**) (via expression of *M. azedarach* genes)	14

Purification of kihadalactone A (**19**) (via expression of *C. sinensis* genes)	15

Purification of azadirone (**18**) (from *A. indica* leaf powder)	15

Purification of MaCYP716AD4 side-product (20) (via expression of M. azedarach genes)	15

**Supplementary Figures**	**17**

Fig. S1. Supplementary limonoid and protolimonoid structures.	17

Fig. S2. Co-expression analysis of *C. sinensis* publicly available microarray expression data from Network inference for Citrus Co-Expression (NICCE) using *Cs*OSC1 as a bait gene.	18

Fig. S3. Hi-C heat map post-scaffolding heatmap of *M. azedarach* genome.	19

Fig. S4. Karyotyping of *M. azedarach.*	20

Fig. S5. Characterization of *Cs*CYP88A51.	21

Fig. S6. Individual activity of *Ma*CYP88A108 and *Ma*MOI2.	22

Fig. S7. Characterisation of *Ma*CYP88A108 and *Ma*MOI2.	23

Fig. S8. Histogram of the number of sterol isomerase genes present in high-quality plant genomes.	24

Fig. S9. Characterization of *Cs*L21AT.	25

Fig. S10. Characterisation of *Ma*L21AT.	26

Fig. S11. Characterization of *Cs*SDR.	27

Fig. S12. Characterisation of *Ma*SDR.	28

Fig. S13. Substrate promiscuity of *Cs*L21AT and *Cs*SDR.	29

Fig. S14. *Ma*SDR and *M*aL21AT can also function on melianol scaffold	30

Fig. S15. 3D models of 21(*S*)-acetyloxy-apo-melianone (**6**) and 21(*R*)-acetyloxy-apo-melianone.	31

Fig. S16. Detection of 21-acetoxyl-apo-melianone (**6**) and *epi*-neemfruitin B (**10**) in *Melia azedarach* samples.	32

Fig. S17. Characterization of *Cs*CYP716AC1.	33

Fig. S18. Characterization of *Cs*CYP88A37.	34

Fig. S19. Oxidation of 21-acetoxy-apo-melianone (**6**) by either *Cs*CYP88A37 or *Cs*CYP716AC1.	35

Fig. S20. Characterisation of *Ma*CYP88A164.	36

Fig. S21. Oxidation by *Cs*CYP88A37 or *Cs*CYP716AC1 requires *Cs*SDR.	37

Fig. S22. Characterization of *Cs*L1AT.	38

Fig. S23. Characterisation of *Ma*L1AT.	39

Fig. S24. Characterization of *Cs*L1AT in the absence of *Cs*CYP716AC1 or *Cs*CYP88A37.	40

Fig. S25. Characterization of *Cs*L7AT.	41

Fig. S26. Characterisation of *Ma*L7AT.	42

Fig. S27. Accumulation of 1,21-diacetoxyl (**11**) and 1,7-diacetoxyl protolimonoid intermediates with the introduction of *C*sL1AT and *Cs*L7AT.	43

Fig. S28. Characterization of *Cs*AKR.	44

Fig. S29. Characterisation of *Ma*AKR.	45

Fig. S30. Characterization of *Cs*CYP716AD2.	46

Fig. S31. Characterization of *Ma*CYP716AD4.	47

Fig. S32. Hypothetical reaction scheme for the action of CYP716ADs and LFSs via Baeyer-Villiger type route.	48

Fig. S33. Characterization of *Cs*LFS.	49

Fig. S34. Characterisation of *Ma*LFS.	50

Fig. S35. Detection of kihadalactone A (**19**) but not azadirone (**18**) in Rutaceae and agro-infiltrated *N. benthamiana* extracts.	51

Fig. S36. Azadirone (**18**) in agro-infiltrated *N. benthamiana* and Meliaceae extracts.	52

Fig. S37. Compatibility of *C. sinensis* and *M. azedarach* pathways.	53

Fig. S38. *Cs*L7AT is required for furan formation.	54

Fig. S39. *Ma*CYP716AD4 side-product (formed in the absence of C7-acetoxyl group).	55

Fig. S40. Proposed limonoid biosynthetic pathway in Rutaceae and Meliaceae plants.	56

Fig. S41. Alignment indicating the conserved active site residues between human sterol isomerase, *Cs*MOI1 and *Cs*MOI2.	57

Fig. S42. Genomic location and expression patterns of sterol isomerases in *M. azedarach*.	58

**Supplementary Tables**	**59**

Table S1. Summary of *M. azedarach* genome assembly and annotation.	59

Table S2. Summary of paired end reads generated for *M. azedarach* RNA-seq.	61

Table S3. ^13^C & ^1^H δ assignments of apo-melianol (**3**) produced using heterologously expressed genes from *M. azedarach* (C-21 epimeric mixture)	62

Table S4. ^13^C & ^1^H δ assignments of (**6**) produced using heterologously expressed genes from *C. sinensis*.	63

Table S5. ^13^C & ^1^H δ assignments of (**4’**) produced using heterologously expressed genes from *C. sinensis.*	64

Table S6. ^13^C & ^1^H δ assignments of 21(*S*)-acetoxyl-apo-melianone (**6**) produced using heterologously expressed genes from *M. azedarach.*	65

Table S7. ^13^C δ comparison with the literature for 21(*S*)-acetoxyl-apo-melianone (**6**).	66

Table S8. ^13^C & 1H δ assignments of 1-hydroxyl luvungin A (**9**) produced using heterologously expressed genes from *C. sinensis*.	67

Table S9. ^13^C & ^1^H δ partial assignments of degraded luvungin A (**7**) produced using heterologously expressed genes from *C. sinensis.*	68

Table S10. Gene ID of active *Melia azedarach* limonoid biosynthetic genes in this study.	69

Table S11. ^13^C & ^1^H δ assignments of *epi*-neemfruitin B (**10**) produced using heterologously expressed genes from *M. azedarach.*	70

Table S12. ^13^C δ comparison with the literature for *epi*-neemfruitin B (**10**) to neemfruitin B.	71

Table S13. ^13^C & ^1^H δ assignments of AKR product (**14**) produced using heterologously expressed genes from *M. azedarach*.	72

Table S14. ^1^H δ assignments of the furan moiety for kihadalactone A (**19**) produced using heterologously expressed genes from *C. sinensis.*	73

Table S15. ^13^C δ comparison with literature values for azadirone (**18**)	74

Table S16. ^13^C & ^1^H δ assignments of *Ma*CYP716AD4 side-product (C24 epimeric mixture) (20) produced using heterologously expressed genes from *M. azedarach.*	75

Table S17. Gene ID/Accession numbers of active Citrus limonoid biosynthetic genes and other Citrus genes in this study.	76

Table S18. Full length CDS and peptide sequence of *Ma*AKR (transcriptome derived).	77

Table S19. List of primer pairs used to clone genes from *C. sinensis.*	78

Table S20. List of primer pairs used to clone genes from *M. azedarach*.	79

Table S21. Isolera™ Prime fractionation conditions for purification of products of heterologously expressed *M. azedara*ch enzymes.	81

Table S22. Full length cloned nucleotide sequence of *MaMOI2*	82

**Captions for Data S1**	**83**

Data S1. NMR spectra for all isolated compounds	83

**References**	**84**

### Materials and Methods

#### Generation of *Melia azedarach* genome assembly, annotation and RNA-seq dataset

Two *Melia azedarach* plants (individuals ’02’ and ’11), purchased in 2016 (Crûg Farm Plants) and maintained (as described (*17*)) in a John Innes Centre greenhouse, were utilized for all sequencing described. Raw reads and genome assembly have been submitted to NCBI under bio-project number XXXXXX.

HMW gDNA (average 58 Kbp in length) was extracted from *M. azedarach* leaves (individual ‘11’) using the modified CTAB protocol which includes the addition of proteinase K and RNase A (Qiagen) (*47*). From this, the Earlham Institute constructed a 20-30 Kbp PacBio shotgun library which was sequenced over 10 SMRT cells on a Sequel instrument. The resultant filtered subreads (over two million with an average length of 13 Kbp) were *de novo* assembled, utilizing the hierarchical genome assembly process 4 (HGAP-4, PacBio) tool to create a draft genome with a total length of 230 Mbp (550 contigs). The proximo Hi-C Plant Kit (Phase Genomics) was used for chromatin cross-linking and subsequent extraction of DNA from *M. azedarach* leaves (individual ‘11’), following this, Hi-C (*48*) was performed by Phase Genomics. The proximal tool was then used to generate a pseudo-chromosome level assembly based on chromatin interactions from the Hi-C analysis and the draft *M. azedarach* genome. A mis-assembly within the draft genome (contig 000011F) was identified during this process and subsequently split, which resulted in the generation of 14 pseudo-chromosomes in the final assembly. Karyotyping was performed on young *M. azedarach* root tips (individual ‘11’). The preparation of mitotic metaphase spreads was carried out as described previously (*49*). Chromosomes were counterstained with DAPI (1 μg/ml). Images were acquired using a Leica DM5500B microscope equipped with a Hamamatsu ORCA-FLASH4.0 camera and controlled by Leica LAS X software v2.0.

Seven different tissues (n=4) were harvested for RNA extraction from *Melia azedarach* plants; upper leaves, lower leaves, petiole (including rachis) and roots of a high salannin-producing individual ‘11’ and upper leaves, lower leaves and petiole (including rachis) of a low salannin individual ’02’. Tissues were immediately flash frozen in liquid nitrogen before being ground to a fine powder using a pre-cooled pestle and mortar. All tissues were harvested on the same day and extractions were performed in technical replicates. RNA extraction was performed using the MacKenzie-modified RNeasy Plant Mini Kit (Qiagen) protocol (*50*), with DNAase (Promega) treatment, performed on column. The Earlham Institute generated high-throughput Illumina stranded RNA libraries (150bp, paired end) of each of the 28 samples, which were multiplexed and sequenced over two lanes of a HiSeq 4000 instrument (Illumina). This generated over 635 million paired end reads (an average of 91 million per tissue (Table S2)).

This RNA-seq dataset was utilized to by the Earlham Institute to generate a high quality structural genome annotation for *M. azedarach*, using their specialist plant genome annotation pipeline (including both Mikado (*51*) and Portcullis (*52*) tools), shown to be capable of annotating a diverse range of plant species (*53*, *54*). Functional annotation was generated using the Assignment of Human Readable Descriptions (*55*) (AHRD) V.3.3.3 tool. AHRD was provided with results of BLAST V2.6.0 (*56*) searches (e-value = 1e-5) against reference proteins from TAIR (*57*), UniProt (*58*), Swiss-Prot and TREMBL (*59*) datasets, along with interproscan (*60*) results.

#### Transcriptome data mining and analysis of *Citrus* dataset

Publicly available gene expression data from a collection of 297 Citrus datasets were downloaded from the Network Inference for Citrus Co-Expression (NICCE) (*22*). The dataset consisted of normalized expression data collected from multiple sources, tissues, and treatments (multiple Citrus spp., fruit, leaf, biotic stress, abiotic stress and age). Linear regression analysis to calculate Pearson’s R coefficient on normalized expression levels was performed using *CsOSC1* as the bait gene (Fig. S2). As additional genes were characterized, these were then used as bait genes along with previously characterized genes (*17*). These included using *CsCYP71CD1*, *CsCYP71BQ4*, *CsCYP88A51*, and *CsL21AT* as bait genes. The obtained list was ranked by decreasing Pearson’s R coefficient (PCC). The top microarray probes (per bait gene list) were then mapped to the respective *Citrus sinensis* genes. Candidate genes were then annotated both via Pfam assignment and via the best blastx hit using the *Arabidopsis thaliana* proteome as a reference. The final list of candidates was further refined as needed to only include candidates with Pfam assignments belonging to desired biosynthetic genes.

#### Mining of resources for gene expression analysis in *M. azedarach*

To process raw RNA-seq reads generated for *M. azedarach* and generate read counts, STAR V.2.5 (*61*) was used to align all reads to the *M. azedarach* genome annotation (pooling all reads per replicate (directional and lane)) and Samtools V.1.7 (*62*) was used to index the subsequent alignment. The featureCounts tool of subread V1.6.0 (*63*) was used to generate raw read counts by counting the number of reads overlapping with genes in each alignment.

Raw read counts were analyzed in R using DEseq2 v1.22.1 (*64*). Genes with zero counts were removed from the analysis, normalization was performed based on library size (to account for differences in number of reads sequenced for each replicate (*65*, *66*)) and subsequent counts were log_2_ transformed with a pseudo count of one. The resultant library-normalized log_2_ read counts were used for downstream analyses. Separately, differential expression analysis (to identify a subset of genes considered differentially rather than constitutively expressed) was performed by importing the raw read counts into an EdgeR (*25*) object and removing genes with low coverage (less than one count per million in more than four samples). Normalization (by library size) was performed using the ‘trimmed mean of M-values’ method. Finally to identify differentially expressed genes, a genewise negative binomial generalized linear model (glmQLFit) was used with pairwise comparisons between all sample types.

Using these differentially expressed genes as a subset, log_2_ library-normalized counts (generated by DEseq2 V1.22.1 (*64*)) for the 28 replicates were used to calculate Pearson’s correlation coefficients (PCCs) for each of the known melianol biosynthetic genes *MaOSC1, MaCYP71CD2* and *MaCYP71BQ5*. Genes were ranked based on their average PCC value against these three genes and then filtered to select only the genes with one of the following interpro annotations of biosynthetic interest; IPR005123 (Oxoglutarate/iron-dependent dioxygenase), IPR020471 (Aldo/keto reductase), IPR002347 (Short-chain dehydrogenase/reductase SDR), IPR001128 (Cytochrome P450), IPR003480 (Transferase) and IPR007905 (Emopamil-binding protein).

Although at rank 84 in this analysis (**Figure 2C**), *MaAKR* can be considered co-expressed, it is not as strongly co-expressed as other functional genes, and was in fact first identified due to its close homology to the functional Citrus gene *CsAKR*. Further, the gene prediction for *MaAKR* in the *M. azedarach* genome is truncated (lacking 38 terminal amino acids due to two point mutations). To identify a full-length homolog of *CsAKR* in *M. azedarach, de novo* transcriptome assembly was performed using Trinity v2.4.0 (*67*) following a standard protocol (*68*) and incorporating all petiole replicates from *M. azedarach* (individual ‘11’ (pooled)). Transdecoder v5.5.0 (*68*) was used to generate structural annotations for this transcriptome. Subsequently the truncated *MaAKR* (Table S10) sequence identified in the genome was used as a BLASTp query to identify the full length *MaAKR* sequence (Table S18).

#### Cloning of candidate genes from *C. sinensis* and *M. azedarach*

mRNA from *Citrus sinensis* var. Valencia (Sweet orange) fruit buds (green immature fruit 1~3 cm in diameter) from one-year old plants were isolated using the Spectrum Plant Total RNA Kit (Sigma-Aldrich) following the manufacturer’s instructions. Tissues were flash-frozen in liquid nitrogen and ground using a pestle and mortar. cDNA was generated using Super Script IV First Strand Synthesis System (Invitrogen). Candidate genes from *C. sinensis* were cloned (via Gibson assembly) into pEAQ-HT vectors (*69*), and transferred into *Agrobacterium tumerificans* (strain *GV3101*) following methods which have been previously described (*70*). Candidate genes from *M. azedarach* were amplified from leaf and petiole cDNA, cloned (via gateway cloning) into pEAQ-HT-DEST1 vectors (*69*) and transferred into *Agrobacterium tumerificans* (strain *LBA4404*) following methods which have been previously described (*17*). Primers used for cloning of functional genes from *C. sinensis* and *M. azedarach* are listed in Table S19 and Table S20, respectively.

#### Characterization of *C. sinensis* and *M. azedarach* candidate genes through transient co-expression in N*icotiana benthamiana*

To understand the function of enzymes of interest, candidate genes from *C. sinensis* and *M. azedarach* were tested via co-expressing various combinations of candidate genes with the previously characterized melianol biosynthetic genes (*AiOSC1/CsOSC1, MaCYP71CD2/CsCYP71CD1* and *MaCYP71BQ5/CsCYP71BQ4* (*17*)). This was performed by agroinfiltration of *A. tumefaciens* strains harboring the genes of interest in pEAQ vectors, following methods previously described (*17*, *71*). In addition to the limonoid biosynthetic genes, *Avena strigosa tHMGR* (encoding a truncated feedback insensitive HMG CoA-reductase that boosts triterpene yield (*71*)) was infiltrated in combination with *M. azedarach* candidate genes, while *A. thaliana* HMG CoA-reductase was used in combination with *C. sinensis* candidate genes.

#### Extraction and analysis of limonoids and protolimonoids from Rutaceae species and *N. benthamiana* expressing candidate *C. sinensis* biosynthetic genes

*N. benthamiana* leaf tissue was collected 5-days post *Agrobacterium* infiltration using a 1 cm DIA leaf disc cutter. Each biological replicate consisted of 4 leaf discs from the same leaf (approx. 0.04 g FW leaves). Citrus plant materials or leaf discs were lyophilized overnight and placed inside a 2 mL safe-lock microcentrifuge tube (Eppendorf). 500 μL of methanol (Fisher Scientific, ACS & HPLC grade) was added to each sample, and these were then homogenized in a ball mill (Retsch MM 400) using 5 mm stainless steel beads and milled at 25 Hz for 2 min. After homogenization, the samples were centrifuged at 13200 rpm for 10 min. Supernatants were filtered using either 0.20 or 0.45 μm PTFE filters (GE) before being subjected to LC-MS analysis.

LC-MS was carried using electrospray ionization (ESI) on positive mode on an Agilent 1260 HPLC coupled to an Agilent 6520 Q-TOF mass spectrometer. Separation was carried out using a 5 μm, 2 × 100 mm Gemini NX-C18 column (Phenomenex) using 0.1% formic acid in water (A) versus 0.1% formic acid in acetonitrile (B) run at 400 μL/min, room temperature. The following gradient of solvent B was used: 3% 0-1 min, 3%-30% 1-3 min, 30%-97% 30-18 min, 97% 18-22 min, 97%-3% 22-23 min and 3% 23-29 min. MS spectra was collected at m/z 50 -1400. The ESI source was set as follows: 350 °C gas temperature, 10 L/min drying gas, 35 psi nebulizer, 3500 V VCap, 150 V fragmentor 65 V skimmer and 750 V octupole 1 RF Vpp.

MS/MS data (100-1700 m/z, 1.5 spectra/sec) was collected using the same instrument, column and gradient under AutoMS2 acquisition mode, with a medium isolation width (~4 m/z) and fixed collision energies of 40 eV.

In addition, seeds of *Phellodendron amurense* (amur cork tree) were purchased from eBay and 2~3 seeds were homogenized in a ball mill (Retsch MM 400) using 5 mm stainless steel beads and milled at 25 Hz for 2 min in 2 mL ethyl acetate solvent. The extract were air dried, redissolved in equal volume of methanol, and filtered using 0.45 μm PTFE filters (GE) before subjecting to LC-MS analysis.

#### Extraction and analysis of limonoids and protolimonoids from Meliaceae species and *N. benthamiana* expressing candidate Meliaceae biosynthetic genes

For each sample, 10 mg of freeze-dried plant material was weighed and then homogenized using Tungsten Carbide Beads (3 mm; Qiagen) with a TissueLyser (1000 rpm, 2 min). Samples were agitated at 18 °C for 20 min in 500 µl methanol (100%). Samples were transferred to a 0.22 µM filter mini-column (Geneflow) and filtered by centrifugation before being transferred to a glass analysis vial.

Unless otherwise stated, all UHPLC-MS experiments described relating to Meliaceae material and genes were performed with positive mode electrospray ionization (Dual AJS ESI) on an LC/Q-TOF instrument (6546, Agilent), with separation by on an 1290 infinity LC system equipped with a DAD (Agilent). 1 ul of sample was injected for separation on a Kinetex 2.6 µm XB-C18 100 Å 2.1 x 50 mm column (Phenomenex) using 0.1% formic acid in water (A) versus acetonitrile (B) at 500µl/min and 40°C. Separation was performed using the following gradient of solvent B: 37% 0-1 min (first minute of flow diverted to waste), 37-67% 1-11 min, 67-100% 11-11.5 min, 100% 11.5-13.5 min, 100-37% 13.5-14 min and 37% 14-15 min. Full MS spectra were collected (m/z 100-1000, 1 spectra/sec). Spray chamber and source parameters were as follows; 325 °C gas temperature, 10 L/min drying gas, 20 psi nebulizer, 3500 V VCap, 120 V fragmentor 45 V skimmer and 750 V octupole 1 RF Vpp. Reference masses used for calibration were 121.05087300 and 922.00979800. In addition DAD spectra (200-400 nm, 2 nm step) were collected.

In addition to metabolite extraction from infiltrated *N. benthamiana* and the *Melia azedarach* trees maintained at JIC, extraction and analysis was also performed on dried leaf material from 13 Meliaceae species (*Carapa guianensis, Cipadessa fruticosa, Dysoxylum spectabile, Khaya nyasica, Malleastrum mandenense, Melia azedarach, Nymania capensis, Toona sinensis, Trichilia havanensis, Turraea floribunda, Turraea obtusifolia, Turraea sericea* and *Turraea vogelioides*) sourced from Kew Gardens in 2017 (Nagoya Protocol compliant) and stored at -70 °C.

#### General considerations for the purification and characterisation of limonoid intermediates from *N. benthamiana* expressing Citrus biosynthetic genes

Approximately 500 g of leaves from 60-100 infiltrated plants were cut into small pieces of approximately 0.25 cm^2^ in area. Leaves were immediately flash frozen and lyophilized to complete dryness. Dried leaves were then grinded to powder using a mortar and pestle. Leaf powder was then placed in a 4 L flask with a magnetic stir bar and extracted using EtOAc for 72 h at room temperature with constant stirring (1 g FW leaves per 12.5 mL). Extracts were filtered using vacuum filtration and dried using rotary evaporation. Flash chromatography was performed using a 7 cm DIA column loaded with silica (SiliaFlash® P60). Hexane (Fisher Scientific, ACS & HPLC grade) and ethyl acetate were used as running solvents. 500 mL fractions were collected via isocratic elution (60% hexane, 40% ethyl acetate). Fractions were analyzed via LC-MS, and those containing the compound of interest were pooled and dried using rotary evaporation. The dried sample was then resuspended in approximately 1 mL of DMSO. The samples were then further purified using an Isolera Prime Biotage using a Sfår C18 Duo 12g column. Fractions were collected using water (A) and acetonitrile (B) as solvents. The following gradient of solvent B was used: 30% for 3 column volumes (CV), 30 – 80% for 25 CV, 80 – 100% 2 CV. Active fractions, as verified by LC-MS, were then dried to completion using rotary evaporation or lyophilization. For Citrus intermediates ^1^H NMR and ^13^C NMR spectra were acquired using a Varian Inova 600 MHz spectrometer at room temperature. Shifts are referenced to the residual solvent peak (CDCl_3_, Acros Organics) and reported downfield in ppm using Me_4_Si as the 0.0 ppm internal reference standard.

#### General considerations for the purification and characterization of limonoid intermediates from *N. benthamiana* expressing *M. azedarach* biosynthetic genes and *A. indica*

To enable the purification of heterologously produced intermediates from *N. benthamiana*, large-scale vacuum infiltration of the relevant *A. tumefaciens* strains was performed as previously described (*72*, *73*), using 100-130 large-sized *N. benthamiana* plants. Once harvested and freeze-dried, a preliminary triterpene extraction was performed on the leaf material using a previously described method (*73*). Briefly, a speed extractor (Bucchi) was used to perform high temperature (100 °C) and pressure (130 bar) extraction from leaf material with ethyl acetate. Unless otherwise specified, the ambersep 900 hydroxide form beads (Sigma-Aldrich) recommended to remove chlorophylls (*73*) were not used, due to the presence of acetate groups in the compounds being isolated.

All Preparative HPLC was performed on an Agilent Technologies infinity system equipped with a 1290 infinity II fraction collector, a 1290 infinity II preparative pump and column oven, a 1260 infinity II quaternary pump, a 1260 infinity II Diode Array Detector (DAD), a 1260 infinity II ELSD and an infinity lab LC/MSD XT. Separation for preparative HPLC was performed on a 250 x 21.2 mm Luna® 5 µM C18(2) 100 Å column (Phenomonex), at 25 ml/min, with a collection:detector split of 1000:1 and the quaternary pump providing a make-up flow at 1.2 ml/min for the detectors. All preparative runs included a minimum of 3 minutes post-time at starting solvent percentage. Unless otherwise stated, MS data was collected via MM-ES+APCI scan mode, collecting data after 1.5 min with a mass range 200-1200 and collection of [M] or [M+H]^+^ masses.

#### General considerations for NMR characterizations

Coupling constants are reported as observed and not corrected for second order effects. Assignments were made via a combination of ^1^H, ^13^C, DEPT-135, DEPT-edited HSQC, HMBC and 2D NOESY or ROESY experiments. Where signals overlap ^1^H δ is reported as the center of the respective HSQC crosspeak. Multiplicities are described as, s = singlet, d = doublet, dd = doublet of doublets, dt = doublet of triplets, t = triplet, q = quartet, quint = quintet, tquin = triplet of quintets, m = multiplet, br = broad, appt = apparent.

#### Purification of *apo*-melianol (3) (via expression of *M. azedarach* genes)

Using vacuum infiltration 115 large *N. benthamiana* plants were infiltrated with equal volumes of *A. tumefaciens* strains harboring pEAQ-HT-DEST1 expression constructs of the following genes: *AstHMGR*, *AiOSC1, MaCYP71CD2, MaCYP71BQ5, MaCYP88A108* and *MaMOI2*. Leaves were harvested and freeze-dried six days after infiltration, yielding 159.9 g of dried leaf material. Following the preliminary triterpene extraction method described above, and for this compound utilizing the ambersep 900 hydroxide form beads to remove chlorophyll, successive rounds of fractionation were performed utilizing an Isolera Prime (Biotage) as described in (SI Table S21). Fractions containing the target were pooled, and to achieve final purification, subject to semi-preparative UHPLC, performed on an Agilent Technologies 1290 Infinity II system equipped with an Agilent Technologies 1290 infinity II Diode Array Detector (DAD), Agilent 1260 Infinity Evaporative Light Scattering Detector (ELSD) and an Agilent 1260 infinity II fraction collector. The sample was dissolved in a minimal volume of acetonitrile and injected in 200 µl aliquots. Separation was performed on a 250 x 10 mm S-5 µM 12 nm Pack pro C18 column (YMC) using water (A) versus 95% acetonitrile (B) at 4 ml/min and 40ºC with the following gradient of solvent B; 68% 0-30 min, 68-100% 30-32min, 100% 32-37 min, 100-41% 37-39 min and 41% 39-44 min. The fraction collector was programmed to collect between 22-25 min (with a maximum peak duration of 2 min) and to be triggered (threshold and peak) by detection of a peak from either the DAD or ELSD detector. DAD was set to collect signals with a wavelength of 205 nm and bandwidth of 4 nm. Fractions collected within this region (over 11 runs) were pooled and dried down. This yielded 13.1 mg of a (**3**) as a white powder on which NMR was performed in CDCl_3_ (Table S3).

#### Purification of (6) (via expression of *C. sinensis* genes)

62 *N. benthamina* plants (5-6 week old) were vacuum infiltrated using *A. tumefaciens* mediated transient expression using equal volume of infiltrated strains (OD per strain = 0.2) harboring *AtHMGR, CsOSC1, CsCYP71CD1, CsCYP71BQ4, CsCYP88A51, CsMOI2, CsL21AT*, and *CsSDR*. 871.09 g of leaves were harvested 6 days post-infiltration, dried (yielding 106.89 g) and extracted in ethyl acetate following the standard procedure outlined above. Isolation and NMR analysis of (**6**) (Table S4) was subsequently performed following the standard methods outlined above.

#### Purification of (4’) (via expression of *C. sinensis* genes)

43 *N. benthamina* plants (6-7 week old) were vacuum infiltrated using *A. tumefaciens* mediated transient expression using equal volume of infiltrated strains (OD per strain = 0.2) harboring *AtHMGR, CsOSC1, CsCYP71CD1, CsCYP71BQ4, CsCYP88A51, CsMOI1, CsL21AT* and *CsSDR*. 865.1 g of leaves were harvested 6 days post-infiltration, dried (yielding 105.5 g) and extracted in ethyl acetate following the standard procedure outlined above. Isolation and NMR analysis of (**4’**) (Table S5) was subsequently performed following the standard methods outlined above

#### Purification of 21(*S*)-acetoxyl-*apo*-melianone (6) (via expression of *M. azedarach* genes)

Using vacuum infiltration (*72*, *73*), 121 large *N. benthamiana* plants were infiltrated with equal volumes of *A. tumefaciens* strains harboring pEAQ-HT-DEST1 expression constructs of *AstHMGR, AiOSC1, MaCYP71CD2, MaCYP71BQ5, MaCYP88A108, MaMOI2, MaL21AT* and *MaSDR*. One week after infiltration, leaves were harvested and freeze-dried yielding 150.1 g of dried material. Following the preliminary extraction of triterpenes described above, successive rounds of fractionation were performed utilizing an Isolera Prime (Biotage) (Table S21). Fractions containing the target were pooled and final purification was achieved by re-crystallisation. Briefly, hot ethanol (70 ºC) was added dropwise to the sample (heated to 70 ºC) until all solids had dissolved. The sample was then covered and left at room temperature for crystals to form. Crystals were washed in cold ethanol under vacuum and then filtered by dissolving in methanol to allow collection. Initial recrystallisation was performed in triplicate, yielding ~300 mg of pale yellow product. The recrystallisation was repeated using this pale yellow product, to yield 77.25 mg of white product (**6**). 5 mg of product redissolved in CDCl_3_ for NMR (Table S6).

#### Purification of (9) (via expression of *C. sinensis* genes)

63 *N. benthamina* plants (5-6 week old) were vacuum infiltrated using *A. tumefaciens* mediated transient expression using equal volume of infiltrated strains (OD per strain = 0.2) harboring *AtHMGR, CsOSC1, CsCYP71CD1, CsCYP71BQ4, CsCYP88A51, CsMOI2, CsL21AT, CsSDR, CsCYP716AC1* and *CsCYP88A37*. 864.2 g of leaves were harvested 6 days post-infiltration, dried (yielding 89.38 g) and extracted in ethyl acetate following the standard procedure outlined above. This resulted in the isolation of 20.3 mg of (9). NMR analysis was performed following the standard methods outlined above.

#### Purification of epi-neemfruitin B (10) (via expression of *M. azedarach* genes)

Using vacuum infiltration (*72*, *73*), 143 large *N. benthamiana* plants were infiltrated with equal volumes of A. tumefaciens strains harboring pEAQ-HT-DEST1 expression constructs of the following genes: *AstHMGR, AiOSC1, MaCYP71CD2, MaCYP71BQ5, MaCYP88A108, MaMOI2, MaL21AT, MaSDR, MaCYP88A164* and *MaL1AT*. Eight days after infiltration, leaves were harvested and freeze-dried, yielding 112.5g of dried material. Following the preliminary extraction of triterpenes described above, successive rounds of fractionation were performed utilizing an Isolera Prime (Biotage) (Table S21). Fractions containing the target were pooled and dissolved in minimal volume of methanol (3 ml) for final purification via injection (500-1200 µl injection) onto a preparative HPLC. Separation was achieved using water (A) versus 95% acetonitrile (B) with the following gradient of solvent B; 42% 0-1 min, 42-73% 1-1.5 min, 73-100 1.5-11.5 min, 100% 11.5-16.5 min and 100-42% 16.5-17 min. Fractions were collected between 8-11 minutes triggered by detection of an MS peak with a mass of 526 [M] (threshold 5,000) and DAD peak (threshold of 5, wavelength 205 nm). Fractions were pooled and dried to yield 4 mg of a pale yellow product (**10**), which was dissolved in minimal ethanol and treated with activated charcoal to remove coloured impurities. This yielded 2.25 mg of purified product, which was dissolved in CDCl_3_ for NMR (Table S11).

#### Purification of the MaL7AT and MaAKR product (14) (via expression of *M. azedarach* genes)

Using vacuum infiltration (*72*, *73*), 110 medium/large *N. benthamiana* plants were infiltrated with equal volumes of *A. tumefaciens* strains harboring pEAQ-HT-DEST1 expression constructs of *AstHMGR, AiOSC1, MaCYP71CD2, MaCYP71BQ5, MaCYP88A108, MaMOI2, MaL21AT, MaSDR, MaCYP88A164, MaL1AT, MaL7AT* and *MaAKR.* Six days after infiltration, leaves were harvested and freeze-dried yielding 140.4 g of dried material. Following the preliminary extraction of triterpenes described above, initial fractionation was then performed utilizing an Isolera Prime (Biotage) (Table S21). Fractions containing the target compound were then subject to liquid-liquid partitioning (80% methanol:hexane, in triplicate). The 80% methanol fractions were pooled and re-dissolved in a minimal volume methanol (10 ml) for final purification via injection (250-1000 µl) onto preparative HPLC. Separation was performed using water (A) versus 95% acetonitrile (B) with the following gradient of solvent B: 42%-100% 0-15 min, 100% 15-19 min and 100-42% 19-19.5 min. Fractions were collected between 9-11.5 minutes triggered by a peak of mass 570 [M+ACN+H]^+^ (threshold 5,000). The [M+ACN+H]^+^ adduct mass was used as an inputted mass rather than [M] or [M+H]^+^ due to the high accumulation of acetonitrile adducts for this intermediate in this system. Ten fraction collecting runs were performed and the pooled fractions yielded ~20 mg of product. Initially, 4 mg of product was dissolved in CDCl_3_ for NMR, however this was converted to the known protolimonoid, gradifoliolenone (36) (Fig. S1), in solution. A further 4 mg of product was dissolved in pyridine-d_5_ however a suspected rotamer effect was observed. Therefore NMR characterisation was finally performed by dissolving 5 mg of product in benzene-d_6_ (Table S13). The fact that gradifoliolenone has previously been reported as isolated from nature, whilst the true product (**14**) has not, may suggest that some of the protolimonoid E-ring structures isolated to date could be the result of spontaneous or extraction method-derived modifications, and may be red-herrings in terms of their involvement in limonoid biosynthesis.

#### Purification of kihadalactone A (19) (via expression of *C. sinensis* genes)

63 *N. benthamina* plants (5-6 week old) were vacuum infiltrated using *A. tumefaciens* mediated transient expression using equal volume of infiltrated strains (OD per strain = 0.2) harboring *AtHMGR, CsOSC1, CsCYP71CD1, CsCYP71BQ4, CsCYP88A51, CsMOI2, CsL21AT, CsSDR, CsCYP716AC1, CsCYP88A37, CsL1AT, CsL1A7* and *CsLFS*. 705.7 g of leaves were harvested 6 days post-infiltration, dried (yielding 79.15 g) and extracted in ethyl acetate following the standard procedure outlined above. Isolation and NMR analysis of (**19**) (Table S14) was subsequently performed following the standard methods outlined above.

#### Purification of azadirone (18) (from *A. indica* leaf powder)

224.9 g of neem (*A. indica*) leaf powder (purchased from H&C Herbal Ingredients Expert) was extracted following the preliminary triterpene extraction method described above. Following this the extract was partitioned between ethyl acetate (800 ml) and water (800 ml) which yielded 21.2 g of crude extract. Initial fractionation was then performed utilizing an Isolera Prime (Biotage) (Table S21) following a method adapted from previous reports of azadirone isolation from *A. indica* fruits (*74*). Fractions containing azadirone were then dissolved in a minimal volume of methanol (with dropwise addition of ethyl acetate), before being filtered, through both a Sep-Pak vac 3cc C18 cartridge (Waters) and a minisart highflow PES 0.22 µM syringe filter (Sartorius), before injection (7000 µl) onto a preparative HPLC system. MS was collected via MM-ES+APCI in SIM mode, detecting and collecting for a mass of 437.2 [M+H]^+^. Fractions were collected between 14-20 min (threshold 5,000). Initial separation was performed using water (A) versus acetonitrile (B) with the following gradient of solvent B; 65% 0-1.5 min, 60-100% 1.5-26.5 min, 100%, 26.5-30 min and 100-65% 30-30.5 min. After 3 runs, fractions of azadirone (**18**) with a reasonable level of purity were pooled, yielding ~1 mg of purified product, which was dissolved in CDCl_3_ for NMR (Table S15).

#### Purification of MaCYP716AD4 side-product (20) (via expression of *M. azedarach* genes)

Using vacuum infiltration (*72*, *73*), 120 medium/large *N. benthamiana* plants were infiltrated with equal volumes of *A. tumefaciens* strains harboring pEAQ-HT-DEST1 expression constructs of *AstHMGR, AiOSC1, MaCYP71CD2, MaCYP71BQ5, MaCYP88A108, MaMOI2, MaL21AT, MaSDR, MaCYP88A164, MaL1AT, MaAKR* and *MaCYP716AD4*. Eight days after infiltration, leaves were harvested and freeze-dried yielding 123.1 g of dried material. Following the preliminary extraction of triterpenes described above, initial fractionation was then performed utilizing an Isolera Prime (Biotage) (Table S21). Fractions containing the target were then dissolved in a minimal volume of 80% acetonitrile (6 ml) before injection (500-1500 µl) onto a preparative HPLC system. For this intermediate MS was collected via MM-ES+APCI in SIM mode, detecting and collecting for a mass of 503.4 [M+H]^+^. Fractions were collected between 1.5-10 min (threshold 5,000). Initial separation was performed using water (A) versus acetonitrile (B) with the following gradient of solvent B: 60% 0-0.5 min, 60-75% 0.5-10 min, 75-100% 10.-10.5 min, 100% 10.5-15 min and 100-60% 15-15.5 min. After 9 runs, fractions containing target were pooled and further purified by a second round of preparative HPLC, using the same instrument settings, but a different gradient consisting of water (A) versus methanol (B) with the following gradient of solvent B: 67% 0-0.5 min, 67-77% 0.5-20 min, 77-100% 20-20.5 min, 100% 20.5-24.5 min and 100-67% 24.5-25 min. After two injections, fractions containing the target were pooled, yielding ~0.6 mg of purified product (**20**), which was dissolved in benzene-d_6_ for NMR (Table S16).

### **Supplementary Figures**

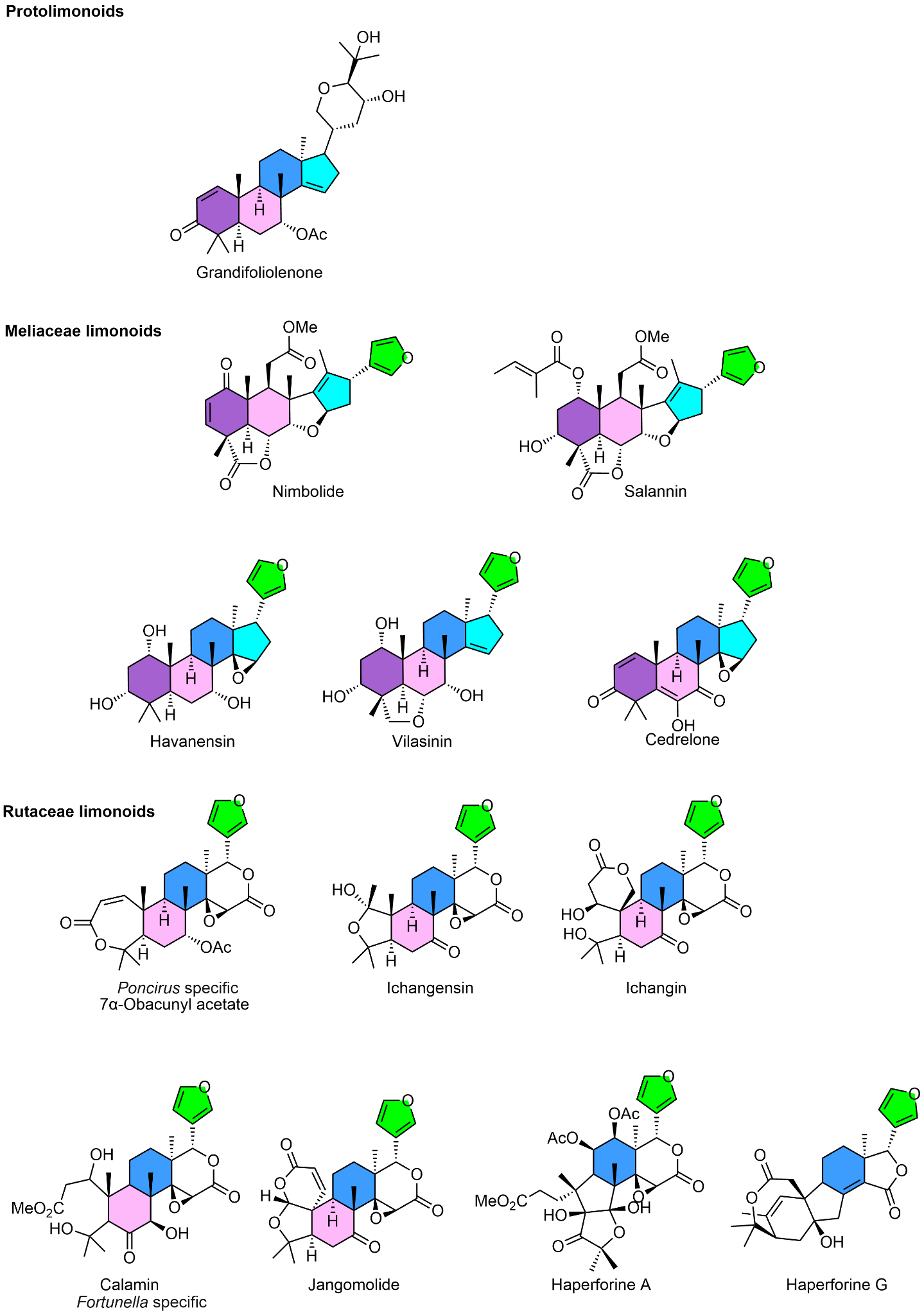

##### Fig. S1. Supplementary limonoid and protolimonoid structures.

Structures of additional limonoids and protolimonoids relevant to the main text. Ring A-D and the furan moiety are colored to show the cleavage and conservation of each ring.

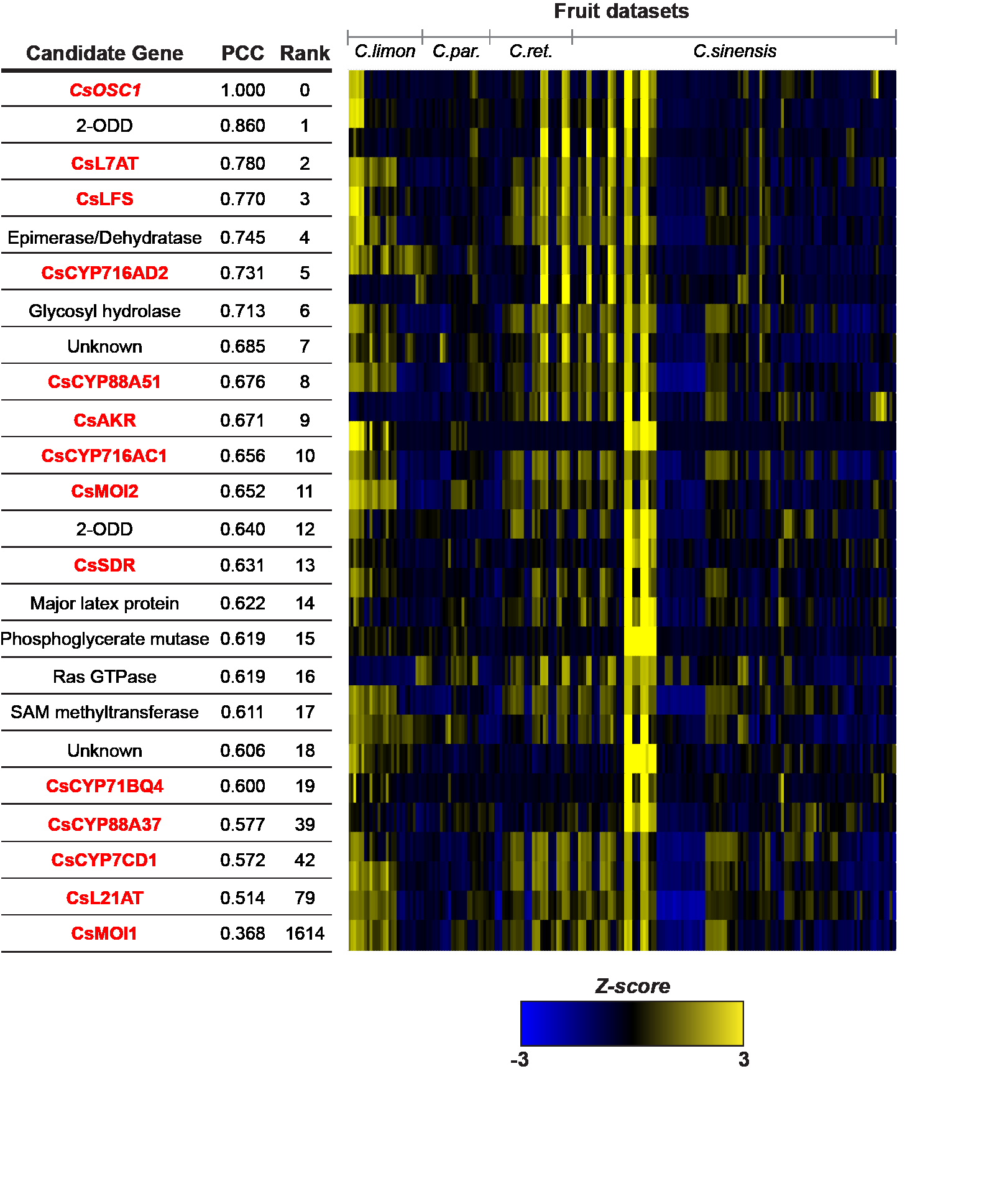

##### Fig. S2. Co-expression analysis of *C. sinensis* publicly available microarray expression data from Network inference for Citrus Co-Expression (NICCE) using *CsOSC1* as a bait gene.

Linear regression analysis was used to rank the top 20 genes based on Pearson’s correlation coefficient (PCC) to *CsOSC1*. Heat map displays Z-score calculated from log_2_ normalized expression across fruit datasets. Genes in red indicate candidates characterized in this study or our previous work (*17*). CYPs and acetyltransferases in the top 100 were first selected to screen for their activities by co-expression with *Cs*OSC1, *Cs*CYP71CD1 and *Cs*CYP71BQ4.

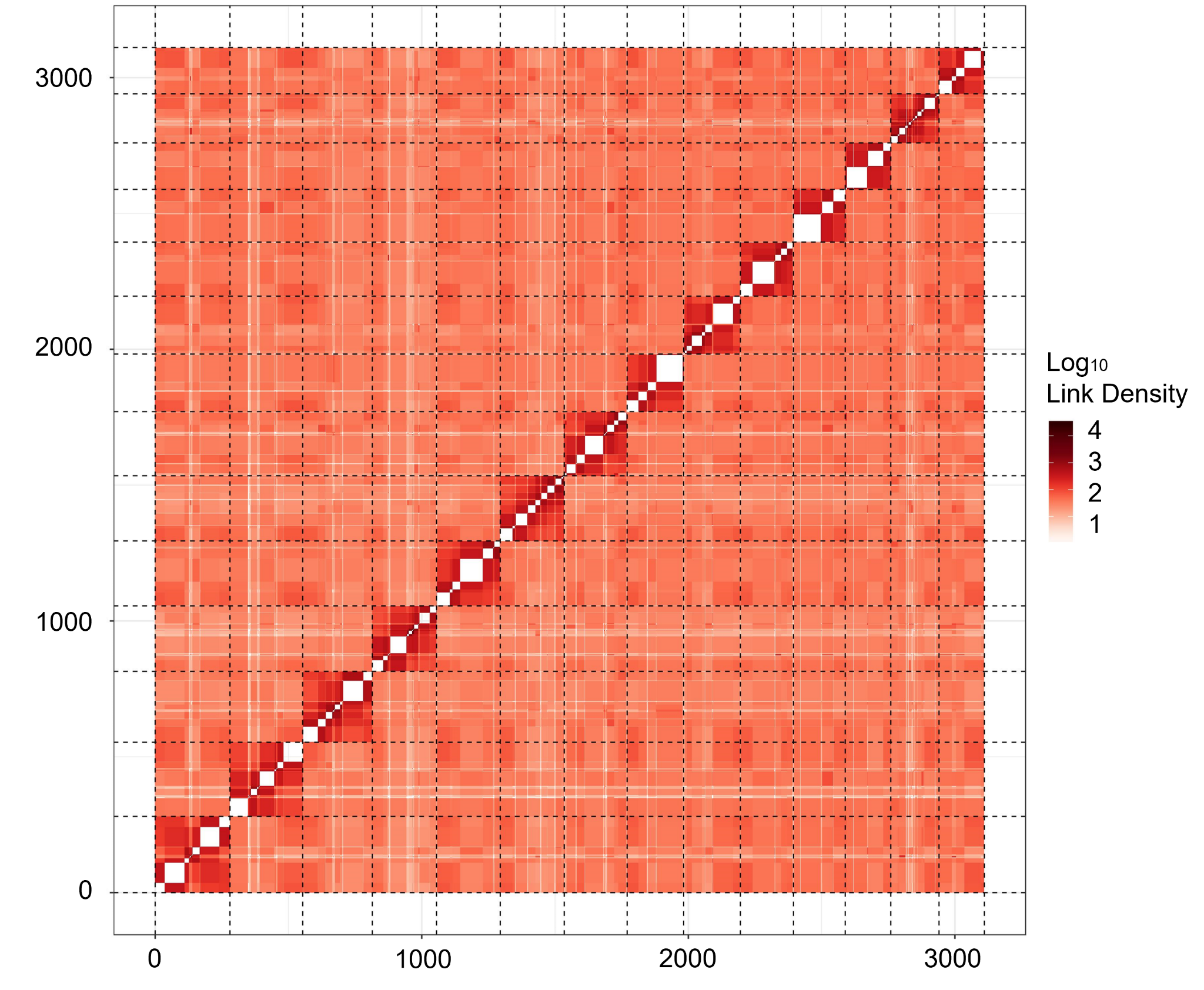

##### Fig. S3. Hi-C heat map post-scaffolding heatmap of *M. azedarach* genome.

Analysis and generation of heatmap was performed by Phase Genomics. The genome has been divided into 3,000 bins (length = 75,470 bp) for this analysis. The density of Hi-C links is plotted (red). Links between the same contig are not shown (white). White boxes therefore indicate draft assembly contigs.

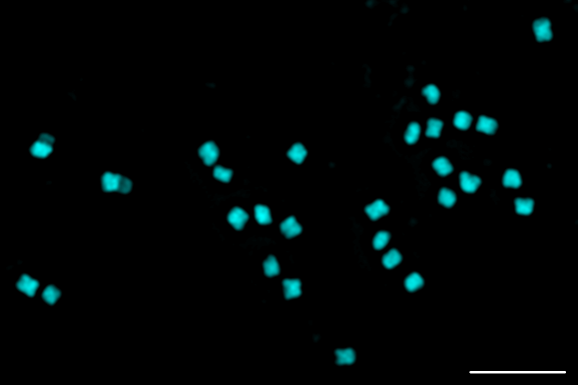

##### Fig. S4. Karyotyping of *M. azedarach.*

Representative image of a mitotic metaphase spread of *M. azedarach* (individual ‘11’) showing 28 chromosomes (2n=28). Scale bar = 5µm.

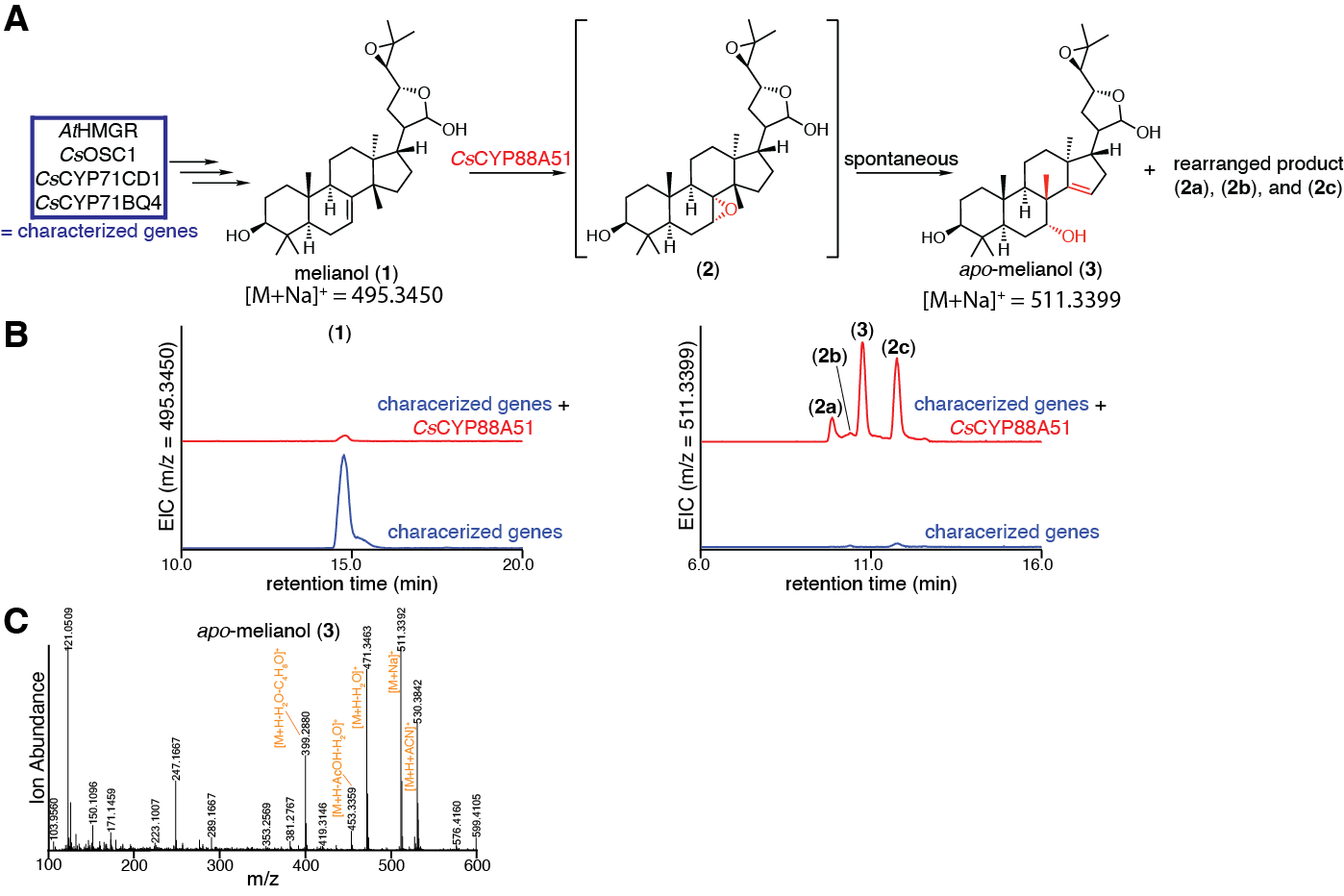

##### Fig. S5. Characterization of *Cs*CYP88A51.

(A) Predicted function of *Cs*CYP88A51 in converting (**1**) to an unstable epoxide intermediate (**2**), which spontaneously rearranges into uncharacterized products (**2a**), (**2b**) and (**2c**). (B) Extracted ion chromatograms (EICs) for extracts of *N. benthamiana* agro-infiltrated with the characterized genes listed in panel A either alone (blue) or with the addition of *Cs*CYP88A51 (red). EICs are displayed for masses of 495.3450 (calculated mass for (**1**) [M+Na]^+^) or 511.3399 (calculated mass for (**2a-c**) and (**3**) [M+Na]^+^). (C) Mass spectrum of (**3**) in panel B. Note that [M+Na]^+^ doesn’t fragment well in MSMS and the parent peak [M+H]^+^ is too low to be useful for MSMS analysis. Representative EICs and mass spectra are displayed for experiments of n=6.

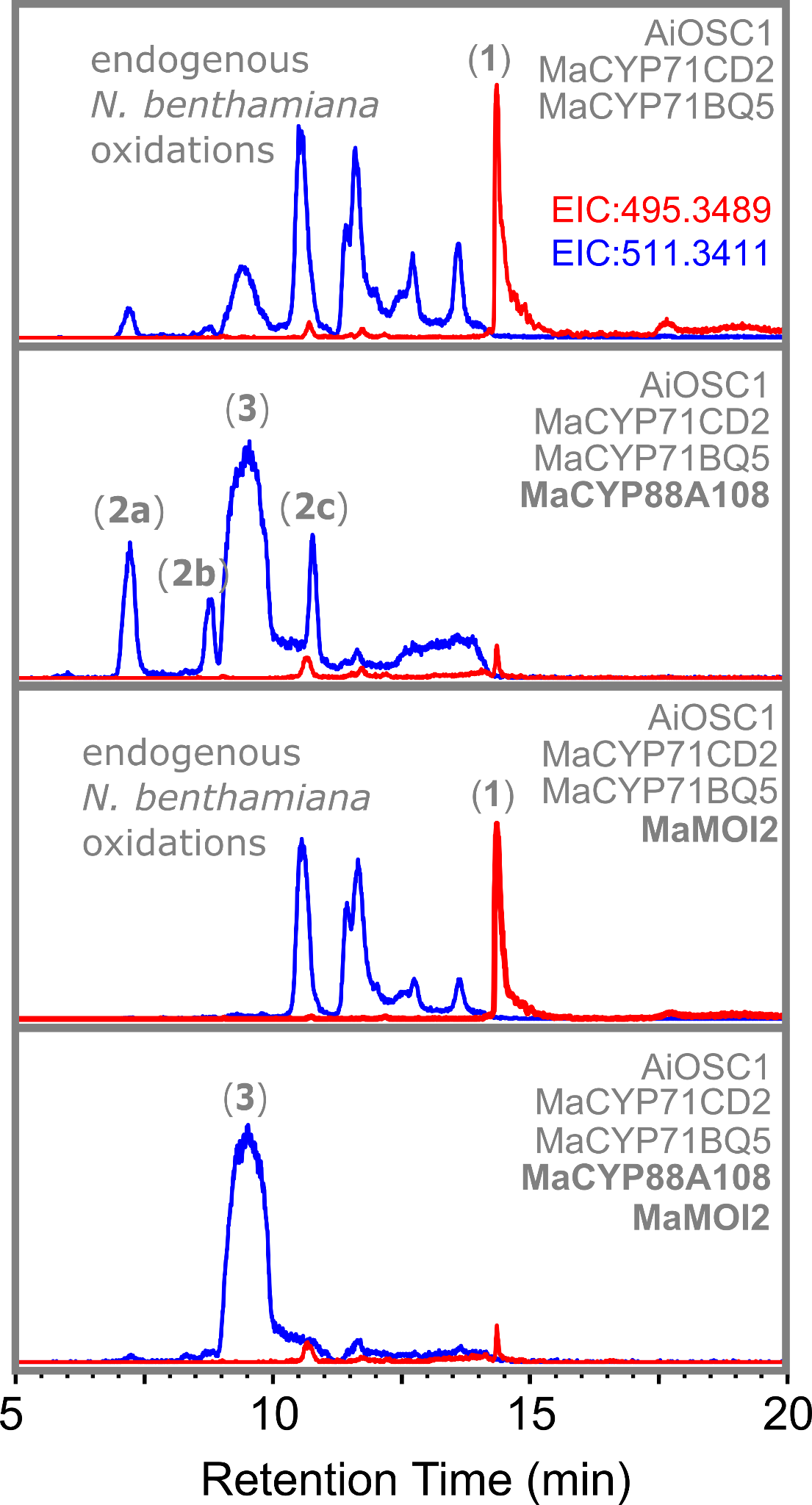

##### Fig. S6. Individual activity of *Ma*CYP88A108 and *Ma*MOI2.

Extracted ion chromatograms (EICs) of methanol extracts from agroinfiltrated *N. benthamiana* leaves expressing *MaMOI2* in combination with melianol biosynthetic genes (*AiOSC1*, *MaCYP71CD2* and *MaCYP71BQ5*), with and without *MaCYP88A108*. EICs displayed are for the masses of [melianol (**1**)+Na]^+^=495.3489 (red, calculated mass) and [melianol(**1**)+O+Na]^+^=511.3411 (blue, calculated mass). For these LCMS traces, analysis was performed using an UHPLC-IT-TOF (Shimadzu) instrument following a method and methanol gradient previously described for the analysis of protolimonoids (*17*). Characterisation of *Ma*CYP88A108 and *Ma*MOI2 expressed together is available (Fig. S7).

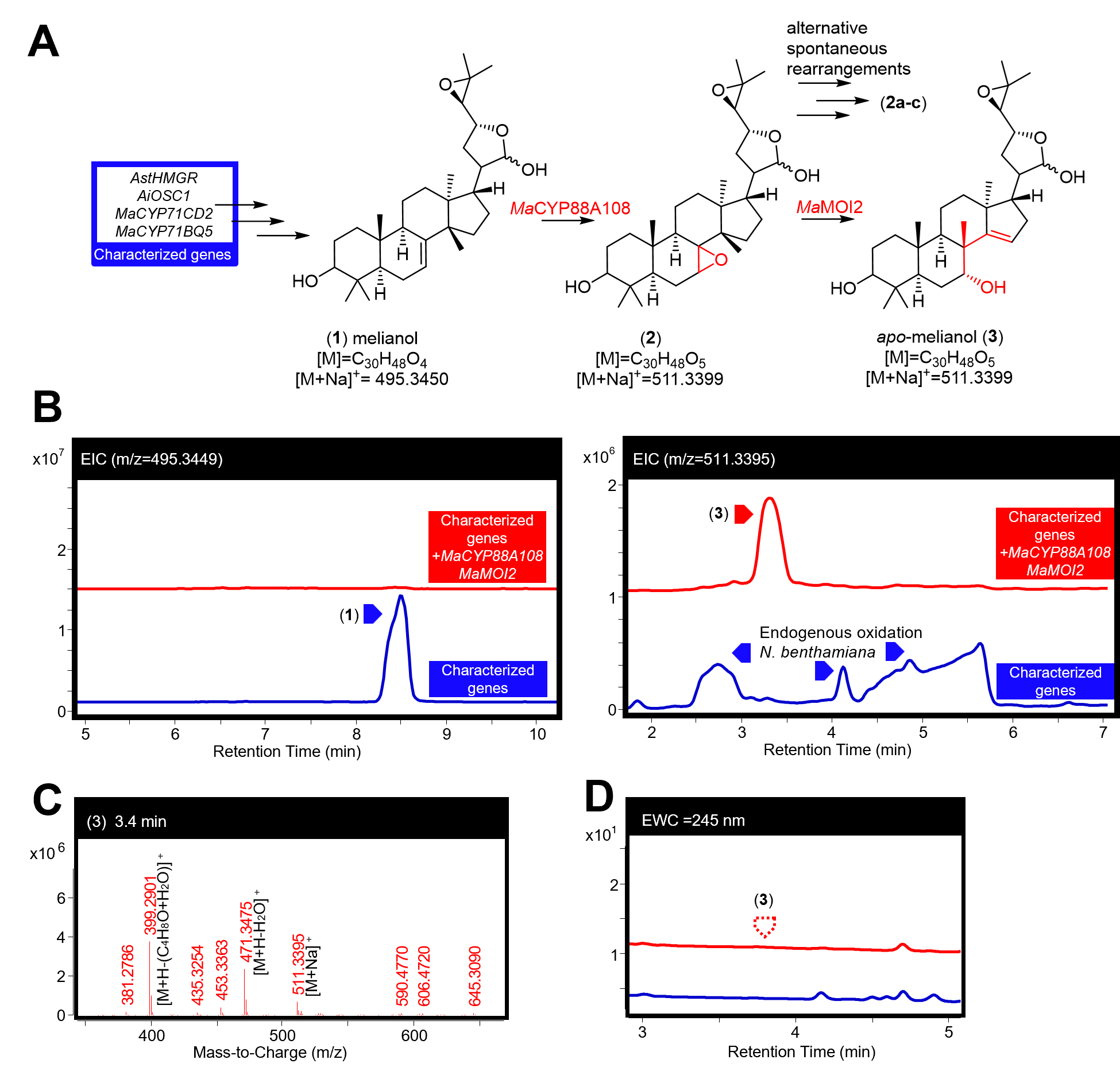

##### Fig. S7. Characterisation of *Ma*CYP88A108 and *Ma*MOI2.

(A) Function of *Ma*CYP88A108 and *Ma*MOI2 in converting melianol (**1**) to the epimeric mixture *apo*-melianol (**3**), confirmed by NMR (Table S3). Calculated masses for the major observed adducts are listed. (B) Extracted ion chromatograms (EICs) for extracts of *N. benthamiana* agro-infiltrated with the characterized genes listed in panel A either alone (blue) or with the addition of *MaCYP88A108* and *MaMOI2* (red). The EICs are displayed for masses of 495.3449 (observed mass for [(**1**)+Na]^+^) and 511.3395 (observed mass for [(**3**)+Na]^+^). (C) Mass spectra of (**3**) being heterologously produced in *N. benthamiana*, the main observed adduct ([M+Na]^+^) and fragments (including loss of water [M+H-H_2_O]^+^, and loss of water and four-carbon epoxide containing fragment [M+H-(H_2_O+C_4_H_8_O)]^+^) are labeled. (D) Extracted wavelength chromatograms (EWCs), of 245 nm (width of 4nm) for the extracts displayed in panel B. Due to the lack of an enone system in (**3**) no UV peak is observed. Representative traces and spectra are displayed (n=6). Traces showing individual activity of *Ma*CYP88A108 and *Ma*MOI2 are available (Fig. S6).

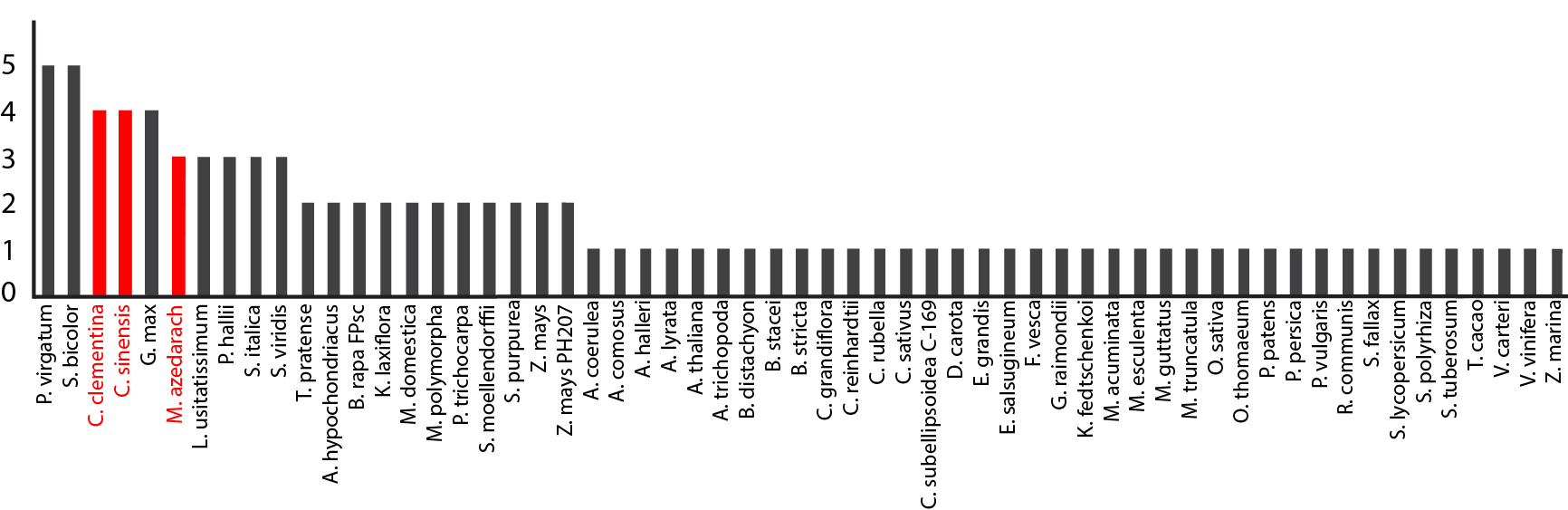

##### Fig. S8. Histogram of the number of sterol isomerase genes present in high-quality plant genomes.

Plant genomes from high-quality and annotated genomes were downloaded from Phytozome (*75*). Sterol isomerases sequences were identified by pFAM assignment to EBP (PF05241).

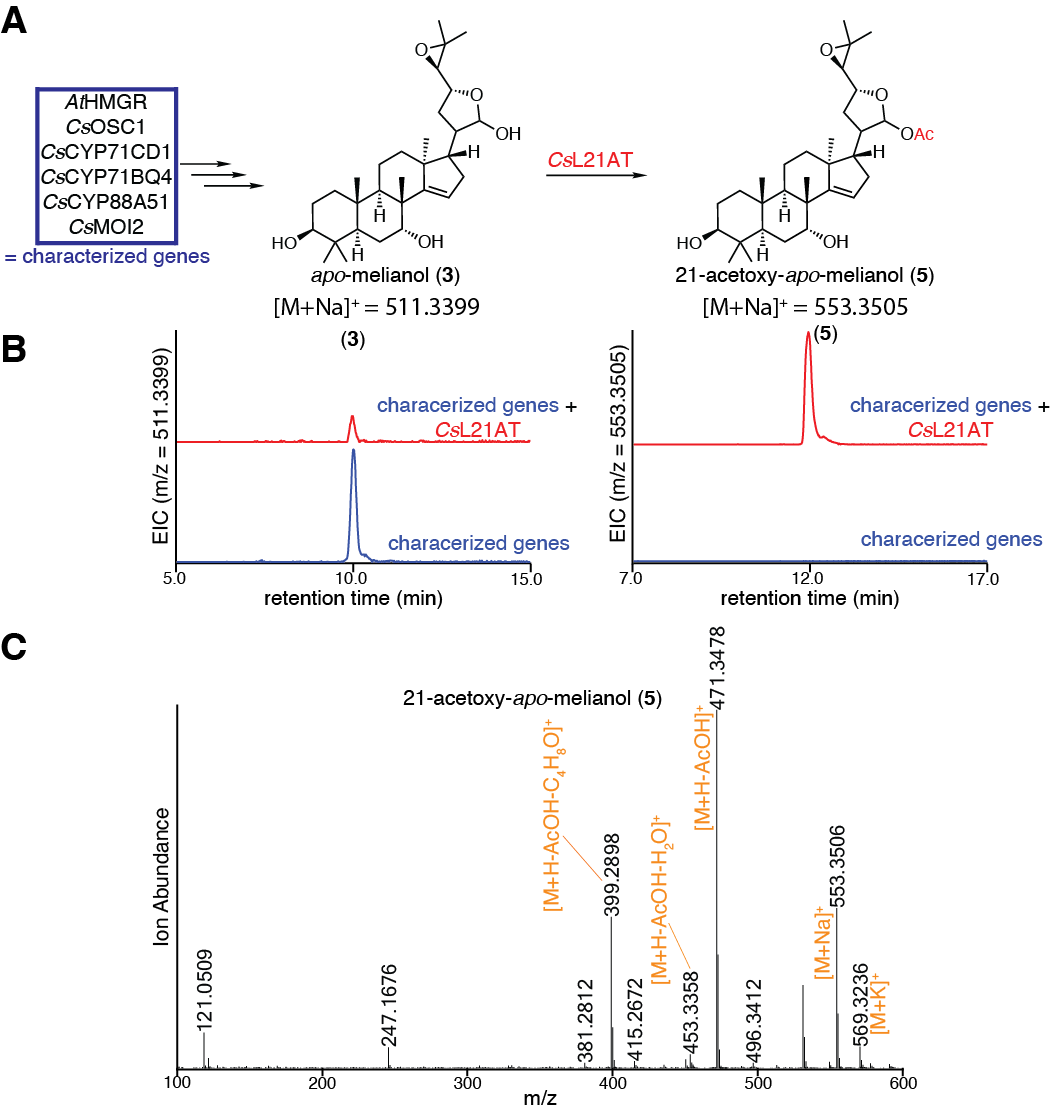

##### Fig. S9. Characterization of *Cs*L21AT.

(A) Predicted function of *Cs*L21AT in converting *apo-*melianol (**3**) to 21-acetoxyl-*apo-*melianol (**5**). (B) Extracted ion chromatograms (EICs) for extracts of *N. benthamiana* agro-infiltrated with the characterized genes listed in panel A, either alone (blue) or with the addition of *Cs*L21AT (red). EICs are displayed for masses of 511.3399 (calculated mass for (**3**) [M+Na]^+^) and 553.3505 (calculated mass for (**5**) [M+Na]^+^). **(C)** Mass spectrum of (**5**) heterologously produced in *N. benthamiana* as shown in panel B. Note that [M+Na]^+^ doesn’t fragment well in MSMS and the parent peak [M+H]^+^ is too low to be useful for MSMS analysis. Representative EICs and mass spectra are displayed (n=6).

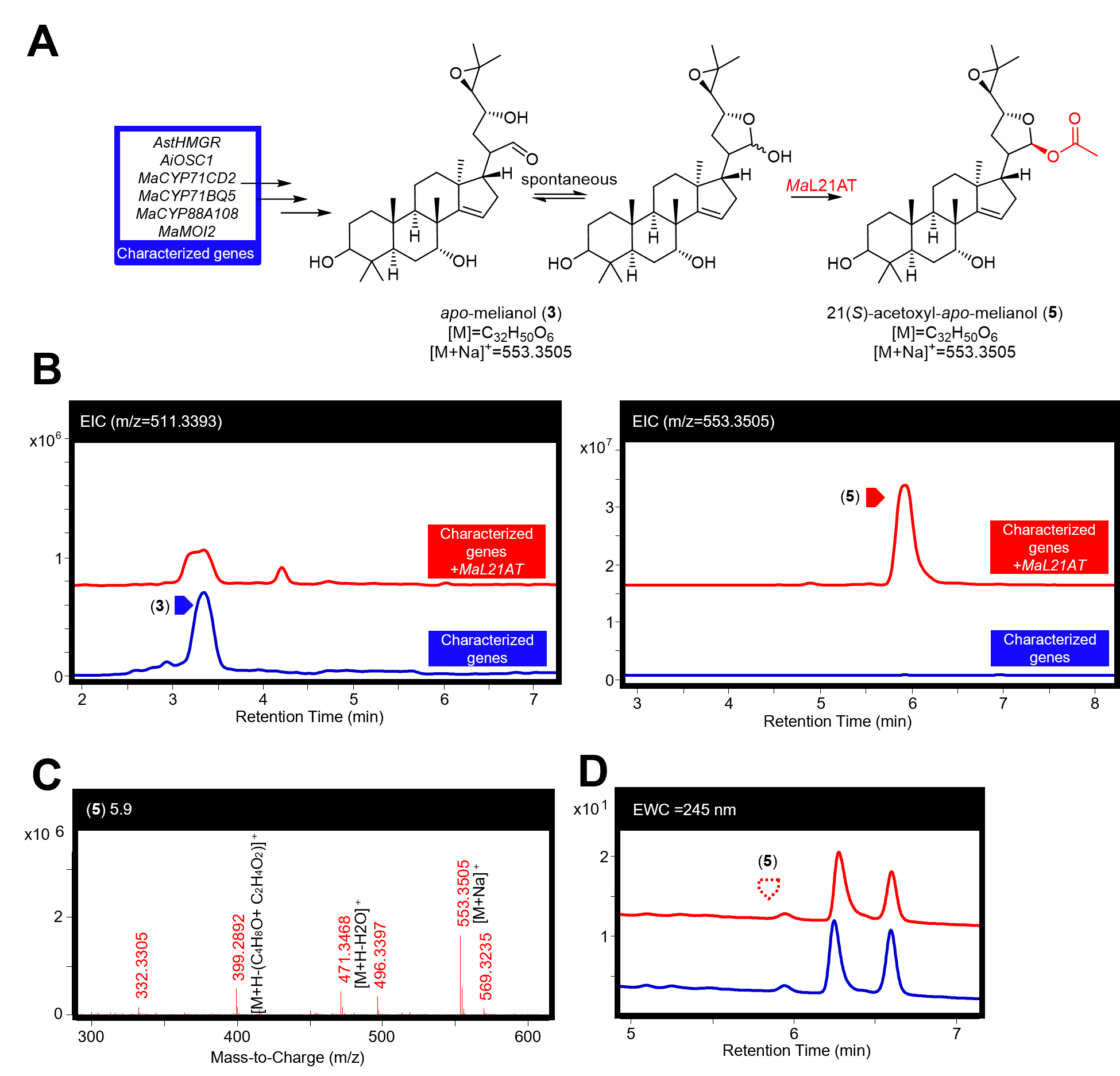

##### Fig. S10. Characterisation of *Ma*L21AT.

(A) Function of *Ma*L21AT in producing 21-acetoxyl-*apo*-melianol (**5**) (confirmed by NMR of later products (Fig. S15, Table S6 to S7)) from *apo*-melianol (**3**). (B) Extracted ion chromatograms (EICs) for extracts of *N. benthamiana* agro-infiltrated with the characterized genes listed in panel A either alone (blue), or with the addition of *Ma*L21AT (red). The EICs are displayed for masses of 511.3393 (observed mass for [(**3**)+Na]^+^) and 553.3505 (observed mass for [(**5**)+Na]^+^). (C) Mass spectra for (**5**) being heterologously produced in *N. benthamiana,* the main observed adduct ([M+Na]^+^) and fragments (including loss of acetic acid [M+H-C_2_H_4_O_2_]^+^ and loss the four-carbon epoxide containing fragment and an acetic acid [M+H-(C_4_H_8_O+C_2_H_4_O_2_)]^+^) are labeled. (D) Extracted wavelength chromatograms (EWCs), of 245 nm (width of 4nm), for the extracts displayed in panel B. Due to the lack of an enone system in (**5**) no UV peak is observed. Representative traces and spectra are displayed (n=6).

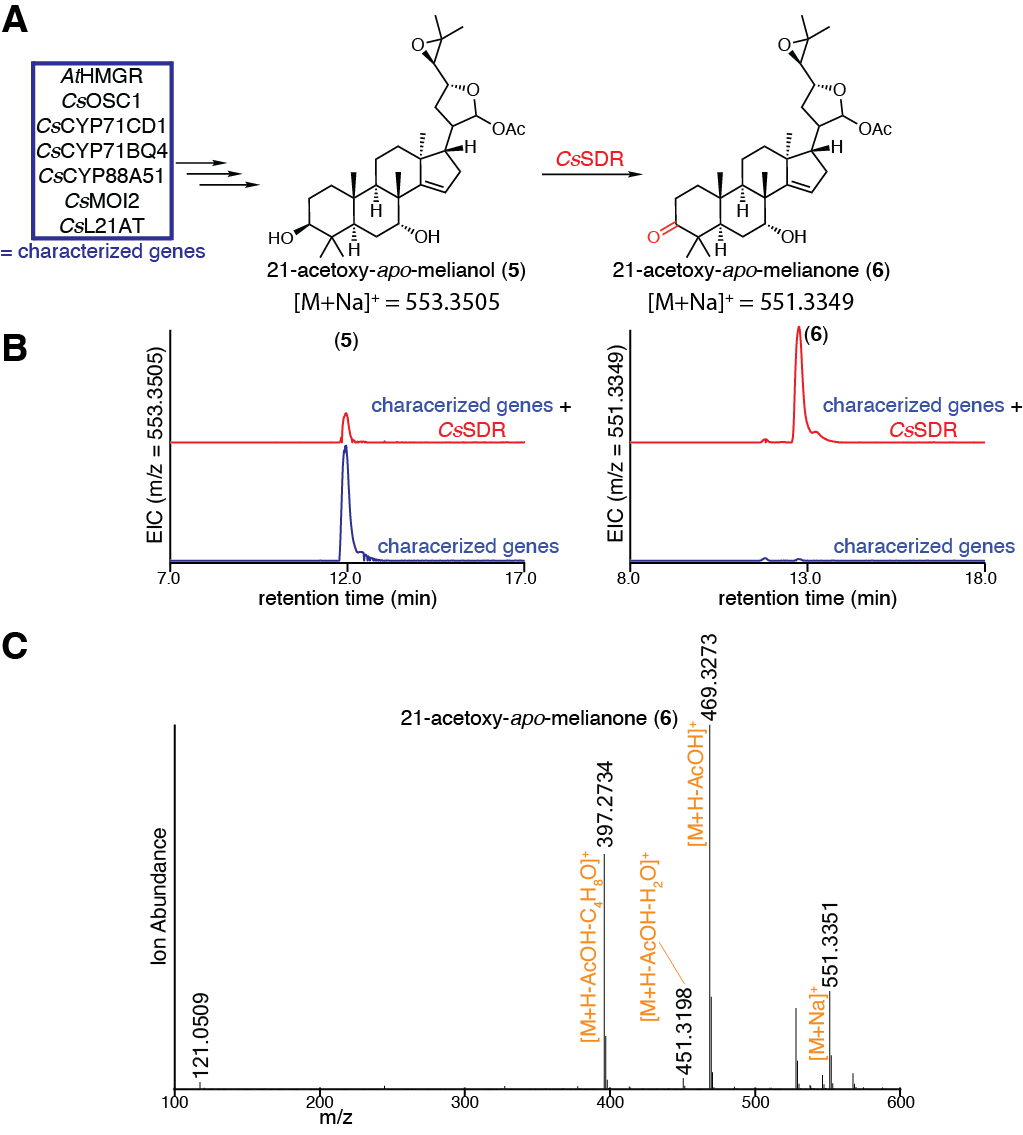

##### Fig. S11. Characterization of *Cs*SDR.

(A) Predicted function of *Cs*SDR in converting 21-acetoxyl-*apo-*melianol (**5**) to 21-acetoxyl-*apo-*melianone (**6**). (B) EICs for extracts of *N. benthamiana* agro-infiltrated with the characterized genes listed in panel A, either alone (blue) or with the addition of *Cs*SDR (red). EICs are displayed for masses of 553.3505 (calculated mass for (**5**) [M+Na]^+^) or 551.3349 (calculated mass for (**6**) [M+Na]^+^). (C) Mass spectrum of (**6**) heterologously produced in *N. benthamiana* as shown in panel B. Note that [M+Na]^+^ doesn’t fragment well in MSMS and the parent peak [M+H]^+^ is too low to be useful for MSMS analysis. Representative EICs and mass spectra are displayed (n=6).

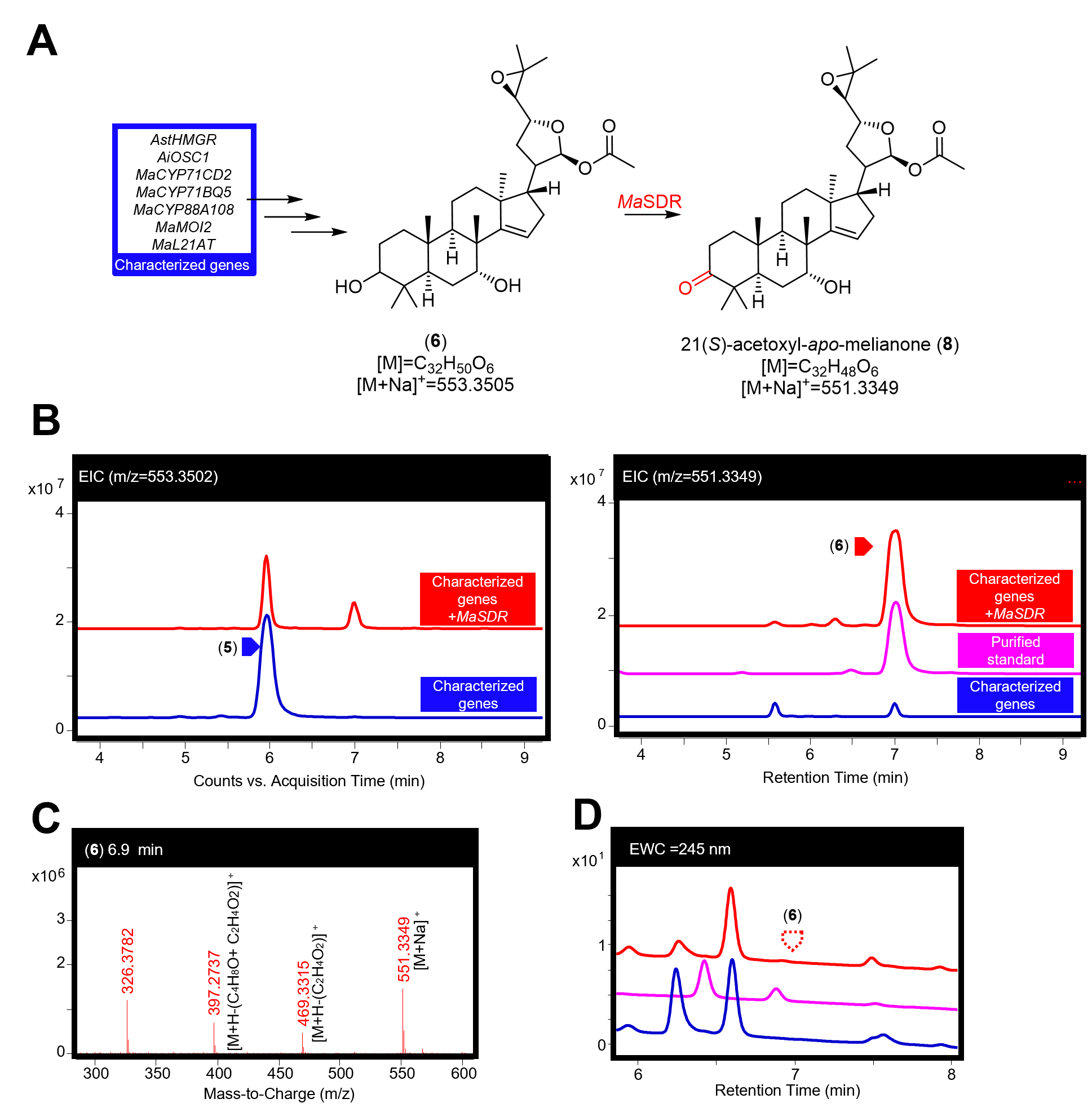

##### Fig. S12. Characterisation of *Ma*SDR*.*

(A) Function of *Ma*SDR in producing 21(*S*)-acetoxyl-*apo*-melianone (**6**) (confirmed by NMR (Fig. S15, Table S6 to S7)) from 21-acetoxyl-*apo*-melianol (**5**). (B) Extracted ion chromatograms (EICs) for extracts of *N. benthamiana* agro-infiltrated with the characterized genes listed in panel A, either alone (blue) or with the addition of *MaSDR* (red), along with a purified standard (pink). The EICs are displayed for masses of 553.3502 (observed mass for [(**5**)+Na]^+^) and 551.3349 (observed mass [(**6**)+Na]^+^). (C) Mass spectra for (**6**) being heterologously produced in *N. benthamiana,* the main observed adduct ([M+Na]^+^) and fragments (including loss of acetic acid [M+H-C_2_H_4_O_2_]^+^ and loss the four-carbon epoxide containing fragment and an acetic acid [M+H-(C_4_H_8_O+C_2_H_4_O_2_)]^+^) are labeled. (D) Extracted wavelength chromatograms (EWCs), of 245 nm (width of 4nm), for extracts displayed in panel B. Due to the lack of an enone system in (**6**) no UV peak is observed. Traces of purified standards have been scaled for comparison. Representative traces and spectra are displayed (n=6).

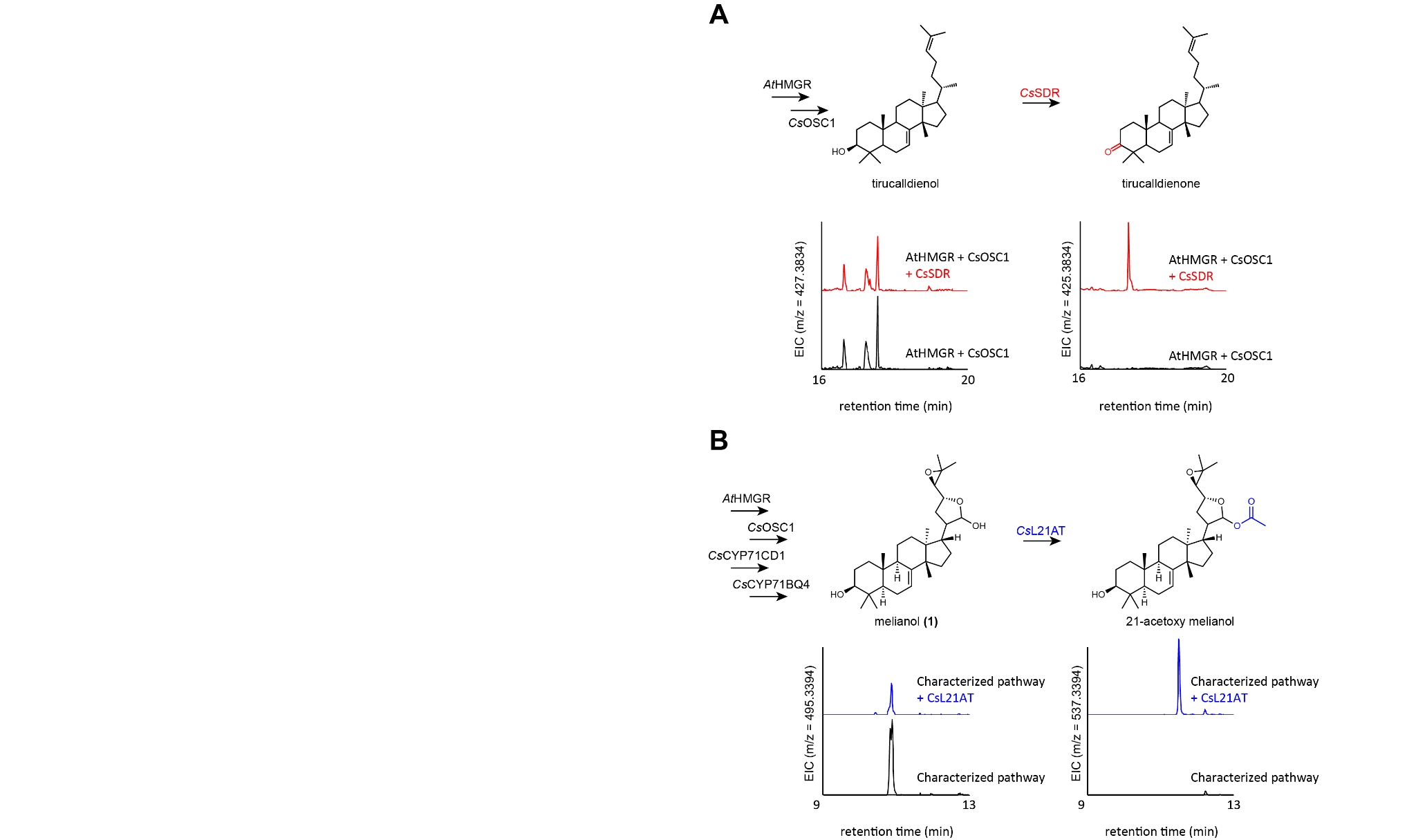

##### Fig. S13. Substrate promiscuity of *Cs*L21AT and *Cs*SDR.

(A) Representative extracted ion chromatograms (EICs) for extracts of *N. benthamiana* agro-infiltrated with *At*HMGR and *Cs*OSC1 with and without *Cs*SDR. (B) Representative EICs for extracts of *N. benthamiana* agro-infiltrated with *At*HMGR, *Cs*OSC1, *Cs*CYP71CD1 and *Cs*CYP71BQ4 with and without *Cs*L21AT. Representative EICs are displayed (n=3).

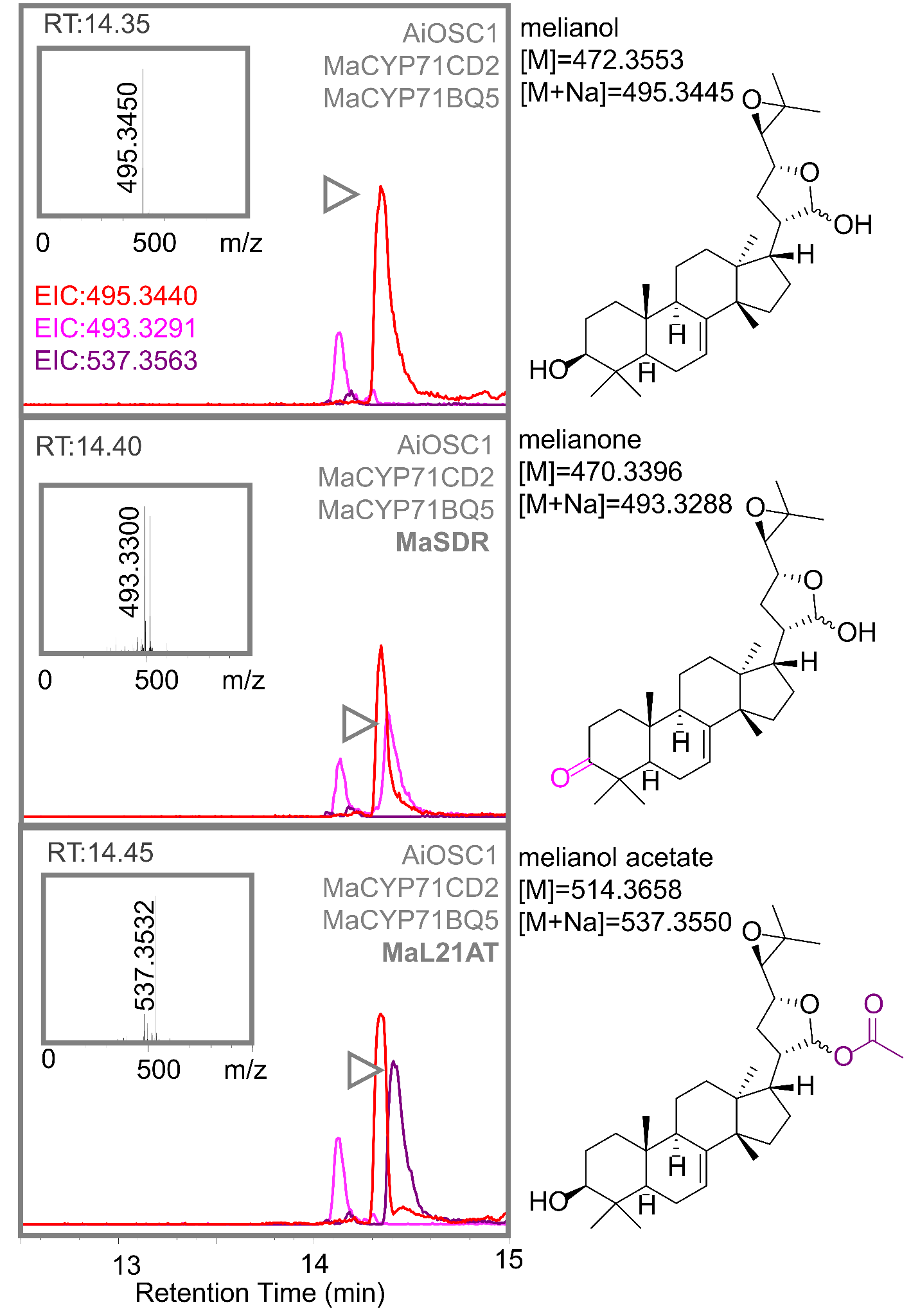

##### Fig. S14. *Ma*SDR and *Ma*L21AT can also function on melianol scaffold

UHPLC-IT-TOF generated extracted ion chromatograms (EICs) of methanol extracts from agro-infiltrated *N.benthamiana* leaves expressing *Ma*SDR and *Ma*L21AT in combination with melianol biosynthetic enzymes (*Ai*OSC1, *Ma*CYP71CD2 and *Ma*CYP71BQ5). EICs displayed are for the following observed adducts: [melianol (**1**)+Na]^+^=495.3440 (red), [melianol (**1**)-2H+Na]^+^=493.3291 (pink) and [melianol (**1**)+CH3COO+Na]^+^=537.3563 (purple). Mass spectra of new peaks (highlighted with gray arrow) are given. UHPLC-IT-TOF analysis performed using the methanol gradient previously described for the Shimadzu IT-TOF instrument (*17*). Predicted structures for labeled peaks are also provided (with exact mass and calculated sodium adduct). Representative EICs are displayed (n=3).

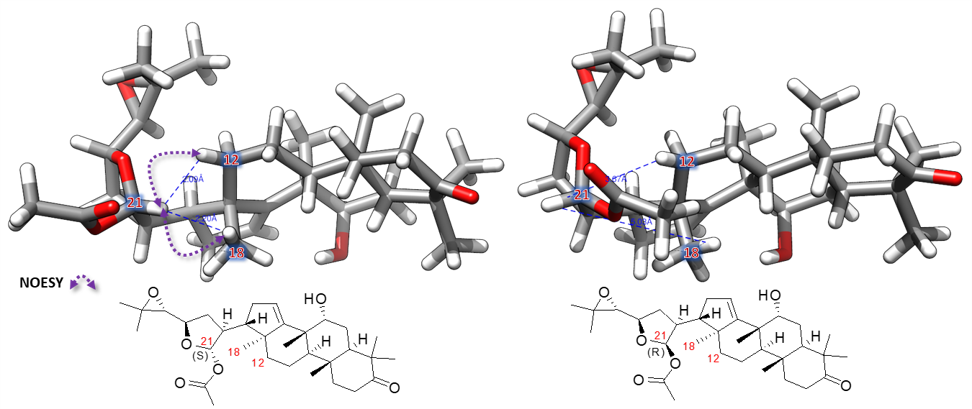

##### Fig. S15. 3D models of 21(*S*)-acetyloxy-apo-melianone (6) and 21(*R*)-acetyloxy-apo-melianone.

Geometry optimized by molecular dynamics (forcefield: MMFF94, number of steps: 500, algorithm: steepest descent and convergence: 10e-7, run by AvogadroV 1.1.1). NOEs between C21-H, C18-H3 and C12-H2 observed in 2D NOESY experiments are consistent with assignment as 21(*S*)-acetyloxy-apo-melianone. Full assignment is given (Table S6).

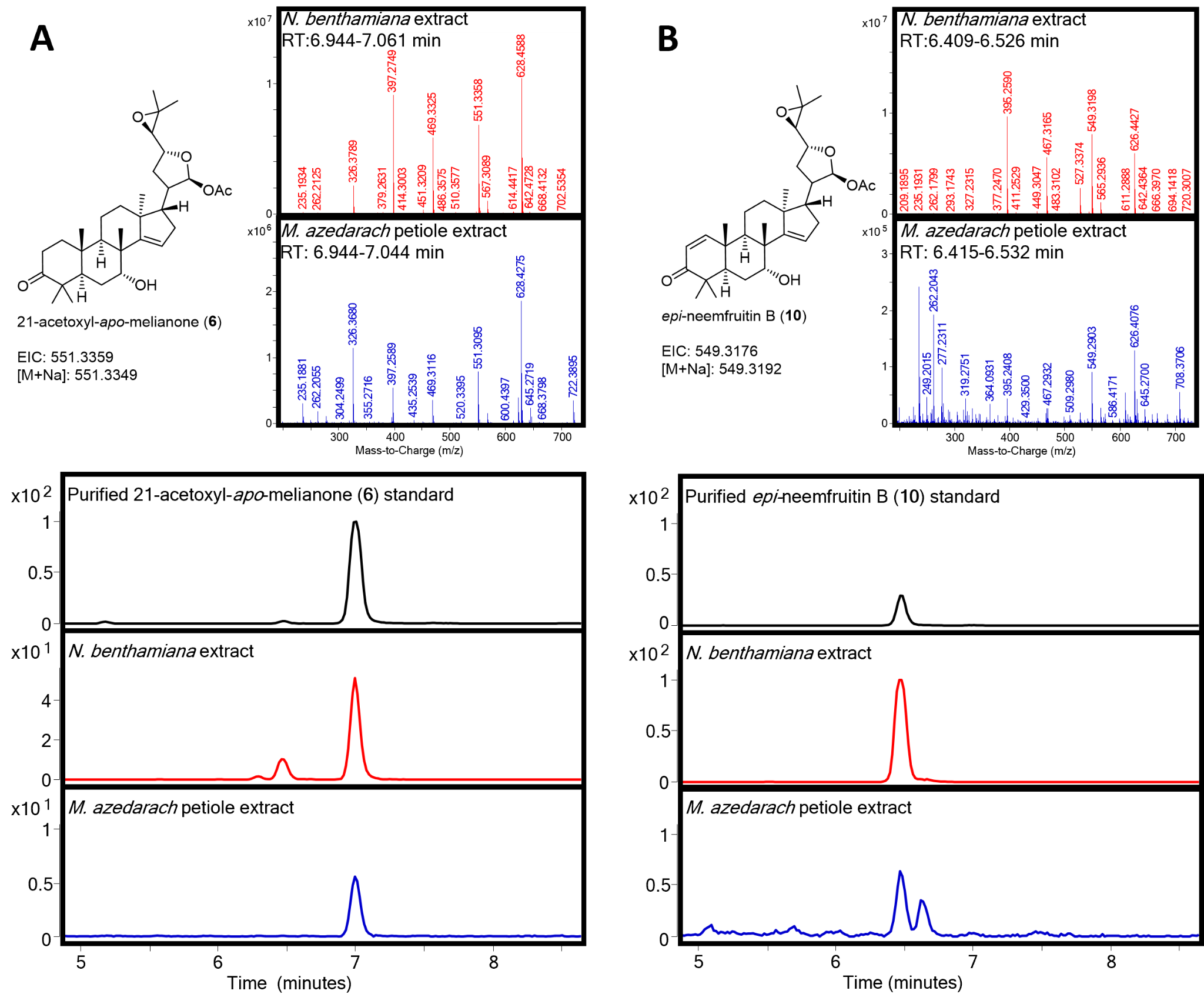

##### Fig. S16. Detection of 21-acetoxyl-*apo*-melianone (6) and *epi-*neemfruitin B (10) in *Melia azedarach* samples.

(A) Structure, mass spectra and extracted ion chromatograms (EICs) comparing extracts from *N. benthamiana* expressing 21-acetoxyl-*apo*-melianone (**6**) biosynthetic enzymes (*Ai*OSC1*, Ma*CYP71CD2, *Ma*CYP71BQ5, *Ma*CYP88A108*, Ma*MOI2, *Ma*L21AT and *Ma*SDR) to extracts from *M. azedarach* petiole tissues (individual 11). EIC of purified 21-acetoxyl-*apo*-melianone (**6**) (Table S6) is also displayed. (B) Structure, mass spectra and extracted ion chromatograms (EICs) comparing extracts from *N. benthamiana* expressing *epi*-neemfruitin B (**10**) biosynthetic enzymes (the enzymes described in panel (A) with addition of *Ma*CYP88A164 and *Ma*L1AT) to extracts from *M. azedarach* petiole tissues (individual 11). EIC of *epi-*neemfruitin B (**10**) is also displayed (Table S11). Representative EICs and spectra are displayed (n=3).

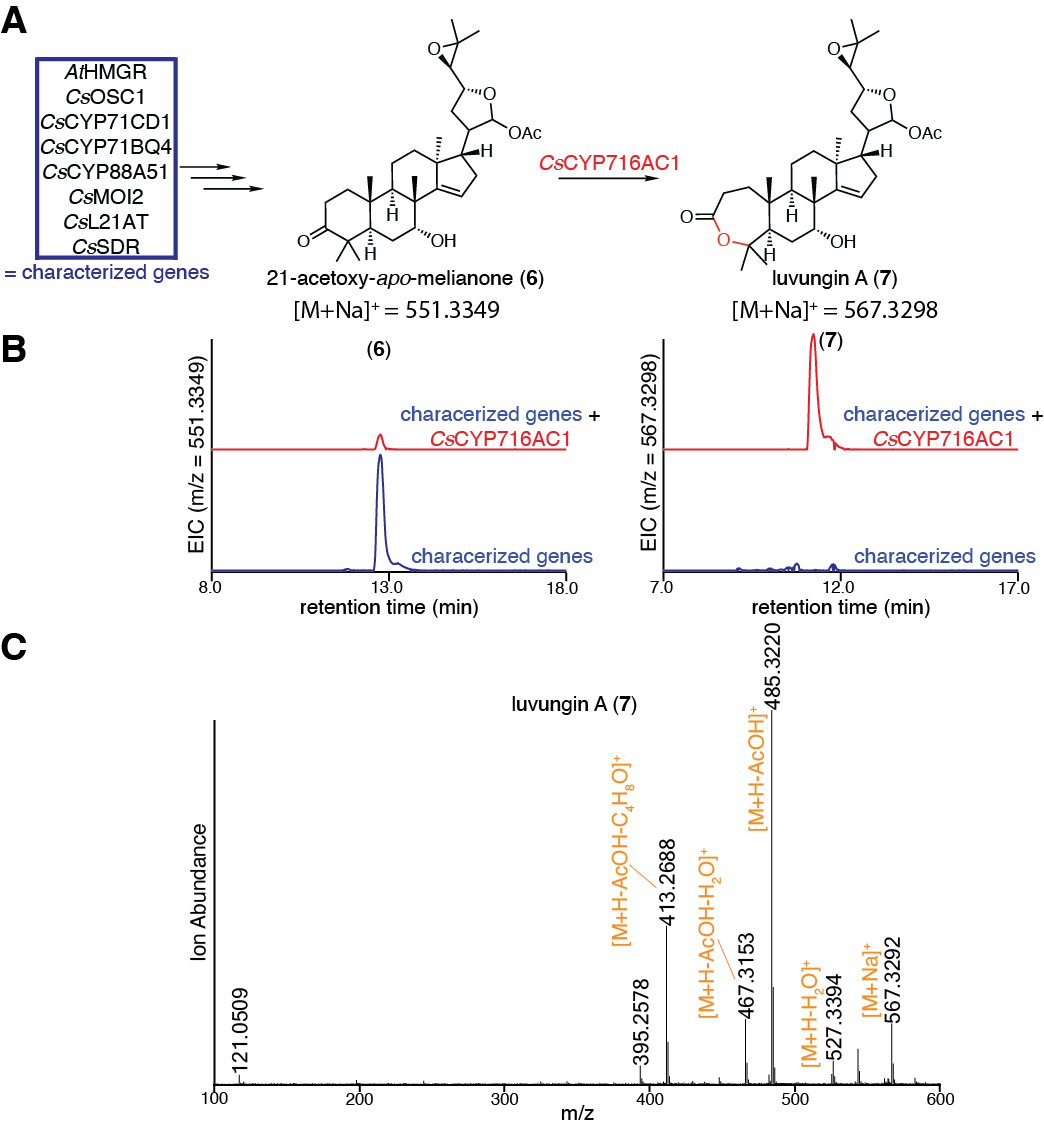

##### Fig. S17. Characterization of *Cs*CYP716AC1.

(A) Predicted function of *Cs*CYP716AC1 in converting 21-acetoxyl-*apo*-melianone (**6**) to luvungin A (**7**). (B) Extracted ion chromatograms (EICs) for extracts of *N. benthamiana* agro-infiltrated with the characterized genes listed in panel A, either alone (blue) or with the addition of *Cs*CYP716AC1 (red). EICs are displayed for masses of 551.3349 (calculated mass for (**6**) [M+Na]^+^) or 567.3298 (calculated mass for (**7**) [M+Na]^+^). (C) Mass spectrum of (**7**) heterologously produced in *N. benthamiana* as shown in panel B. Note that [M+Na]^+^ doesn’t fragment well in MSMS and the parent peak [M+H]^+^ is too low to be useful for MSMS analysis. Representative EICs and mass spectra are displayed (n=6).

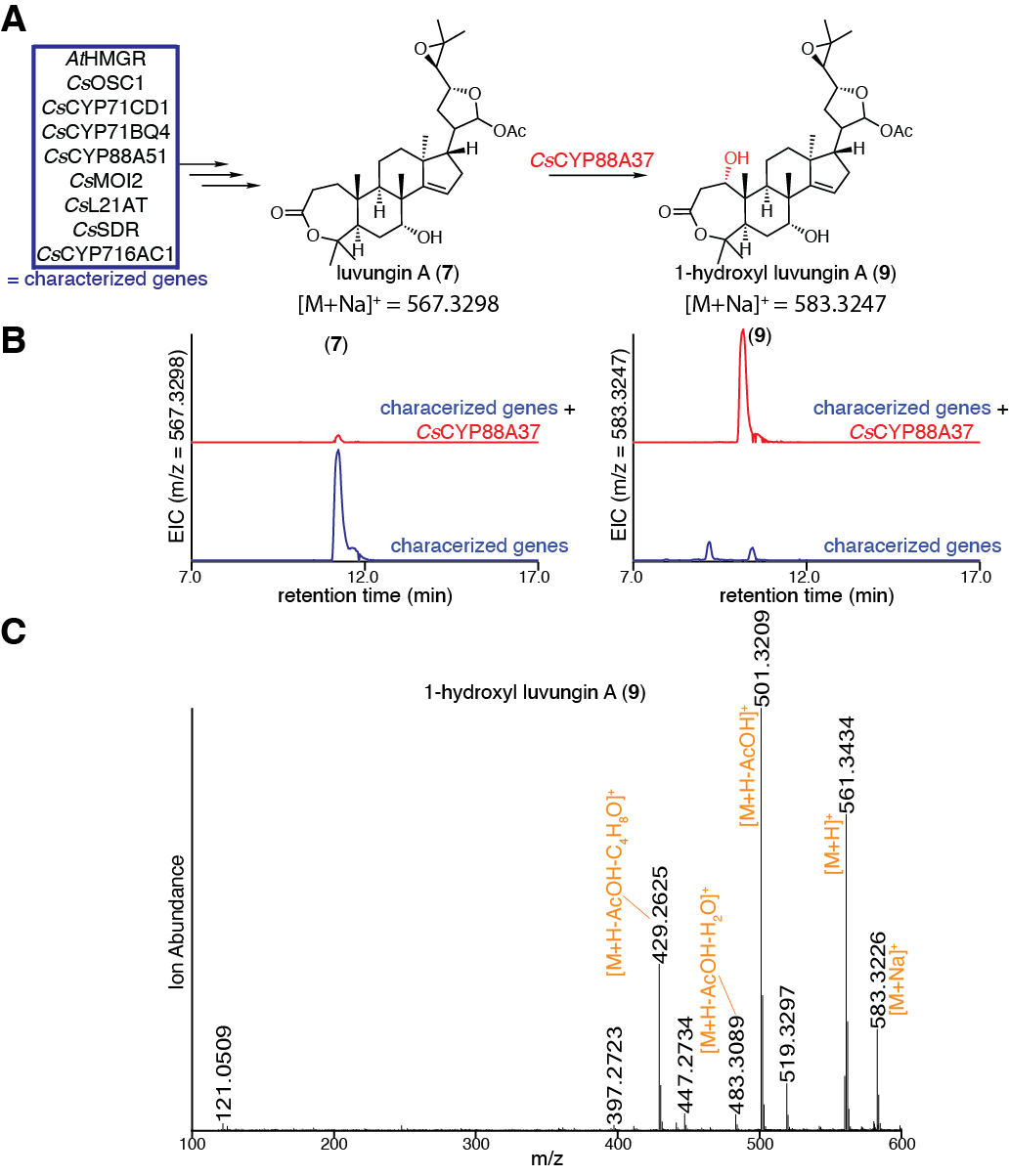

##### Fig. S18. Characterization of *Cs*CYP88A37.

(A) Predicted function of *Cs*CYP88A37 in converting luvungin A (**7**) to 1-hydroxyl luvungin A (**9**). (B) Extracted ion chromatograms (EICs) for extracts of *N. benthamiana* agro-infiltrated with the characterized genes listed in panel A, either alone (blue) or with the addition of *Cs*CYP88A37 (red). EICs are displayed for masses of 567.3298 (calculated mass for (**7**) [M+Na]^+^) or 583.3247 (calculated mass for (**9**) [M+Na]^+^). **(C)** Mass spectrum of (**9**) heterologously produced in *N. benthamiana* as shown in panel B. Representative EICs and mass spectra are displayed (n=6).

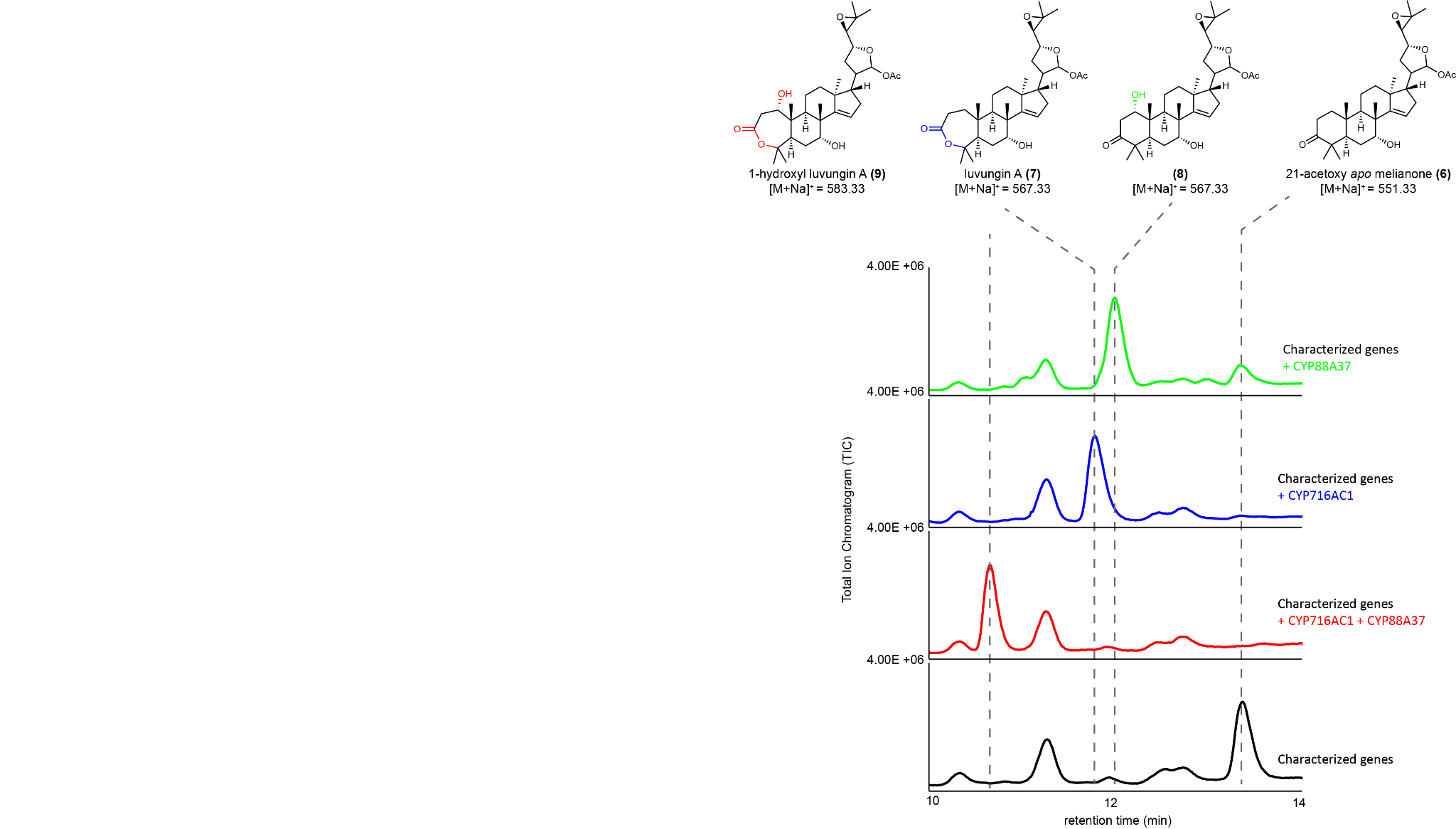

##### Fig. S19. Oxidation of 21-acetoxy-*apo*-melianone (6) by either *Cs*CYP88A37 or *Cs*CYP716AC1.

Predicted structures and representative total ion chromatograms (TICs) for extracts of *N. benthamiana* agro-infiltrated with characterized enzymes (*At*HMGR, *Cs*OSC1, *Cs*CYP71CD1, *Cs*CYP71BQ4, *Cs*CYP88A51, *Cs*MOI2, *Cs*L21AT and *Cs*SDR (black)) in combination with *Cs*CYP88A37 and *Cs*CYP716AC1, either alone (green and blue, respectively), or in together (red). Representative TICs displayed for the experiments (n=3).

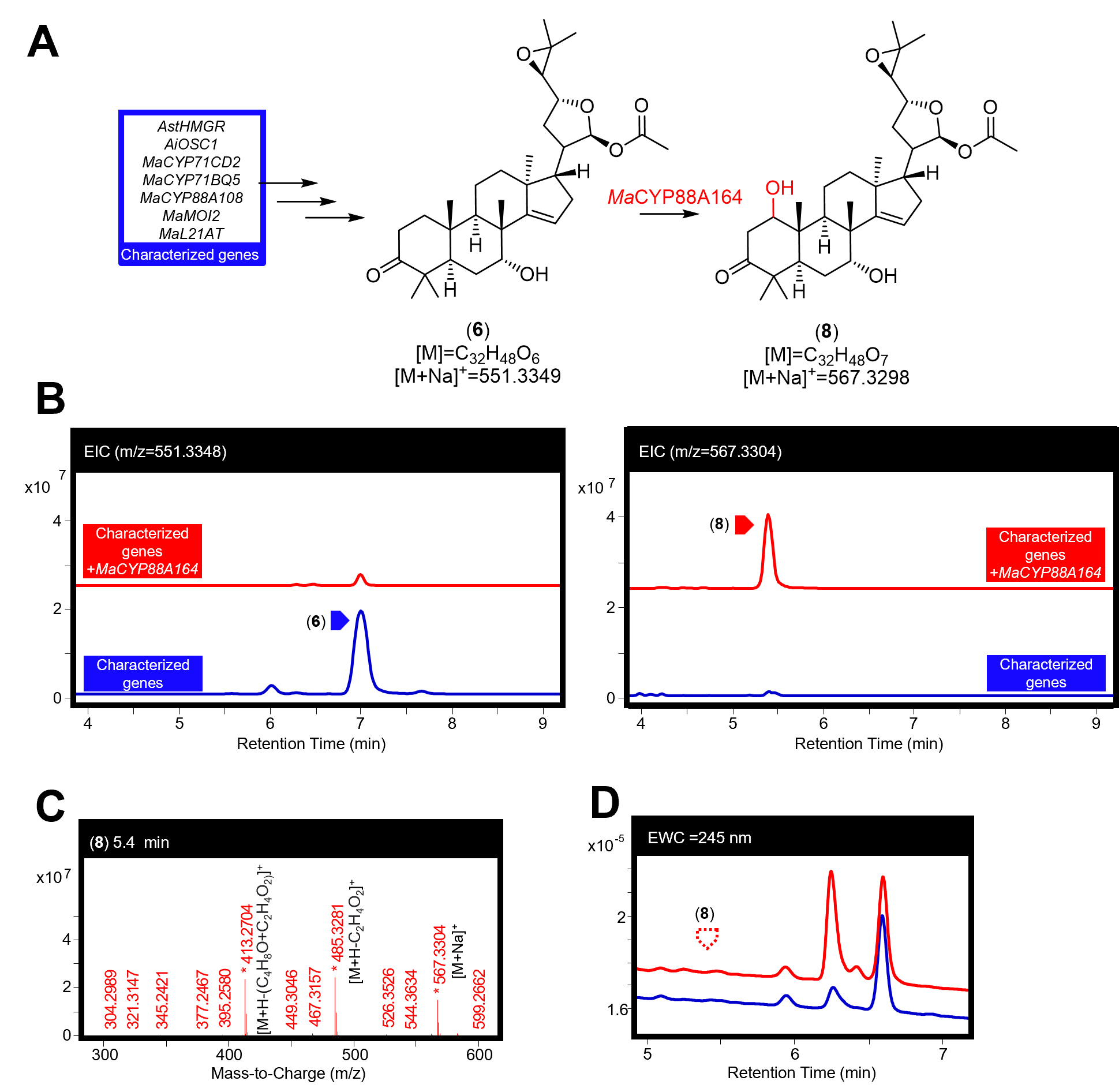

##### Fig. S20. Characterisation of *Ma*CYP88A164.

(A) Function of MaCYP88A164 in producing a 1-hydroxyl-21(*S*)-acetoxyl-*apo*-melianone (**8**) (confirmed by NMR of later products (Table S11, Table S12)) from 21(*S*)-acetoxyl-*apo*-melianone (**6**). (B) Extracted ion chromatograms (EICs) for extracts of *N. benthamiana* agro-infiltrated with the characterized enzymes listed in panel A, either alone (blue) or with the addition of *Ma*CYP88A164 (red). The EICs display masses of 551.3348 (observed mass for [(**6**)+Na]^+^) and 567.3304 (observed mass for [(**8**)+Na]^+^). (C) Mass spectra for (**8**) being heterologously produced in *N. benthamiana,* the main observed adduct ([M+Na]^+^) and fragments (including loss of acetic acid [M+H-C_2_H_4_O_2_]^+^ and loss of the four-carbon epoxide containing fragment and an acetic acid [M+H-(C_4_H_8_O+C_2_H_4_O_2_)]^+^) are labeled. (D) Extracted wavelength chromatograms (EWCs) of 245 nm (width of 4 nm) for extracts displayed in panel B. Due to the lack of an enone system in (**8**) no UV peak is observed. Representative traces and spectra are displayed (n=6).

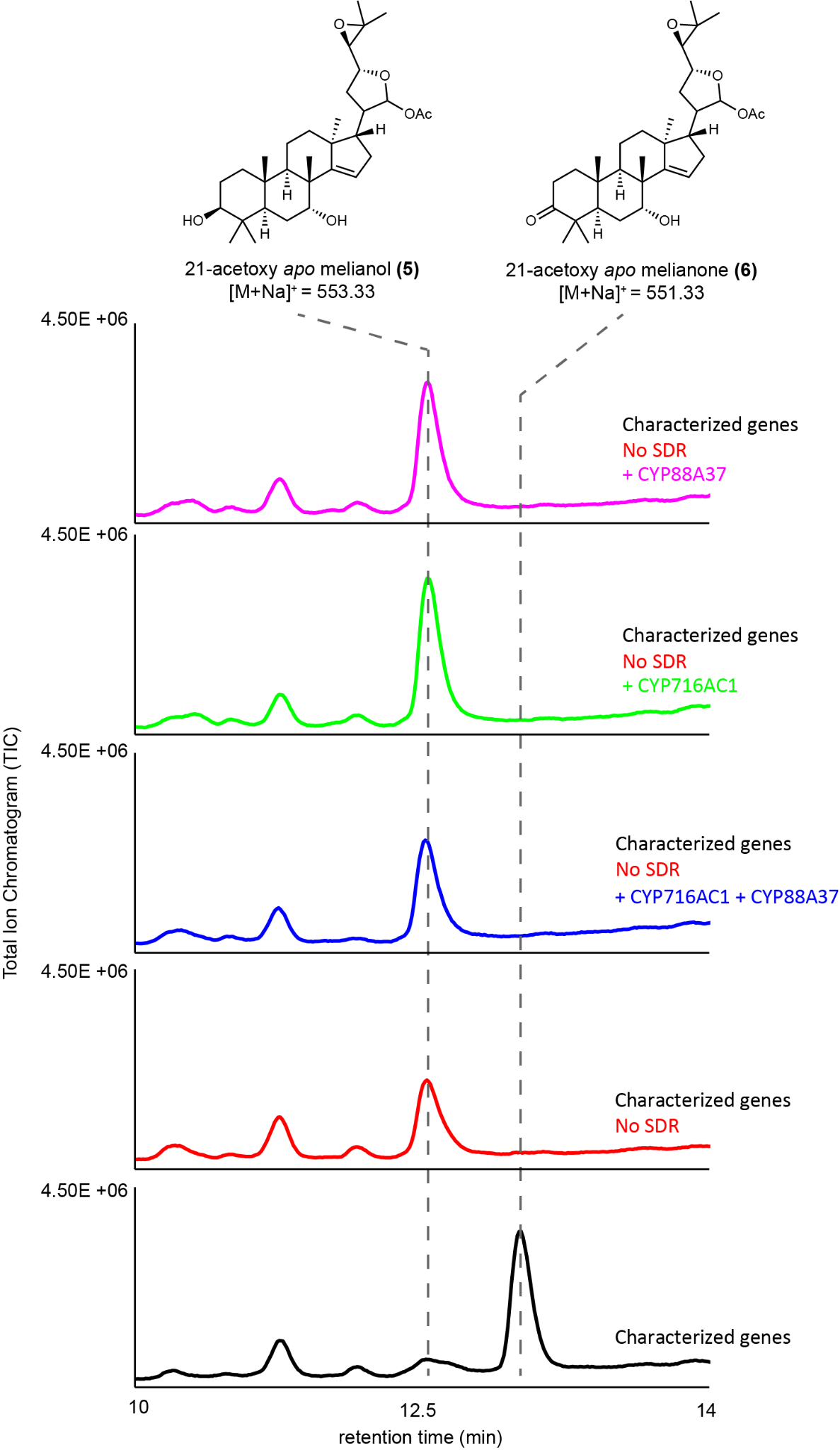

##### Fig. S21. **Oxidation by *Cs*CYP88A37 or *Cs*CYP716AC1 requires *Cs*SDR.**

Predicted structures and representative total ion chromatograms (TICs) for extracts of *N. benthamiana* agro-infiltrated with characterized enzymes (*At*HMGR, *Cs*OSC1, *Cs*CYP71CD1, *Cs*CYP71BQ4, *Cs*CYP88A51, *Cs*MOI2, and *Cs*L21AT) in combination with *CsSDR* (black) or without *Cs*SDR (red) but with the addition of *Cs*CYP88A37 or *Cs*CYP716AC1, either alone (pink and green, respectively) or together (blue). Representative TICs are displayed (n=3).

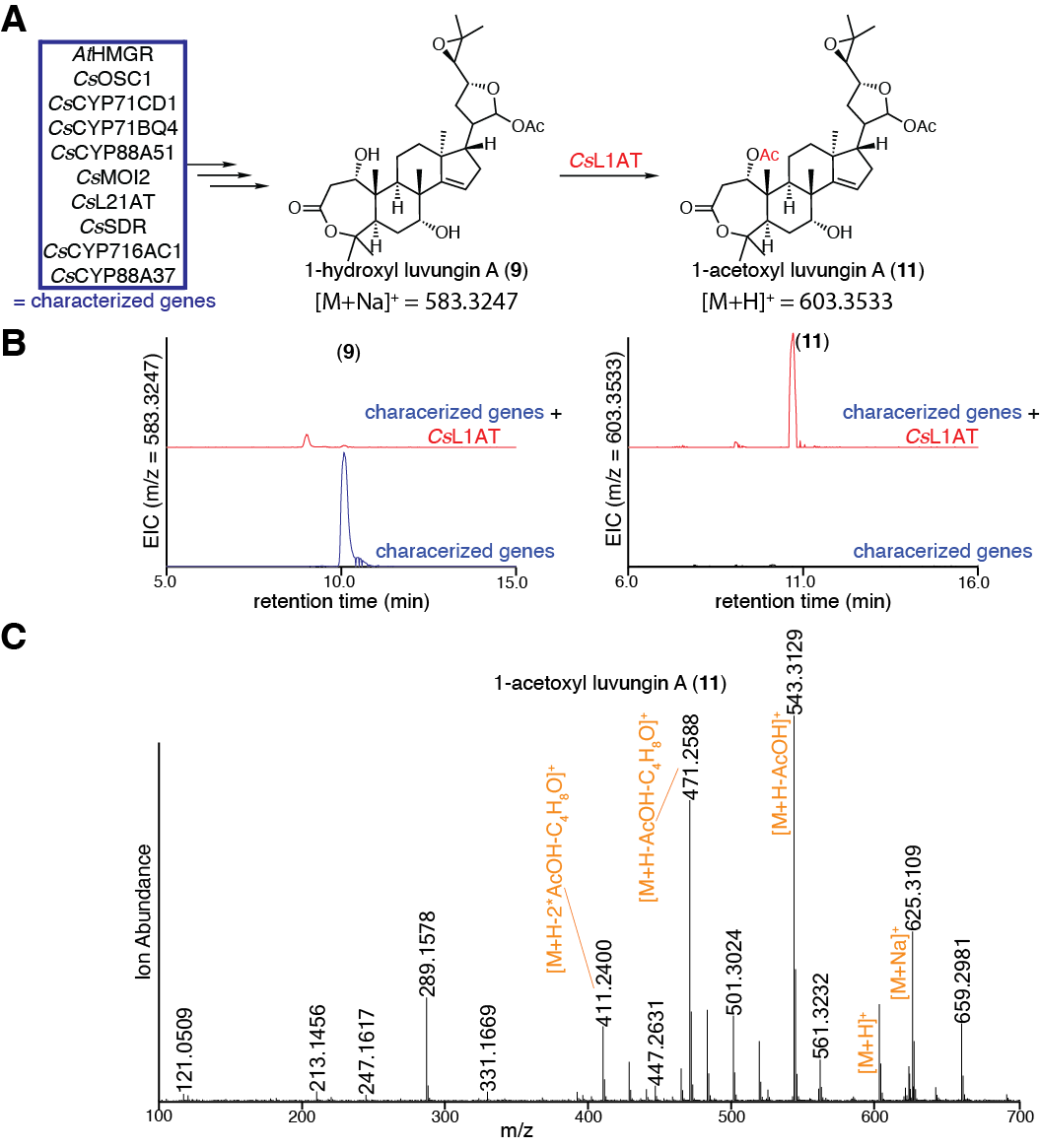

##### Fig. S22. Characterization of *Cs*L1AT.

(A) Predicted function of *Cs*L1AT in converting 1-hydroxyl luvungin A (**9**) to 1-acetoxyl luvungin A (**11**). (B) Extracted ion chromatograms (EICs) for extracts of *N. benthamiana* agro-infiltrated with the characterized genes listed in panel A, either alone (blue) or with the addition of *Cs*L1AT (red). EICs are displayed for masses of 583.3247 (calculated mass for (**9**) [M+Na]^+^) or 603.3533 (calculated mass for (**11**) [M+H]^+^). **(C)** Mass spectrum of (**11**) being heterologously produced in *N. benthamiana* as shown in panel B. Representative EICs and mass spectra are displayed (n=6).

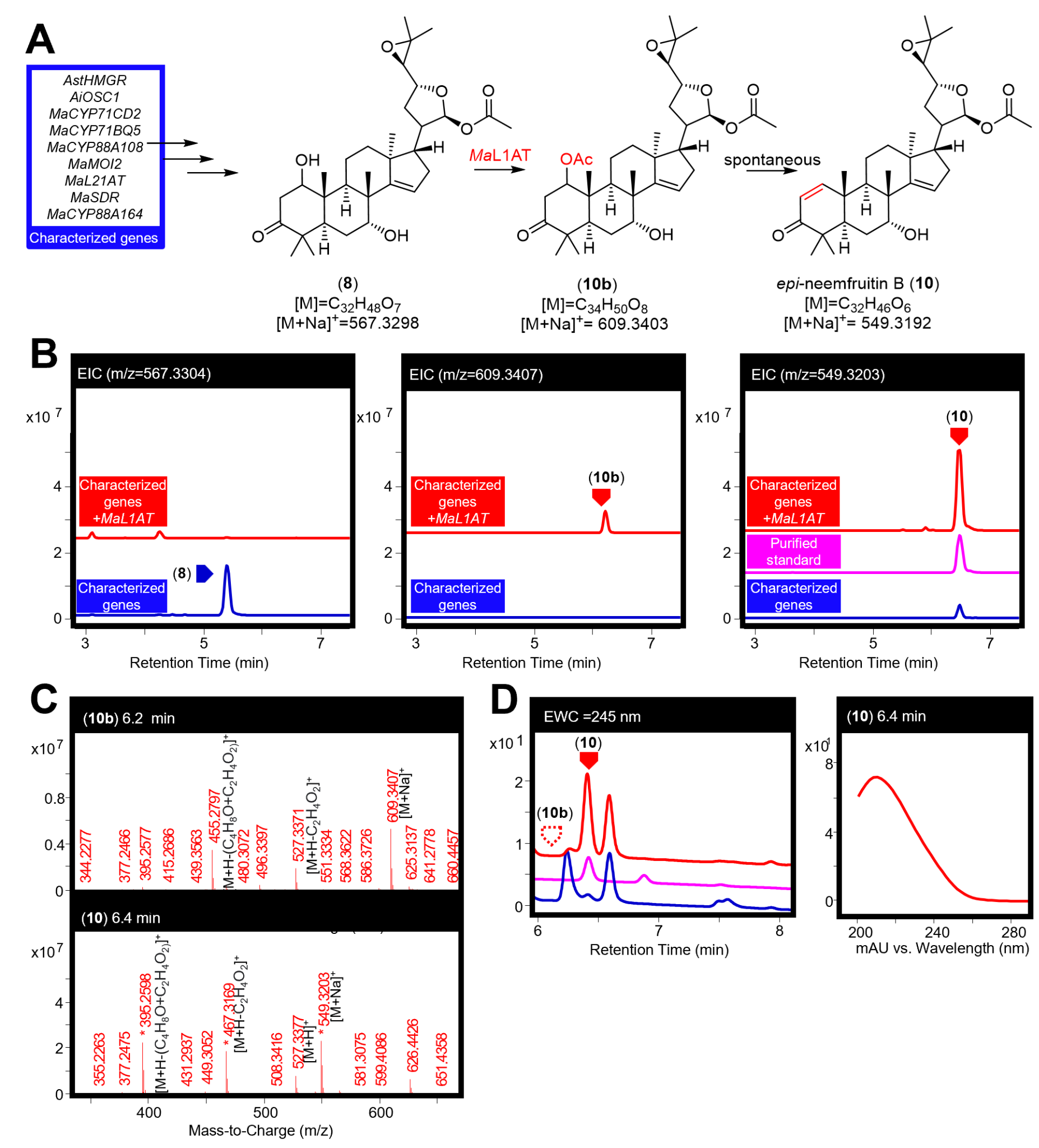

##### Fig. S23. Characterisation of *M*aL1AT.

(A) Function of *Ma*L1AT in producing *epi*-neemfruitin B (**10**) (confirmed by NMR, Table S11 to S12) as a major product, along with 1,21-di-acetoxyl-*apo*-melianone (**10b**), from 1-hydroxyl-21(*S*)-acetoxyl-*apo*-melianone (**8**). (B) Extracted ion chromatograms (EICs) for extracts of *N. benthamiana* agro-infiltrated with the characterized genes listed in panel A, either alone (blue), with *Ma*L1AT (red) or for a purified standard of (**10**) (pink). The EICs display masses of 567.3304 (observed mass for [(**8**)+Na]^+^), 609.3407 (observed mass for [(**10b**)+Na]^+^) and 549.3203 (observed mass [(**10**)+Na]^+^). (C) Mass spectra for (**10b**) and (**10**) being heterologously produced in *N. benthamiana,* the main observed adducts ([M+Na]^+^ and [M+H]^+^) and fragments (including loss of acetic acid [M+H-C_2_H_4_O_2_]^+^ and loss the four-carbon epoxide containing fragment and an acetic acid [M+H-(C_4_H_8_O+C_2_H_4_O_2_)]^+^) are labeled. (D) Extracted wavelength chromatograms (EWCs) of 245 nm (width of 4nm) for extracts displayed in panel B. Due to the lack of an enone system in (**10b**) no UV peak is observed. The UV spectra (mAU) of (**10**) is given. Standards have been scaled. Representative traces and spectra are displayed (n=6).

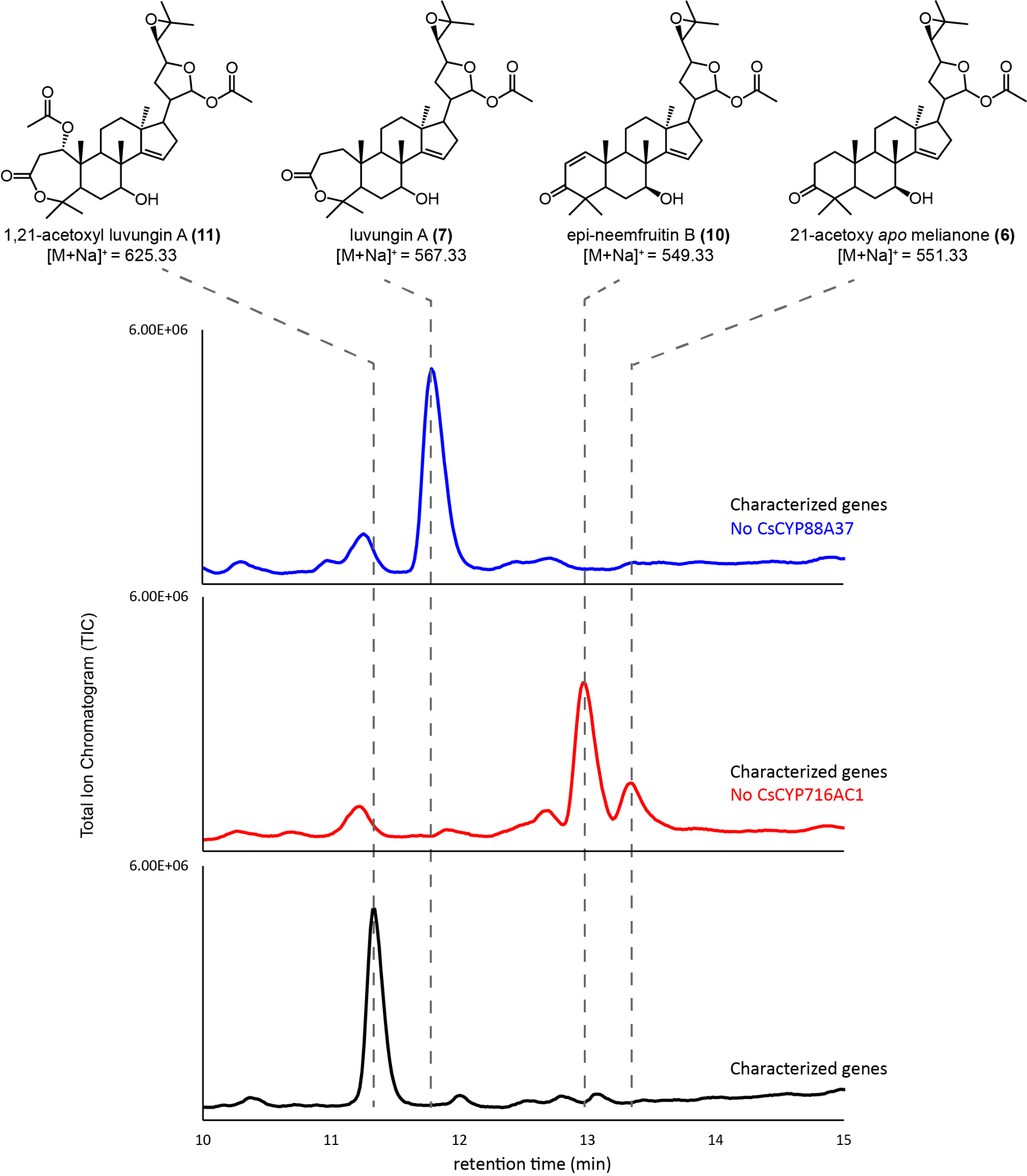

##### Fig. S24. Characterization of *Cs*L1AT in the absence of *Cs*CYP716AC1 or *Cs*CYP88A37.

Predicted structures and representative total ion chromatograms (TICs) for extracts of *N. benthamiana* agro-infiltrated with the following enzymes: *At*HMGR, *Cs*OSC1, *Cs*CYP71CD1, *Cs*CYP71BQ4, *Cs*CYP88A51, *Cs*MOI2, *Cs*L21AT, *Cs*SDR, *Cs*CYP716AC1, *Cs*CYP88A37, and *Cs*L21AT (black). Traces for the same combination of enzymes lacking either *Cs*CYP88A37 (blue) or *Cs*CYP716AC1 (red) are also shown. Representative TICs are displayed (n=3).

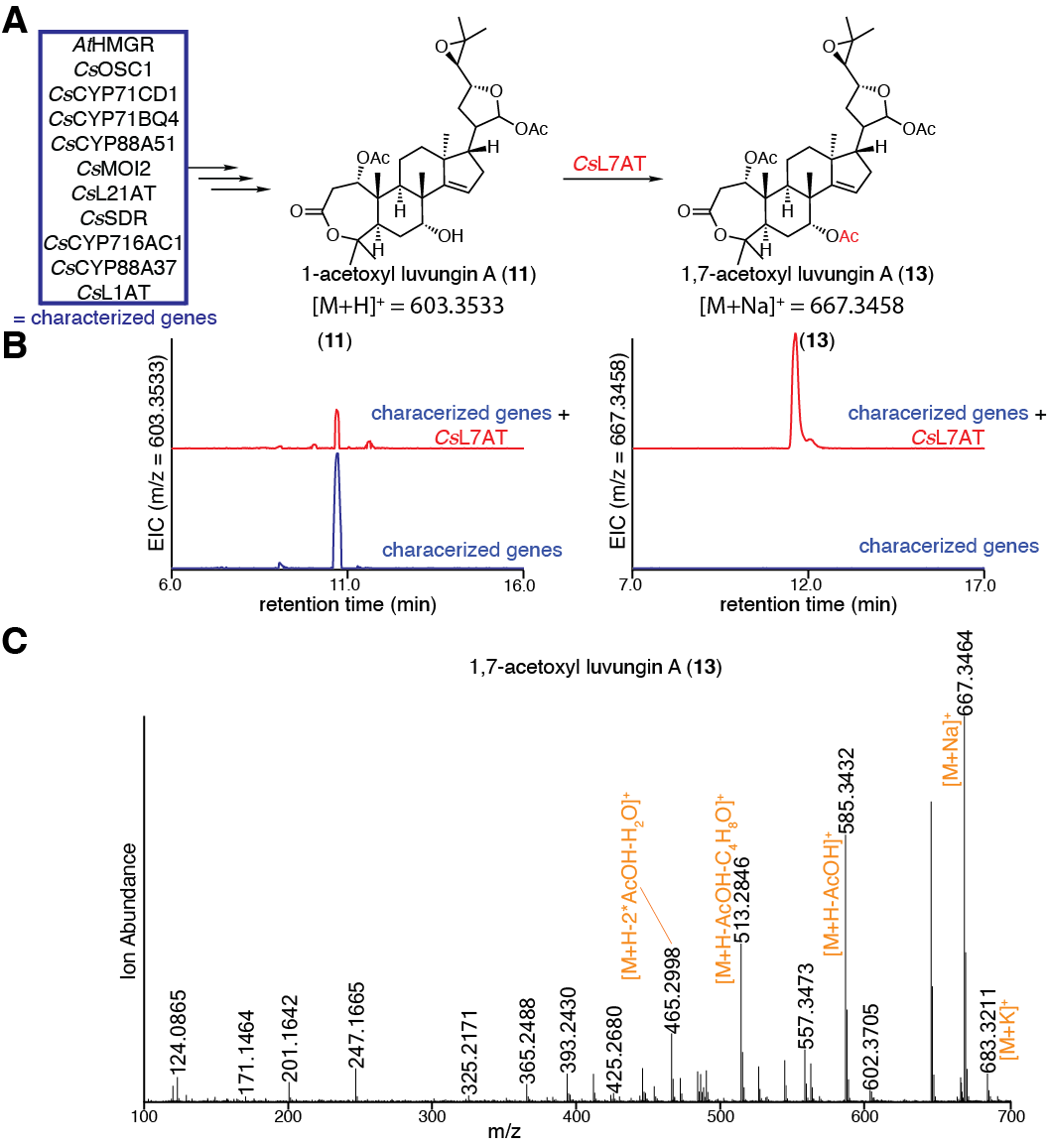

##### Fig. S25. Characterization of *Cs*L7AT.

(A) Predicted function of *Cs*L7AT in converting 1-acetoxyl luvungin A (**11**) to 1,7-acetoxyl luvungin A (**13**). (B) Extracted ion chromatograms (EICs) for extracts of *N. benthamiana* agro-infiltrated with the characterized genes listed in panel A, either alone (blue) or with the addition of *Cs*L7AT (red). EICs are displayed for masses of 603.3533 (calculated mass for (**11**) [M+H]^+^) or 667.3458 (calculated mass for (**13**) [M+Na]^+^). (C) Mass spectrum of (**13**) heterologously produced in *N. benthamiana* as shown in panel B. Representative EICs and mass spectra are displayed (n=6).

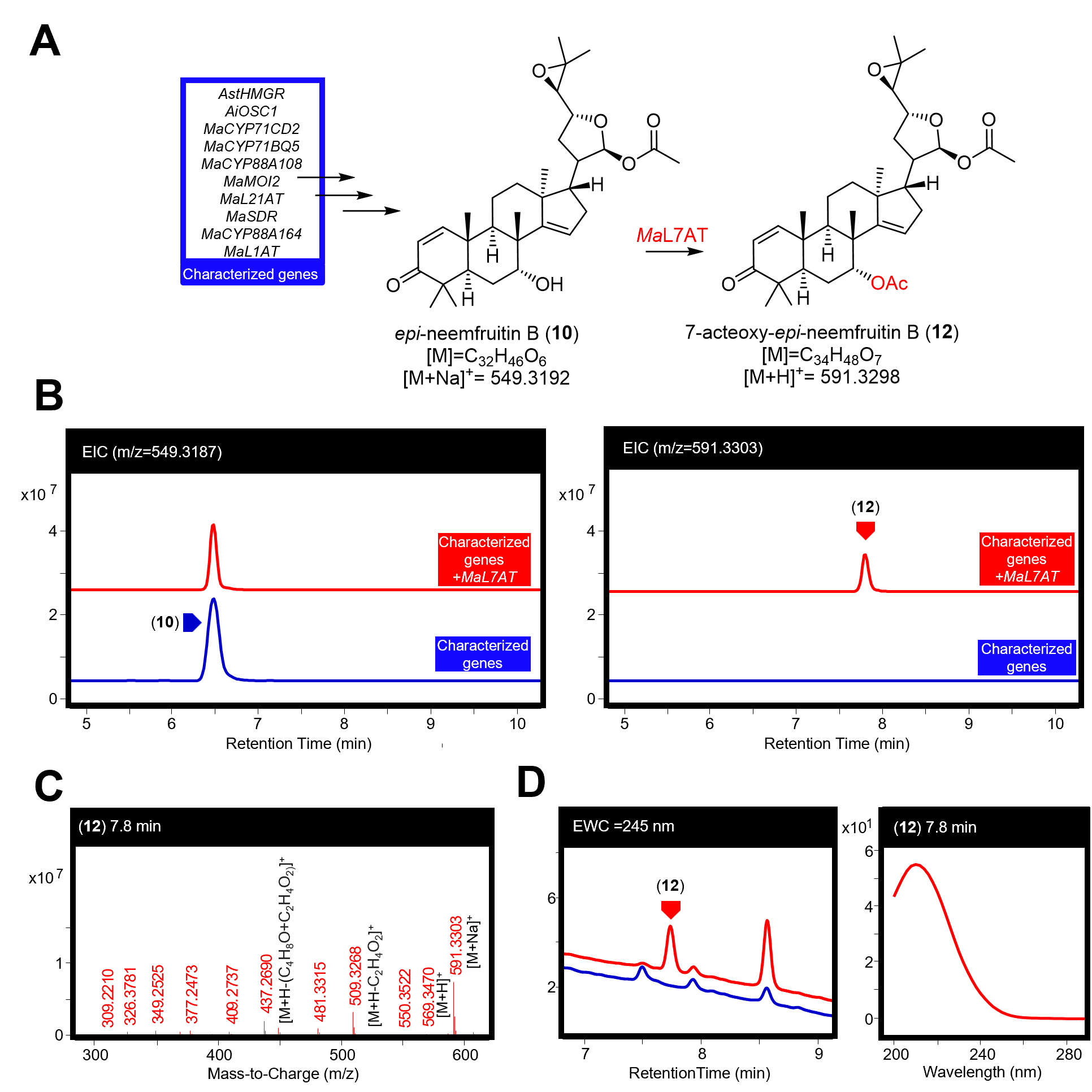

##### Fig. S26. Characterisation of *Ma*L7AT.

(A) Function of *Ma*L7AT in producing a 7-acetoxyl-*epi*-neemfruitin B (**12**) (position confirmed by NMR of later product (**14**) Table S13) from *epi*-neemfruitin B (**10**). (B) Extracted ion chromatograms (EICs) for extracts of *N. benthamiana* agro-infiltrated with the characterized genes listed in panel A, either alone (blue) or with the addition of *MaL7AT* (red). The EICs display masses of 549.3187 (observed mass for [(**10**)+Na]^+^) and 591.3303 (observed mass for [(**12**)+Na]^+^). (C) Mass spectra for (**12**) being heterologously produced in *N. benthamiana,* the main observed adducts ([M+H]^+^ and [M+Na]^+^) and fragments (including loss of acetic acid [M+H-C_2_H_4_O_2_]^+^ or loss the four-carbon epoxide containing fragment and an acetic acid [M+H-(C_4_H_8_O+C_2_H_4_O_2_)]^+^) are labeled. (D) Extracted wavelength chromatograms (EWCs) of 245 nm (width of 4nm) for extracts displayed in panel B. UV spectra (mAU) of (**12**) being heterologously produced in *N. benthamiana* is also given*.* Representative traces and spectra are displayed (n=6).

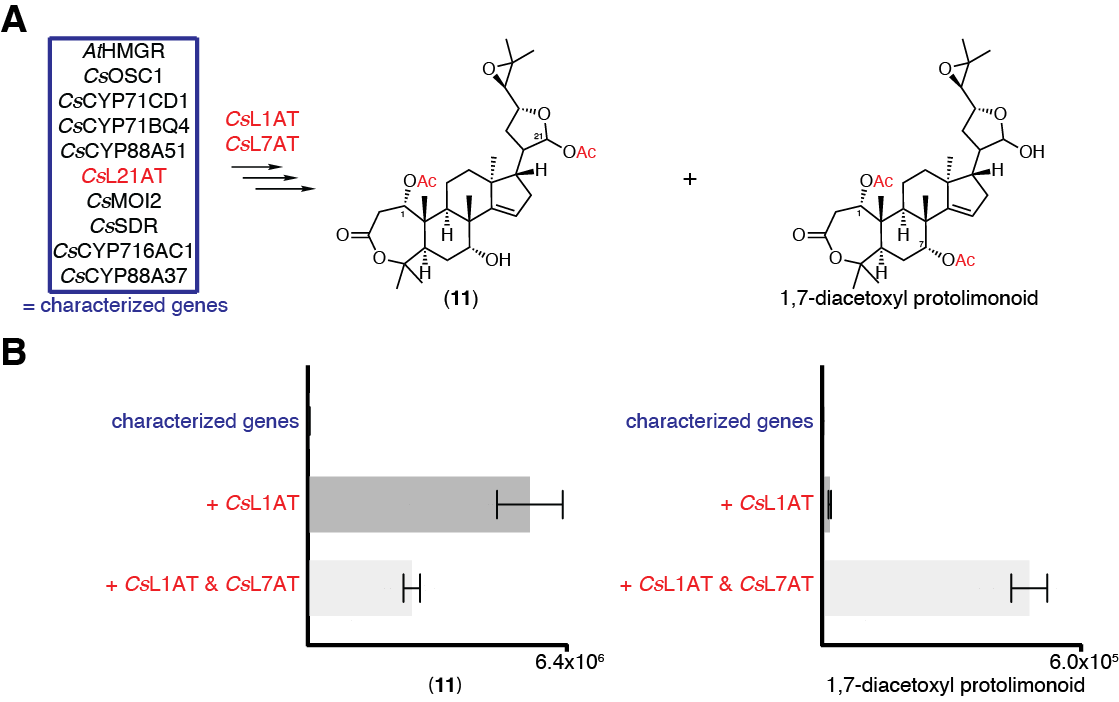

##### Fig. S27. Accumulation of 1,21-diacetoxyl (11) and 1,7-diacetoxyl protolimonoid intermediates with the introduction of *Cs*L1AT and *Cs*L7AT.

(A) Structures of the two predicted diacetoxyl protolimonoids ((**11**) and a 1,7-diacetoxyl protolimonoid), which are produced when all biosynthetic enzymes for the production of (**13**) (enzymes in the characterized genes box with the addition of *Cs*L1AT and *Cs*L7AT) are co-expressed. (B) Integrated peak area of extracted ion chromatogram (EIC) showing the accumulation of (**11**) and of the 1,7-diacetoxyl protolimonoid in *N. benthamiana*. Accumulation is shown for the characterized genes listed in panel A (blue) with the addition of *Cs*L1AT alone or *Cs*L1AT and *Cs*L7AT (red). Values and error bars represent the mean and the standard error of the mean (n=6).

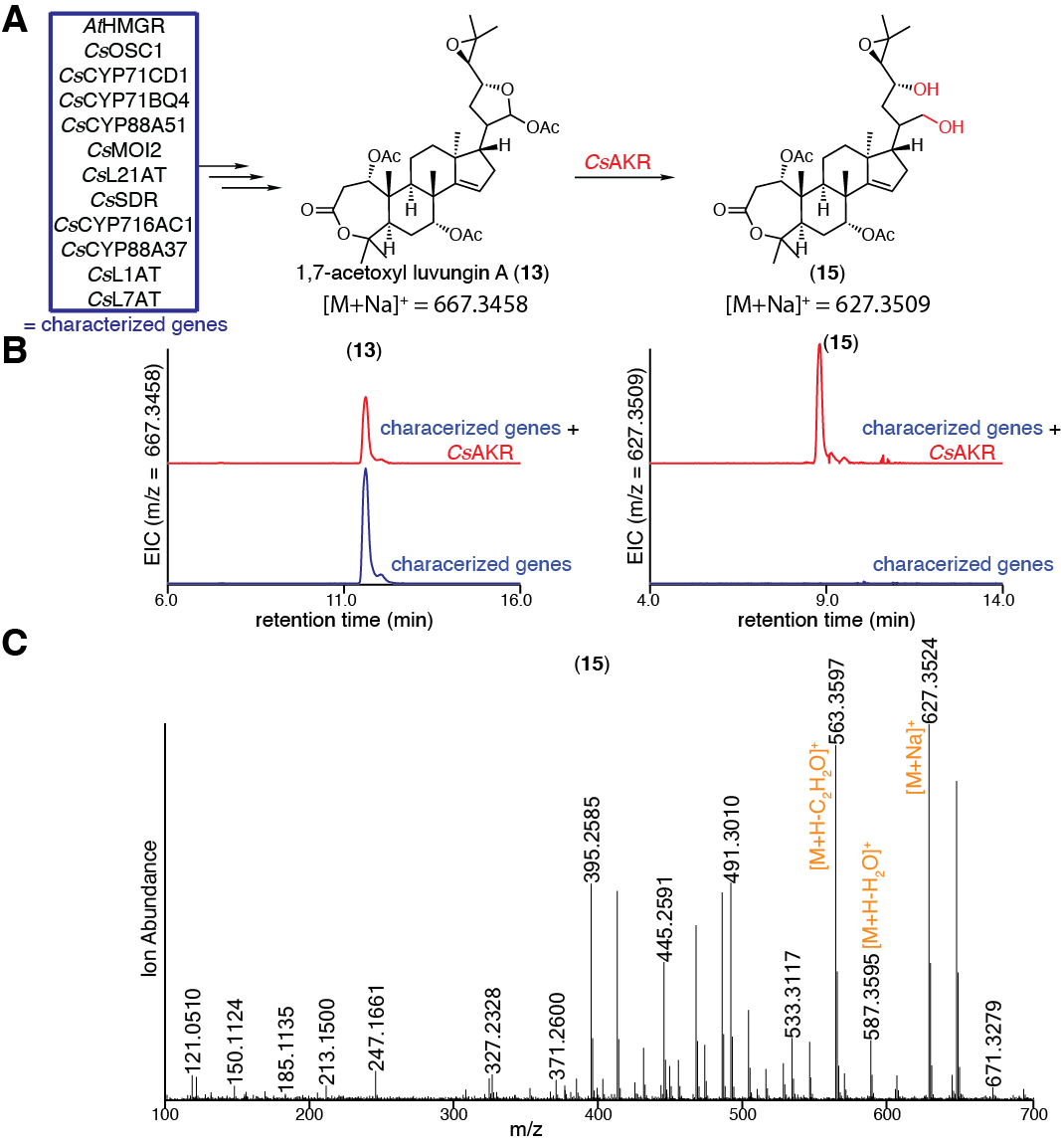

##### Fig. S28. Characterization of *Cs*AKR.

(A) Predicted function of *Cs*AKR in converting 1,7-acetoxyl luvungin A (**13**) to (**15**). (B) Extracted ion chromatograms (EICs) for extracts of *N. benthamiana* agro-infiltrated with the characterized genes listed in panel A, either alone (blue) or with the addition of *Cs*AKR (red). The EICs display masses of 667.3458 (calculated mass for (**13**) [M+Na]^+^) or 627.3509 (calculated mass for (**15**) [M+Na]^+^). (C) Mass spectrum of (**15**) heterologously produced in *N. benthamiana* as shown in panel B. Representative EICs and mass spectra are displayed (n=6).

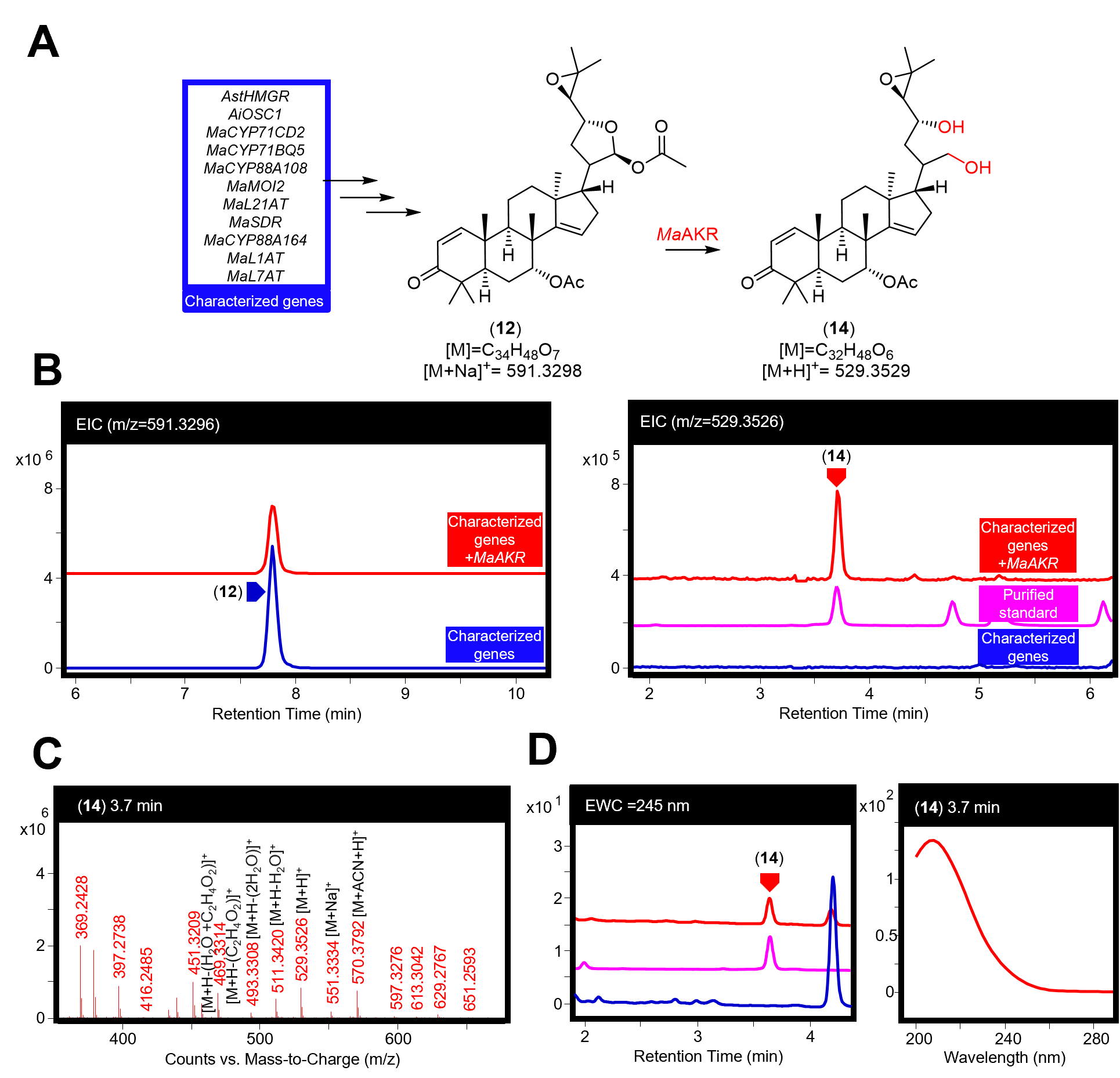

##### Fig. S29. Characterisation of *Ma*AKR.

(A) Function of *Ma*AKR in producing the 21,23 diol (**14**) (confirmed by NMR, Table S13) from 7-acetoxyl-*epi*-neemfruitin B (**12**), although the actual substrate for enzymatic transformation may be an earlier non-C21-acetylated intermediate. (B) Extracted ion chromatograms (EICs) for extracts of *N. benthamiana* agro-infiltrated with the characterized genes listed in panel A, either alone (blue), with the addition of *Ma*AKR (red) or for a purified standard of (**14**) (pink). The EICs display masses of 591.3296 (observed mass for [(**12**)+Na]^+^) and 529.3526 (observed mass of [(**14**)+H]^+^). (C) Mass spectra for (**14**) being heterologously produced in *N. benthamiana,* the main observed adducts ([M+H]^+^, [M+ACN+H]^+^and [M+Na]^+^) and fragments (including loss of one [M+H-H_2_O]^+^ or two [M+H-(2H_2_O)]^+^ water molecules, acetic acid [M+H-C_2_H_4_O_2_]^+^ or water and acetic acid [M+H-(H_2_O+C_2_H_4_O_2_)]^+^) are labeled. The fragments are consistent with the presence of a diol, rather than the precursor hemi-acetal ring, and loss of acetic acid can be explained by the loss of the C7 acetoxyl group. (D) Extracted wavelength chromatograms (EWCs) of 245 nm (width of 4 nm) for extracts displayed in panel B. UV spectra (mAU) of (**14**) being heterologously produced in *N. benthamiana* is also given. Traces of standards have been scaled. Representative traces and spectra are displayed (n=6).

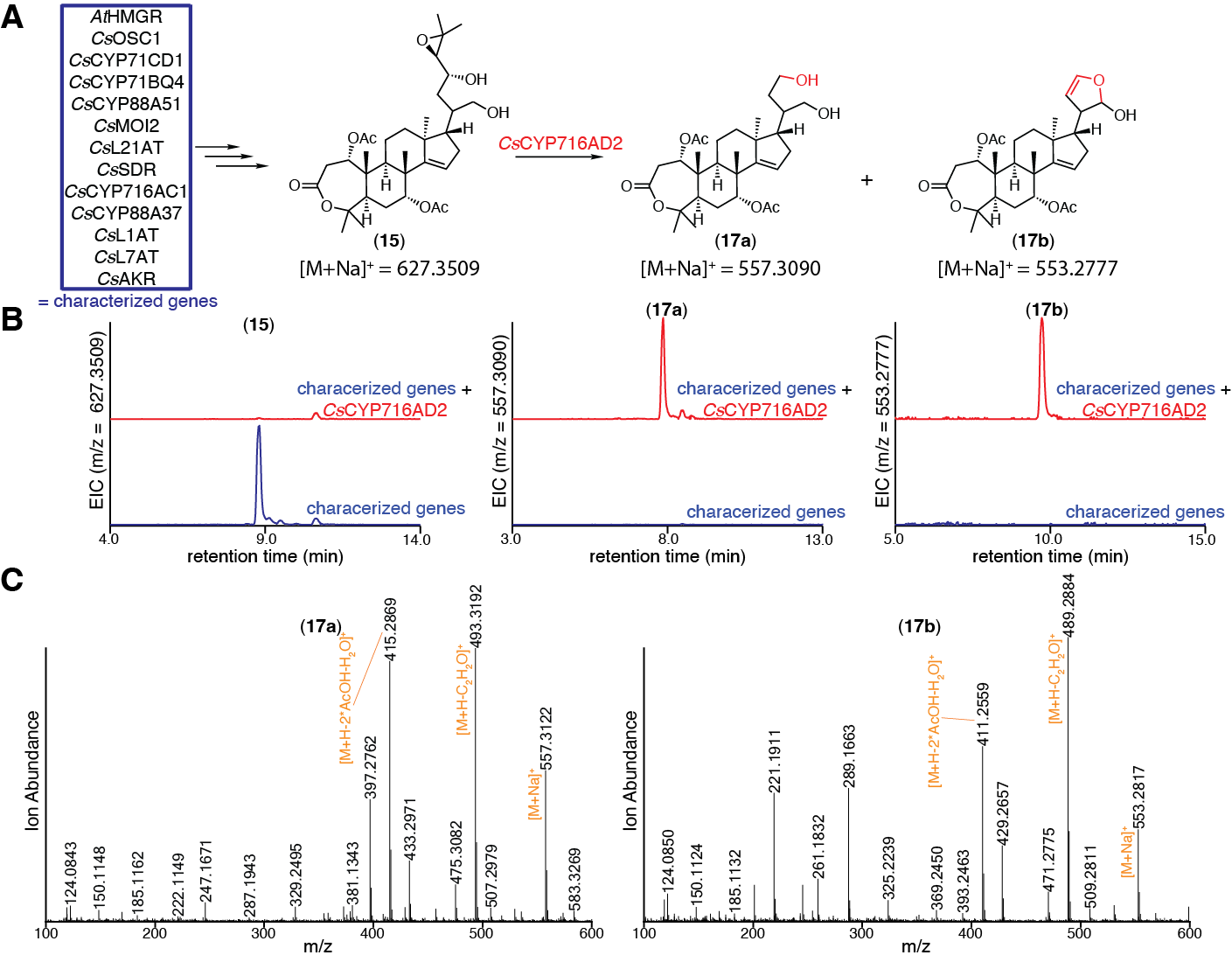

##### Fig. S30. Characterization of *Cs*CYP716AD2.

(A) Predicted function of *Cs*CYP716AD2 in converting (**15**) to (**17a**) and (**17b**). (B) Extracted ion chromatograms (EICs) for extracts of *N. benthamiana* agro-infiltrated with the characterized genes listed in panel A, either alone (blue) or with the addition of *Cs*CYP716AD2 (red). EICs are displayed for masses of 627.3509 (calculated mass for (**15**) [M+Na]^+^), 557.3090 (calculated mass for (**17a**) [M+Na]^+^) and 553.2777 (calculated mass for (**17b**) [M+Na]^+^). (C) Mass spectra of (**17a**) and (**17b**) heterologously produced in *N. benthamiana* are shown in panel B. Representative EICs and mass spectra are displayed (n=6).

##### Fig. S31. Characterization of *Ma*CYP716AD4.

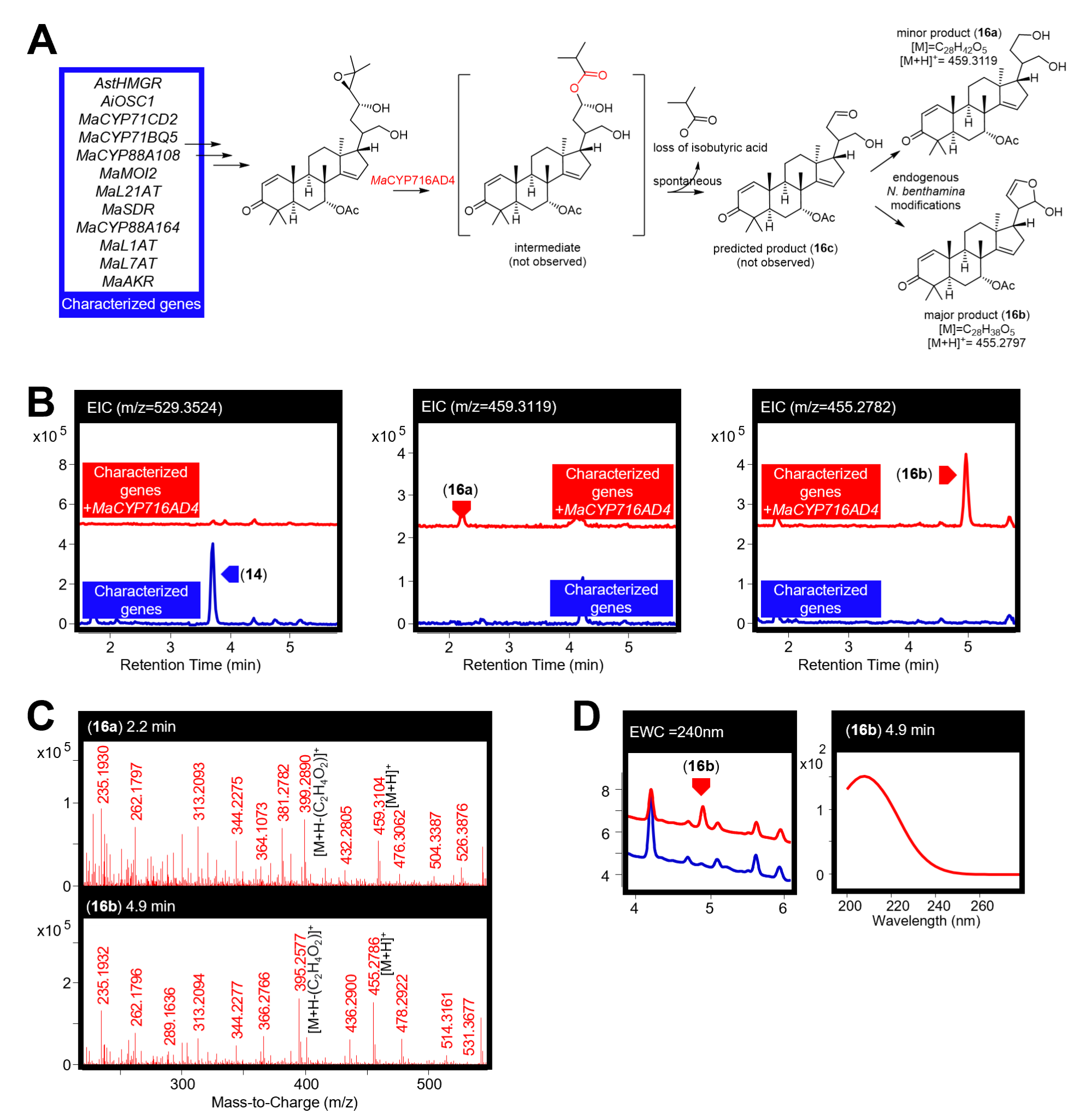

(A) Predicted function of *Ma*CYP716AD4 in converting (**14**) to (**16c**), with observation of products (**16a**) and (**16b**), full predicted mechanism is expanded upon in Fig. S32. (B) Extracted ion chromatograms (EICs) for extracts of *N. benthamiana* agro-infiltrated with the characterized genes listed in panel A, either alone (blue) or with the addition of *Ma*CYP716AD4 (red). EICs are displayed for masses of 529.3524 (observed mass of [(**14**)+H]^+^), 459.3119 (observed mass for [(**16a**)+H]^+^) and of 455.2782 (observed mass for [(**16b**)+H]^+^) . (C) Mass spectra for (**16a**) and (**16b**) being heterologously produced in *N. benthamiana,* the main observed adduct ([M+H]^+^) and fragment (loss of acetic acid [M+H-C_2_H_4_O_2_]^+^, consistent with the loss of C7 acetoxy group) are labeled. (D) Extracted wavelength chromatograms (EWCs) of 245 nm (width of 4 nm) for extracts displayed in panel B. UV spectra (mAU) of (**16b**) being heterologously produced in *N. benthamiana* is also given. No peak was observed for (**16a**), thought to be because of low accumulation. Representative traces and spectra are given (n=6).

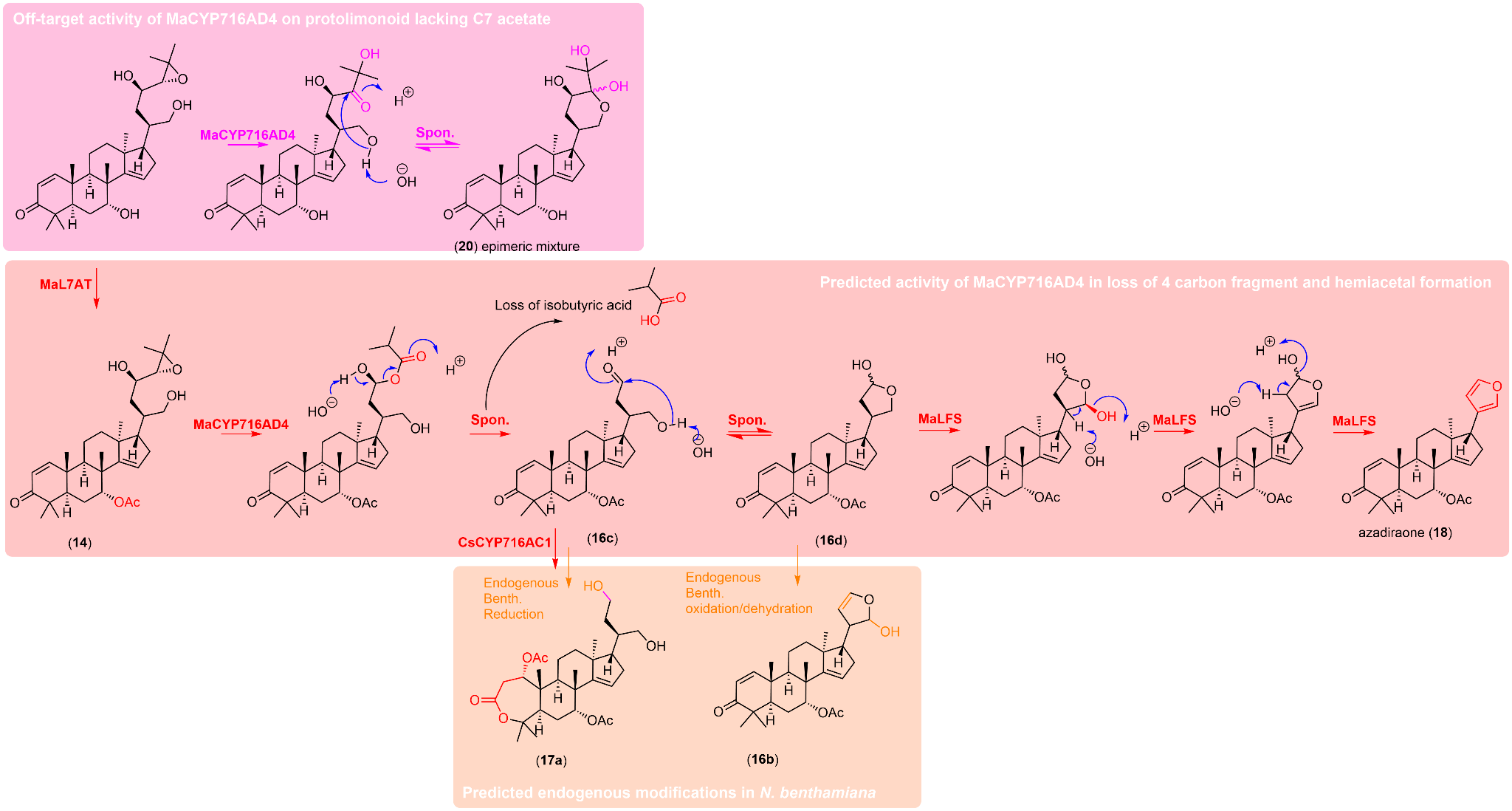

##### Fig. S32. Hypothetical reaction scheme for the action of CYP716ADs and LFSs via Baeyer-Villiger type route.

*Ma*CYP716AD4 and *Cs*CYP716AD2 are speculated to act via a Baeyer-Villiger type route. This would involve the enzymes converting the epoxide of their respective substrates to a ketone at C24 prior to introduction of an ester. The resulting products may then be spontaneously cleaved, with loss of isobutyric acid, resulting in the formation of a 5-membered hemi-acetal ring (**16d**) or (**17d**) which are the true substates for *Ma*LFS and *Cs*LFS, respectively. Evidence to support this proposal comes from a six-membered hemiacetal product of *Ma*CYP716AD4 that is produced only in the absence of acetylation of the C7-hydroxyl group (**20**) (Fig. S39). This product was purified (following large-scale expression in *N. benthamiana* of all genes for (**16**), excluding *Ma*L7AT), and shown by NMR to be an epimeric mixture of (**20**) (Table S16, Fig. S32). Although this exact product has not been isolated from nature before, protolimonoids with similar E-rings have been (*76*). This 6-membered hemiacetal product implies the prior conversion of the epoxide to a C24 ketone by *Ma*CYP716AD4, consistent with that required for the proposed Baeyer-Villiger-type oxidation to take place when the enzyme is presented with the optimal acetylated substate. It is proposed that in the absence of acetylation, C25 hydroxylation is observed instead of the Baeyer-Villiger-type oxidation, perhaps due to different substrate positioning in the active site. The above figure outlines the proposed reactions detailed above, to explain the occurrence of side-product (**20**) (Table S16) when *Ma*CYP716AD4 is expressed in the absence of *Ma*L7AT and the occurrence of proposed observed major products of *Cs*CYP716AD2 and *Ma*CYP716AD4 ((**17a**) and (**16b**) respectively) when expressed in the presence of *Cs*L7AT or *Ma*L7AT. Proposed on target reactions are indicated in red, off-target enzyme activity in pink and suspected endogenous modifications *in planta* by native *N. benthamiana* enzymes in orange.

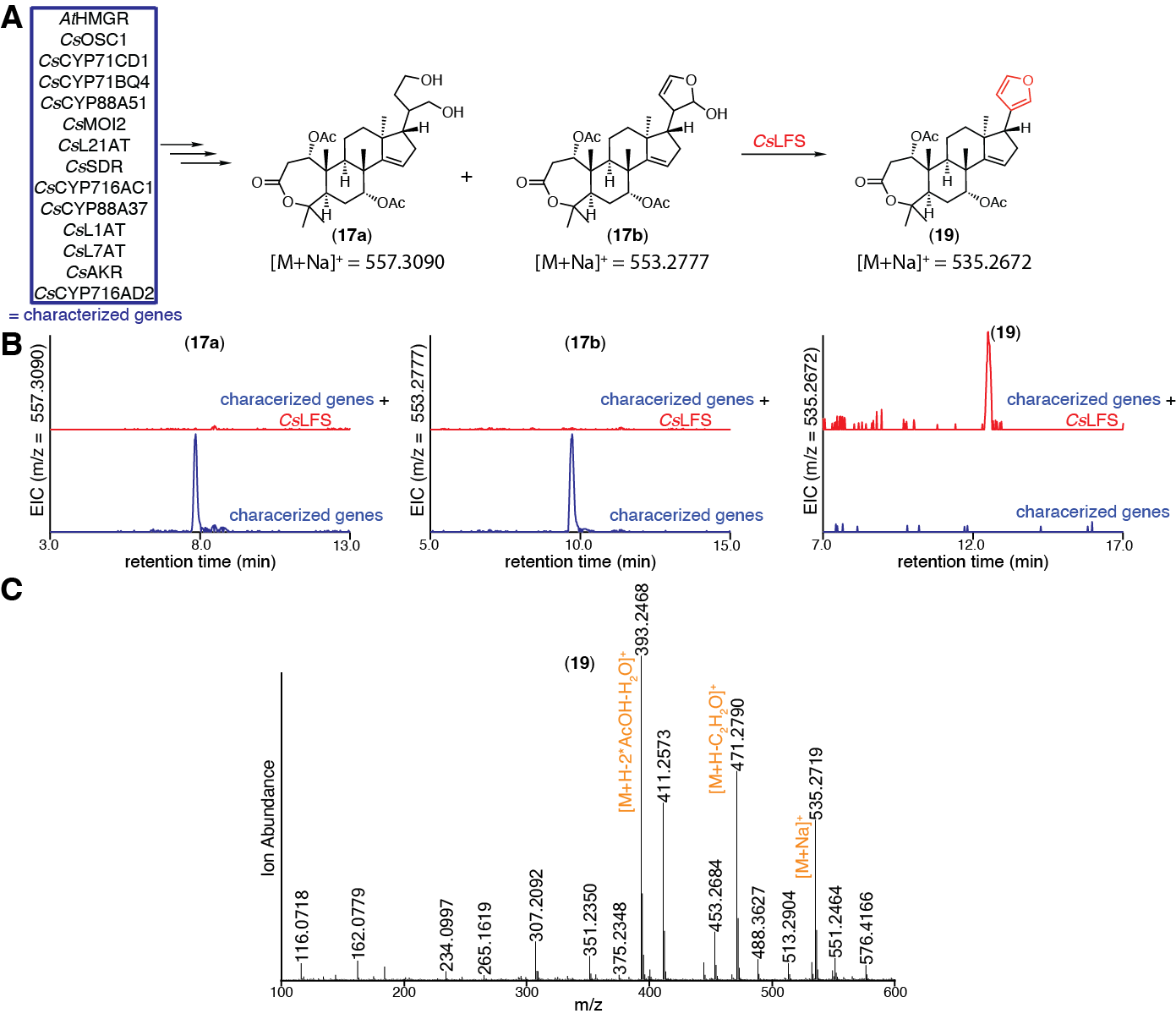

##### Fig. S33. Characterization of *Cs*LFS.

(A) Predicted function of *Cs*LFS in converting (**17a**) and (**17b**) to (**19**). (B) EICs for extracts of *N. benthamiana* agro-infiltrated with the characterized genes listed in panel A, either alone (blue) or with the addition of *Cs*LFS (red). The EICs display masses of 557.3090 (calculated mass for (**17a**) [M+Na]^+^), 553.2777 (calculated mass for (**17b**) [M+Na]^+^), or 535.2672 (calculated mass for (**19**) [M+Na]^+^). (C) Mass spectrum of (**19**) heterologously produced in *N. benthamiana* as shown in panel B. Representative EICs and mass spectra are displayed (n=6).

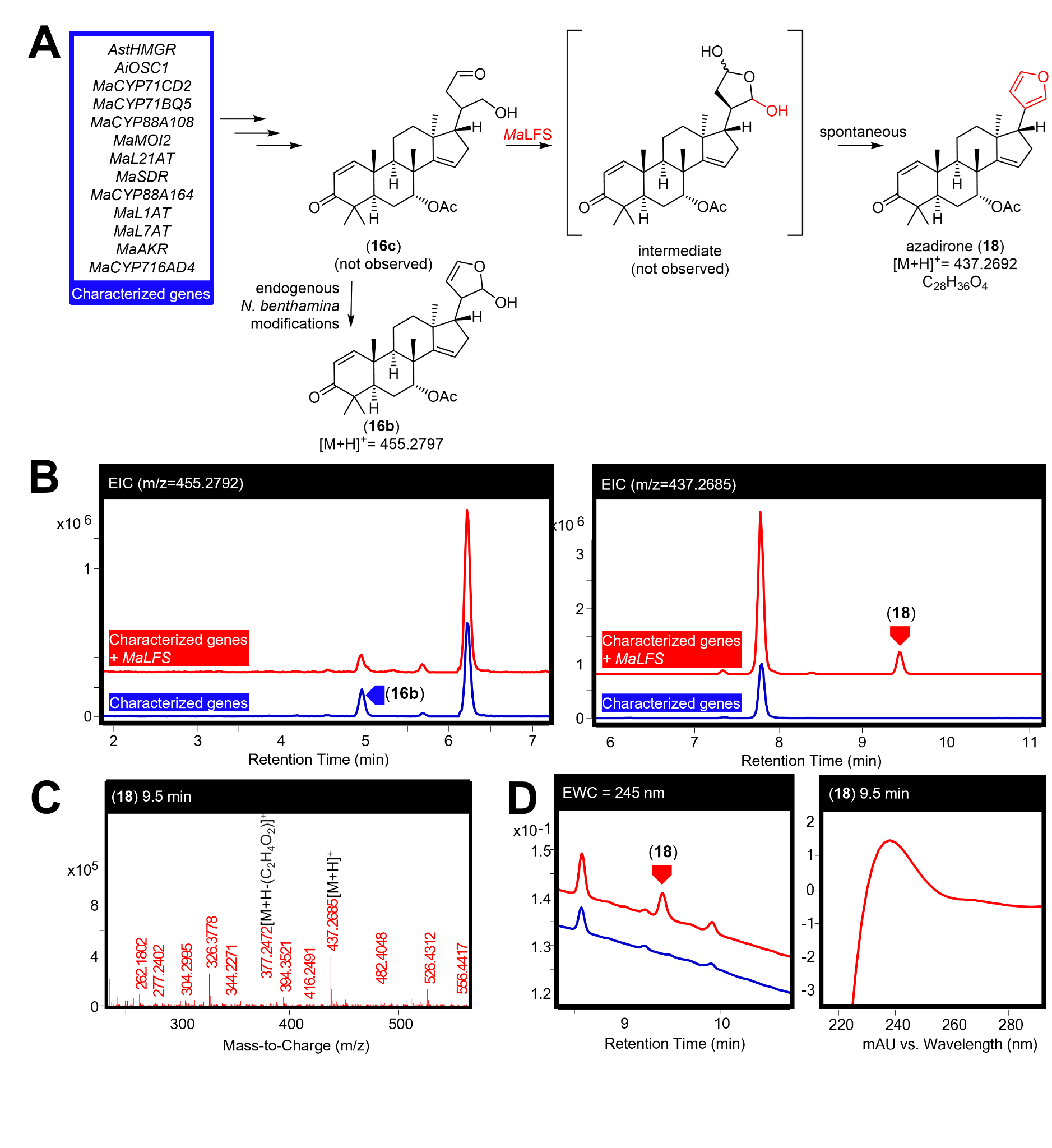

##### Fig. S34. Characterisation of *MaLFS.*

(A) Predicted function of *Ma*LFS in converting (**16c**) to azadirone (**18**) (see Fig. S32. for predicted mechanisms). (B) Extracted ion chromatograms (EICs) for extracts of *N. benthamiana* agro-infiltrated with the characterized genes listed in panel A, either alone (blue) or with the addition of *Ma*LFS (red). The EICs display masses of 455.2792 (observed mass of [(**16b**)+H]^+^) and of 437.2687 (observed mass of [(**18**)+H]^+^). (C) Mass spectra for (**18**) being heterologously produced in *N. benthamiana,* the main observed adduct ([M+H]^+^) and fragment (loss of acetic acid [M+H-C_2_H_4_O_2_]^+^, consistent with the loss of C7 acetoxy group and the literature (*74*)) are labeled. (D) Extracted wavelength chromatograms (EWCs) of 245 nm (width of 4 nm) for extracts displayed in Panel B. UV spectra (mAU) of azadirone (**18**) being heterologously produced in *N. benthamiana* is also given*.* Representative traces and spectra are displayed (n=6).

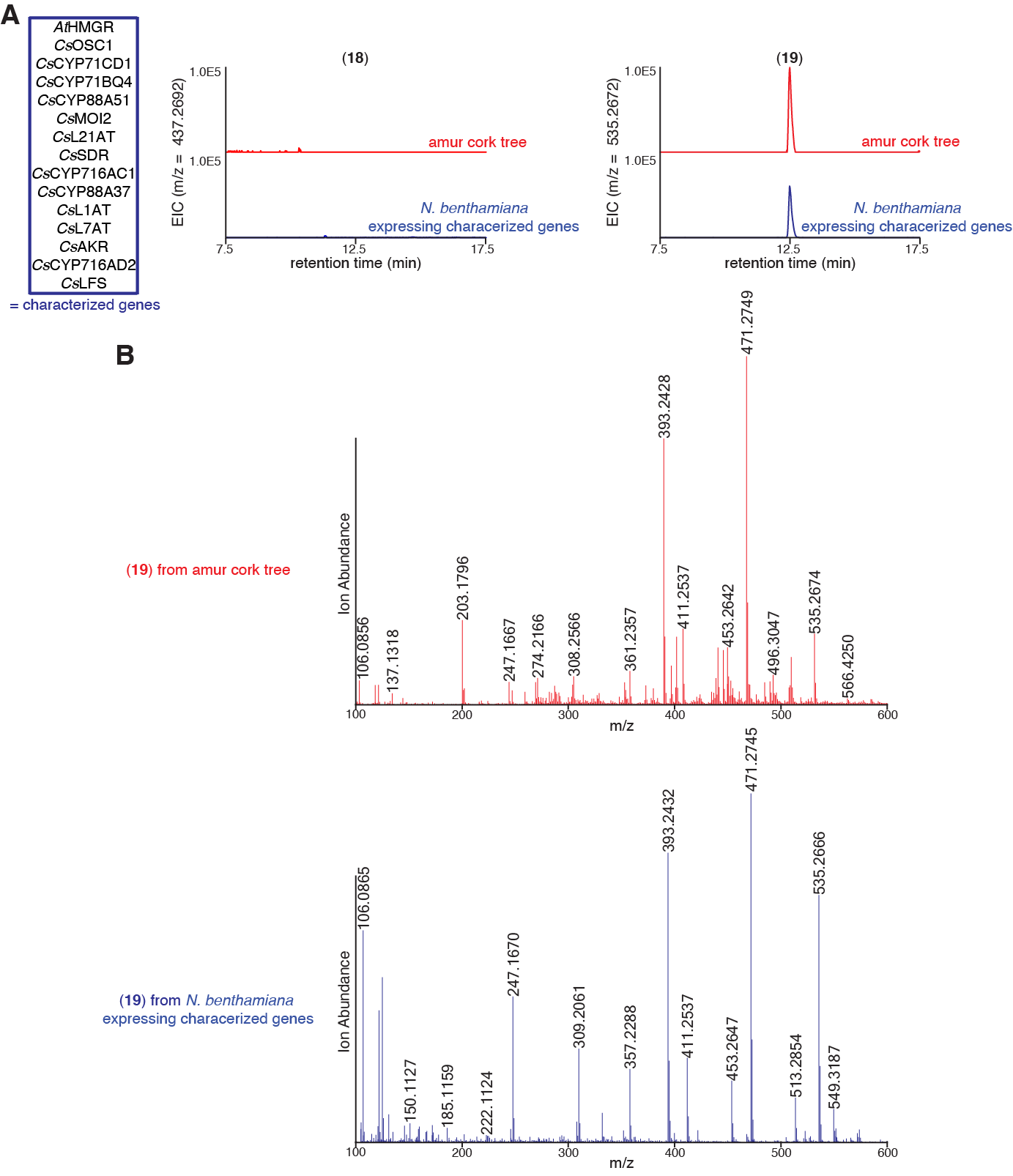

##### Fig. S35. Detection of kihadalactone A (19) but not azadirone (18) in Rutaceae and agro-infiltrated *N. benthamiana* extracts.

(A) Extracted ion chromatograms (EICs) for both *Phellodendron amurense* (amur cork tree) extract (red) and *N. benthamiana* extracts agro-infiltrated with the combinations of genes outlined in the blue box. EICs display masses of [M+H]^+^ = 437.2692 (calculated mass of (**18**)) and [M+Na]^+^=535.2672 (calculated mass of (**19**)). (B) Mass spectrum of (**19**) in *P. amurense* extract (red) or being heterologously produced in *N. benthamiana* (blue) using genes listed in panel A. Representative EICs and mass spectra are displayed, n=3 biological replicates.

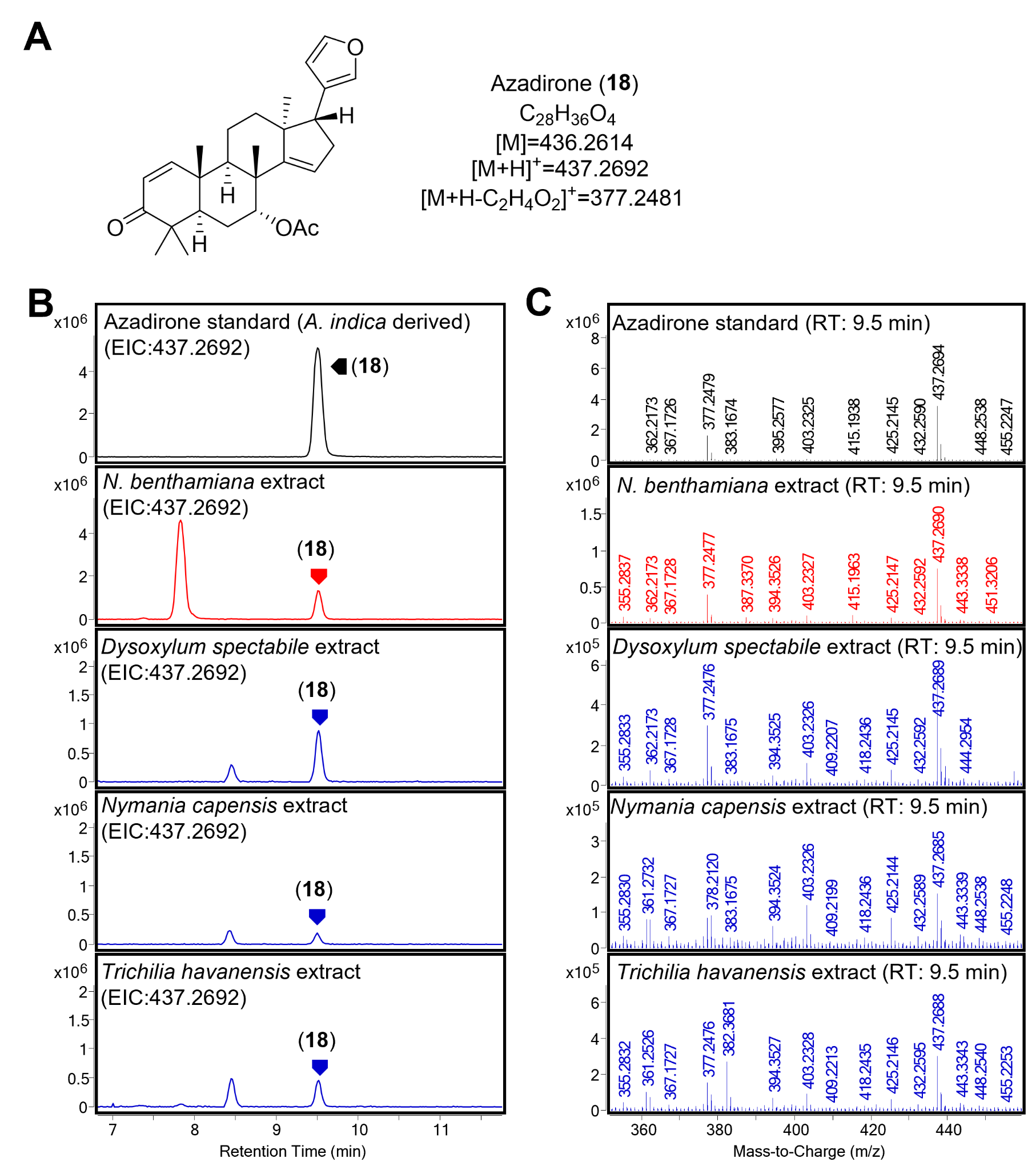

##### Fig. S36. Azadirone (18) in agro-infiltrated *N. benthamiana* and Meliaceae extracts.

(A) Structure, formula and observed adducts of azadirone (**18**), which, as well as being identified in agro-infiltrated *N. benthamiana* extracts, was identified in three (*Trichilia havanensis* (*77*), *Dysoxylum spectabile* and *Nymania capensis*) of the thirteen Meliaceae species sourced from Kew Gardens. (B) Extracted ion chromatograms (EICs) comparing an analytical standard of azadirone (**18**) (black, purified from *A. indica* leaf powder (Table S15)), to extract from *N. benthamiana* expressing azadirone (**18**) biosynthetic enzymes (*Ai*OSC1*, Ma*CYP71CD2, *Ma*CYP71BQ5, *Ma*CYP88A108*, Ma*MOI2, *Ma*L21AT, *Ma*SDR, *Ma*CYP88A164, *Ma*L1AT, *Ma*L7AT, *Ma*AKR, *Ma*CYP716AD4 and *Ma*LFS (red)) and extracts of the three Meliaceae species identified as containing azadirone (**18**) (blue). (C) Mass spectra of azadirone (**18**) peak corresponding to each of the extracts displayed in panel B.

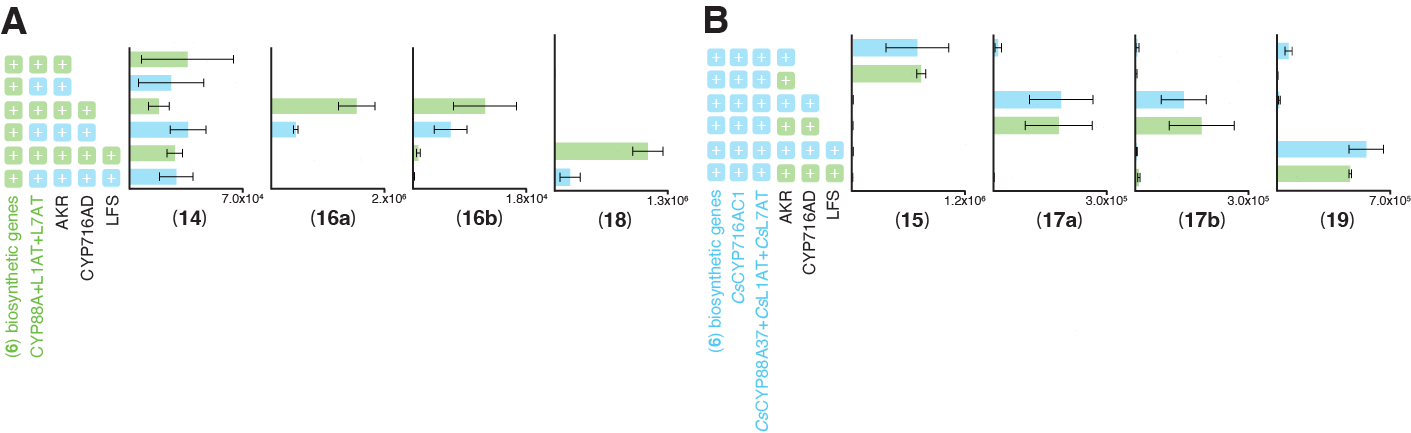

##### Fig. S37. Compatibility of *C. sinensis* and *M. azedarach* pathways.

(A) Integrated peak area of extracted ion chromatograms (EICs) for the last four *M. azedarach* pathway products (**14**, **16a**, **16b**, **18**) being produced by heterologous expression in *N. benthamiana,* either exclusively with enzymes from *M. azedarach* (green), or instead using the relevant *C. sinensis* homologs (blue) (B) Integrated peak area of extracted ion chromatograms (EICs) for the last four *C. sinensis* pathway products (**15**, **17a**, **17b**, **19**) being produced by heterologous expression in *N. benthamiana,* either exclusively with enzymes from *C. sinensis* (blue), or with the relevant *M. azedarach* homologs (green). Biosynthetic enzymes from *M. azedarach* for production of (**6**) are as follows; *Ma*OSC1, *Ma*CYP71CD2, *Ma*CYP71BQ5, *Ma*CYP88A108, *Ma*MOI2, *Ma*L21AT and *Ma*SDR. Biosynthetic enzymes from *C. sinensis* for production of (**6**) are as follows; *Cs*OSC1, *Cs*CYP71CD1, *Cs*CYP71BQ4, *Cs*CYP88A51, *Cs*MOI2, *Cs*L21AT and *Cs*SDR . Additional enzymes used are listed in the figure, CYP88 refers to either *Ma*CYP88A164 or *Cs*CYP88A37. Values and error bars represent the mean and the standard error of the mean; n=3 biological replicates.

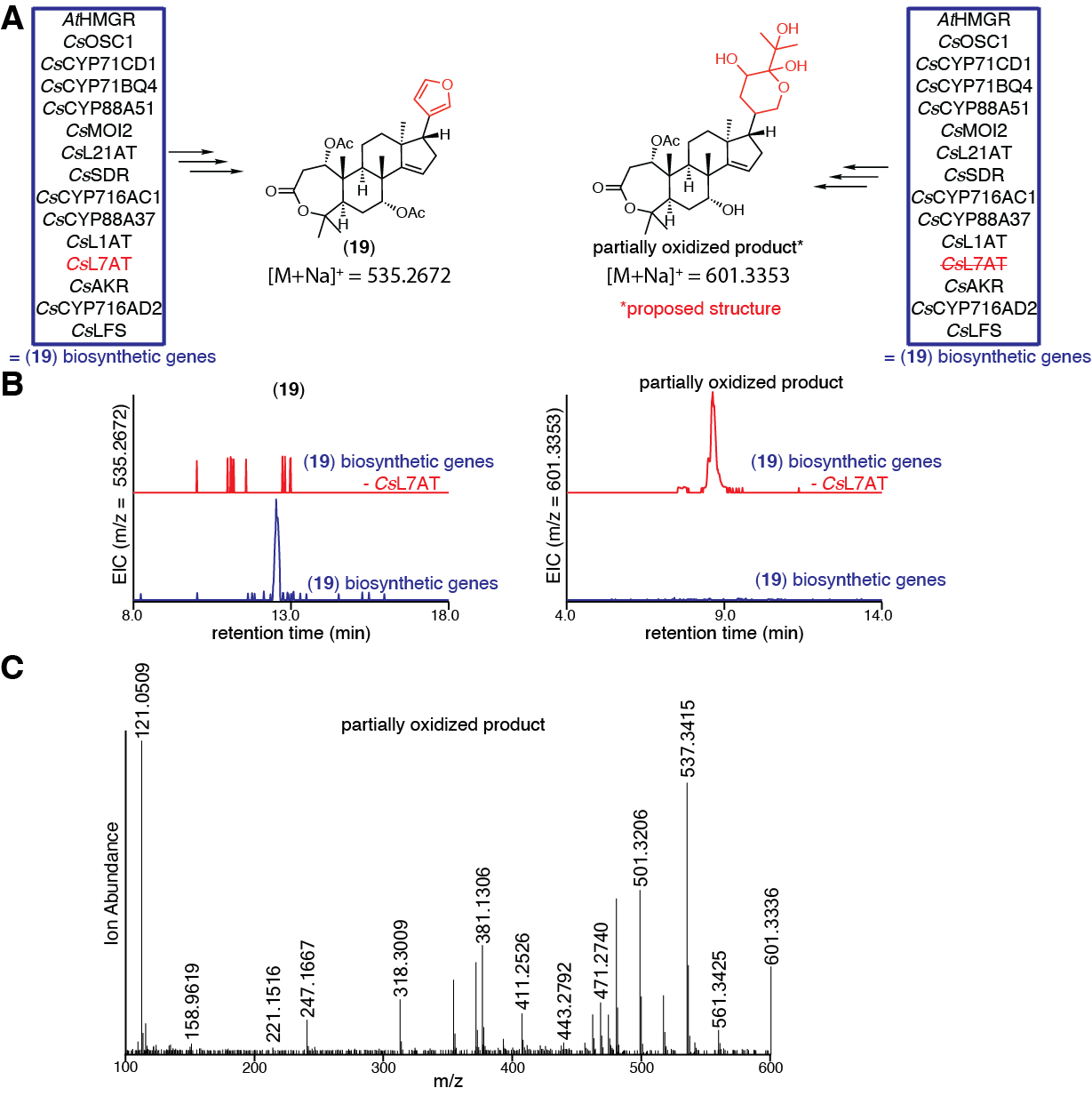

##### Fig. S38. *Cs*L7AT is required for furan formation.

(A) Predicted structures when *Cs*L7AT is either included, or omitted from the set of co-expressed biosynthetic enzymes required for the production of kihadalactone A (**19**). The proposed structure resembles that of (**20**) (Table S16), which was purified from heterologous expression of *M. azedarach* enzymes for (**16**), in the absence of *Ma*L7AT (Fig. S39). (B) Extracted ion chromatograms (EICs) of *N. benthamiana* extracts agro-infiltrated with the combinations of genes outlined in panel A, either with (blue) or without (red) *Cs*L7AT. EICs display masses of [M+Na]^+^=535.2672 (calculated mass of (**19**)) and [M+Na]^+^ = 601.3353 (calculated mass for proposed partially oxidized product). Neither (**19**) nor the corresponding C-7 deacetylated limonoid product was observed in the absence of *Cs*L7AT, however, a new peak of 601.3353 appeared, corresponding to a partially oxidized limonoid presumed to be formed by *Cs*CYP716AD2. (C) Mass spectrum of the observed partially oxidized product heterologously produced in *N. benthamiana* as shown in panel B. Representative EICs and mass spectra are displayed (n=6).

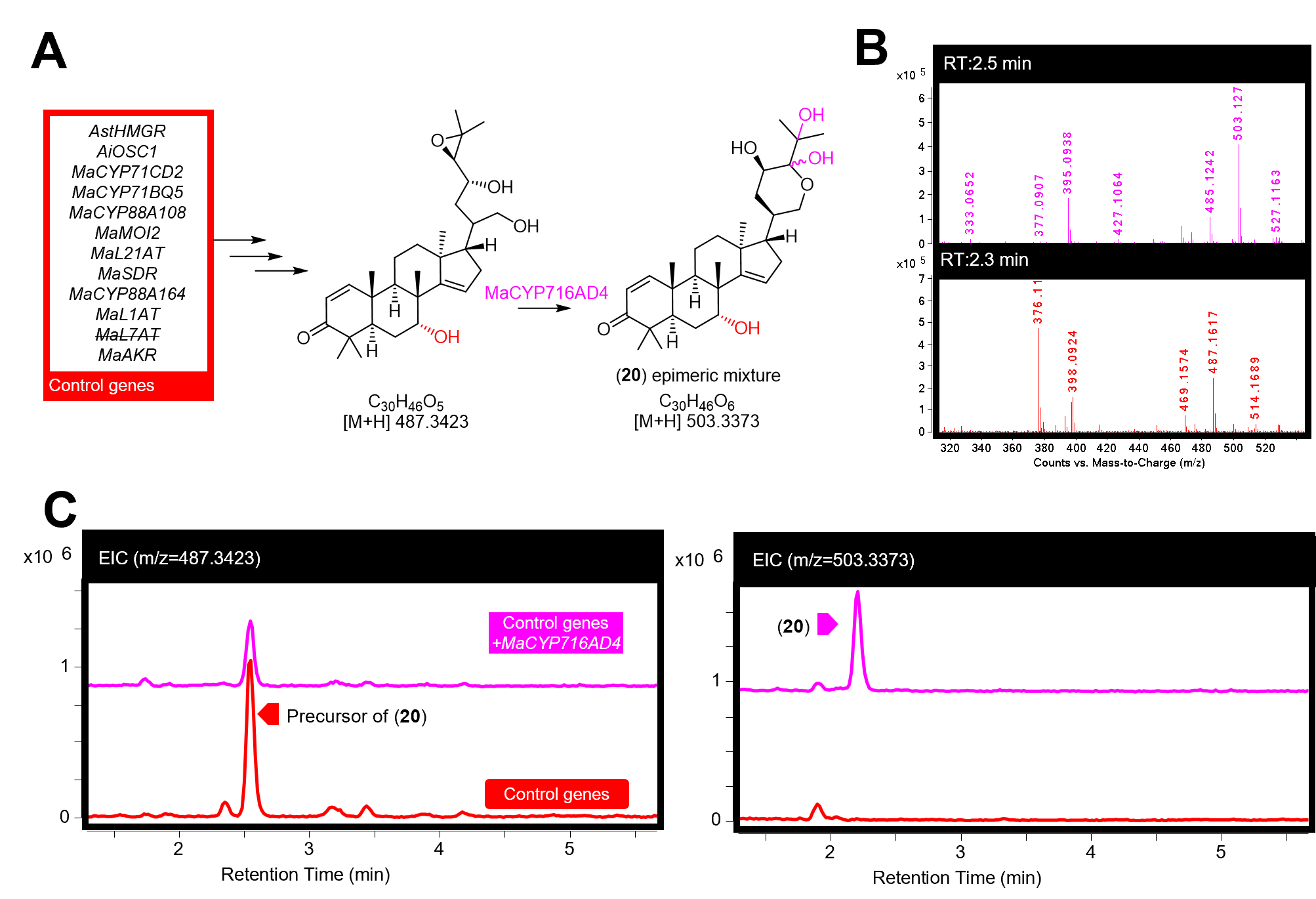

##### Fig. S39. *Ma*CYP716AD4 side-product (formed in the absence of C7-acetoxyl group).

(A) Off-target function of *Ma*CYP716AD4 in producing the side-product (**20**) (NMR confirmed, Table S16). Predicted mechanism is expanded upon in Fig. S32. (B) Mass spectra for (**20**) (pink) and its precursor (red), being heterologously produced in *N. benthamiana,* displaying the main observed adducts (both [M+H]^+^) of 503.1270 and 487.1617, respectively. (C) Extracted ion chromatograms (EICs) for extracts of *N. benthamiana* agro-infiltrated with the control genes (importantly lacking *MaL7AT*) listed in panel A, either alone (red) or with the addition of *Ma*CYP716AD4 (pink). EICs display masses of 487.3423 (predicted mass of the precursor of (**20**) [M+H]^+^) and 503.3373 (predicted mass of (**20**) [M+H]^+^). Representative EICs and mass spectra are displayed (n=3).

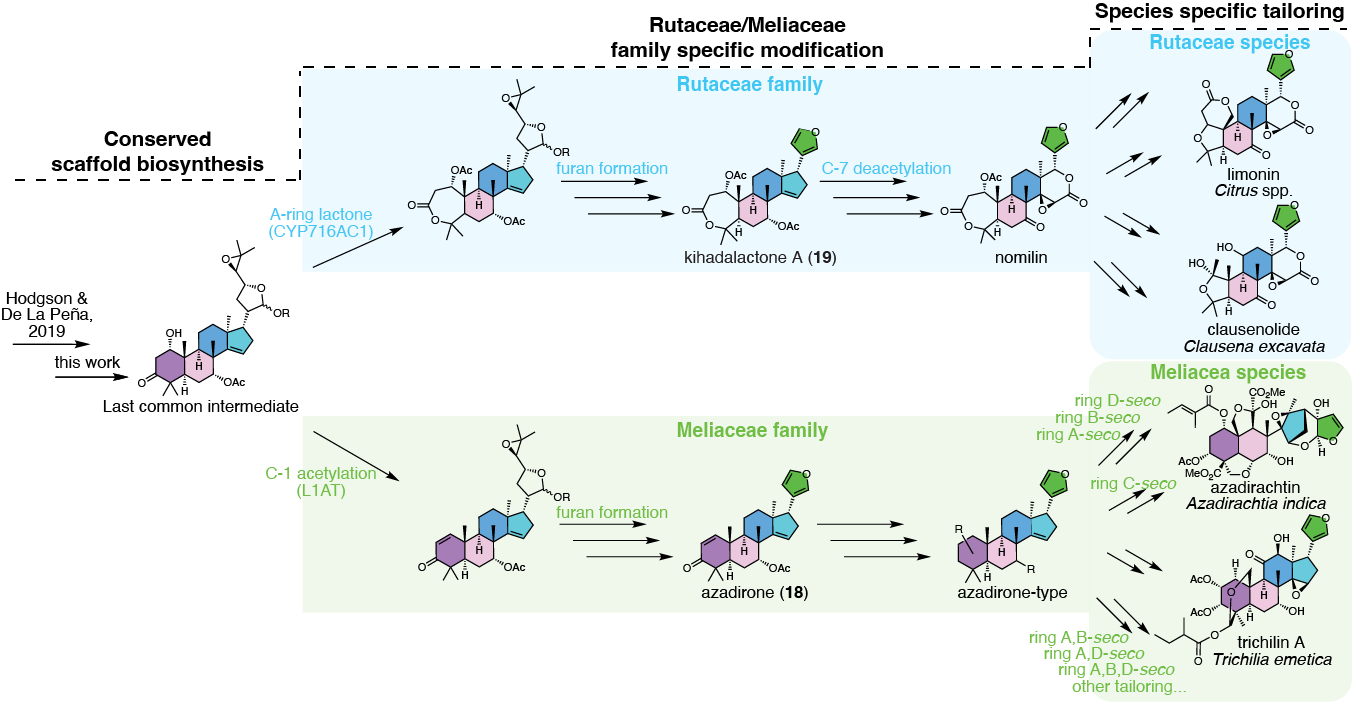

##### Fig. S40. Proposed limonoid biosynthetic pathway in Rutaceae and Meliaceae plants.

A conserved protolimonoid core scaffold biosynthesis is shared between the two families to form the last common biosynthetic intermediate that is structurally similar to (**6**). The pathway diverges with Rutaceae and Meliaceae family specific modifications, notably the A-ring lactone formation by *Cs*CYP716AC1 to yield nomilin- and azadirone-type biosynthetic intermediate. The Melia and Citrus pathways likely go through azadirone (**18**) and kihadalactone A (**19**) as biosynthetic intermediates, respectively, because we have shown that C-7 *O*-acetylation is a prerequisite for furan formation in both pathways. The nomilin- and azadirone-type intermediates can undergo further species-specific tailoring to form structurally diverse limonoids, many of which are species specific (species of isolation is indicated below the molecule name).

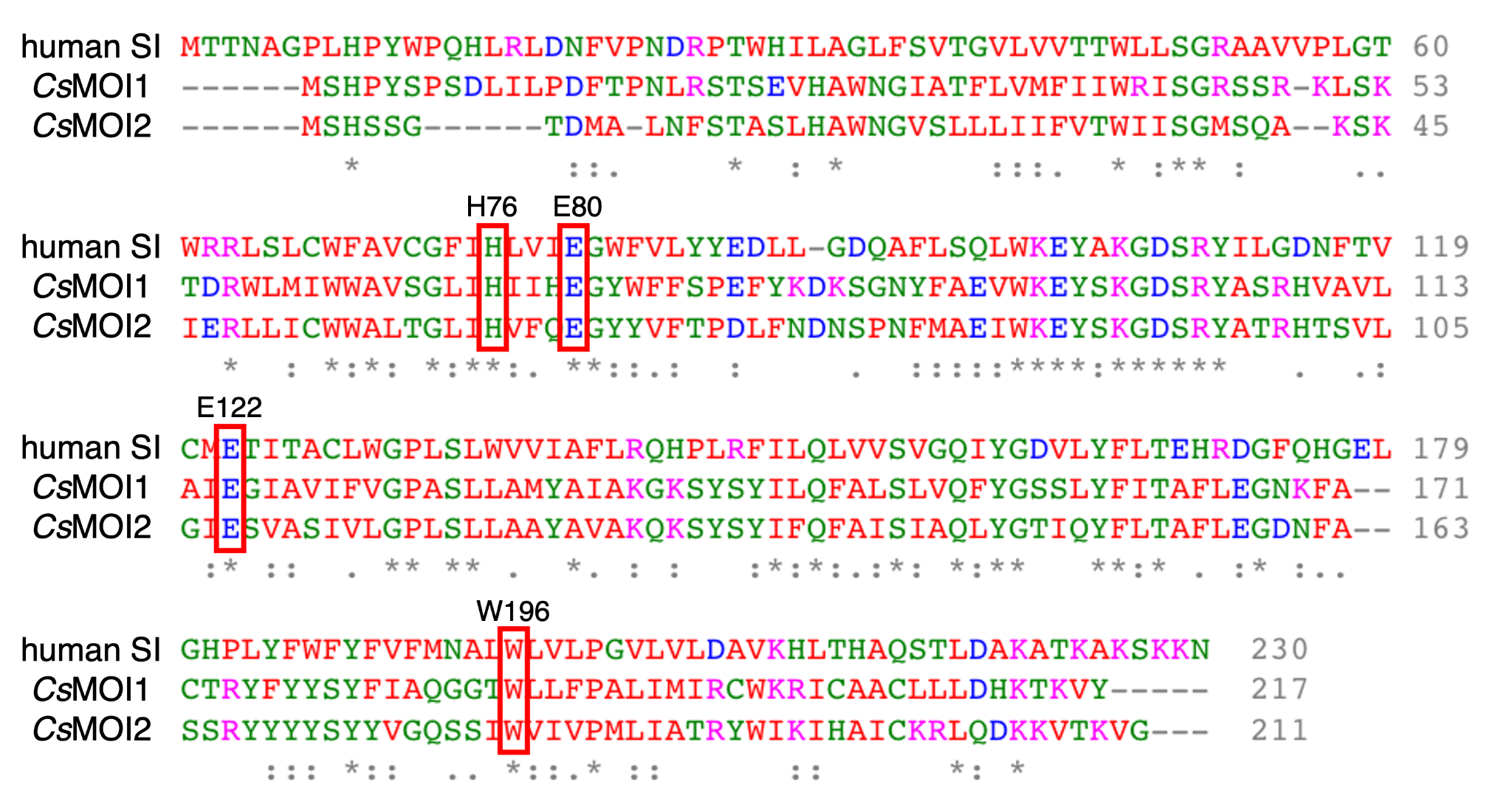

##### Fig. S41. Alignment indicating the conserved active site residues between human sterol isomerase, *Cs*MOI1 and *Cs*MOI2.

The active site of human sterol isomerase (SI) has previously been studied through protein crystal structural analysis and substrate docking (*78*). The residues H76, E80, E122, and W196 (highlighted in red boxes) of human SI were each proposed to be key in stabilizing the carbocation intermediate during isomerization. The conservation of these residues in *Cs*MOI1/2 suggests a similar isomerization mechanism via the formation of a carbocation. To determine how two different types of rearrangements are controlled by *Cs*MOI1 and *Cs*MOI2 despite their conserved active site residues, will require further study on the binding pocket of these enzymes. The protein sequences were aligned through the online Clustal Omega tool.

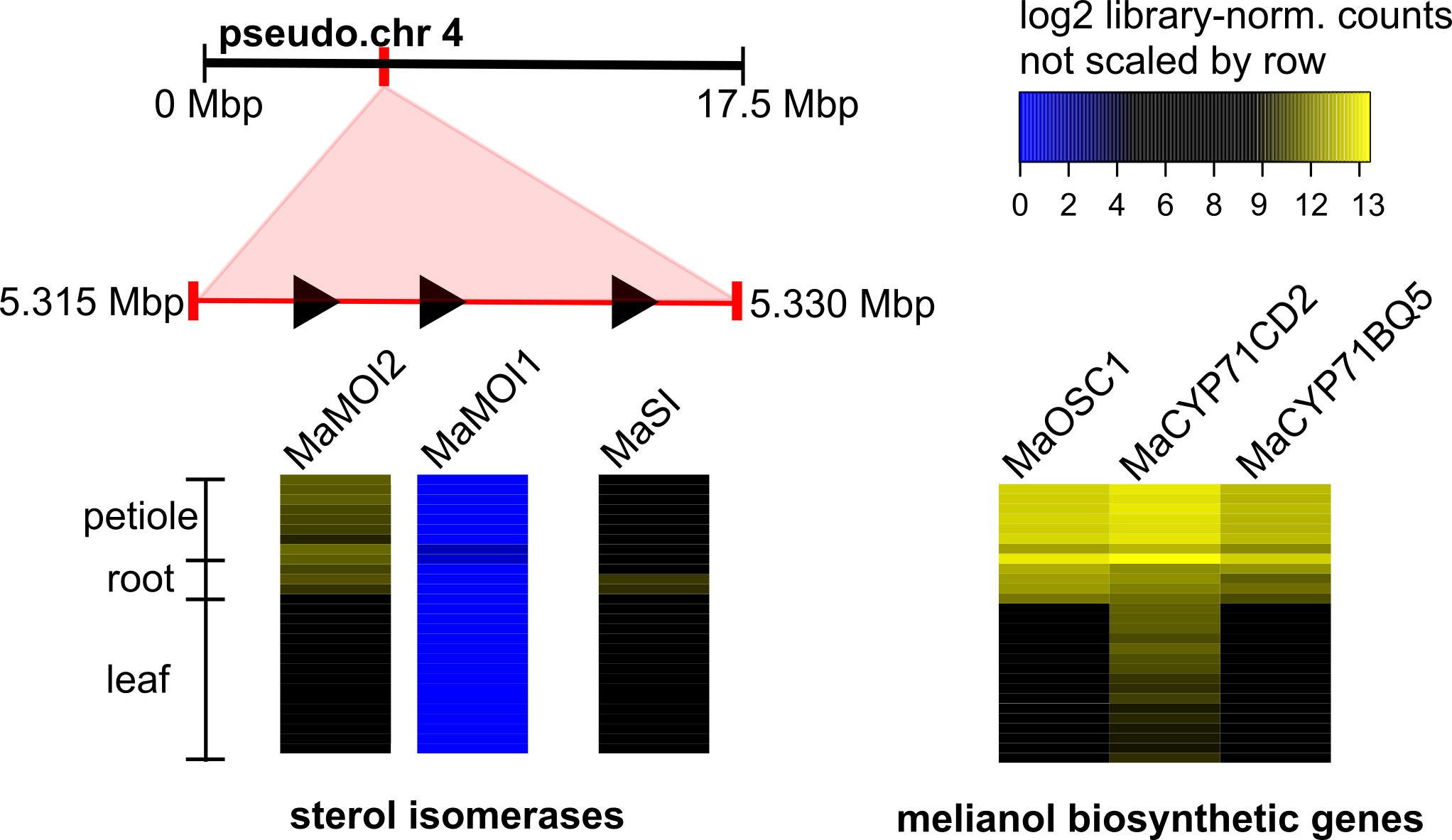

##### Fig. S42. Genomic location and expression patterns of sterol isomerases in *M. azedarach*.

The expression pattern and genomic location (on pseudo-chromosome 4) of all sterol isomerase candidates (IPR007905 (Emopamil-binding protein)) in the *M. azedarach* genome. Gene IDs are provided (Table S10). The expression pattern of melianol biosynthetic genes has been included for comparative purposes. Heatmap was constructed using library normalized log_2_ read counts in Heatmap3 V1.1.1 (*46*) (with no scaling by row). *MaSI* was cloned and found to have no melianol oxide isomerase activity when tested (by co-expression in *N. benthamiana* with melianol biosynthetic genes and *MaCYP88A108*). A protein coding version of *MaMOI1* was not amplifiable based on the *M. azedarach* genome annotation.

### Supplementary Tables

##### Table S1. Summary of *M. azedarach* genome assembly and annotation.

| ***Melia azedarach* genome assembly statistics** | |
| --- | --- |
| Number of contigs | 346 |
| Largest contig | 20,704,184 |
| Total length | 230,865,674 |
| GC (%) | 32.21 |
| N50 | 16,923,081 |
| N75 | 14,637,465 |
| L50 | 7 |
| L75 | 10 |
| Ns per 100 kbp | 9.44 |
| ***Melia azedarach* genome annotation statistics** | |
| **Genes** | |
| Total number of genes | 26,738 |
| Protein coding (high) | 22,785 |
| Transposable element (high) | 1,250 |
| Predicted (low) | 230 |
| Protein coding (low) | 1,651 |
| Transposable element (low) | 822 |
| **Transcripts** | |
| Transcripts per gene | 1.16 |
| Total number of transcripts | 31,048 |
| **CDS** | |
| Transcript mean size CDS (bp) | 1,309.11 |
| Min CDS | 78 |
| Max CDS | 15,903 |
| CDS mean size (bp) | 245.97 |
| Exon mean size (bp) | 312.11 |
| Exons per transcript | 5.71 |
| Total exons | 177,227 |
| Monoexonic transcripts | 5,473 |
| **cDNA** | |
| Transcript mean size cDNA (bp) | 1,781.55 |
| Min cDNA | 114 |
| Max cDNA | 16,537 |
| Intron mean size (bp) | 392.02 |
| 5UTR mean size (bp) | 186.24 |
| 3UTR mean size (bp) | 286.21 |

**Table S1. Summary of *M. azedarach* genome assembly and annotation continued.**

| **BUSCO- assessment** | | |
| --- | --- | --- |
|  | ***Melia azedarach*** | ***Arabidopsis thaliana*** |
| Complete genes (single-copy) | 1,339 | 1,416 |
| Complete genes (2 copies) | 46 | 11 |
| Complete genes (3+ copies) | 7 | 4 |
| Fragmented genes | 20 | 5 |
| Missing genes | 28 | 4 |

*M. azedarach* pseudo-chromosome level genome statistics were generated by QUAST V.4.6.3 (*79*) and are based on contigs of size ≥500 bp. Statistics for *M. azedarach* annotation generated by the Earlham Institute. Genes are classified as either: protein coding, predicted (limited homology support <30%) or transposable element (>40% overlap with interspersed repeats). Genes were assigned a confidence classification of high or low based on their ability to meet specified criteria (>80% coverage to reference proteins or >60% protein coverage with >40% of the structure supported by transcriptome data). Statistics for coding sequences (CDS) and complementary DNA (cDNA) as also included. BUSCO (Benchmarking Universal Single-Copy Orthologs) (*24*) assessment of protein annotation of *M. azedarach* and gold standard *Arabidopsis thaliana*, performed by the Earlham Institute.

##### Table S2. Summary of paired end reads generated for *M. azedarach* RNA-seq.

| **Sample** | **Rep.** | **Lane 1** | **Lane2** | **Total**  **(per rep.)** | **Total**  **(per sample)** |
| --- | --- | --- | --- | --- | --- |
| *M. azedarach* 'individual 11'  Upper Leaf | 1A | 7,312,258 | 7,957,007 | 15,269,265 | 78,676,364 |
|  | 1B | 12,440,818 | 13,367,863 | 25,808,681 |  |
|  | 1C | 9,677,158 | 10,402,466 | 20,079,624 |  |
|  | 1D | 8,501,858 | 9,016,936 | 17,518,794 |  |
| *M. azedarach* 'individual 11'  Lower Leaf | 2A | 14,706,042 | 15,713,081 | 30,419,123 | 95,833,402 |
|  | 2B | 9,952,003 | 10,506,690 | 20,458,693 |  |
|  | 2C | 9,995,844 | 10,724,057 | 20,719,901 |  |
|  | 2D | 11,759,629 | 12,476,056 | 24,235,685 |  |
| *M. azedarach* 'individual 11'  Petiole | 3A | 11,225,462 | 12,293,851 | 23,519,313 | 82,662,893 |
|  | 3B | 8,518,447 | 9,151,386 | 17,669,833 |  |
|  | 3C | 8,723,766 | 9,267,735 | 17,991,501 |  |
|  | 3D | 11,360,248 | 12,121,998 | 23,482,246 |  |
| *M. azedarach* 'individual 11'  Root | 4A | 12,795,130 | 13,497,456 | 26,292,586 | 107,736,216 |
|  | 4B | 9,430,278 | 10,235,484 | 19,665,762 |  |
|  | 4D | 14,075,197 | 14,780,596 | 28,855,793 |  |
|  | 4F | 15,951,734 | 16,970,341 | 32,922,075 |  |
| *M. azedarach* 'individual 02'  Upper Leaf | 5A | 8,425,596 | 8,942,230 | 17,367,826 | 86,473,401 |
|  | 5B | 7,483,256 | 7,905,622 | 15,388,878 |  |
|  | 5C | 15,588,782 | 16,252,245 | 31,841,027 |  |
|  | 5D | 10,597,294 | 11,278,376 | 21,875,670 |  |
| *M. azedarach* 'individual 02'  Lower Leaf | 6A | 7,570,717 | 7,949,090 | 15,519,807 | 83,790,681 |
|  | 6B | 15,757,443 | 16,754,196 | 32,511,639 |  |
|  | 6C | 8,341,297 | 8,628,045 | 16,969,342 |  |
|  | 6D | 9,074,779 | 9,715,114 | 18,789,893 |  |
| *M. azedarach* 'individual 02'  Petiole | 7A | 15,145,250 | 16,073,522 | 31,218,772 | 100,692,927 |
|  | 7B | 11,317,371 | 12,034,239 | 23,351,610 |  |
|  | 7C | 12,710,000 | 13,530,865 | 26,240,865 |  |
|  | 7D | 9,595,536 | 10,286,144 | 19,881,680 |  |

Numbers of paired-end are reported per lane, replicate and sample. Petiole samples include. rachis.

##### Table S3. ^13^C & ^1^H δ assignments of *apo*-melianol (3) produced using heterologously expressed genes from *M. azedarach* (C-21 epimeric mixture)

| **Carbon numbering scheme and selected COSY, HMBC and NOESYS** | | | | | | | | | |
| --- | --- | --- | --- | --- | --- | --- | --- | --- | --- |
| 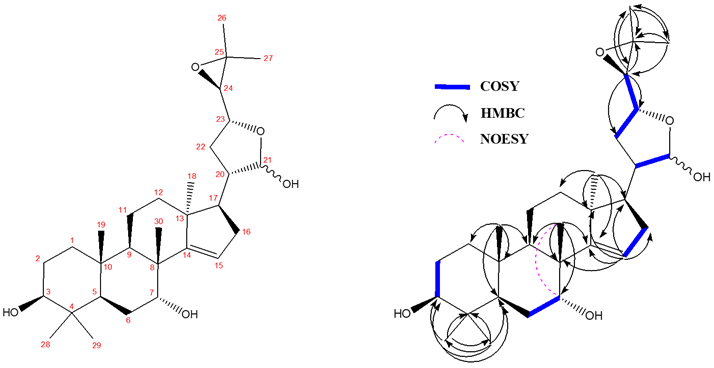 | | | | | | | | | |
| **C** | **^13^C δ**  **(150 MHz)** | | **^1^H δ**  **(600 MHz)** | | **C** | **^13^C δ**  **(150 MHz)** | | **^1^H δ**  **(600 MHz)** | |
| **14** | 162.53 | 162.20 | / | | **1** | 37.91 | 37.88 | 1.61 (1H, m)  1.04 (1H, m) | |
| **15** | 119.52 | 119.04 | 5.47 (1H, m) | 5.46 (1H, m) | **10** | 37.58 | 37.57 | / | |
| **21** | 102.38 | 97.59 | 5.39 (1H, m) | 5.38 (1H, m) | **16** | 35.07 | 34.73 | 2.18 (2H, m) | 2.12 (2H, m) |
| **3** | 78.77 | 78.74 | 3.28 (1H, dd J= 11.3, 4.5) | | **22** | 34.71 | 31.35 | 2.09 (1H, m)  1.41 (1H, m) | 2.02 (1H, m)  1.74 (1H, m) |
| **23** | 78.37 | 77.24 | 3.91 (1H, m) | 3.96 (1H, m) | **12** | 33.06 | 32.80 | 1.82 (1H, m)  1.52 (1H, m) | 1.98 (1H, m)  1.46 (1H, m) |
| **7** | 72.33 | 72.32 | 3.93 (1H, m) | | **28** | 27.67 | | 0.99 (3H, s) | |
| **24** | 67.62 | 65.21 | 2.83  (1H, d J= 7.5) | 2.70  (1H, d J= 7.5) | **30** | 27.65 | 27.63 | 1.058 (3H, s) | 1.056 (3H, s) |
| **25** | 58.09 | 57.27 | / | | **2** | 27.14 | | 1.64 (1H, m)  1.58 (1H, m) | |
| **17** | 57.51 | 52.73 | 1.74 (1H, m) | 2.01 (1H, m) | **26** | 25.02 | 24.92 | 1.33 (3H, s) | 1.32 (3H, s) |
| **20** | 47.80 | 45.48 | 2.39 (1H, m) | 1.49 (1H, m) | **6** | 23.68 | 23.66 | 1.85 (1H, m)  1.74 (1H, m) | |
| **13** | 47.09 | 46.70 | / | | **18** | 19.79 | 19.45 | 1.02 (3H, s) | 1.09 (3H, s) |
| **5** | 46.53 | 46.49 | 1.50 (1H, m) | 1.48 (1H, m) | **27** | 19.35 | 19.21 | 1.32 (3H, s) | |
| **8** | 44.26 | 44.24 | / | | **11** | 16.34 | 16.29 | 1.69 (1H, m)  1.52 (1H, m) | |
| **9** | 41.78 | 41.74 | 1.93 (1H, m) | 1.91 (1H, m) | **29** | 15.45 | | 0.79 (3H, s) | |
| **4** | 38.36 | | / | | **19** | 15.41 | 15.39 | 0.89 (3H, s) | |

NMR spectra were recorded in CDCl_3_, referenced to TMS and characterization was performed following the general considerations outlined.

##### Table S4. ^13^C & ^1^H δ assignments of (6) produced using heterologously expressed genes from *C. sinensis.*

| **Carbon numbering scheme and selected COSY and HMBC** | | | | | | | |
| --- | --- | --- | --- | --- | --- | --- | --- |
| 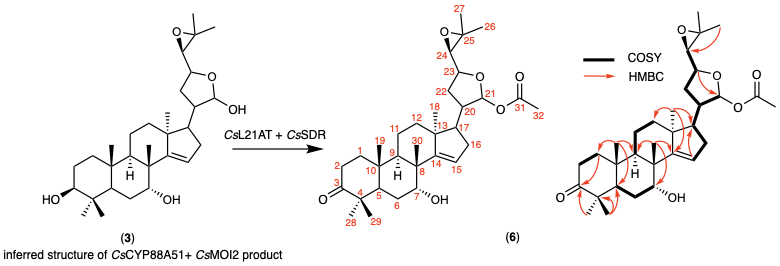 | | | | | | | |
| **C** | **^13^C δ (ppm)** | **^1^H δ**  **(ppm, J in Hz)** | | **C** | **^13^C δ**  **(ppm)** | **^1^H δ**  **(ppm, J in Hz)** | |
| **1** | 38.58 | 1.49 (1H, m) | 1.82 (1H, m) | **17** | 52.73 | 1.91 (1H, dt J = 8.1, 10.4) | |
| **2** | 34.03 | 2.41 (1H, ddd J = 3.8, 7.5, 15.8) | 2.51 (1H, ddd J = 7.5, 10.3, 15.7) | **18** | 19.82 | 1.02 (3H, s) | |
| **3** | 217.31 | / | | **19** | 15.04 | 0.98 (3H, s) | |
| **4** | 47.00 | / | | **20** | 44.33 | 2.35 (1H, dddd J = 4.1, 7.0, 10.9, 12.5) | |
| **5** | 46.71 | 2.07 (1H, m) | | **21** | 96.70 | 6.24 (1H, d J = 4.1) | |
| **6** | 24.92 | 1.77 (1H, m) | 1.82 (1H, m) | **22** | 31.50 | 1.68 (1H, m) | 2.07 (1H, m) |
| **7** | 72.05 | 3.95 (1H, appt t = 2.8) | | **23** | 79.86 | 3.91 (1H, dt J = 9.9, 7.2) | |
| **8** | 44.17 | / | | **24** | 66.81 | 2.65 (1H, d J = 7.5) | |
| **9** | 40.91 | 1.99 (1H, dd J = 7.6, 12.0) | | **25** | 57.25 | / | |
| **10** | 37.30 | / | | **26** | 19.45 | 1.27 (3H, s) | |
| **11** | 16.38 | 1.54 (1H, m) | 1.68 (1H, m) | **27** | 25.03 | 1.31 (3H, s) | |
| **12** | 32.45 | 1.29 (1H, m) | 1.59 (1H, m) | **28** | 21.28 | 1.03 (3H, s) | |
| **13** | 46.62 | / | | **29** | 26.35 | 1.08 (3H, s) | |
| **14** | 161.70 | / | | **30** | 27.36 | 1.08 (3H, s) | |
| **15** | 119.72 | 5.49 (1H, dd J = 1.9, 3.4) | | **31** | 170.05 | / | |
| **16** | 35.19 | 2.2 (2H, m) | | **32** | 21.61 | 2.04 (3H, s) | |

NMR spectra were recorded in CDCl_3_, referenced to TMS and characterization was performed following the general considerations outlined. Literature comparison is also given (Table S7).

##### Table S5. ^13^C & ^1^H δ assignments of (4’) produced using heterologously expressed genes from *C. sinensis.*

| **Carbon numbering scheme and selected COSY and HMBC** | | | | | | | |
| --- | --- | --- | --- | --- | --- | --- | --- |
| 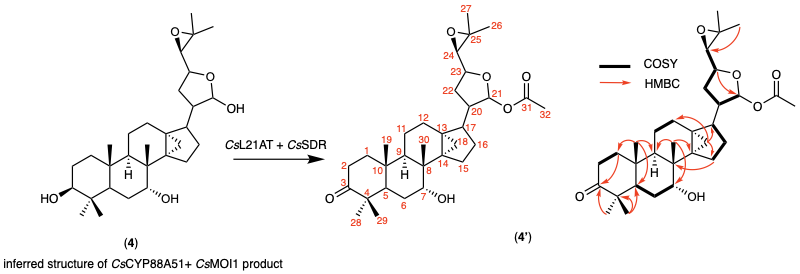 | | | | | | | |
| **C** | **^13^C δ (ppm)** | **^1^H δ**  **(ppm, J in Hz)** | | **C** | **^13^C δ**  **(ppm)** | **^1^H δ**  **(ppm, J in Hz)** | |
| **1** | 39.24 | 1.42 (1H, dt  J = 13.2, 8.6) | 1.78 (1H, dt J = 13.2, 8.6) | **17** | 44.89 | 2.08 (1H, m) | |
| **2** | 33.91 | 2.43 (2H, dd J = 6.2, 8.6) | | **18** | 13.61 | 0.44  (1H, d J = 5.0) | 0.69  (1H, d J = 5.0) |
| **3** | 217.68 | / | | **19** | 15.74 | 0.92 (3H, s) | |
| **4** | 46.78 | / | | **20** | 48.03 | 2.02 (1H, m) | |
| **5** | 45.69 | 2.07 (1H, m) | | **21** | 97.55 | 6.25 (1H, d J = 3.8) | |
| **6** | 25.5 | 1.64 (2H, m) | | **22** | 30.64 | 1.63 (1H, m) | 2.01 (1H, m) |
| **7** | 73.88 | 3.76 (1H, appt t J = 2.7) | | **23** | 79.91 | 3.86 (1H, ddd J = 6.2, 7.5, 9.6) | |
| **8** | 36.88 | / | | **24** | 66.78 | 2.64 (1H, d J = 7.6) | |
| **9** | 42.96 | 1.32 (1H, m) | | **25** | 57.21 | / | |
| **10** | 36.86 | / | | **26** | 24.99 | 1.29 (3H, s) | |
| **11** | 16.66 | 1.26 (1H, m) | 1.33 (1H, m) | **27** | 19.40 | 1.24 (3H, s) | |
| **12** | 25.26 | 1.64 (1H, m) | 1.72 (1H, m) | **28** | 26.71 | 1.06 (3H, s) | |
| **13** | 28.75 | / | | **29** | 21.09 | 0.99 (3H, s) | |
| **14** | 38.75 | / | | **30** | 19.53 | 1.03 (3H, s) | |
| **15** | 26.32 | 1.56 (1H, dd J = 8.3, 12.6) | 1.92 (1H, m) | **31** | 170.01 | / | |
| **16** | 27.56 | 0.93 (1H, m) | 1.68 (1H, m) | **32** | 21.66 | 2.06 (3H, s) | |

NMR spectra were recorded in CDCl_3_, referenced to TMS and characterization was performed following the general considerations outlined.

##### Table S6. ^13^C & ^1^H δ assignments of 21(*S*)-acetoxyl-*apo*-melianone (6) produced using heterologously expressed genes from *M. azedarach.*

| **Carbon numbering scheme and selected COSY and HMBC** | | | | | | |
| --- | --- | --- | --- | --- | --- | --- |
| 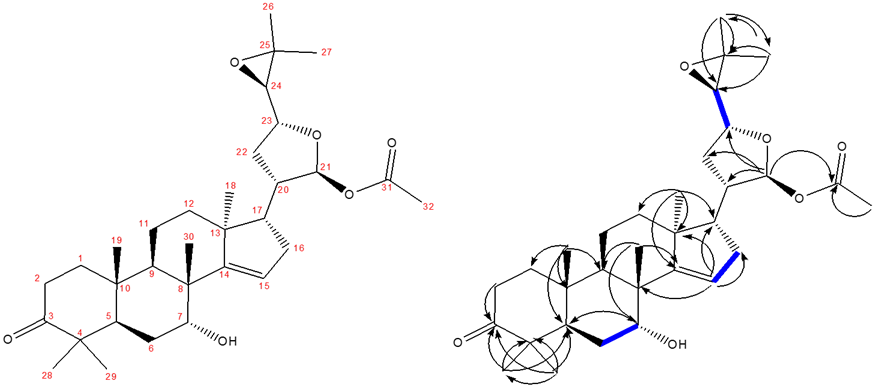 | | | | | | |
| **C** | **^13^C δ**  **(150 MHz)** | **^1^H δ**  **(600 MHz)** | **C** | **^13^C δ**  **(150 MHz)** | **^1^H δ**  **(600 MHz)** | |
| **3** | 217.27 | / | **1** | 38.46 | 1.85 (1H, m) | 1.51 (1H, m) |
| **31** | 169.97 | / | **10** | 37.19 | / | |
| **14** | 161.62 | / | **16** | 35.09 | 2.22 (2H, m) | |
| **15** | 119.61 | 5.51 (1H, m) | **2** | 33.93 | 2.53 (1H, m) | 2.43 (1H, m) |
| **21** | 96.59 | 6.26 (1H, d J= 4.1) | **12** | 32.32 | 1.61 (1H, m) | 1.30 (1H, m) |
| **23** | 79.75 | 3.93 (1H, ddd, J= 7.6, 7.8, 10) | **22** | 31.40 | 2.08 (1H, m) | 1.70 (1H, m) |
| **7** | 71.95 | 3.96 (1H, appt t, J= 2.9) | **30** | 27.26 | 1.10 (3H, s) | |
| **24** | 66.71 | 2.67 (1H, d, J= 7.6) | **28** | 26.25 | 1.10 (3H, s) | |
| **25** | 57.17 | / | **26** | 24.93 | 1.33 (3H, s) | |
| **17** | 52.62 | 1.93 (1H, m) | **6** | 24.80 | 1.84 (1H, m) | 1.79 (1H, m) |
| **4** | 46.91 | / | **32** | 21.52 | 2.06 (3H, s) | |
| **13** | 46.61 | / | **29** | 21.18 | 1.05 (3H, s) | |
| **5** | 46.51 | 2.08 (1H, m) | **18** | 19.71 | 1.03 (3H, s) | |
| **20** | 44.21 | 2.37 (1H, m) | **27** | 19.35 | 1.29 (3H, s) | |
| **8** | 44.07 | / | **11** | 16.27 | 1.71 (1H, m) | 1.56 (1H, m) |
| **9** | 40.80 | 2.01 (1H, m) | **19** | 14.95 | 1.00 (3H, s) | |

NMR spectra were recorded in CDCl_3_, referenced to TMS and characterization was performed following the general considerations outlined. The compound was assigned as the C21(*S*) epimer on the basis of observed NOEs (Fig. S15), also consistent with the literature (Table S7).

##### Table S7. ^13^C δ comparison with the literature for 21(*S*)-acetoxyl-*apo*-melianone (6).

| **C** | **Literature*** | ***M. azedarach* (rounded)** | **Δ Literature to *M. azedarach*** | ***C. sinensis* (rounded)** | **Δ Literature to *C. sinensis*** |
| --- | --- | --- | --- | --- | --- |
| **3** | **217.2** | 217.3 | 0.1 | 217.31 | 0.11 |
| **31** | **170** | 170 | 0 | 170.05 | 0.05 |
| **14** | **161.5** | 161.6 | 0.1 | 161.7 | 0.2 |
| **15** | **119.6** | 119.6 | 0 | 119.72 | 0.12 |
| **21** | **96.6** | 96.6 | 0 | 96.7 | 0.1 |
| **23** | **79.7** | 79.7 | 0 | 79.86 | 0.16 |
| **7** | **71.9** | 71.9 | 0 | 72.05 | 0.15 |
| **24** | **66.7** | 66.7 | 0 | 66.81 | 0.11 |
| **25** | **57.1** | 57.2 | 0.1 | 57.25 | 0.15 |
| **17** | **52.6** | 52.6 | 0 | 52.73 | 0.13 |
| **4** | **46.9** | 46.9 | 0 | 47 | 0.1 |
| **5** | **46.5** | 46.5 | 0 | 46.71 | 0.21 |
| **13** | **46.5** | 46.6 | 0.1 | 46.62 | 0.12 |
| **20** | **44.2** | 44.2 | 0 | 44.33 | 0.13 |
| **8** | **44** | 44.1 | 0.1 | 44.17 | 0.17 |
| **9** | **40.8** | 40.8 | 0 | 40.91 | 0.11 |
| **1** | **38.5** | 38.5 | 0 | 38.58 | 0.08 |
| **10** | **37.1** | 37.2 | 0.1 | 37.3 | 0.2 |
| **16** | **35.1** | 35.1 | 0 | 35.19 | 0.09 |
| **2** | **33.9** | 33.9 | 0 | 34.03 | 0.13 |
| **12** | **32.3** | 32.3 | 0 | 32.45 | 0.15 |
| **22** | **31.3** | 31.4 | 0.1 | 31.5 | 0.2 |
| **30** | **27.2** | 27.3 | 0.1 | 27.36 | 0.16 |
| **28** | **26.2** | 26.2 | 0 | 26.35 | 0.15 |
| **6** | **24.9** | 24.8 | -0.1 | 24.92 | 0.02 |
| **26** | **24.9** | 24.9 | 0 | 25.03 | 0.13 |
| **32** | **21.5** | 21.5 | 0 | 21.61 | 0.11 |
| **29** | **21.1** | 21.2 | 0.1 | 21.28 | 0.18 |
| **18** | **19.7** | 19.7 | 0 | 19.82 | 0.12 |
| **27** | **19.3** | 19.4 | 0.1 | 19.45 | 0.15 |
| **11** | **16.3** | 16.3 | 0 | 16.38 | 0.08 |
| **19** | **14.9** | 14.9 | 0 | 15.04 | 0.14 |

Comparison of the ^13^C δ values for (**6**) from literature (100 mHZ) and this work (150 mHZ), all performed in CDCl_3_. Asterix (*) refers to literature assignment present in (*32*). Full assignment of (**6**) purified from heterologous expression of *M. azedarach* (Table S6) and *C. sinensis* (Table S4) enzymes is provided.

##### Table S8. ^13^C & ^1^H δ assignments of 1-hydroxyl luvungin A (9) produced using heterologously expressed genes from *C. sinensis.*

| **Carbon numbering scheme and selected COSY and HMBC** | | | | | | | |
| --- | --- | --- | --- | --- | --- | --- | --- |
| 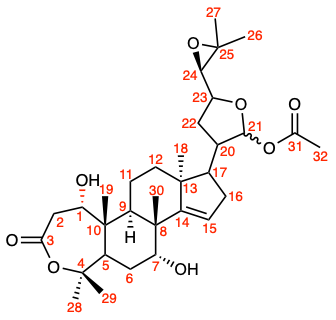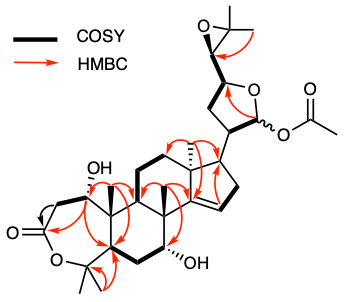 | | | | | | | |
| **C** | **^13^C δ (ppm)** | **^1^H δ**  **(ppm, J in Hz)** | | **C** | **^13^C δ**  **(ppm)** | **^1^H δ**  **(ppm, J in Hz)** | |
| **1** | 68.71 | 3.72 (1H, d J = 7.5) | | **17** | 52.53 | 1.90 (1H, m) | |
| **2** | 39.12 | 2.90  (1H, dd J = 7.5, 15.5) | 3.22  (1H, d J = 15.5) | **18** | 19.20 | 1.04 (3H, s) | |
| **3** | 170.12 | / | | **19** | 15.53 | 1.05 (3H, s) | |
| **4** | 86.37 | / | | **20** | 44.32 | 2.37 (1H, m) | |
| **5** | 41.65 | 2.70 (1H, d = 12.3) | | **21** | 96.76 | 6.24 (1H, d J = 4.0)* | |
| **6** | 27.19 | 1.82 (1H, m) | 1.97 (1H, m) | **22** | 31.48 | 1.55 (1H, m) | 2.07 (1H, m) |
| **7** | 71.53 | 3.87 (1H, br) | | **23** | 79.87 | 3.92 (1H, dt J = 10.1, 7.2) | |
| **8** | 43.88 | / | | **24** | 66.87 | 2.66 (1H, d J = 7.6) | |
| **9** | 33.61 | 2.73 (1H, dd J = 7.7, 11.5) | | **25** | 57.41 | / | |
| **10** | 45.47 | / | | **26** | 25.05 | 1.32 (3H, s) | |
| **11** | 16.43 | 1.45 (1H, m) | 1.85 (1H, m) | **27** | 19.45 | 1.28 (3H, s) | |
| **12** | 32.57 | 1.26 (1H, m) | 1.62 (1H, m) | **28** | 34.39 | 1.46 (3H, s) | |
| **13** | 46.72 | / | | **29** | 23.78 | 1.46 (3H, s) | |
| **14** | 161.92 | / | | **30** | 28.05 | 1.10 (3H, s) | |
| **15** | 119.70 | 5.48 (1H, t J = 2.4) | | **31** | 172.43 | / | |
| **16** | 35.23 | 2.21 (2H, m) | | **32** | 21.64 | 2.04 (3H, s) | |

NMR spectra were recorded in CDCl_3_, referenced to TMS and characterization was performed following the general considerations outlined. 1-hydroxyl-luvungin A (**9**) was purified as a pair of C-21 epimers in a ratio of ~5:1. The most significant difference between the two spectra was the ^1^H δ of C-21 (marked with *). The minor epimer showed a ^1^H δ of 6.28 ppm (d, J = 3.2). The absolute stereochemistry of the epimers were not resolved.

##### Table S9. ^13^C & ^1^H δ partial assignments of degraded luvungin A (7) produced using heterologously expressed genes from *C. sinensis.*.

| **Carbon numbering scheme and selected COSY and HMBC** | | | | | | |
| --- | --- | --- | --- | --- | --- | --- |
| 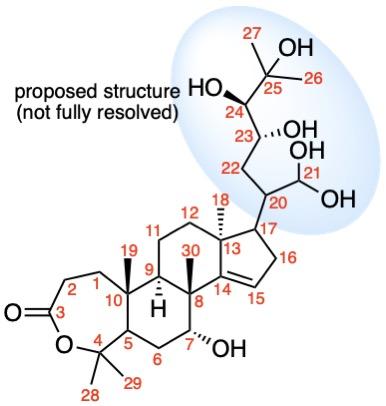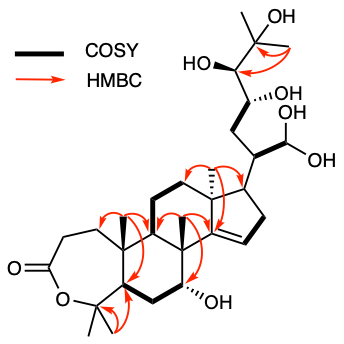 | | | | | | |
| **C** | **^13^C δ (ppm)** | **^1^H δ**  **(ppm, J in Hz)** | | **C** | **^13^C δ**  **(ppm)** | **^1^H δ**  **(ppm, J in Hz)** |
| **1** | 37.58 | 1.76 (2H, m) | | **16** | 35.04 | 2.17 (2H, m) |
| **2** | N/A | N/A | | **17** | 52.92 | N/A |
| **3** | 161.37 | / | | **18** | 20.09 | 1.02 (3H, s) |
| **4** | 85.98 | / | | **19** | 16.42 | 1.10 (3H, s) |
| **5** | 46.31 | 2.37 (1H, m) | | **20** | N/A | 2.15 (1H, m) |
| **6** | N/A | 1.81 (1H, m) | 1.88 (1H, m) | **21** | 97.09 | 5.28 (1H, s) |
| **7** | 71.58 | 3.90 (1H, s) | | **22** | N/A | 1.90 (2H, m) |
| **8** | 43.91 | / | | **23** | 78.50 | 4.46 (1H, m) |
| **9** | 41.41 | 2.65 (1H, m) | | **24** | 75.22 | 3.19 (1H, m) |
| **10** | 40.21 | / | | **25** | 73.17 | / |
| **11** | N/A | 1.58 (1H, m) | 1.75 (1H, m) | **26** | 26.71 | 1.26 (3H, s) |
| **12** | 33.17 | N/A | | **27** | 26.64 | 1.29 (3H, s) |
| **13** | 46.29 | / | | **28** | 32.05 | 1.49 (3H, s) |
| **14** | 161.12 | / | | **29** | 26.00 | 1.43 (3H, s) |
| **15** | 120.04 | 5.48 (1H, s) | | **30** | 26.86 | 1.09 (3H, s) |

NMR spectra were recorded in CDCl_3_, referenced to TMS and characterization was performed following the general considerations outlined. Proposed structure of the degraded product of luvungin (**7**) is shown, the blue shade indicates the uncertain structure moiety. Partial assignment of the degraded product was achieved through comparison with the complete NMR assignment of (**9**) (Table S8). While the exact functional groups on C-21 and C-23~25 couldn’t be fully resolved by the NMR due to overlapped signals and low signal intensities, the higher ^13^C δ of C-24,25 (75.22 and 73.17 ppm) compared to those in (**9**) (66.87 and 57.41 ppm) suggested that the C-24,25 epoxide was opened. HMBC correlation from C-28/29 to C-4 and the presence of C-3 ketone (^13^C δ = 161.12 ppm) were two key pieces of evidence supporting the A-ring lactone structure, which was further corroborated by the complete assignment of (**9**) (Table S8).

##### Table S10. Gene ID of active *Melia azedarach* limonoid biosynthetic genes in this study.

| **#** | **Name** | **GeneID (*M. azedarach* genome)** |
| --- | --- | --- |
| **1** | *MaOSC1** | MELAZ155640_EIv1_0159960.1 |
| **2** | *MaCYP71CD2* | MELAZ155640_EIv1_0070910.1 |
| **3** | *MaCYP71BQ5* | MELAZ155640_EIv1_0148050.1 |
| **4** | *MaCYP88A108* | MELAZ155640_EIv1_0061960.1 |
| **5** | *MaMOI2**** | MELAZ155640_EIv1_0192980.1 |
| **6** | *MaL21AT* | MELAZ155640_EIv1_0142070.1 |
| **7** | *MaSDR* | MELAZ155640_EIv1_0198190.1 |
| **9** | *MaCYP88A164* | MELAZ155640_EIv1_0061950.1 |
| **10** | *MaL1AT* | MELAZ155640_EIv1_0164450.1 |
| **11** | *MaL7AT* | MELAZ155640_EIv1_0235630.1 |
| **12** | *MaAKR*** | MELAZ155640_EIv1_0165520.1 |
| **13** | *MaCYP716AD4* | MELAZ155640_EIv1_0052990.1 |
| **14** | *MaLFS* | MELAZ155640_EIv1_0015190.1 |
|  | *MaMOI1* | MELAZ155640_EIv1_0192990.1 |
|  | *MaSI* | MELAZ155640_EIv1_0193000.1 |
|  | Closest *CsCYP716AC1* homolog**** | MELAZ155640_EIv1_0122250.1 |

Gene name and relevant ID from *M. azedarach* genome for all functional *M.azedarach* genes (numbered in order of reported occurrence) described in this study as well as the additional sterol isomerases and cytochrome p450s mentioned. Asterisks denote the following. (*) indicates that *MaOSC1* is the *Melia azedarach* version of a tirucalla-7,24-dien-3β-ol synthase (previously characterized (*17*)), however the *A. indica* version (AiOSC1, (*17*)) was used for all experimental work in this paper. (**) indicates *MaAKR* was identified as a candidate based on homology to *Cs*AKR, however is truncated in the *M. azedarach* genome annotation (potentially accounting for its lower ranking than other functional genes (**Figure 2C**)), a full-length copy (TRINITY_DN15268_c1_g3_i2.p1, Table S18) was identified in a transcriptome assembly constructed from *M. azedarach* petiole RNA-seq data. (***) indicates that the functional sequence for *MaMOI2,* which was cloned and used in this study, contained the first intron, in addition to the exons. Due to its functionality in *N. benthamiana* it is assumed this intron is spliced out *in planta* to achieve functionality, as the resultant protein without splicing would be truncated, the cloned sequence with intron indicated is available (Table S22). (****) indicates that this gene is truncated and not co-expressed (PCC: -0.137, Rank:15335).

##### Table S11. ^13^C & ^1^H δ assignments of *epi-*neemfruitin B (10) produced using heterologously expressed genes from *M. azedarach.*

| **Carbon numbering scheme and selected COSY and HMBC** | | | | | |
| --- | --- | --- | --- | --- | --- |
| 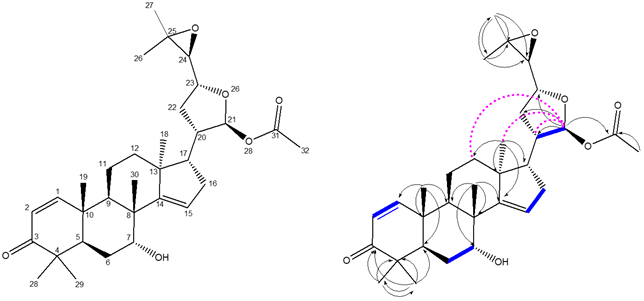 | | | | | |
| **C** | **^13^C δ**  **(150 mHz)** | **^1^H δ**  **(600 mHz)** | **C** | **^13^C δ**  **(150 mHz)** | **^1^H δ**  **(600 mHz)** |
| **3** | 205.18 | / | **20** | 44.32 | 2.38 (1H, m) |
| **31** | 170.13 | / | **10** | 40.34 | / |
| **14** | 161.60 | / | **9** | 36.61 | 2.20 (1H, m) |
| **1** | 158.14 | 7.10 (1H, d J= 10.2) | **16** | 35.23 | 2.23 (2H, m) |
| **2** | 125.71 | 5.83 (1H, d J= 10.2) | **12** | 32.41 | 1.68 (1H, m)  1.36 (1H, m) |
| **15** | 119.85 | 5.52 (1H, m) | **22** | 31.52 | 2.09 (1H, m)  1.71 (1H, m) |
| **21** | 96.72 | 6.27 (1H, d J= 4.1) | **30** | 27.87 | 1.13 (3H, s) |
| **23** | 79.90 | 3.93 (1H, m) | **29** | 27.27 | 1.16 (3H, s) |
| **7** | 71.61 | 3.99 (1H, m) | **26** | 25.07 | 1.33 (3H, s) |
| **24** | 66.83 | 2.67 (1H, d J= 7.60) | **6** | 24.39 | 1.88 (2H, m) |
| **25** | 57.35 | / | **32** | 21.67 | 2.07 (3H, s) |
| **17** | 52.79 | 1.95 (1H, m) | **28** | 21.64 | 1.09 (3H, s) |
| **13** | 46.72 | / | **18** | 19.92 | 1.03 (3H, s) |
| **8** | 44.90 | / | **27** | 19.49 | 1.29 (3H, s) |
| **5** | 44.64 | 2.39 (1H, m) | **19** | 19.04 | 1.16 (3H, s) |
| **4** | 44.34 | / | **11** | 16.42 | 1.96 (1H, m)  1.70 (1H, m) |

NMR spectra were recorded in CDCl_3_, referenced to TMS and characterization was performed following the general considerations outlined. Opposite stereochemistry at C21 to previously reported neemfruitin B assigned due to NOEs observed between C21-H and C18-H3 and C12-H2. This is consistent to those observed for 21(*S*)-acetoxyl-apomelianone (**6**) (Table S6, Fig. S15.) and different to those reported for neemfruitin B (Fig. S10.) (*33*).

##### Table S12. ^13^C δ comparison with the literature for *epi*-neemfruitin B (10) to neemfruitin B.

| **C** | ***epi-*neemfruitin B**  **this work**  **(as reported)** | ***epi-*neemfruitin B**  **this work**  **(rounded)** | **neemfruitin B literature***  **(as reported)** | **Δ** |
| --- | --- | --- | --- | --- |
| **3** | 205.18 | 205.2 | **205.8** | 0.6 |
| **31** | 170.13 | 170.1 | **170.7** | 0.6 |
| **14** | 161.60 | 161.6 | **161.9** | 0.3 |
| **1** | 158.14 | 158.1 | **161.8** | 3.7 |
| **2** | 125.71 | 125.7 | **127.2** | 1.5 |
| **15** | 119.85 | 119.9 | **119.9** | 0.0 |
| **21** | 96.72 | 96.7 | **96.8** | 0.1 |
| **23** | 79.90 | 79.9 | **80.3** | 0.4 |
| **7** | 71.61 | 71.6 | **72** | 0.4 |
| **24** | 66.83 | 66.8 | **67** | 0.2 |
| **25** | 57.35 | 57.3 | **57.8** | 0.5 |
| **17** | 52.79 | 52.8 | **53.1**** | 0.3 |
| **13** | 46.72 | 46.7 | **46.7**** | 0.0 |
| **8** | 44.90 | 44.9 | **44.8** | -0.1 |
| **5** | 44.64 | 44.6 | **44.7** | 0.1 |
| **4** | 44.34 | 44.3 | **44.7** | 0.4 |
| **20** | 44.32 | 44.3 | **44.3** | 0.0 |
| **10** | 40.34 | 40.3 | **40.5** | 0.2 |
| **9** | 36.61 | 36.6 | **36.8** | 0.2 |
| **16** | 35.23 | 35.2 | **35.4** | 0.2 |
| **12** | 32.41 | 32.4 | **33.4** | 1.0 |
| **22** | 31.52 | 31.5 | **32.3** | 0.8 |
| **30** | 27.87 | 27.9 | **27.4** | -0.5 |
| **29** | 27.27 | 27.3 | **27.1** | -0.2 |
| **26** | 25.07 | 25.1 | **25.9** | 0.8 |
| **6** | 24.39 | 24.4 | **24** | -0.4 |
| **32** | 21.67 | 21.7 | **23.2** | 1.5 |
| **28** | 21.64 | 21.6 | **21.3** | -0.3 |
| **18** | 19.92 | 19.9 | **21.2** | 1.3 |
| **27** | 19.49 | 19.5 | **19.5** | 0.0 |
| **19** | 19.04 | 19.0 | **18.9** | -0.1 |
| **11** | 16.42 | 16.4 | **16.8** | 0.4 |

Comparison of ^13^C δ values for neemfruitin B from the literature and for *epi*-neemfruitin B (**10**) for this work. Asterisks refer to the following: (*) literature assignment present in (*33*) and_._ (**) values believed to be mis-assigned in literature. Full-assignment of *epi*-neemfruitin B (**10**) is available (Table S11).

##### Table S13. ^13^C & ^1^H δ assignments of AKR product (14) produced using heterologously expressed genes from *M. azedarach.*

| **Carbon numbering scheme and selected COSY and HMBC** | | | | | |
| --- | --- | --- | --- | --- | --- |
| 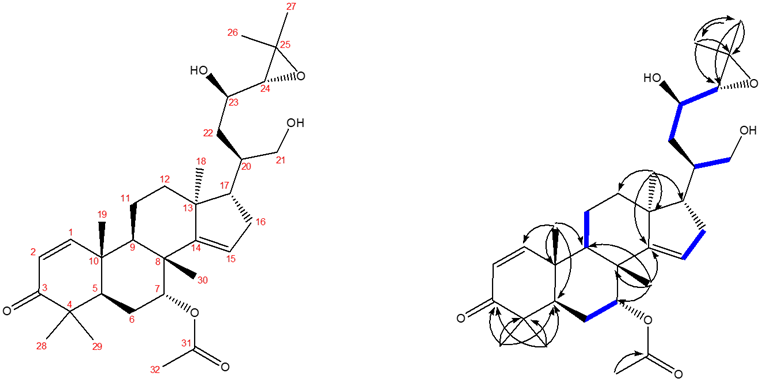 | | | | | |
| **C** | **^13^C δ**  **(150 Mhz)** | **^1^H δ**  **(600 MHz)** | **C** | **^13^C δ**  **(150 Mhz)** | **^1^H δ**  **(600 Mhz)** |
| **3** | 203.06 | / | **20** | 41.34 | 1.81 (1H, m) |
| **31** | 169.22 | / | **10** | 39.83 | / |
| **14** | 160.00 | / | **9** | 39.07 | 2.07 (1H, m) |
| **1** | 157.23 | 6.69 (1H, d J= 10.2) | **22** | 36.17 | 1.60 (1H, m)  1.47 (1H, m) |
| **2** | 125.88 | 5.90 (1H, d J= 10.2) | **16** | 35.43 | 2.08 (1H, m)  1.86 (1H, m) |
| **15** | 119.19 | 5.34 (1H, m) | **12** | 34.87 | 1.84 (1H, m)  1.52 (1H, m) |
| **7** | 74.54 | 5.29 (1H, m) | **30** | 27.39 | 0.91 (3H, s) |
| **23** | 71.28 | 3.48 (1H, m) | **29** | 27.34 | 1.15 (3H, s) |
| **24** | 67.86 | 2.64 (1H, d J= 7.9) | **26** | 24.87 | 1.09 (3H, s) |
| **21** | 64.58 | 3.94 (1H, dd J= 11.0, 3.1)  3.54 (1H, dd J= 11.0, 6.5) | **6** | 24.15 | 1.67 (1H, m)  1.57 (1H, m) |
| **25** | 58.85 | / | **28** | 21.42 | 1.00 (3H, s) |
| **17** | 56.20 | 1.73 (1H, m) | **32** | 20.83 | 1.65 (3H, s) |
| **13** | 46.86 | / | **18** | 20.05 | 1.00 (3H, s) |
| **5** | 46.65 | 2.19 (1H, dd J= 13.1, 2.5) | **27** | 19.36 | 1.10 (3H, s) |
| **4** | 44.26 | / | **19** | 18.98 | 0.81 (3H, s) |
| **8** | 43.01 | / | **11** | 16.96 | 1.52 (1H, m)  1.31 (1H, m) |

NMR spectra were recorded in benzene-d_6_, referenced to 7.16 and 128.06, following the general considerations outlined.

##### Table S14. ^1^H δ assignments of the furan moiety for kihadalactone A (19) produced using heterologously expressed genes from *C. sinensis.*

| **Carbon numbering scheme and selected COSY** | | | | |
| --- | --- | --- | --- | --- |
| 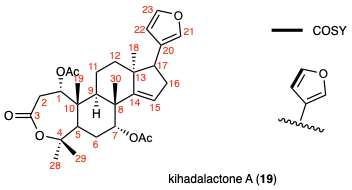 | | | | |
| **C** | **^1^H δ**  **(ppm, J in Hz)** | | **^1^H δ literature** | **Δ** |
| **21** | 7.19 (1H, m) | | 7.23 | -0.04 |
| **22** | 6.25 (1H, m) | | 6.26 | -0.01 |
| **23** | 7.37 (1H, appt t J = 1.7) | | 7.37 | 0.00 |

NMR spectra were recorded in CDCl_3_, referenced to TMS and characterization was performed following the general considerations outlined. While complete ^1^H δ assignment of kihadalactone A (**19**) was hampered by its low yield and co-eluting impurities, the signature furan moiety for limonoids was clearly distinguishable from other peaks on NMR ^1^H spectrum, and the assignment for furan moiety is shown here. The chemical shifts and coupling constant are consistent with literature values (*34*), supporting the presence of (**19**).

##### Table S15. ^13^C δ comparison with literature values for azadirone (18)

| **Carbon numbering scheme** | | | |
| --- | --- | --- | --- |
| 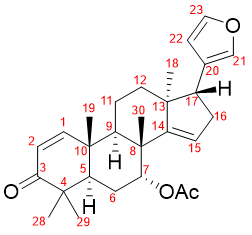 | | | |
| **C** | **Azadirone**  **this work**  **(as reported)** | **Azadirone literature**  **(as reported)** | **Δ** |
| **3** | 204.64 | 204.58 | 0.06 |
| **7αCOCH_3_** | 170.16 | 170.11 | 0.05 |
| **14** | 158.85 | 158.78 | 0.07 |
| **1** | 158.2 | 158.18 | 0.02 |
| **23** | 142.59 | 142.52 | 0.07 |
| **21** | 139.71 | 139.63 | 0.08 |
| **2** | 125.5 | 125.41 | 0.09 |
| **20** | 124.59 | 124.52 | 0.07 |
| **15** | 119.11 | 119.01 | 0.1 |
| **22** | 111.06 | 111 | 0.06 |
| **7** | 74.52 | 74.42 | 0.1 |
| **17** | 51.63 | 51.53 | 0.1 |
| **13** | 47.18 | 47.1 | 0.08 |
| **5** | 46.15 | 46.05 | 0.1 |
| **4** | 44.16 | 44.07 | 0.09 |
| **8** | 42.81 | 42.73 | 0.08 |
| **10** | 39.96 | 39.87 | 0.09 |
| **9** | 38.66 | 38.55 | 0.11 |
| **16** | 34.38 | 34.3 | 0.08 |
| **12** | 32.99 | 32.89 | 0.1 |
| **30** | 27.35 | 27.28 | 0.07 |
| **28** | 27.07 | 26.99 | 0.08 |
| **6** | 23.81 | 23.73 | 0.08 |
| **18** | 21.32 | 21.26 | 0.06 |
| **7αCOCH_3_** | 21.18 | 21.13 | 0.05 |
| **29** | 20.64 | 20.56 | 0.08 |
| **19** | 19.08 | 19.02 | 0.06 |
| **11** | 16.51 | 16.43 | 0.08 |

Comparison of ^13^C δ values for azadirone (**18**) isolated for *A. indica* leaf powder in this work (150 mHZ) with the literature assignment (*80*) (100 mHZ).

##### Table S16. ^13^C & ^1^H δ assignments of MaCYP716AD4 side-product (C24 epimeric mixture) (20) produced using heterologously expressed genes from *M. azedarach.*

| **Carbon numbering scheme and selected COSY and HMBC** | | | | | | | | | |
| --- | --- | --- | --- | --- | --- | --- | --- | --- | --- |
| 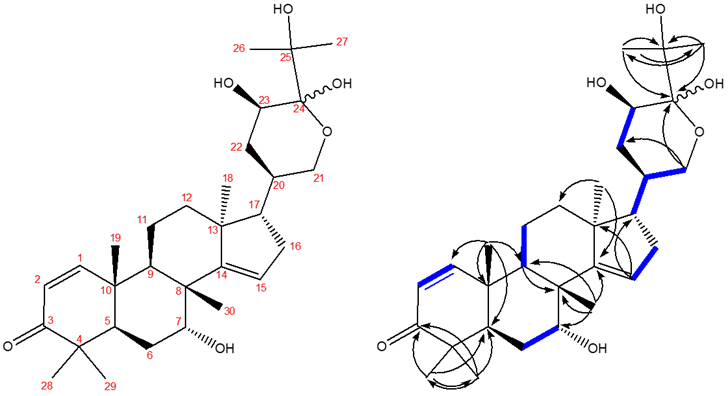 | | | | | | | | | |
| **C** | **^13^C δ**  **(150 Mhz)** | | **^1^H δ**  **(600 MHz)** | | **C** | **^13^C δ**  **(150 Mhz)** | | **^1^H δ**  **(600 Mhz)** | |
| **3** | 203.33 | | / | | **10** | 40.13 | 40.11 | / | / |
| **14** | 161.57 | 162.01 | / | / | **9** | 37.04 | 37.15 | 2.11 (1H, m) | |
| **1** | 157.00 | 157.04 | 6.64 (1H, d J=10.2) | 6.66 (1H, d J=10.2) | **16** | 34.02 | 35.03 | 1.96 (1H, m)  1.58 (1H, m) | 2.06 (1H, m)  1.68 (1H, m) |
| **2** | 125.98 | 125.97 | 5.92 (1H, d J=10.2) | 5.91 (1H, d J=10.2) | **12** | 34.14 | 34.54 | 1.74 (2H, m) | 1.79 (1H, m) |
| **15** | 119.78 | 119.91 | 5.09 (1H, brd J=2.4) | 5.11 (1H, brd J=2.4) | **22** | 33.52 | 32.50 | 1.68 (2H, m) | 1.79 (1H, m)  1.67 (1H, m) |
| **24** | 97.77 | 96.39 | / | / | **20** | 30.21 | | 2.20 (1H, m) | |
| **25** | 76.19 | 76.90 | / | / | **29** | 27.58 | | 1.33 (3H, s) | 1.32 (3H, s) |
| **7** | 71.77 | 71.90 | 3.74 (1H, brm)* | | **30** | 27.54 | | 0.85 (3H, s) | 0.84 (3H, s) |
| **23** | 67.77 | 64.23 | 3.87 (1H, m)** | 3.98 (1H, aptq J=5.5)*** | **6** | 24.80 | 24.81 | 1.79 (1H, m)  1.59 (1H, m) | |
| **21** | 65.50 | 62.24 | 3.81 (1H, dd J=11.4, 5.0)  3.60 (1H, t J= 11.4) | 3.90 (1H, dd J=11.5, 2.5)  3.52 (1H, brd J=11.5) | **26** | 24.74 | | 1.43 (3H, s) | 1.32 (3H, s) |
| **17** | 57.36 | 52.15 | 1.22 (1H, m) | 1.98 (1H, m) | **27** | 23.28 | 24.32 | 1.18 (3H, s) | 1.28 (3H, s) |
| **13** | 46.79 | 46.72 | / | / | **28** | 21.77 | 21.74 | 1.11 (3H, s) | 1.10 (3H, s) |
| **5** | 44.92 | 44.98 | 2.58 (1H, brdd J=13.0, 2.2) | | **18** | 19.35 | 19.40 | 0.86 (3H, s) | 0.68 (3H, s) |
| **8** | 44.86 | | / | | **19** | 18.95 | | 0.87 (3H, s) | 0.85 (3H, s) |
| **4** | 44.43 | | / | | **11** | 16.51 | 16.60 | 1.51 (1H, m)  1.30 (1H, m) | |

NMR spectra were recorded in benzene-d_6_, referenced to 7.16 and 128.06, following the general considerations outlined. Isolated product is a C24 epimeric mixture ca. 125 : 68 ratio. The δ for most abundant epimer is reported where a difference is observed. Asterisks indicate the following COSY coupling to OH; (*) δ 1.88, (**) δ 2.85 and (***) δ 2.61.

##### Table S17. Gene ID/Accession numbers of active *Citrus* limonoid biosynthetic genes and other *Citrus* genes in this study.

|  | **Gene Name** | **Gene ID** |
| --- | --- | --- |
| **1** | *CsOSC1* | XM_006468053 |
| **2** | *CsCYP71CD1* | XM_006467236 |
| **3** | *CsCYP71BQ4* | XM_006469432 |
| **4** | *CsCYP88A51* | XM_006485364 |
| **5** | *CSMOI1* | XM_006478528 |
| **6** | *CsMOI2* | XM_006494479 |
| **7** | *CsMOI3* | XM_006471624 |
| **8** | *CsL21AT* | XM_006482023 |
| **9** | *CsSDR* | XM_006481636 |
| **10** | *CsCYP716AC1* | XM_006464942 |
| **11** | *CsCYP88A37* | XM_006485365 |
| **12** | *CsL1AT* | XM_006478966 |
| **13** | *CsL7AT* | Cs1g05840.1 |
| **14** | *CsAKR* | XM_006492221 |
| **15** | *CsCYP716AD2* | XM_006494121 |
| **16** | *CsLFS* | Cs5g20040.1 |
| **17** | *CsSI* | XM_006478527 |

The 12 genes cloned and characterized from C. sinensis with gene ID either NCBI: PRJNA86123 (*81*) or NICCE (*22*).

##### Table S18. Full length CDS and peptide sequence of MaAKR (transcriptome derived).

| **CDS** | >MaAKR  ATGGCGAAAACAGTGAGCATTCCTTCTGTAACCCTAGGCTCAACAGGCATAACCATGCCCCTTGTTGGGTTCGGAACGGTGGAATATCCTTTATGTGAATGGTTTAAAGACGCCGTTCTCCATGCAATCAAACTCGGATACAGACACTTCGATACTGCTTCAACTTACCCTTCAGAACAGCCTCTTGGTGAAGCCATCACCGAAGCTCTCCGCCTCGGCCTCATAAAATCCCGCGACGAGCTCTTCATCACTTCCAAGCTCTGGCTCACCGATTCCTTCCCTGACCGCGTCATCCCGGCGCTGAAGAAATCTCTCAAGAATATGGGATTGGAGTACTTGGATTGTTATCTGATTCATTTTCCGGTGTGTTTGATTCCGGAGGCGACGTATCCGGTGAAGAAGGAGGATATTCGTCCGATGGATTTTGAGGGTGTGTGGGCTGCAATGGAGGAATGTCAAAAGCTTGGTCTTACCAAAACCATTGGAGTAAGCAACTTTACTGCCAAAAAACTCGAGAGGATACTTGCTACTGCAAAAATCCTTCCGGCTGTCAATCAGGTGGAGATGAACCCAGTATGGCAACAAAAGAAGCTGAGGCAGTTTTGTGAAGAAAAAGGCATACATTTCTCAGCTTTCTCTCCATTAGGAGCCGTAGGAACAGACTGGGGACATAATCGAGTCATGGAATGTGAGGTGCTGAAAGAGATTGCAAAAGCTAAAGGAAAATCACTTGCTCAGATTGCAATCCGTTGGGTTTACCAACAAGGAGTGAGTGTGATTACAAAGAGCTTTAACAAACAAAGAATGGAAGAGAACCTGGACATATTTGACTGGAAGTTGACTCCTGAAGAGCTACACAAGATTGATCAAATTCCACAGTATAGAGGAAGTCGTGGTGAGACTTTTGTTTCAGAAAATGGTCCTTACAAAACTCTTGAAGAAATGTGGGACGGAGAGATTTAA |
| --- | --- |
| **peptide** | >MaAKR  MAKTVSIPSVTLGSTGITMPLVGFGTVEYPLCEWFKDAVLHAIKLGYRHFDTASTYPSEQPLGEAITEALRLGLIKSRDELFITSKLWLTDSFPDRVIPALKKSLKNMGLEYLDCYLIHFPVCLIPEATYPVKKEDIRPMDFEGVWAAMEECQKLGLTKTIGVSNFTAKKLERILATAKILPAVNQVEMNPVWQQKKLRQFCEEKGIHFSAFSPLGAVGTDWGHNRVMECEVLKEIAKAKGKSLAQIAIRWVYQQGVSVITKSFNKQRMEENLDIFDWKLTPEELHKIDQIPQYRGSRGETFVSENGPYKTLEEMWDGEI* |

Coding sequence (cds) of cloned and full-length version of *MaAKR*, which was identified as a candidate based on homology to *Cs*AKR, however was truncated in the *M. azedarach* genome annotation. Therefore the full-length copy identified above, was sourced in a transcriptome assembly constructed *de novo* from *M. azedarach* petiole RNA-seq data (Table S2) using trinity (*67*, *68*) .

##### Table S19. List of primer pairs used to clone genes from *C. sinensis*.

| **Gene** | **Use** | **Primer Sequence (5’ to 3’)** |
| --- | --- | --- |
| *CsCYP88A51* | Fwd | ***ATTCTGCCCAAATTCGCGACCGGT***ATGGATTCGAATTTTTTGTGG |
|  | Rev | ***GAAACCAGAGTTAAAGGCCTCGAG***TCATCCGACCCTAATGACTTTTGC |
| *CsMOI1* | Fwd | ***ATTCTGCCCAAATTCGCGACCGGT***ATGAGTCATCCATATTCG |
|  | Rev | ***GAAACCAGAGTTAAAGGCCTCGAG***TCAATAAACTTTGGTCTTG |
| *CsMOI2* | Fwd | ***ATTCTGCCCAAATTCGCGACCGGT***ATGAGCCATTCATCTGGG |
|  | Rev | ***GAAACCAGAGTTAAAGGCCTCGAG***TCAACCAACCTTGGTCACC |
| *CsMOI3* | Fwd | ***ATTCTGCCCAAATTCGCGACCGGT***ATGAGTCATCCCTATTCGCC |
|  | Rev | ***GAAACCAGAGTTAAAGGCCTCGAG***TCAATAAACTTTGCTCTTGTGGTC |
| *CsL21AT* | Fwd | ***ATTCTGCCCAAATTCGCGACCGGT***ATGGATCTCCAAATCACCTGC |
|  | Rev | ***GAAACCAGAGTTAAAGGCCTCGAG***TCAAAATATGCTTGGATTAGGGGAAG |
| *CsSDR* | Fwd | ***ATTCTGCCCAAATTCGCGACCGGT***ATGAACGGCCCTTCCTCTG |
|  | Rev | ***GAAACCAGAGTTAAAGGCCTCGAG***TTACTTGATAAGACCGTAAGCCC |
| *CsCYP716AC1* | Fwd | ***ATTCTGCCCAAATTCGCGACCGGT***ATGGAATTCATTATCCTTTCCTTACTTCTTC |
|  | Rev | ***GAAACCAGAGTTAAAGGCCTCGAG***TTAATTGTTGGGATAGAGGCGAACTGG |
| *CsCYP88A37* | Fwd | ***ATTCTGCCCAAATTCGCGACCGGT***ATGGAGTTAGATTTCTCATGG |
|  | Rev | ***GAAACCAGAGTTAAAGGCCTCGAG***TTACTTGAACCCGACTACTTTTGC |
| *CsL1AT* | Fwd | ***ATTCTGCCCAAATTCGCGACCGGT***ATGGAGATCAATAACGTTTCTTCAG |
|  | Rev | ***GAAACCAGAGTTAAAGGCCTCGAG***TTAAATTAAGCTTGTATCAATAGAAGC |
| *CsL7AT* | Fwd | ***ATTCTGCCCAAATTCGCGACCGGT***ATGGAGCCTGAAATACTTTCCATAG |
|  | Rev | ***GAAACCAGAGTTAAAGGCCTCGAG***TTACCACAATGGGCATGGATC |
| *CsAKR* | Fwd | ***ATTCTGCCCAAATTCGCGACCGGT***ATGGGGACGGCCATTCCAGAG |
|  | Rev | ***GAAACCAGAGTTAAAGGCCTCGAG***TTAAATTTCTCCATCCCATATTTCCTCCACAGTTCT |
| *CsCYP716AD2* | Fwd | ***ATTCTGCCCAAATTCGCGACCGGT***ATGGAGCTCCTCCTCCTCC |
|  | Rev | ***GAAACCAGAGTTAAAGGCCTCGAG***CTAATTCTCATAGGCATAGGGATAGAGG |
| *CsLFS* | Fwd | ***ATTCTGCCCAAATTCGCGACCGGT***ATGGCTGATCATTCAACAGTAAATGG |
|  | Rev | ***GAAACCAGAGTTAAAGGCCTCGAG***TTAAACAGCTTTGTTGTCTTTCAC |
| *CsSI* | Fwd | ***ATTCTGCCCAAATTCGCGACCGGT***ATGAGCCATCCGTATGTGC |
|  | Rev | ***GAAACCAGAGTTAAAGGCCTCGAG***TCAGCGAACTTTATTCTTCTTCTGC |

Nucleotides emphasized in bold and italics consist of the 5’ overlaps designed for Gibson assembly using pEAQ-HT vector. All other nucleotides represent sequences that hybridize to the gene of interest.

##### Table S20. List of primer pairs used to clone genes from *M. azedarach.*

| **Target** | **Use** | **Sequence** |
| --- | --- | --- |
| candidate CYP88A165 | Fwd | GGGGACAAGTTTGTACAAAAAAGCAGGCTTCATGGAGTTAGATATCTTGTGG |
|  | Rev | GGGGACCACTTTGTACAAGAAAGCTGGGTTTCATTTGAGCTTGATGACTTT |
| candidate AKR | Fwd | GGGGACAAGTTTGTACAAAAAAGCAGGCTTAATGGGTGCAGTGCCTGAG |
|  | Rev | GGGGACCACTTTGTACAAGAAAGCTGGGTATTATAACTCTGCATCAAGCTG |
| candidate 2-ODD | Fwd | GGGGACAAGTTTGTACAAAAAAGCAGGCTTAATGGCAGAACGGATTGATGG |
|  | Rev | GGGGACCACTTTGTACAAGAAAGCTGGGTATCAATATTTTGTGACGTCTATTAC |
| MaSDR | Fwd | GGGGACAAGTTTGTACAAAAAAGCAGGCTTAATGAACAGTTATTCATCCGCG |
|  | Rev | GGGGACCACTTTGTACAAGAAAGCTGGGTATTAATTGATAAGATTATAAGCTTTC |
| MaL21AT | Fwd | GGGGACAAGTTTGTACAAAAAAGCAGGCTTAATGAATCTCCGAATCACTTCC |
|  | Rev | GGGGACCACTTTGTACAAGAAAGCTGGGTATCAAAGTATGGTGGGATTAGG |
| MaCYP88A108* | Fwd | GGGGACAAGTTTGTACAAAAAAGCAGGCTTAATGGAGCTAAATTTCCTGTGG |
|  | Rev | GGGGACCACTTTGTACAAGAAAGCTGGGTATCAGAAGTTCTTGACCTTGATG |
| candidate AKR | Fwd | GGGGACAAGTTTGTACAAAAAAGCAGGCTTAATGGAAGCTTTGCATCTTGG |
|  | Rev | GGGGACCACTTTGTACAAGAAAGCTGGGTATTATAACTCTGCATCAAGCTG |
| MaL1AT | Fwd | GGGGACAAGTTTGTACAAAAAAGCAGGCTTAATGGAGCTCAAGATTGTTTCTTC |
|  | Rev | GGGGACCACTTTGTACAAGAAAGCTGGGTATTAAATTATGCTTGTATCAACAGAGG |
| candidate CYP714E96 | Fwd | GGGGACAAGTTTGTACAAAAAAGCAGGCTTAATGTGCACTTCTTTAACTTTGGGG |
|  | Rev | GGGGACCACTTTGTACAAGAAAGCTGGGTATCAAATCCTCTTGACATGGAG |
| MaCYP716AD4 | Fwd | GGGGACCACTTTGTACAAGAAAGCTGGGTATTATTTGTTGTAGGGATATAGGCG |
|  | Rev | GGGGACAAGTTTGTACAAAAAAGCAGGCTTAATGGAGCTCTTCCTACCC |
| MaL7AT | Fwd | GGGGACAAGTTTGTACAAAAAAGCAGGCTTAATGGAGCCTGAAATAATTTCC |
|  | Rev | GGGGACCACTTTGTACAAGAAAGCTGGGTATCACATGGGACTTGGG |

**Table S20. List of primer pairs used to clone genes from *M. azedarach* (continued).**

| MaLFS | Fwd | GGGGACAAGTTTGTACAAAAAAGCAGGCTTAATGGCGGATCATCTGACTGC |
| --- | --- | --- |
|  | Rev | GGGGACCACTTTGTACAAGAAAGCTGGGTATTATGCTTTCTTTCCCACAG |
| candidate transferase | Fwd | GGGGACAAGTTTGTACAAAAAAGCAGGCTTAATGGAAATCAAAATTATTTC |
|  | Rev | GGGGACCACTTTGTACAAGAAAGCTGGGTATTATAATCTAGCCTTTTTTGAC |
| candidate transferase | Fwd | GGGGACAAGTTTGTACAAAAAAGCAGGCTTAATGGAAATGGAAATC |
|  | Rev | GGGGACCACTTTGTACAAGAAAGCTGGGTATTAGTTGGAAGAAGC |
| MaCYP88A164 | Fwd | GGGGACAAGTTTGTACAAAAAAGCAGGCTTAATGGGCTCAGATTTGTTGTGG |
|  | Rev | GGGGACCACTTTGTACAAGAAAGCTGGGTATCATTTAAGCTTAACGATTCTTGC |
| MaMOI2 | Fwd | GGGGACAAGTTTGTACAAAAAAGCAGGCTTAATGAGCGACTCATCATCTG |
|  | Rev | GGGGACCACTTTGTACAAGAAAGCTGGGTATCAGCGAACTTTGGTCTTG |
| MaAKR (genome) | Fwd | GGGGACAAGTTTGTACAAAAAAGCAGGCTTAATGCTAAAGACGATTG |
|  | Rev | GGGGACCACTTTGTACAAGAAAGCTGGGTATCATTCAGGAGTCAAC |
| MaAKR (transcriptome) | Fwd | GGGGACAAGTTTGTACAAAAAAGCAGGCTTAATGGCGAAAACAGTG |
|  | Rev | GGGGACCACTTTGTACAAGAAAGCTGGGTATTAAATCTCTCCGTCCC |
| MaMOI1 | Fwd | GGGGACAAGTTTGTACAAAAAAGCAGGCTTAATGAGCCATCCATATTCG |
|  | Rev | GGGGACAAGTTTGTACAAAAAAGCAGGCTTAATGACTAACCATCCATACG |
| MaSI | Fwd | GGGGACCACTTTGTACAAGAAAGCTGGGTATTAATTGGTCTTACACTTC |
|  | Rev | GGGGACCACTTTGTACAAGAAAGCTGGGTATCAGCGAACTTTGGTC |
| pDNR207  (attL sites) | Fwd | TCGCGTTAACGCTAGCAT |
|  | Rev | GTAACATCAGAGATTTTGAGACAC |
| pEAQ-HT-  DEST1  (attB sites) | Fwd | GGGGACAAGTTTGTACAAAAAAGCAGGCTTA |
|  | Rev | GGGGACCACTTTGTACAAGAAAGCTGGGTA |

All primers used in this study for the cloning of genes from *M. azedarach.* The gene or target and use forward or reverse) are all listed. Asterix (*) indicates a gene previously cloned (*17*), and here re-cloned due to extended 5’ coding sequence in the new *M. azedarach* genome.

##### Table S21. Isolera™ Prime fractionation conditions for purification of products of heterologously expressed *M. azedarach* enzymes.

| **Product** | **Cartridge/phase** | **Solvent system** | **Gradient (B %)** | **CV** | **Yield of product (mg)** |
| --- | --- | --- | --- | --- | --- |
| (**3**) | SNAP Ultra 50g (Normal) | A: Hexane B: Ethyl acetate | 6-100% | 28 | 200 |
|  | SNAP KP-Sil 25g (Normal) | A: Dichloromethane B: Methanol | 0-10% | 106 | 130 |
|  | SNAP Ultra 10g (x2) (Normal) | A: Dichloromethane B: Methanol | 4-5% | 74 | 40 |
| (**6**) | SNAP Ultra 50g (Normal) | A: Hexane  B: Ethyl acetate | 6-100% | 33 (x2) | 980 |
|  | SNAP KP-Sil 25g (Normal) | A: Hexane  B: Ethyl acetate | 25-55% | 117 | 570 |
|  | Sfär Silica D Duo 25g (Normal) | A: Hexane  B: Ethyl acetate | 28% | 46 | 470 |
|  |  |  | 28-38% | 22 |  |
|  | Sfär Silica D Duo 25g (Normal) | A: Dichloromethane  B: Methanol | 0-5% | 73 | 295 |
|  |  |  | 5-7% | 17 |  |
|  | SNAP Ultra 10g (Normal) | A: Dichloromethane  B: Methanol | 5% | 9 | 220 |
| (**10**) | Sfär Silica D Duo 200g (Normal) | A: Hexane  B: Ethyl acetate | 6-100% | 19 | 300 |
|  | SNAP KP-Sil 10g  (Normal) | A: Hexane  B: Ethyl acetate | 50-75% | 200 | 70 |
| (**14**) | Sfär Silica D Duo 200g (Normal) | A: Hexane  B: Ethyl acetate | 6-100% | 19.3 | 200 |
| (**18**) | Sfär Silica D Duo 200g (Normal) | A: Hexane  B: Ethyl acetate | 5%  5-10%  10%  10-15%  15%  15-20%  20-25%  25% | 1.5  1.5  1.5  1.5  1.5  1.5  1.5  1.5 | 400 |
| (**20**) | Sfar C18 D- Duo 120g  (Reverse) | A: Water  B: Acetonitrile | 30-100  100 | 13.4  1.2 | - |

Details of conditions used for Isolera™ Prime fractionation including: phase, column, solvent system, percentage gradient of solvent B, column volume (CV) and dry weight of resulting extract (yield). All samples were dry-loaded onto Isolera™ Prime (Biotage) using Silica gel or Celite® (Sigma-Aldrich) for normal or reverse phase respectively.

##### Table S22. Full length cloned nucleotide sequence of MaMOI2

| **Cloned nucleotide sequence** | >MaMOI2  ATGAGCGACTCATCATCTGTTCCCGTGGATTTTGTGCTAAACTTCTCAACTGCCGCCTTGCATGCTTGGAATGGCCTCAGTTTATTCTTAATCGTCTTCATCTCCTGGTTTATCTCCG***GTATGTCTGCTTATTAATCTATTAAGTACACTTCGTATATAATTCTACCTCAATCATATGTAGTTTATTGTTTGACGTGTATATCATATATCTACATATATATACGTTTGCATGAATTGATCATTGCTTGCAG***GGTTGACACAGGCGAAAACAAAAATGGACAGAGTGGTATTATGCTGGTGGGCTCTCACTGGCCTTATTCATGTCTTTCAAGAGGGTTATTATGTTTTCACTCCAGATTTATTTAAAGACGATTCTCCTAATTTTATGGCTGAAATTTGTAAGTACAATATACACATATGTGTGTATATATACATATATATATATGATTCACAATATTTATTATCTAAAGAAATGGGATATATATAAATTAAACATAAACCTGCAGGGAAAGAATACAGCAAAGGTGATTCAAGATATGCAACAAGACACACTTCAGTTCTTACCATCGAATCGATGGCTTCAGTTGTTCTGGGACCTCTTAGCCTTCTAGCAGCGTATGCTTTAGCTAAAGCGAAGTCATACAACTACATTCTTCAGTTTGGAGTCTCAATTGCGCAGCTGTATGGGGCTTGTCTATATTTCCTAAGTGCTTTCCTGGAGGGGGATAATTTTGCTTCTTCTCCGTATTTTTACTGGGCATATTACGTTGGACAAAGTAGCATCTGGGTTATAGTACCAGCACTCATAGCTATACGTTGCTGGAAAAAAATCAATGCTATTTGCTATCTTCAAGACAAGAAGAACAAGACCAAAGTTCGCTGA |
| --- | --- |

The sequence (generated by sanger sequencing) of the cloned version of *MaMOI2*, which differs from predicted sequence due to the retention of the first intron (Table S10), which is assumed to be removed by splicing in *N. benthamiana.* Intron is highlighted in bold italics.

### Captions for Data S1

#### Data S1. NMR spectra for all isolated compounds

Copies of 1D NMR (including ^1^H, ^13^C and DEPT-135 NMR) and 2D NMR (including DEPT-edited-HSQC, HMBC, COSY and NOESY or ROESY) spectra for the products isolated from heterologous expression in *N. benthamiana* of limonoid biosynthetic genes from *C. sinensis* ((**6**), (**4’**), (**9**) and (**19**)) and from *M. azedarach* ((**3**), (**6**), (**10**), (**14**) and (**20**)). Along with the ^13^C NMR spectra of (**18**) isolated from *A. indica.*
