## Supplemental NMR Data for "Complex scaffold remodeling in plant triterpene biosynthesis"

Data S1- NMR spectra for all isolated compounds

**Contents**

Figure 1. NMR spectra of apo-melianol (**3**) (*M. azedarach*).	**3**

Figure 2. NMR Assignment of (**6**) (*C. sinensis*).	**3**

Figure 3. ^1^H spectra for (**6**) (*C. sinensis*).	**4**

Figure 4. ^13^C spectra for (**6**) (*C. sinensis*).	**5**

Figure 5. COSY spectra for (**6**) (*C. sinensis*).	**6**

Figure 6. HSQC spectra for (**6**) (*C. sinensis*).	**7**

Figure 7. HMBC spectra for (**6**) (*C. sinensis*).	**8**

Figure 8. NMR Assignment of (**4’**) (*C. sinensis*).	**9**

Figure 9. ^1^H spectra for (**4’**) (*C. sinensis*).	**10**

Figure 10. ^13^C spectra for (**4’**) (*C. sinensis*).	**11**

Figure 11. COSY spectra for (**4’**) (*C. sinensis*).	**12**

Figure 12. HSQC spectra for (**4’**) (*C. sinensis*).	**13**

Figure 13. HMBC spectra for (**4’**) (*C. sinensis*).	**14**

Figure 15. NMR assignment of degraded luvungin A (**7**) (*C. sinensis*).	**16**

Figure 16. ^1^H spectra for degraded (**7**) (*C. sinensis*).	**17**

Figure 17. ^13^C spectra for degraded (**7**) (*C. sinensis*).	**18**

Figure 18. COSY spectra for degraded (**7**) (*C. sinensis*).	**19**

Figure 19. HSQC spectra for degraded (**7**) (*C. sinensis*).	**20**

Figure 20. HMBC spectra for degraded (**7**) (*C. sinensis*).	**21**

Figure 21. NMR assignment of 1-hydroxyl luvungin A (**9**) (*C. sinensis*).	**22**

Figure 22. ^1^H spectra for (**9**) (*C. sinensis*).	**23**

Figure 23. ^13^C spectra for (**9**) (*C. sinensis*).	**24**

Figure 24. COSY spectra for (**9**) (*C. sinensis*).	**25**

Figure 25. HSQC spectra for (**9**) (*C. sinensis*).	**26**

Figure 26. HMBC spectra for (**9**) (*C. sinensis*).	**27**

Figure 27. NMR spectra of epi-neemfruitin B (**10**) (*M. azedarach*).	**28**

Figure 28. NMR spectra of AKR Product (**14**) (*M. azedarach*).	**29**

Figure 29. Partial NMR assignment for kihadalactone A (**19**).	**30**

Figure 30. ^1^H spectra for kihadalactone A (**19**).	**31**

Figure 32. NMR spectra of *Ma*CYP716AD4 side-product (**20**) (*M. azedarach*).	**33**

Figure 33. ^13^C NMR spectra of analytical standard of azadirone (**18**).	**34**

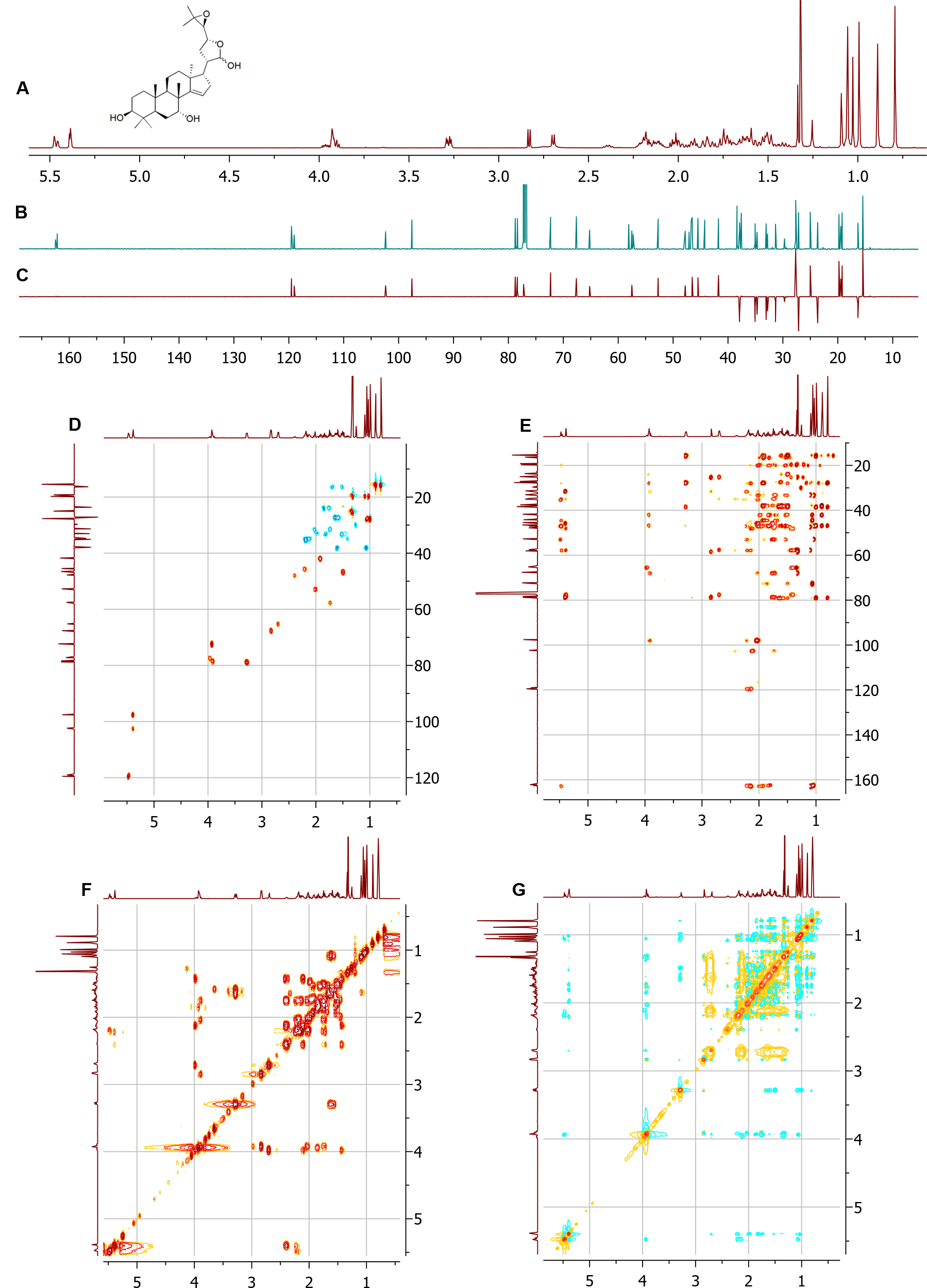

### Figure 1. NMR spectra of apo-melianol (3) (*M. azedarach*).

NMR spectra of C21 epimeric mixture ([CDCl3], δ (ppm)). (A) 1H. (B) 13C. (C) DEPT-135. (D) DEPT-edited-HSQC. (E) HMBC. (F) COSY. (G) NOESY*.*

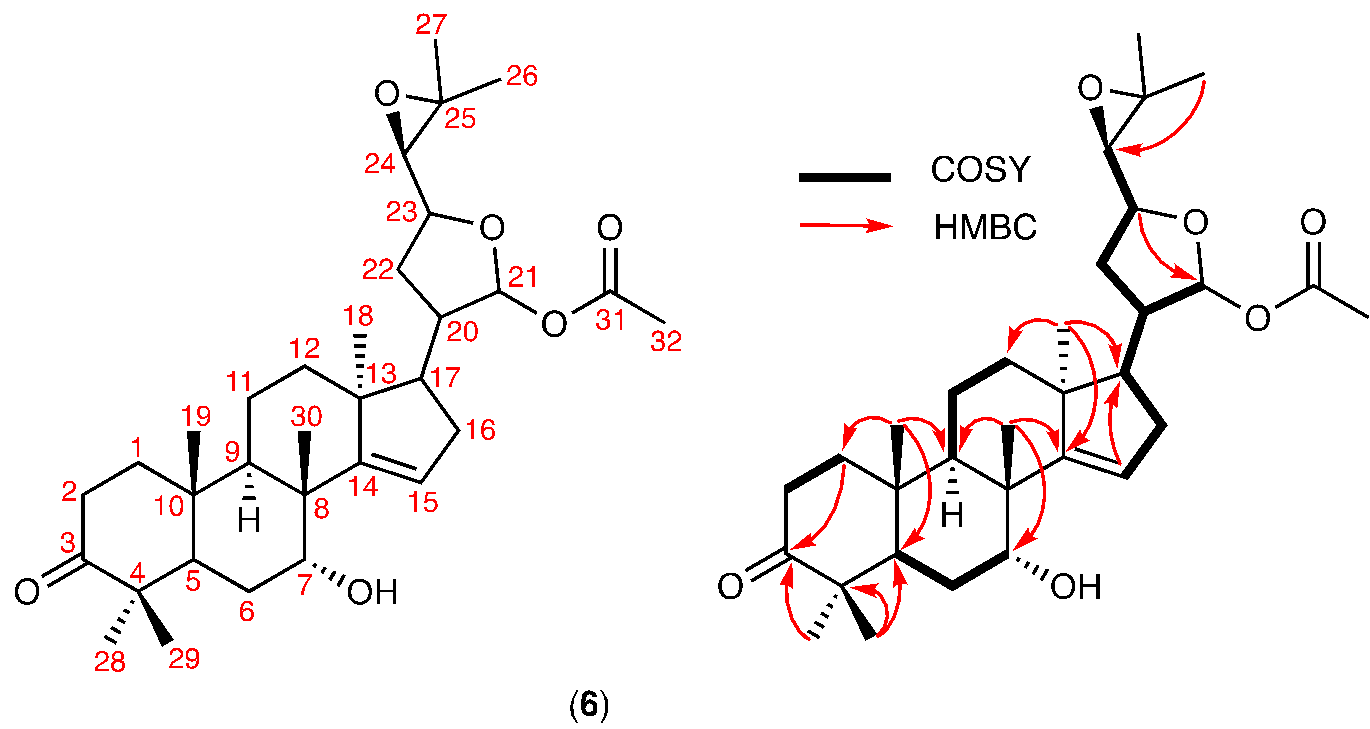

### Figure 2. NMR Assignment of (6) (*C. sinensis*).

Assignment based on NMR spectra ([CDCl_3_], δ (ppm)) listed in Figure 3-7.

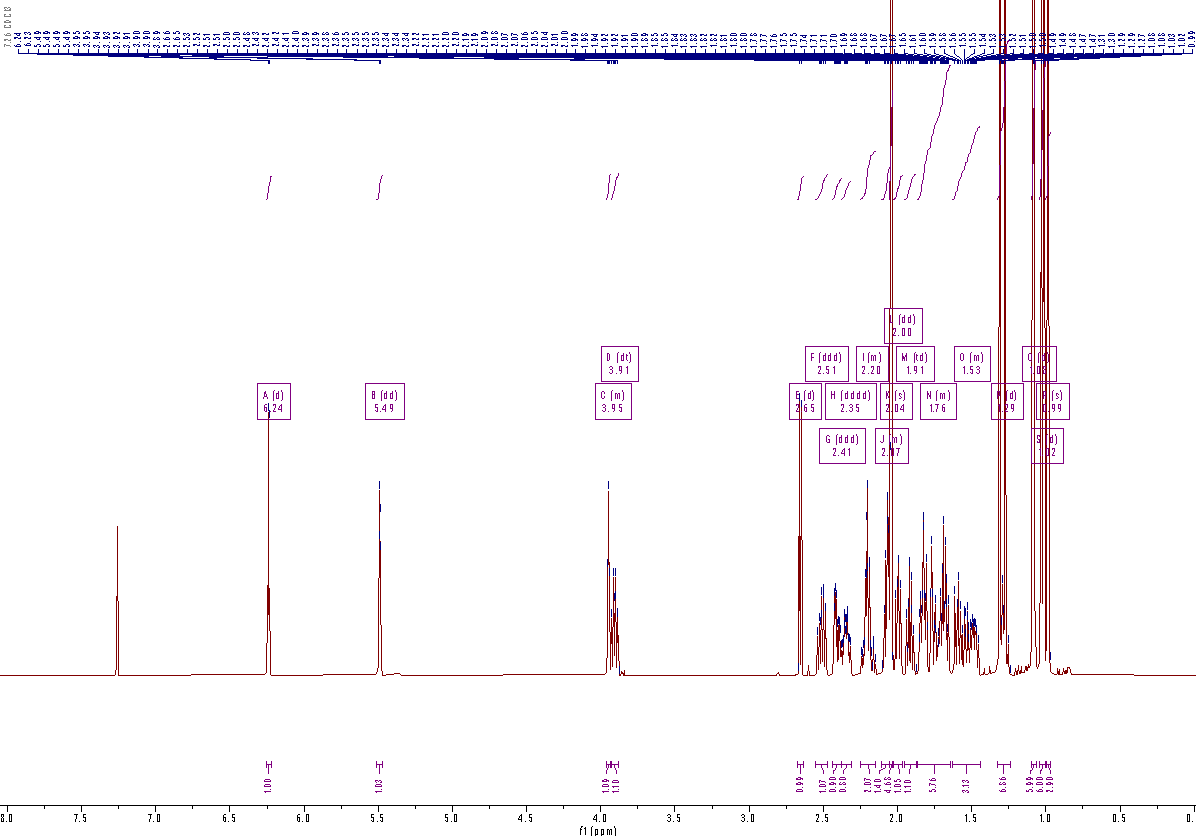

### Figure 3. ^1^H spectra for (6) (*C. sinensis*).

.

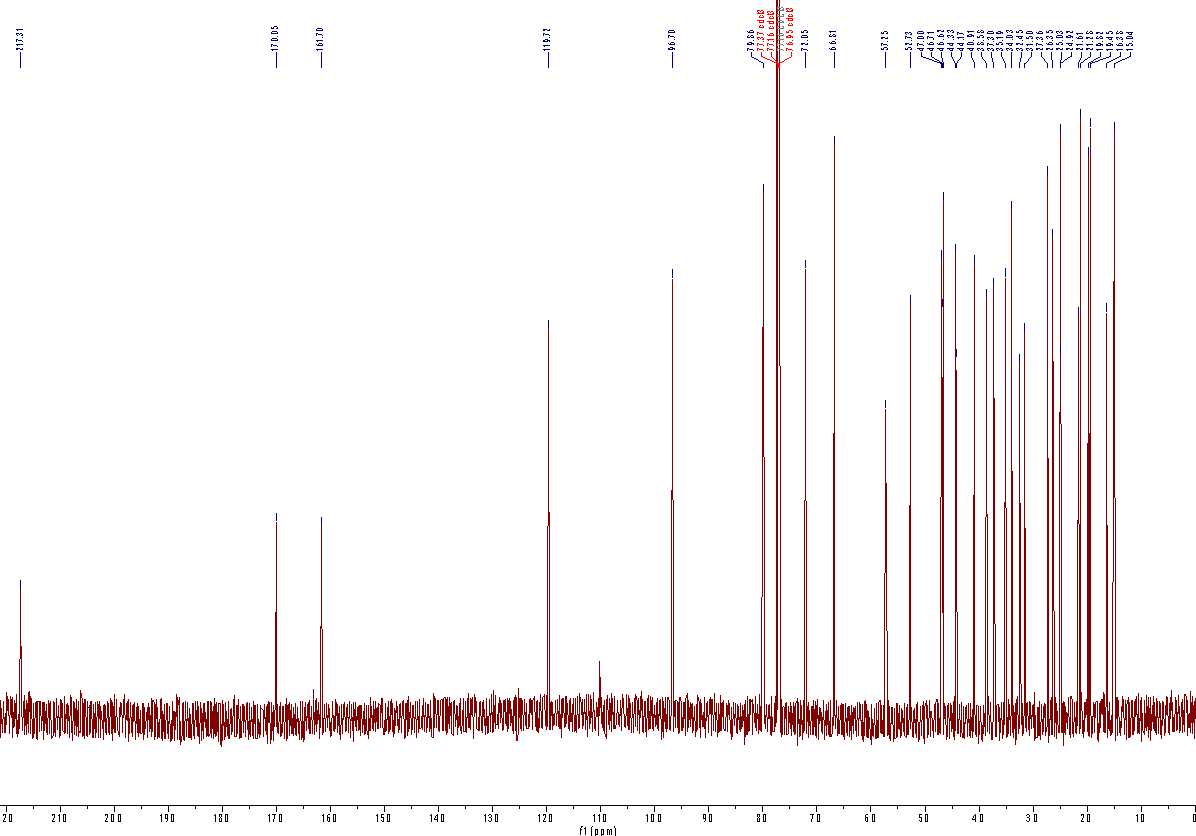

### Figure 4. ^13^C spectra for (6) (*C. sinensis*).

.

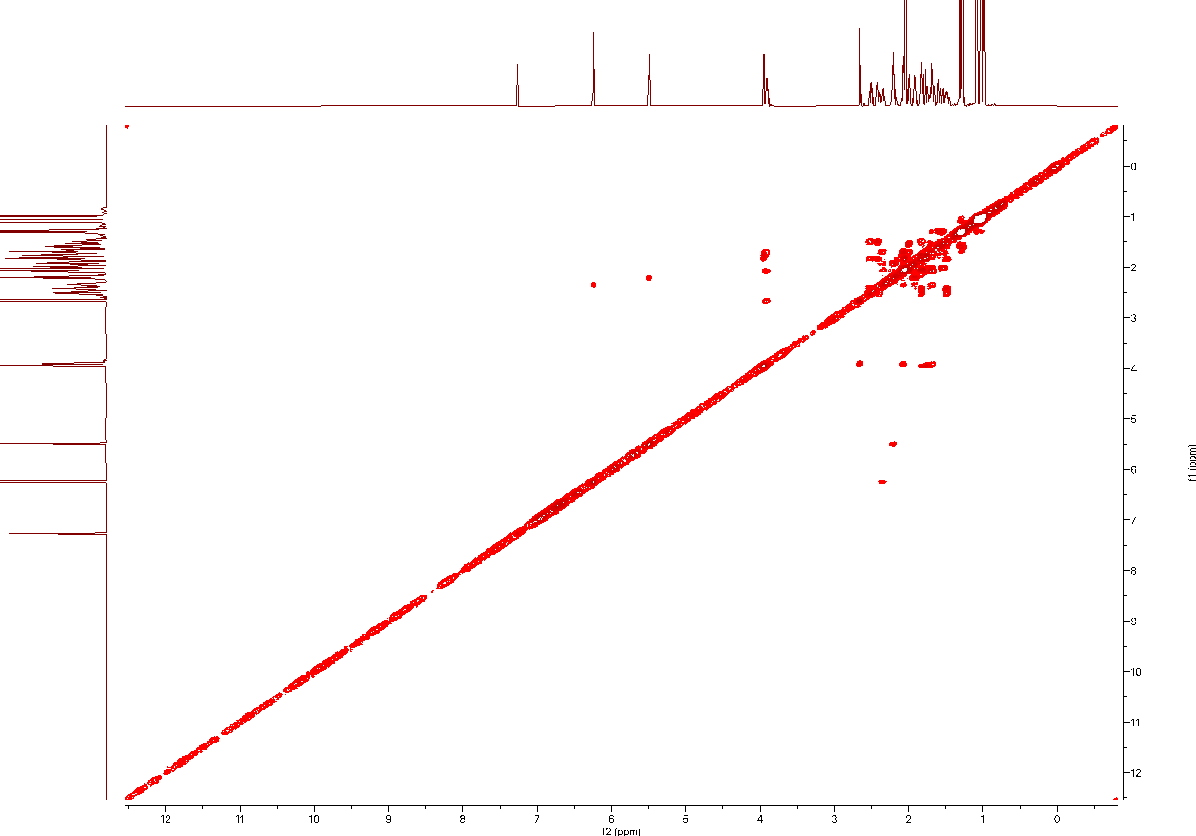

### Figure 5. COSY spectra for (6) (*C. sinensis*).

.

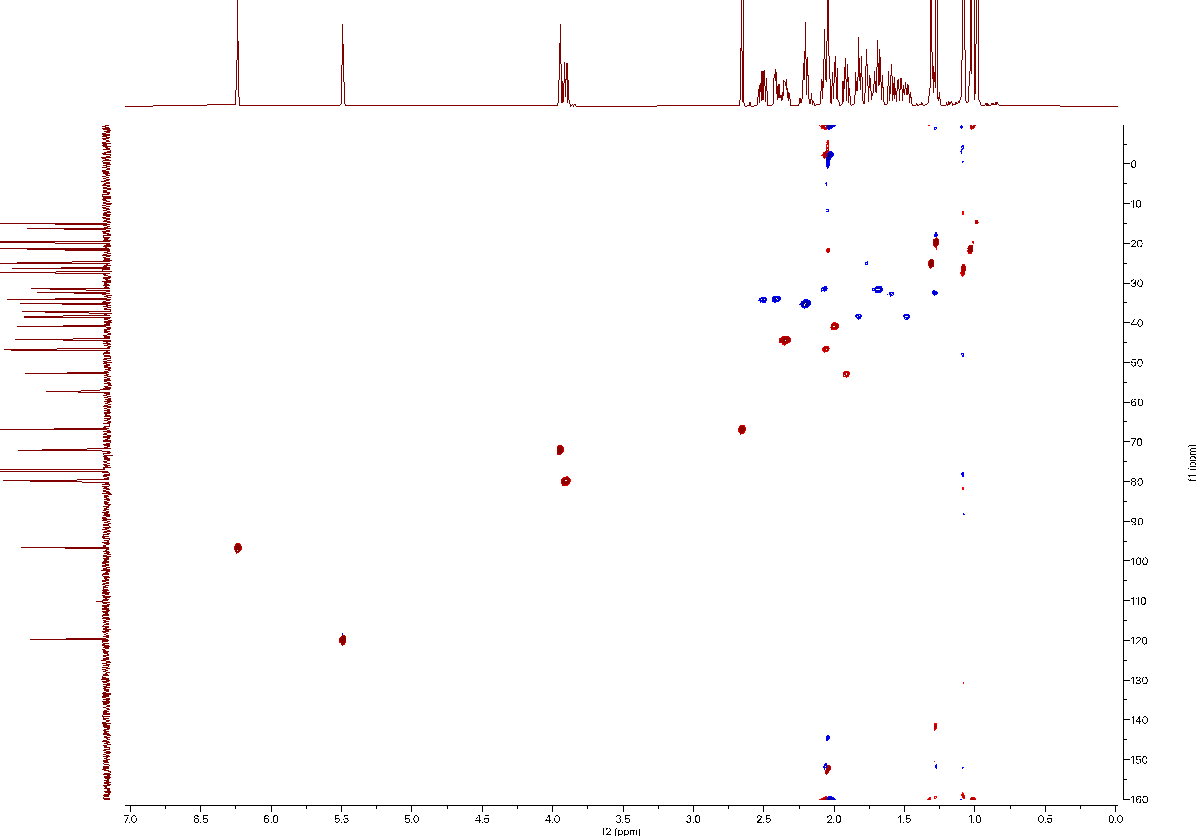

### Figure 6. HSQC spectra for (6) (*C. sinensis*).

.

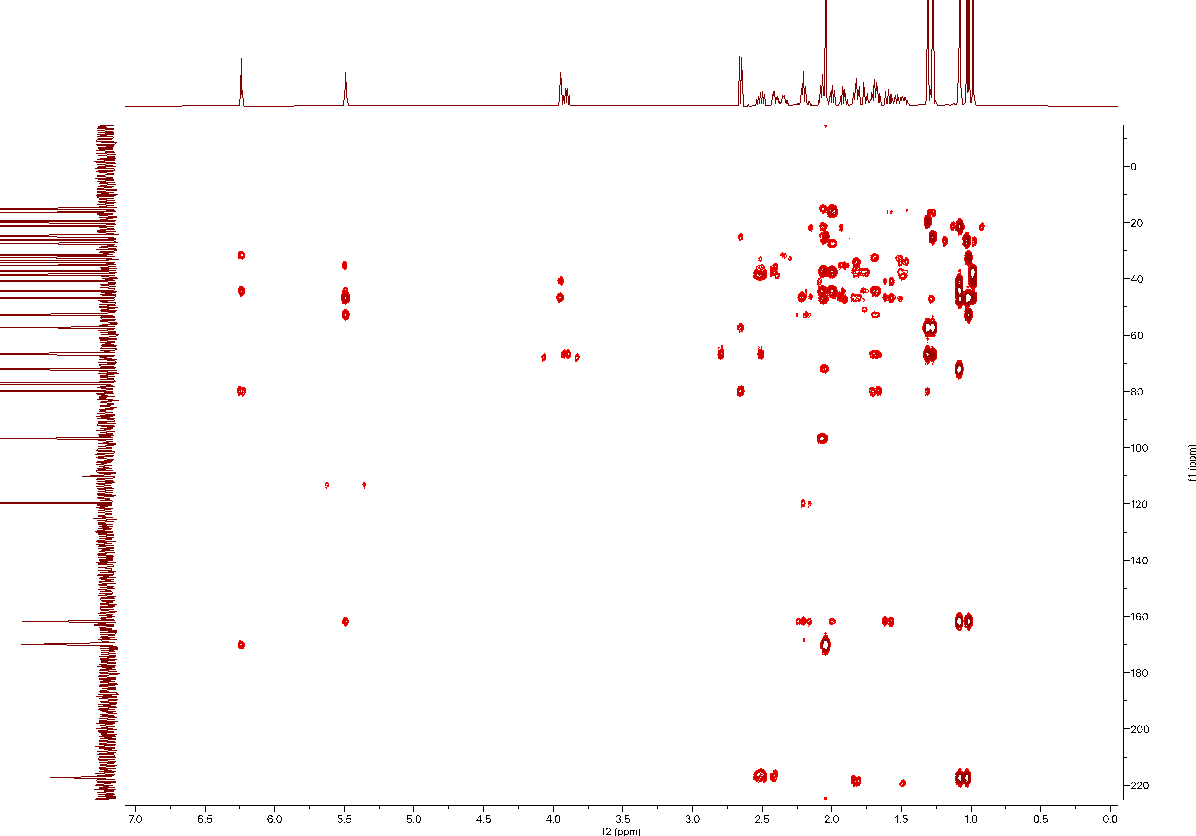

### Figure 7. HMBC spectra for (6) (*C. sinensis*).

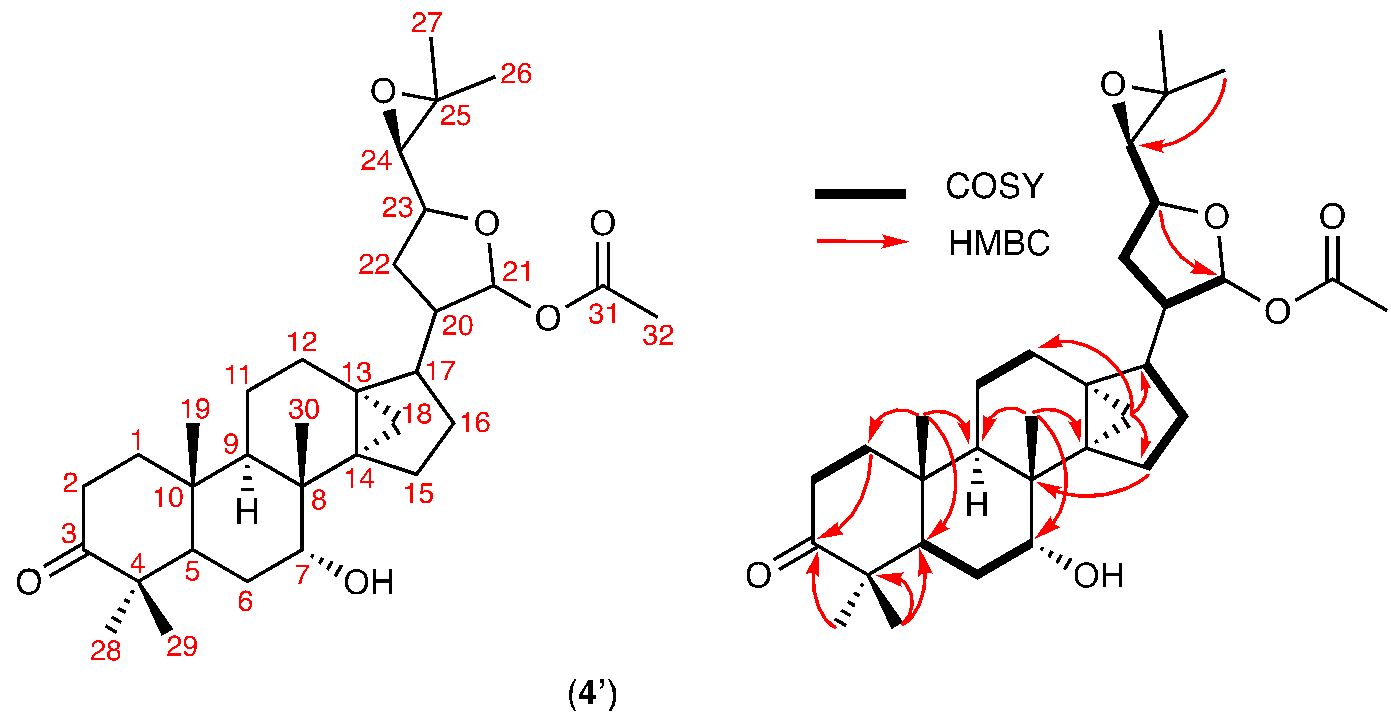

### Figure 8. NMR Assignment of (4’) (*C. sinensis*).

Assignment based on NMR spectra ([CDCl_3_], δ (ppm)) listed in Figure 9-13.

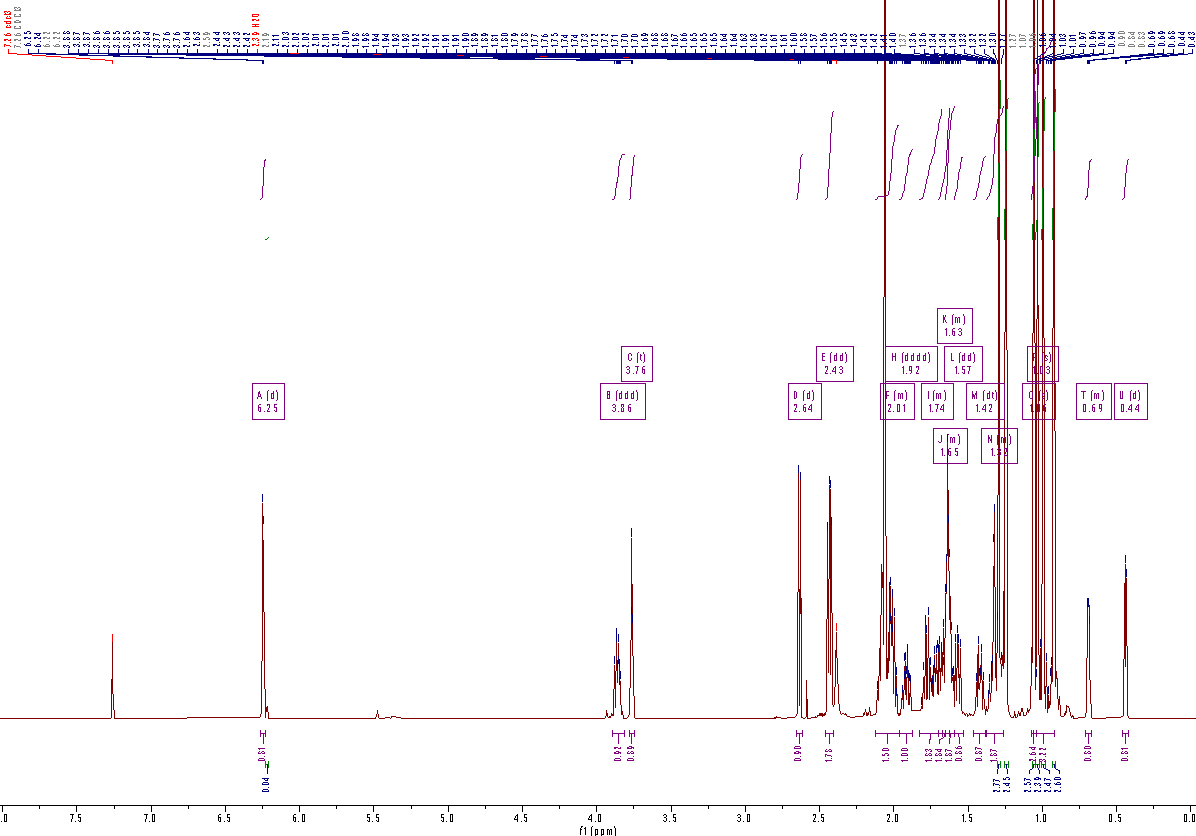

### Figure 9. ^1^H spectra for (4’) (*C. sinensis*).

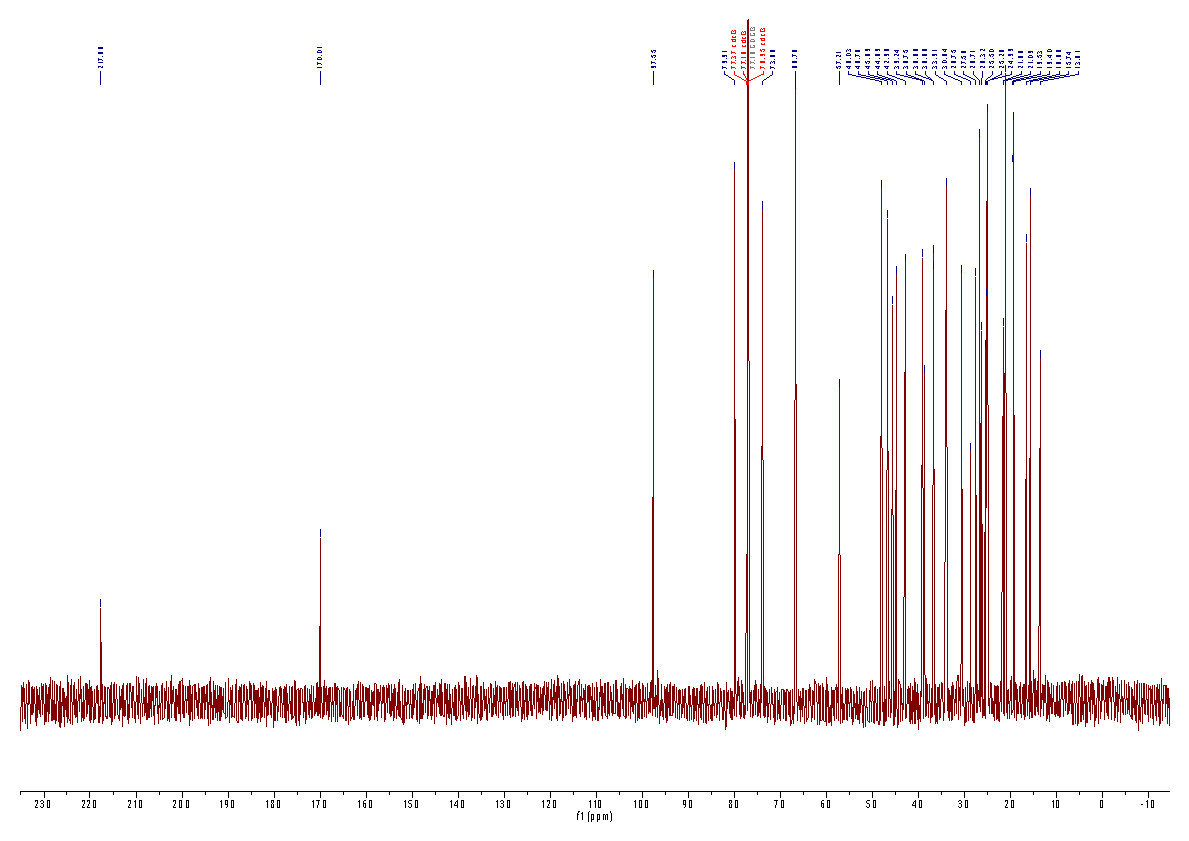

### Figure 10. ^13^C spectra for (4’) (*C. sinensis*).

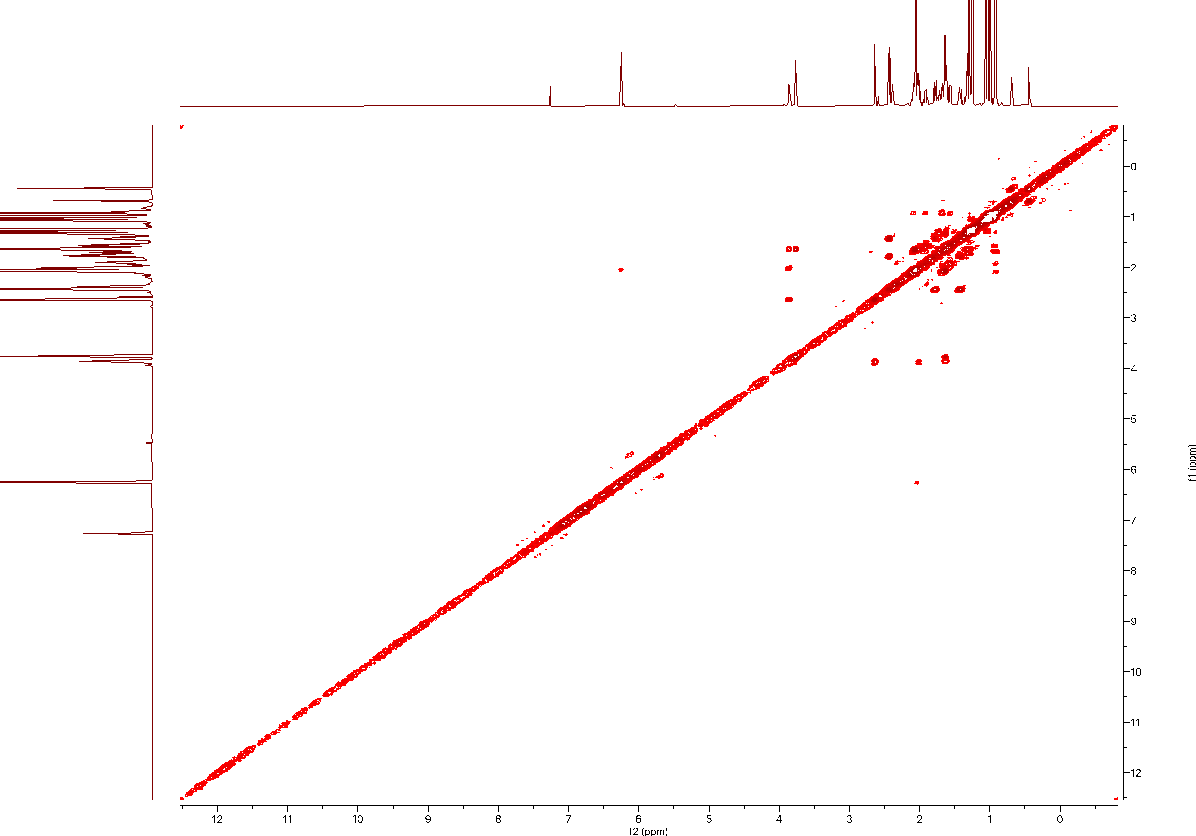

### Figure 11. COSY spectra for (4’) (*C. sinensis*).

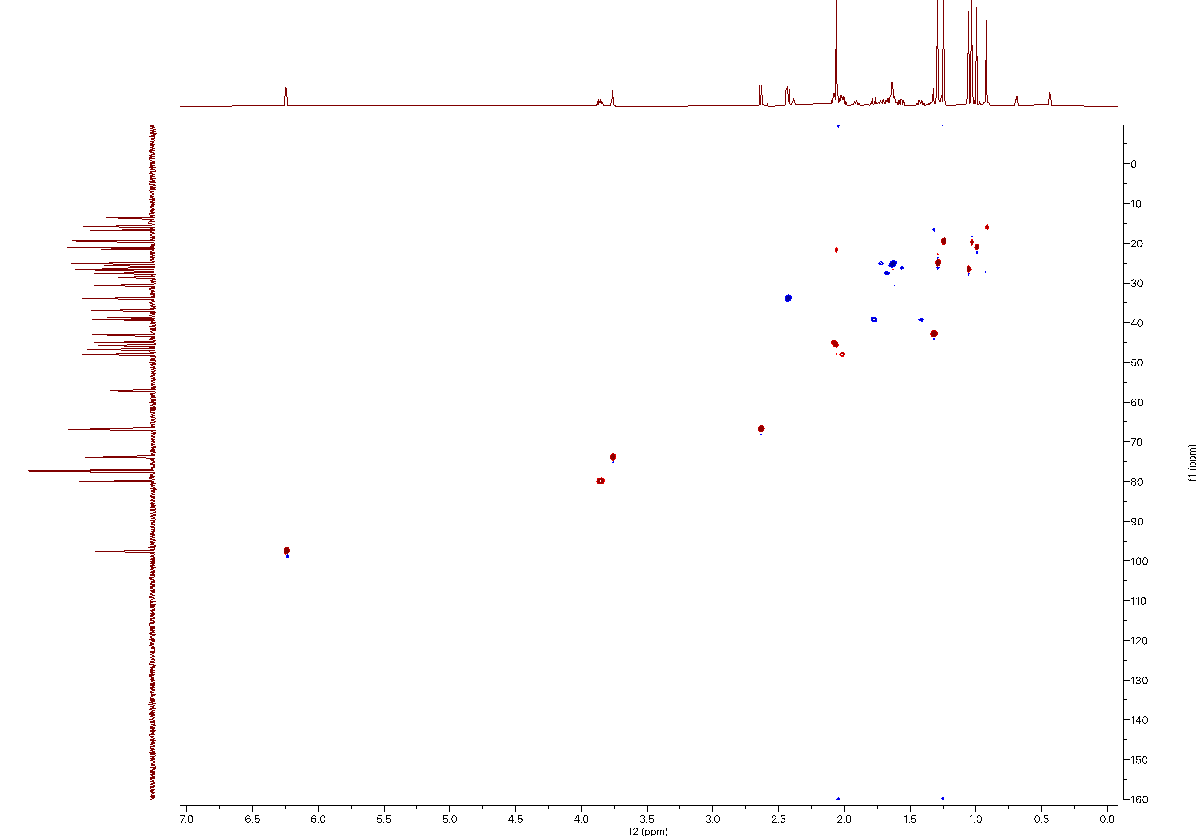

### Figure 12. HSQC spectra for (4’) (*C. sinensis*).

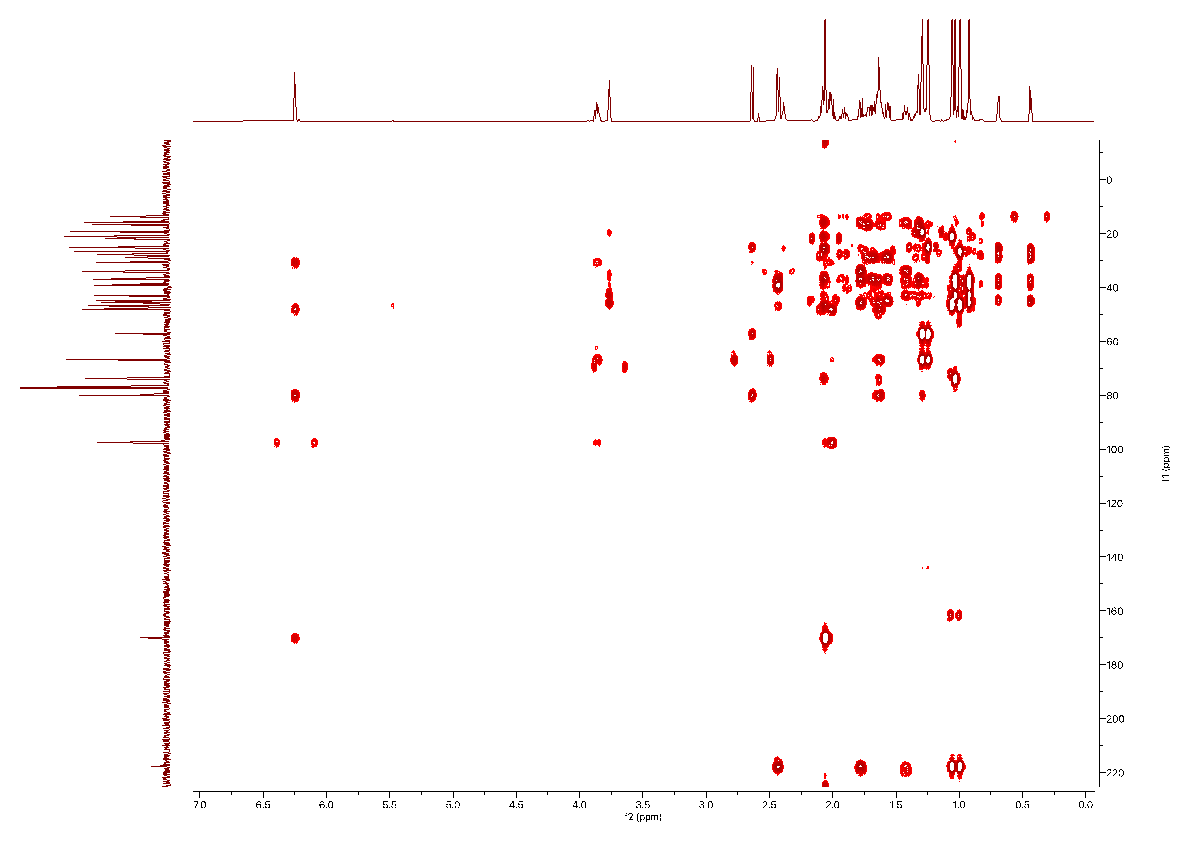

### Figure 13. HMBC spectra for (4’) (*C. sinensis*).

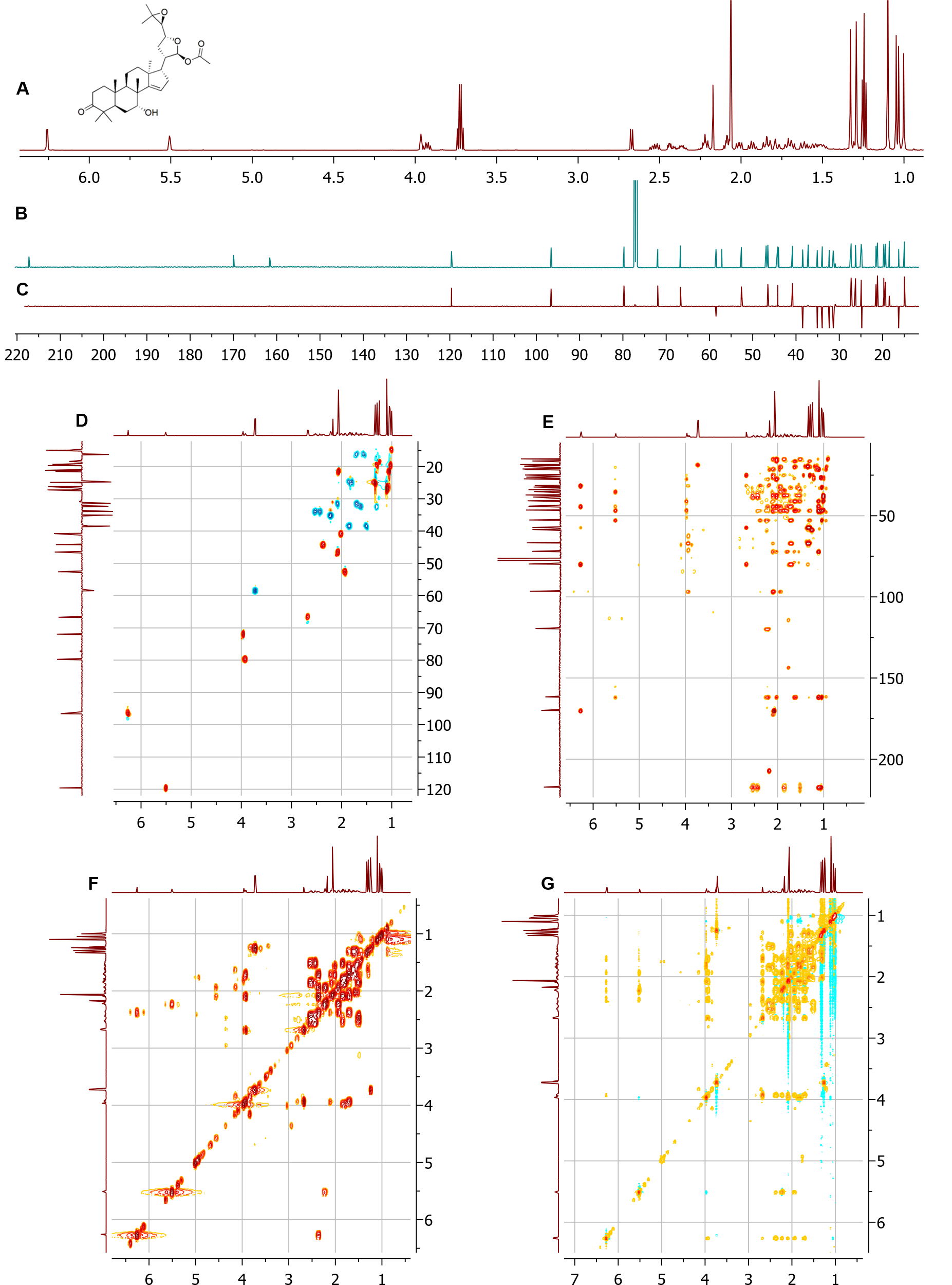

**Figure 14. NMR spectra of 21(*S*)-acetoxyl-apo-melianone (6) (*M. azedarach*).**

NMR spectra **(**[CDCl_3_], δ (ppm)).(A) ^1^H. (B) ^13^C. (C) DEPT-135. (D) DEPT-edited-HSQC. (E) HMBC. (F) COSY. (G) NOESY.

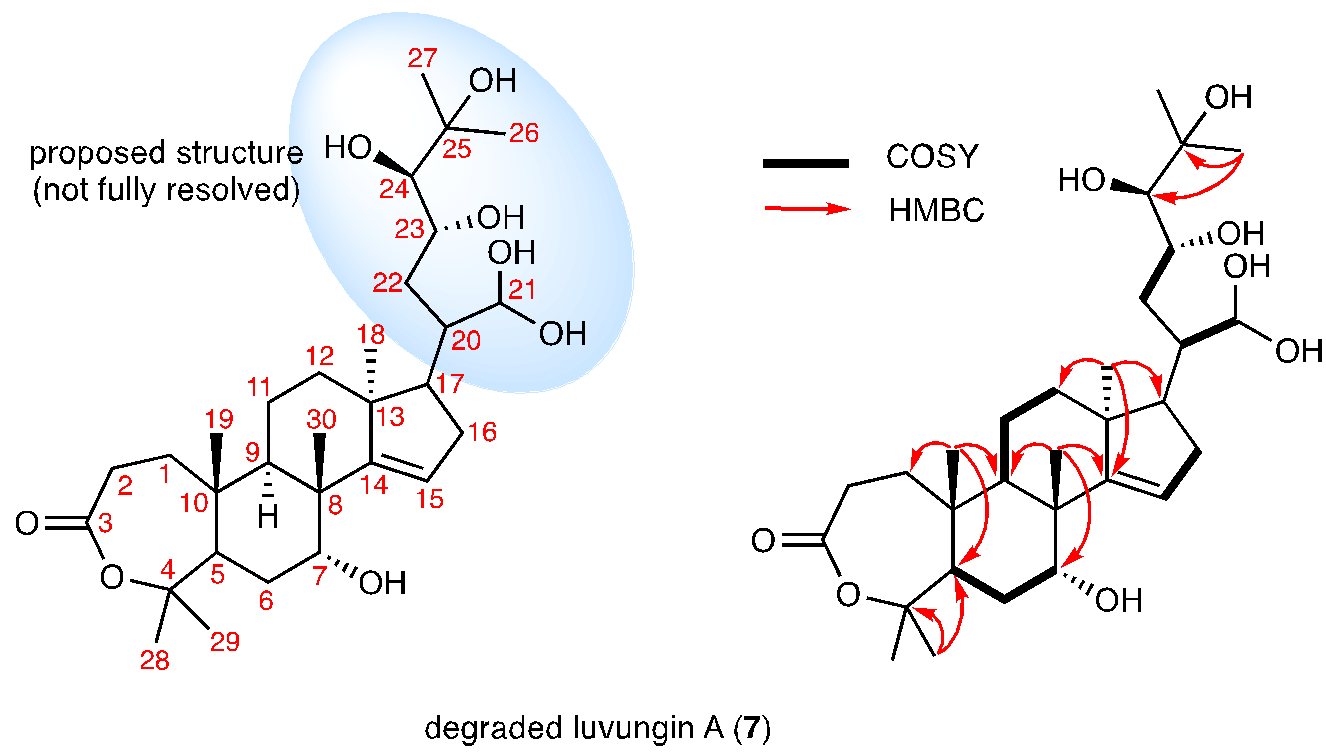

### Figure 15. NMR assignment of degraded luvungin A (7) (*C. sinensis*).

Assignment based on NMR spectra ([CDCl_3_], δ (ppm)) listed in Figure 16-20.

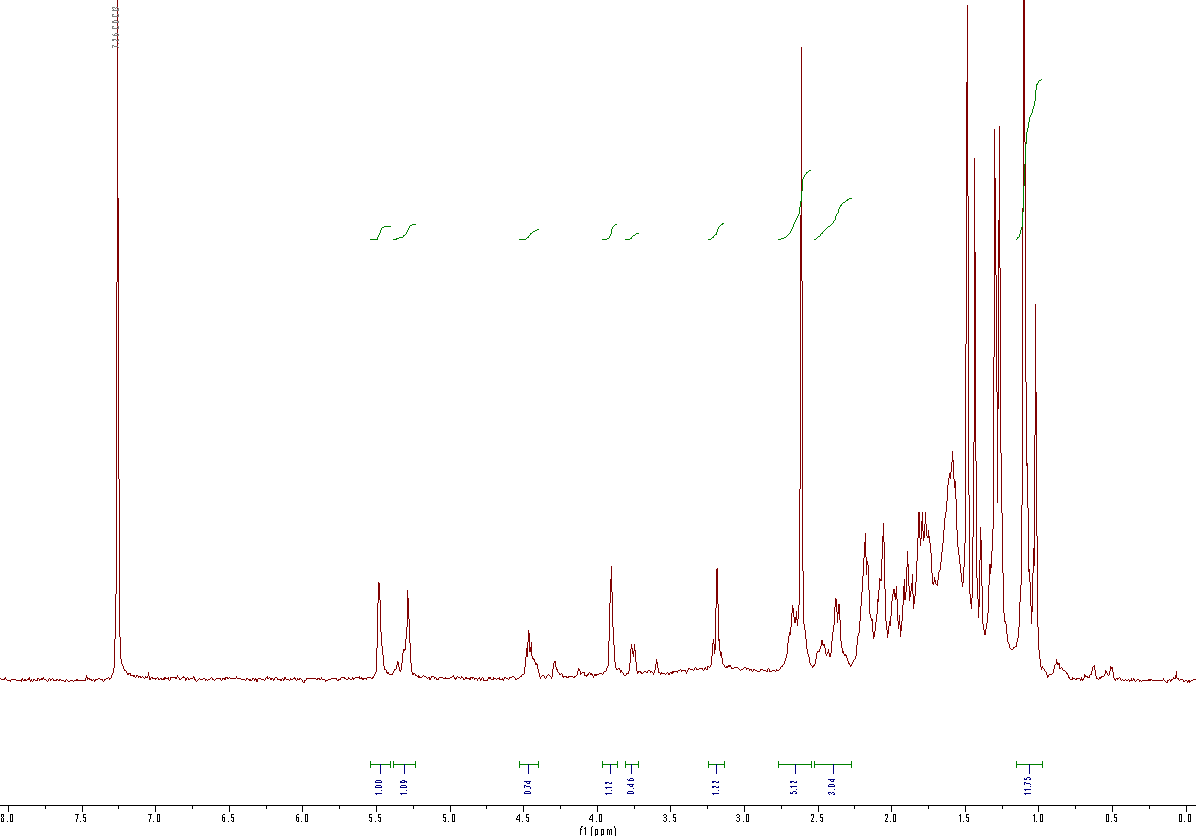

### Figure 16. ^1^H spectra for degraded (7) (*C. sinensis*).

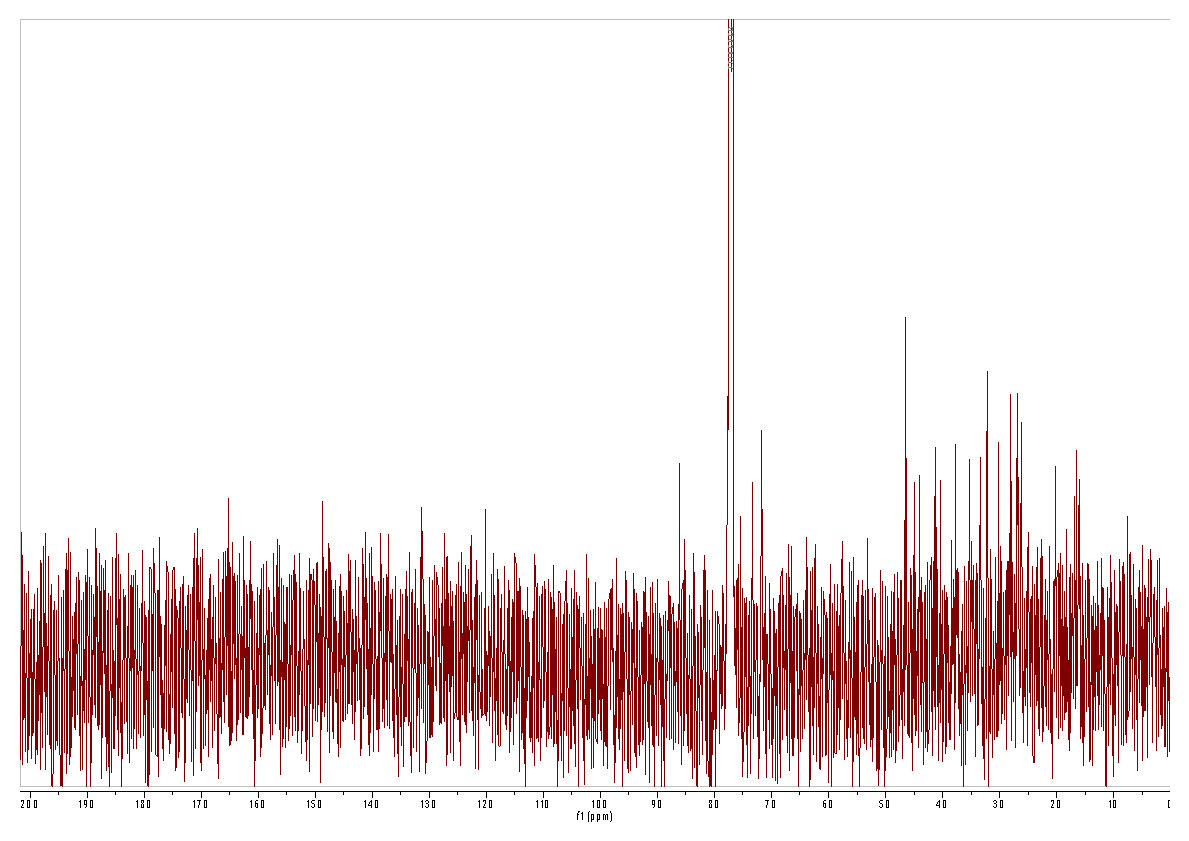

### Figure 17. ^13^C spectra for degraded (7) (*C. sinensis*).

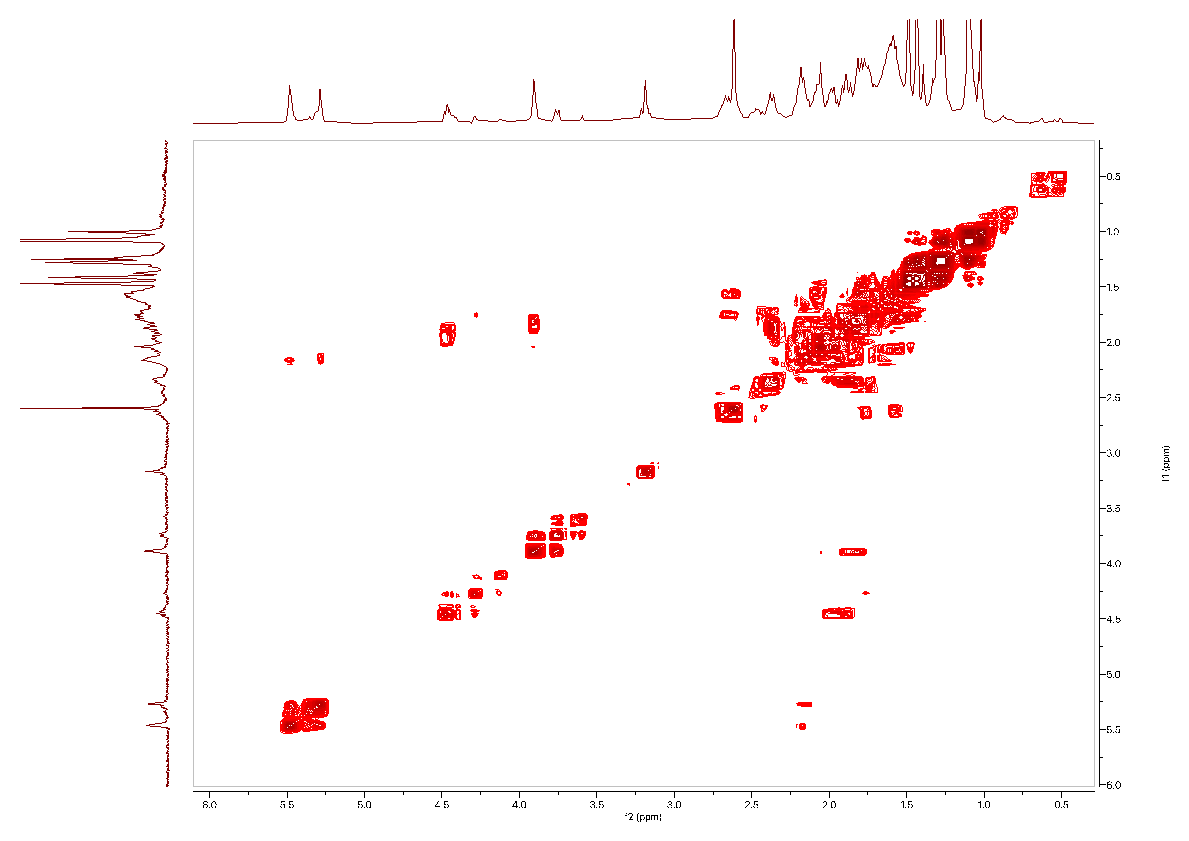

### Figure 18. COSY spectra for degraded (7) (*C. sinensis*).

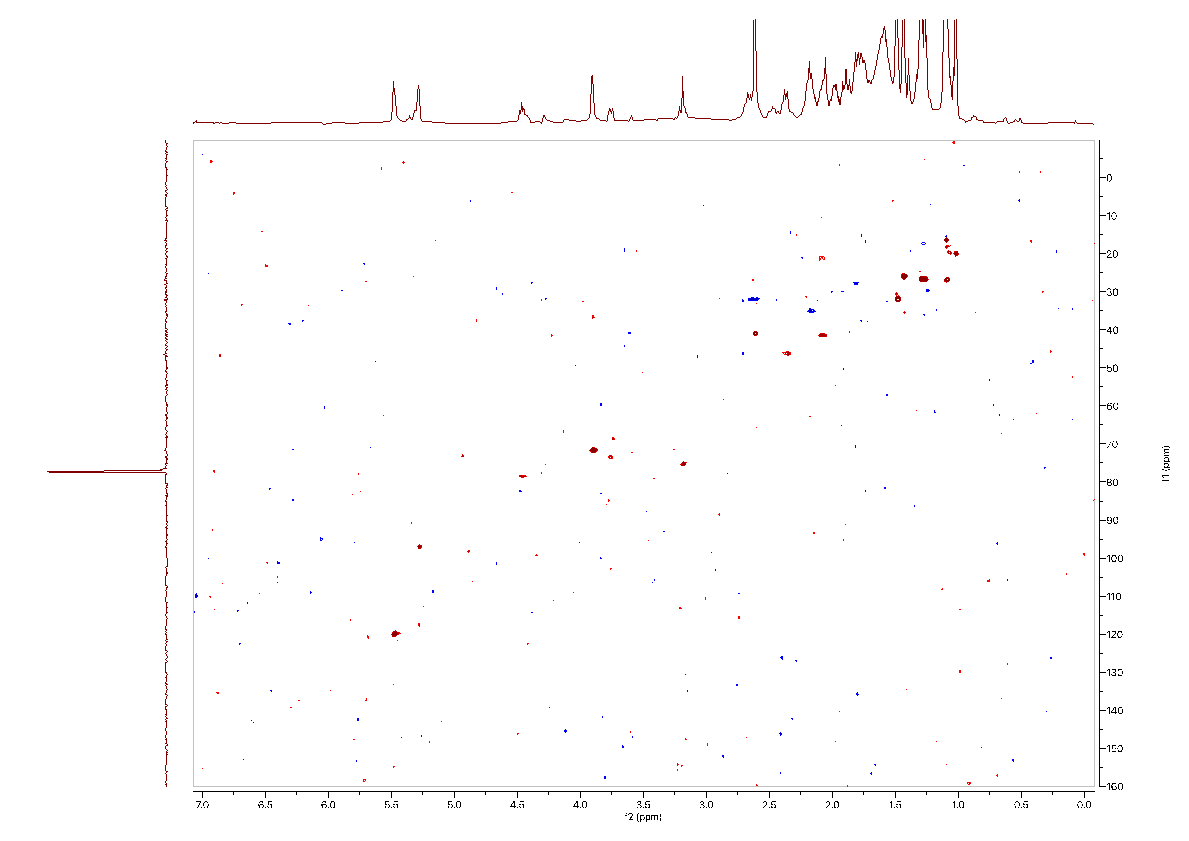

### Figure 19. HSQC spectra for degraded (7) (*C. sinensis*).

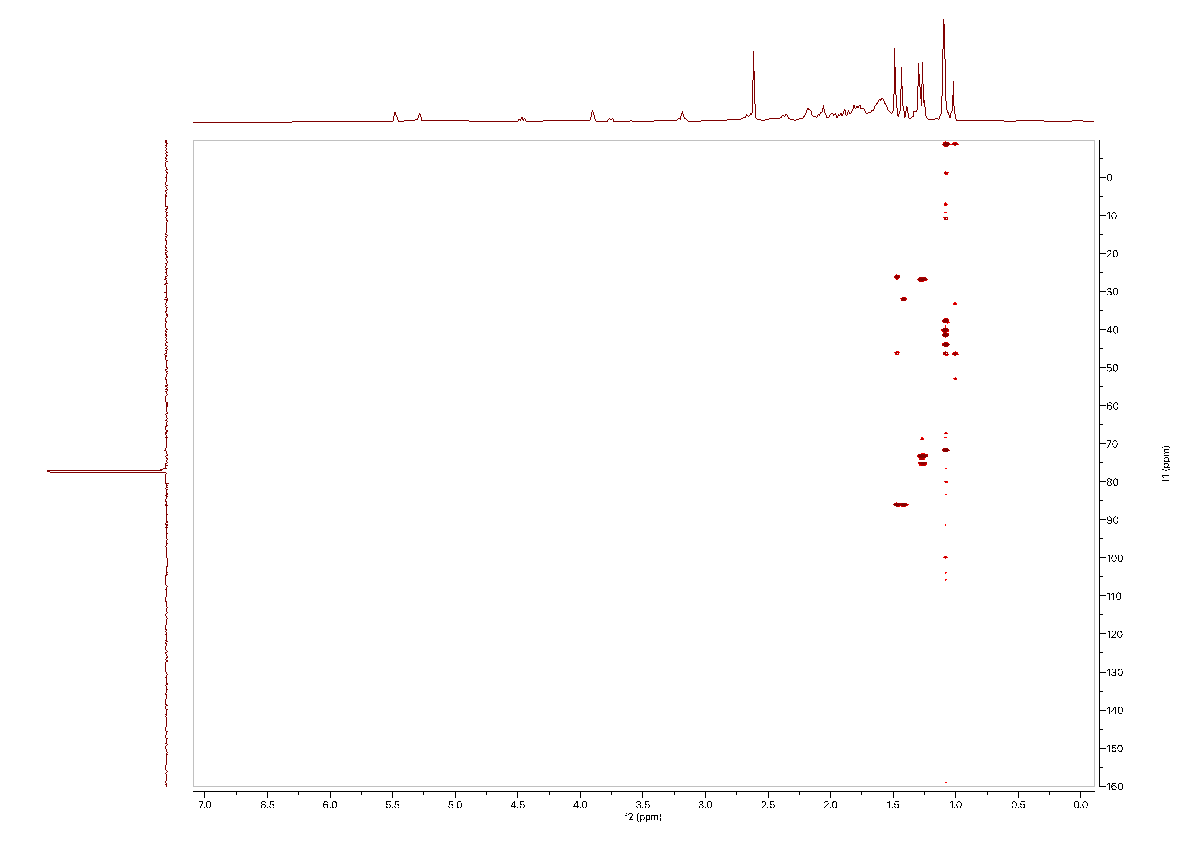

### Figure 20. HMBC spectra for degraded (7) (*C. sinensis*).

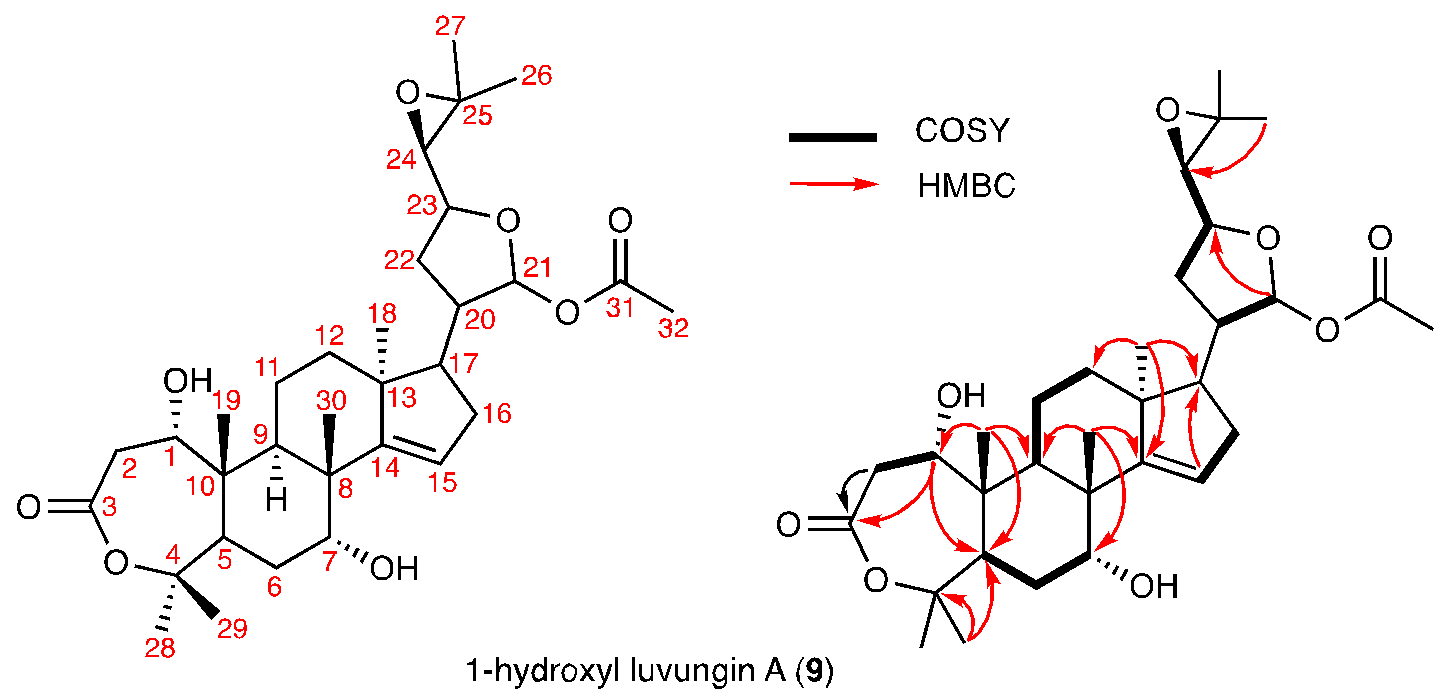

### Figure 21. NMR assignment of 1-hydroxyl luvungin A (9) (*C. sinensis*).

Assignment based on NMR spectra ([CDCl_3_], δ (ppm)) listed in Figure 22-26.

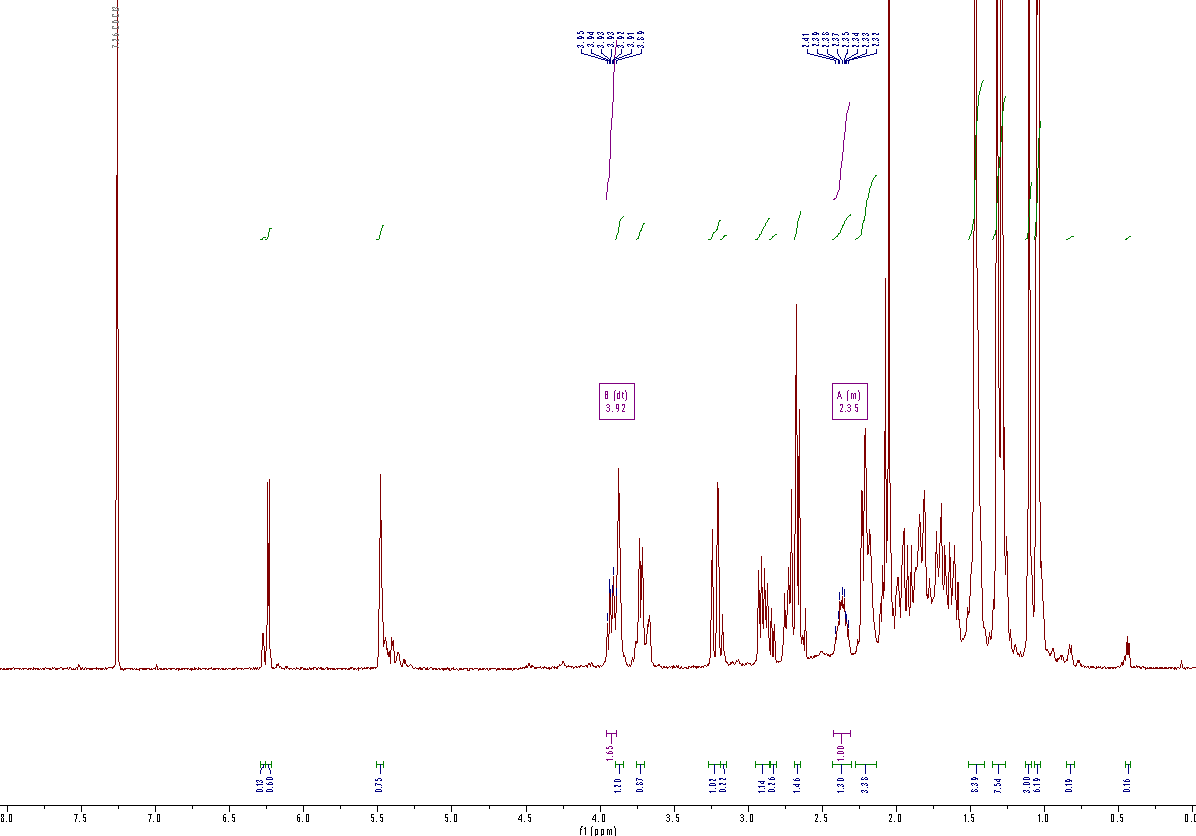

### Figure 22. ^1^H spectra for (9) (*C. sinensis*).

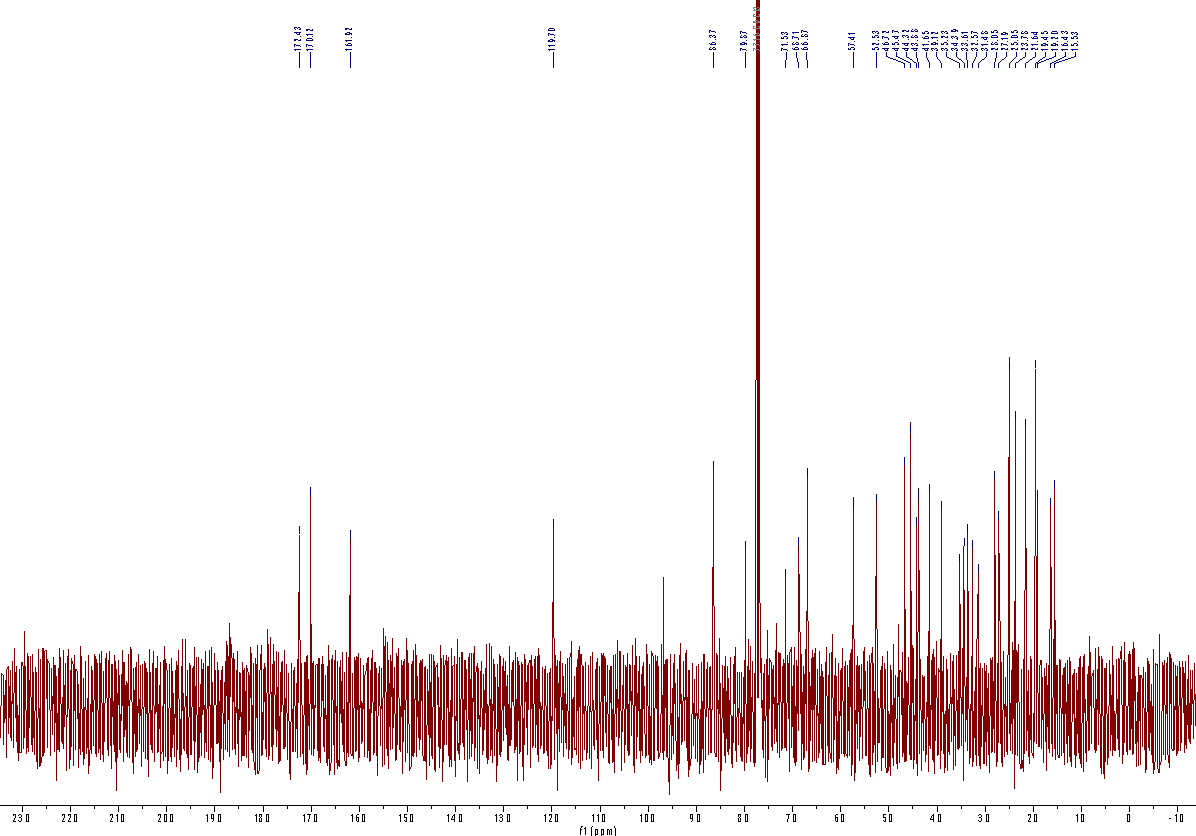

### Figure 23. ^13^C spectra for (9) (*C. sinensis*).

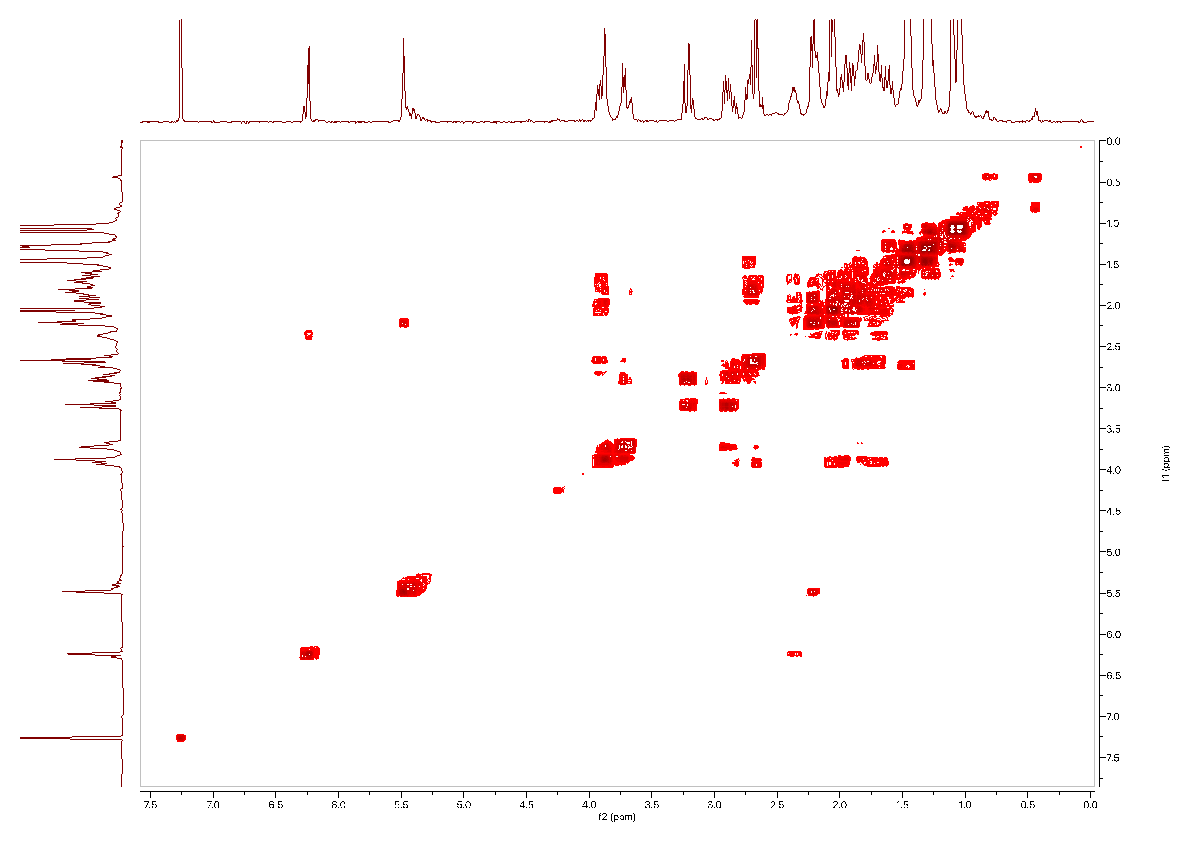

### Figure 24. COSY spectra for (9) (*C. sinensis*).

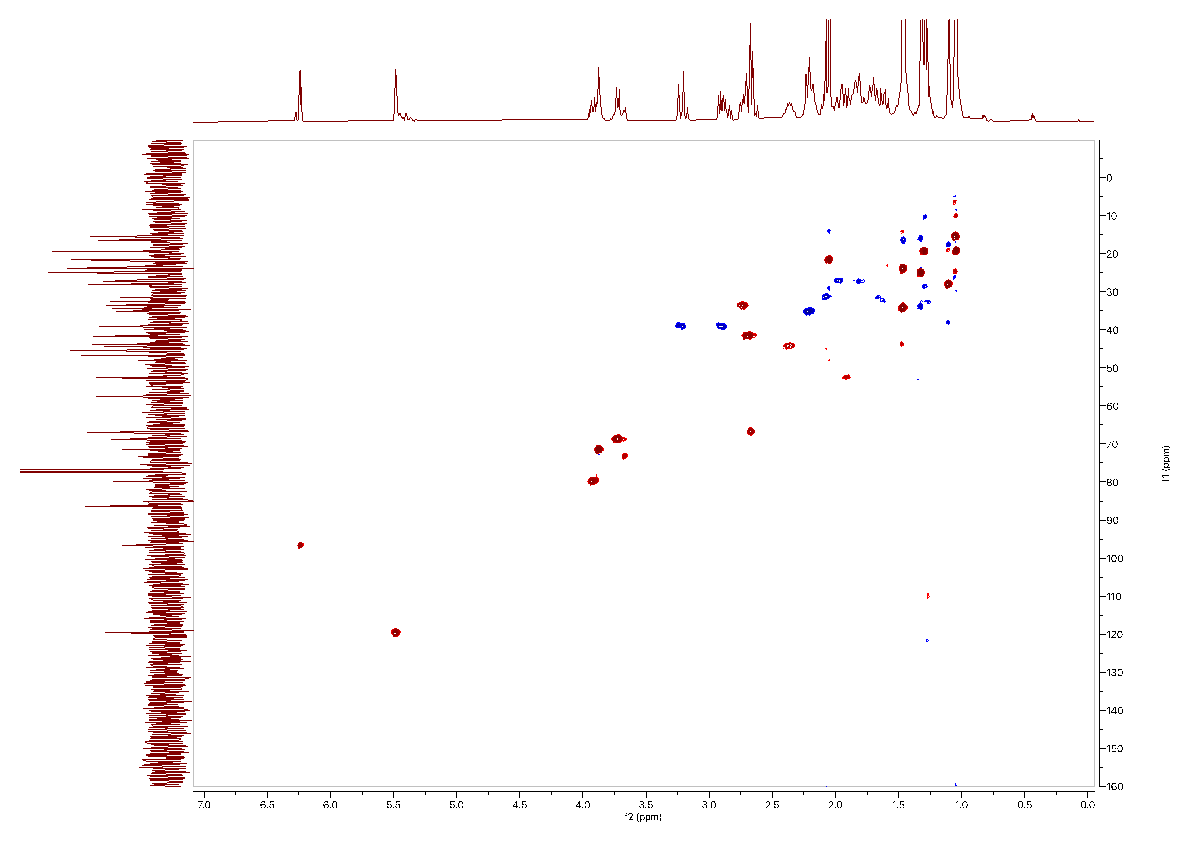

### Figure 25. HSQC spectra for (9) (*C. sinensis*).

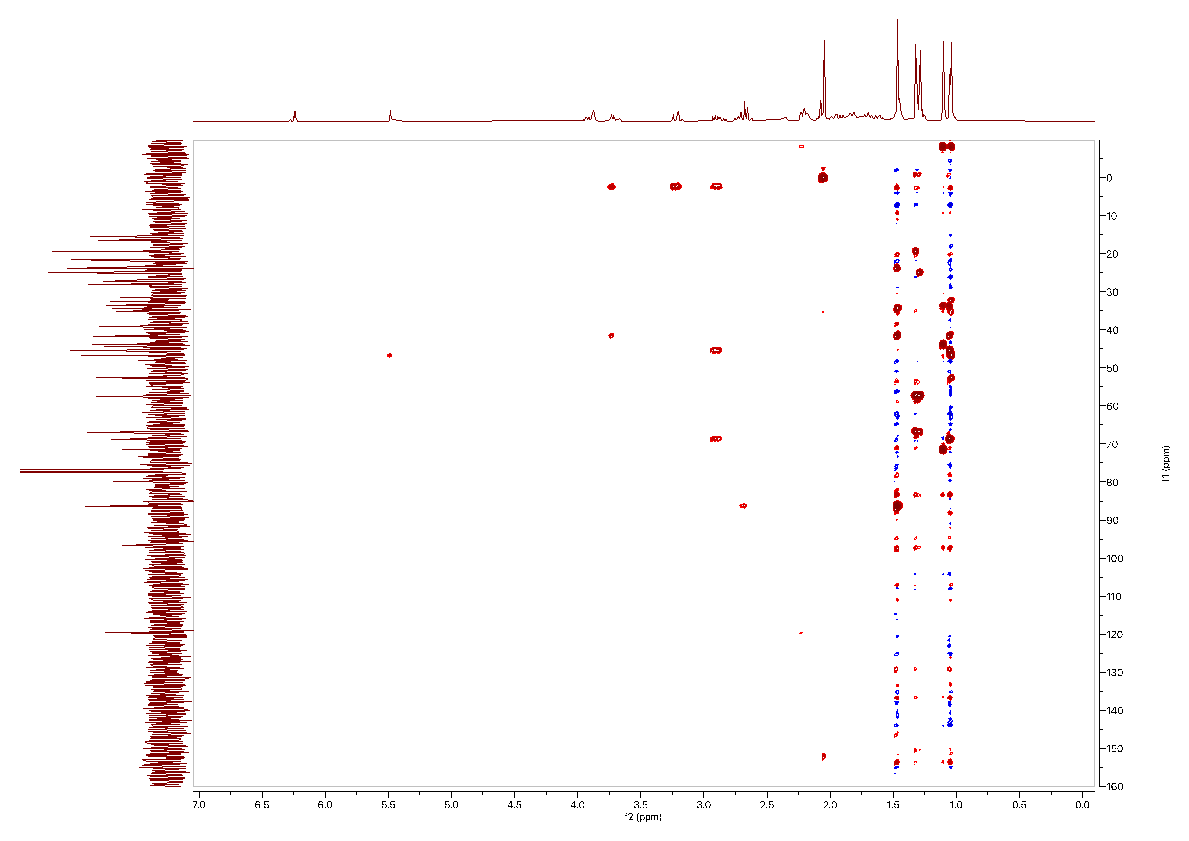

### Figure 26. HMBC spectra for (9) (*C. sinensis*).

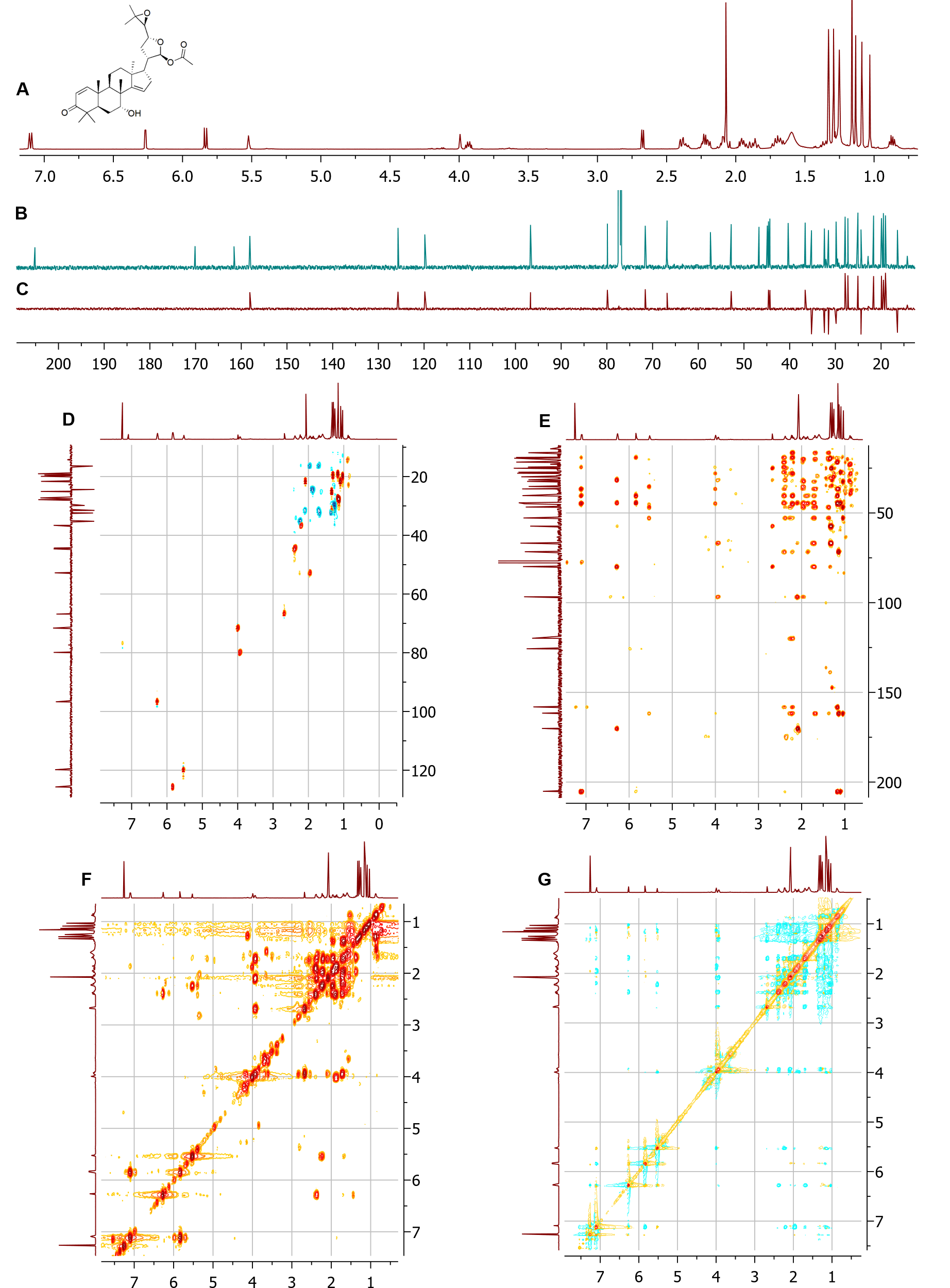

### Figure 27*.* NMR spectra of epi-neemfruitin B (10) *(M. azedarach).*

NMR spectra ([CDCl_3_], δ (ppm)). (A) ^1^H. (B) ^13^C. (C) DEPT-135. (D) DEPT-edited-HSQC. (E) HMBC. (F) COSY. (G) ROESY.

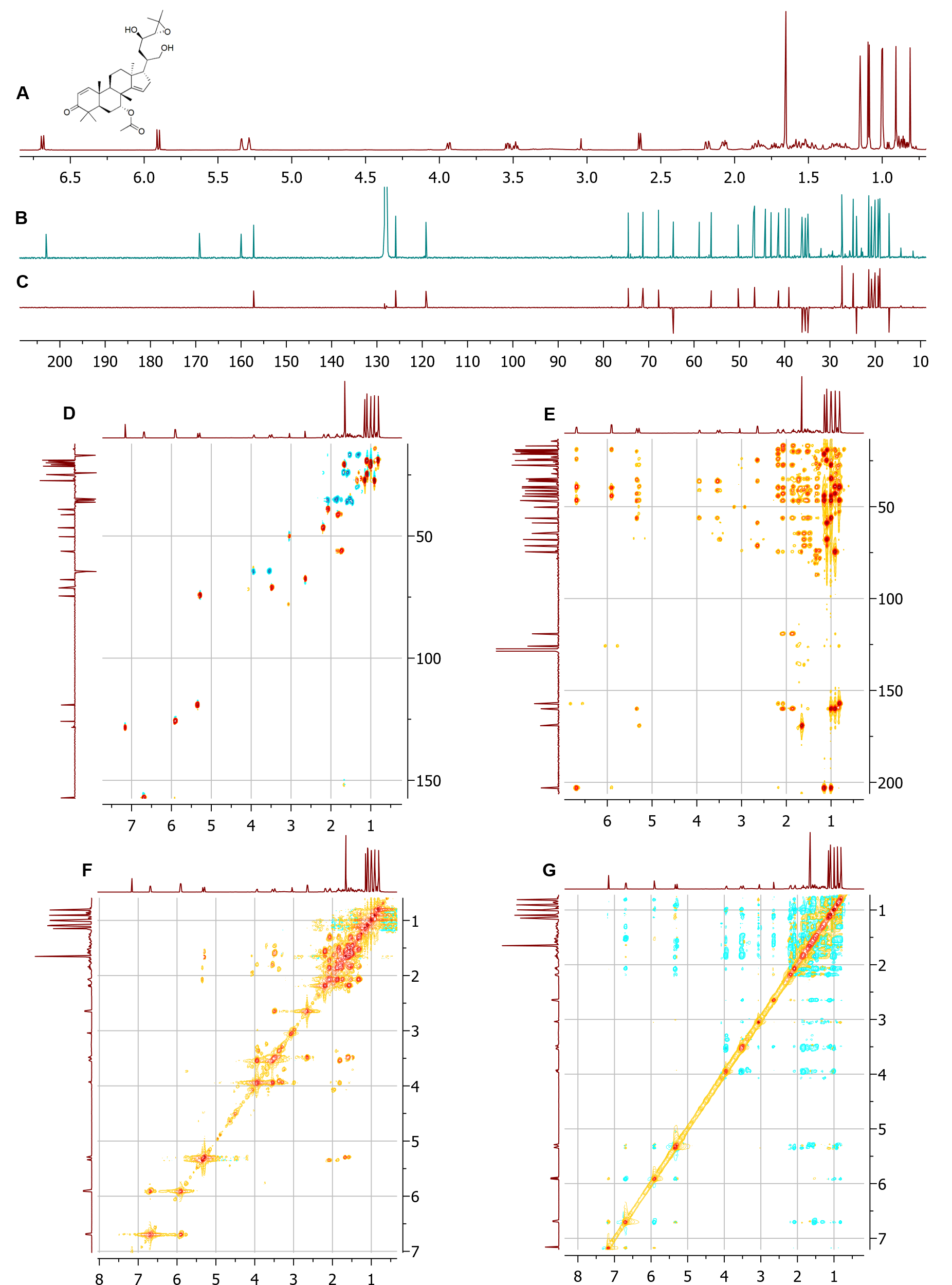

### Figure 28*.* NMR spectra of AKR Product (14) (M. azedarach).

NMR spectra ([Benzene-d_6_], δ (ppm)). (A) ^1^H. (B) ^13^C. (C) DEPT-135. (D) DEPT-edited-HSQC. (E) HMBC. (F) COSY. (G) ROESY.

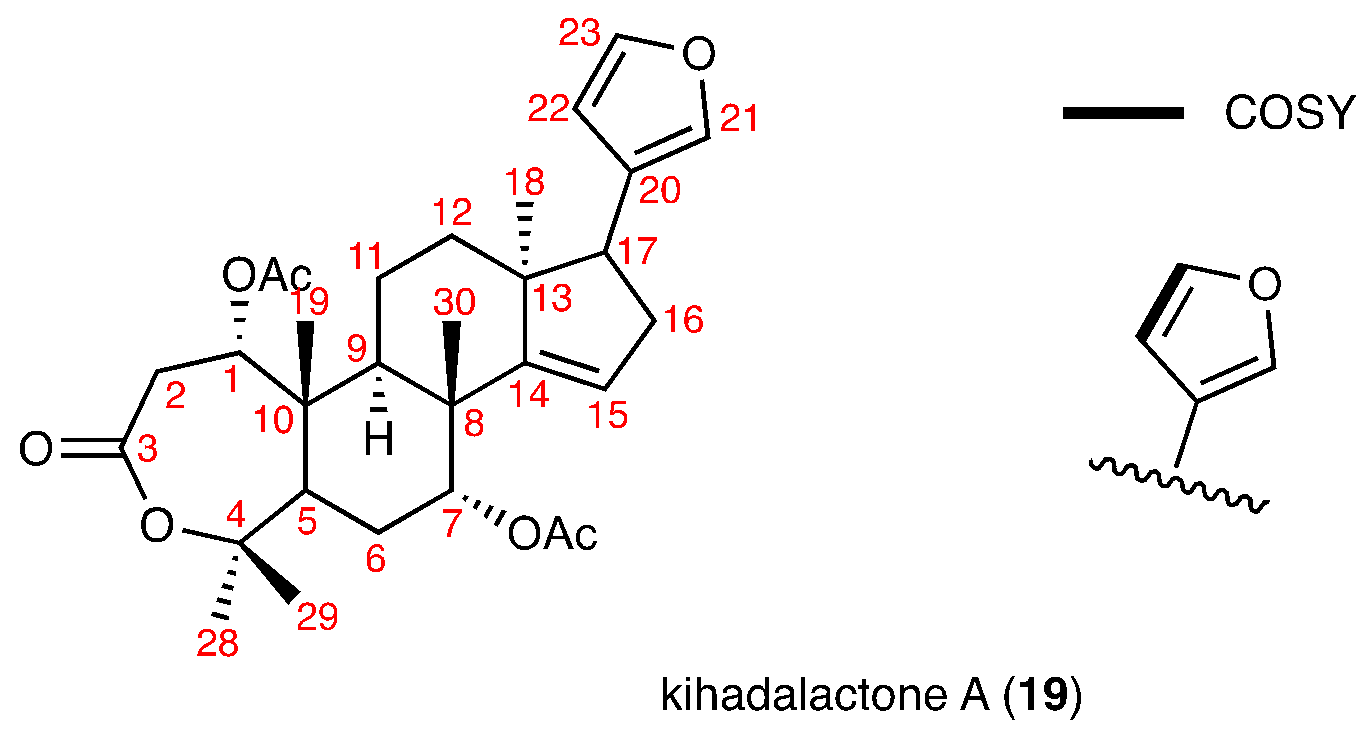

### Figure 29. Partial NMR assignment for kihadalactone A (19).

Assignment based on NMR spectra ([CDCl_3_], δ (ppm)) listed in Figure 30-31.

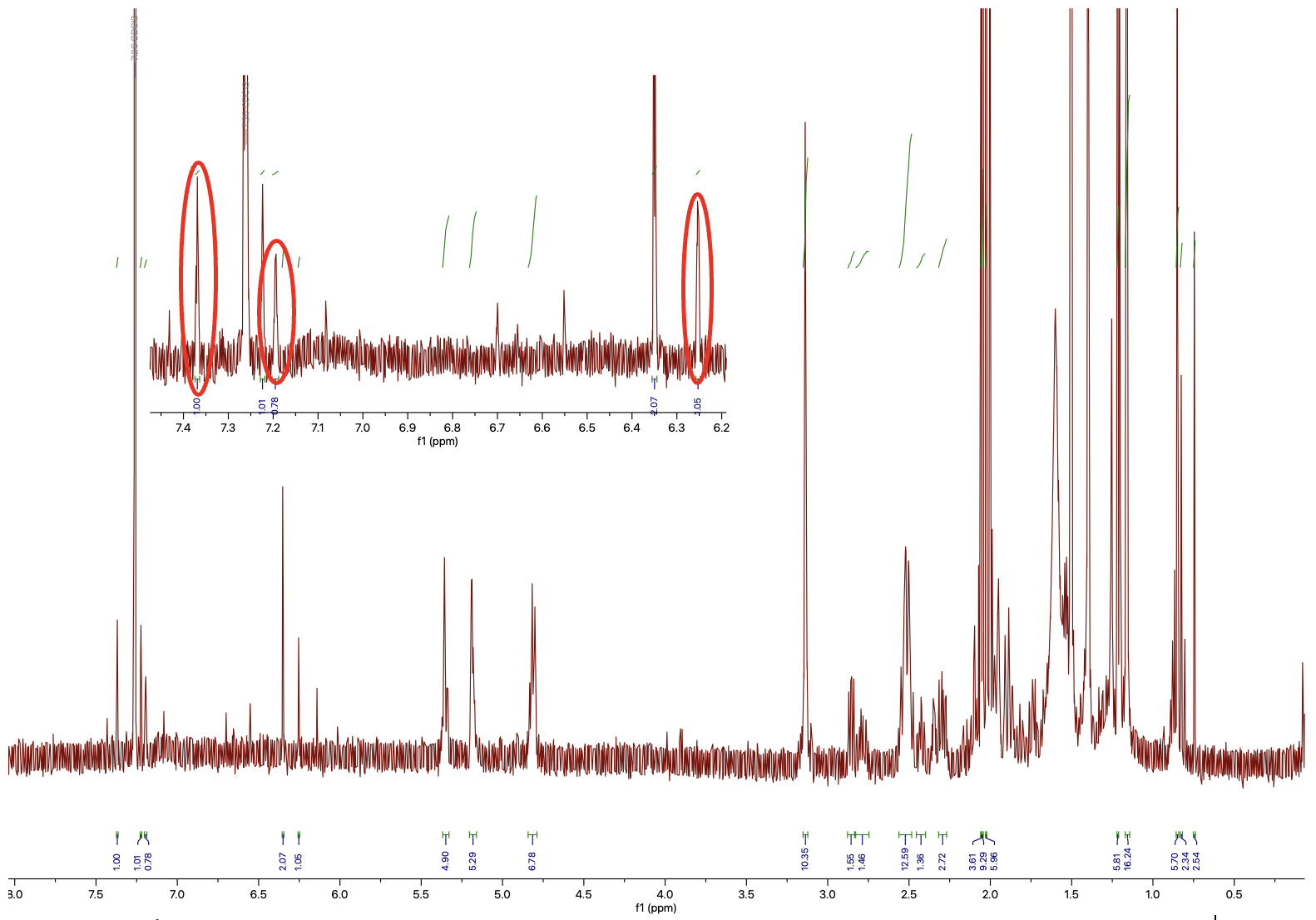

### **Figure 30. 1H spectra for kihadalactone A (19).**

The sample contained non-furan limonoid impurities such as (**17**). The signature furan protons of (**19**) are circled in red.

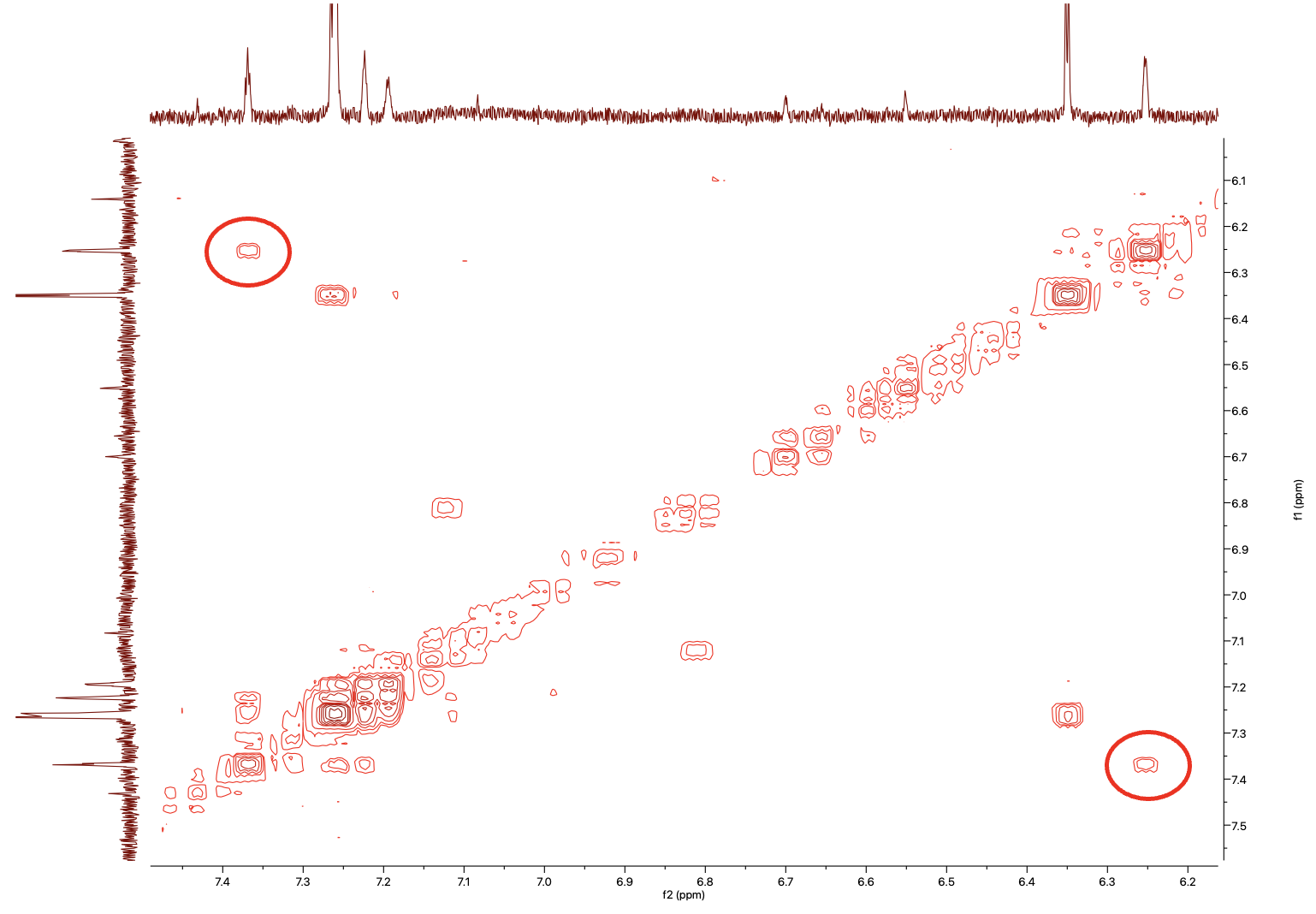

**Figure 31. COSY spectra for kihadalactone A (19).**

The sample contained non-furan limonoid impurities such as (**17**). The signature furan proton correlations of (**19**) are circled in red.

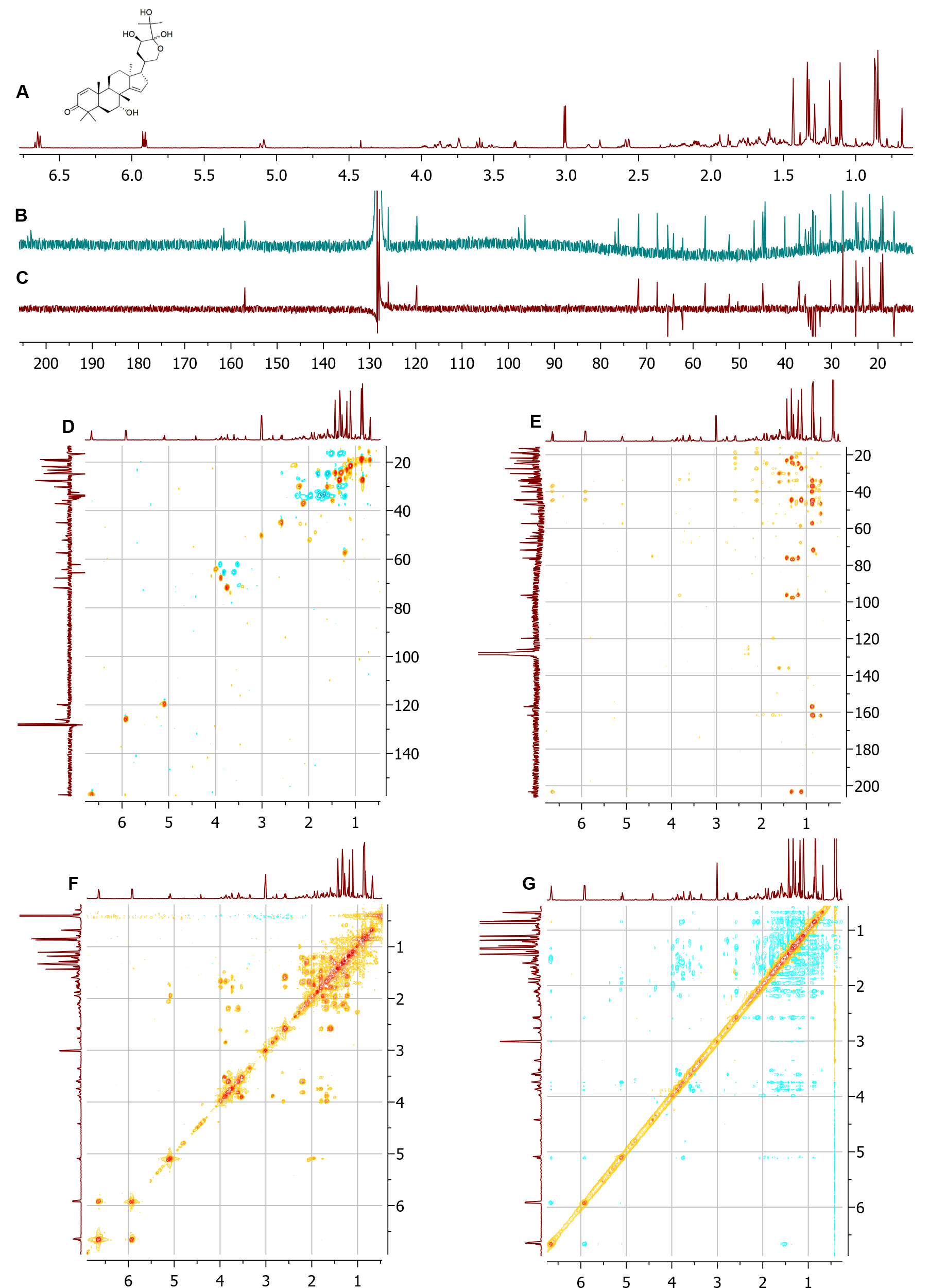

### Figure 32. NMR spectra of MaCYP716AD4 side-product (20) (*M. azedarach).*

NMR spectra of epimeric mixture ([Benzene-d_6_], δ (ppm)). (A) ^1^H. (B) ^13^C. (C) DEPT-135. (D) DEPT-edited-HSQC. (E) HMBC. (F) COSY. (G) ROESY.

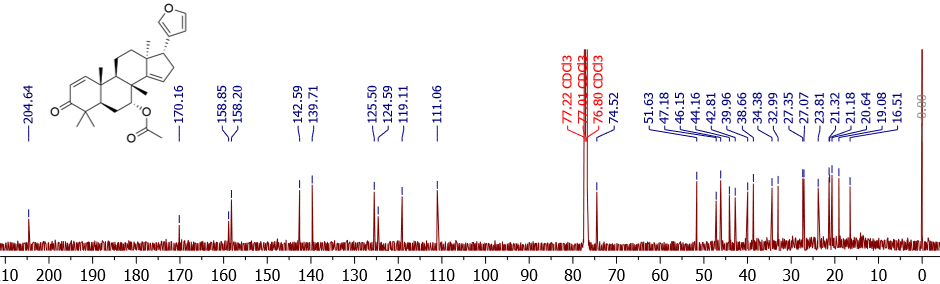

### Figure 33. ^13^C NMR spectra of analytical standard of azadirone (18).

NMR spectra [CDCl3], δ (ppm) of azadirone isolated from *A. indica*.
